# Time-calibrated phylogenomics reveals relationships and parallel evolution of neoteny within Elateriformia (Coleoptera) despite pervasive homoplasy

**DOI:** 10.64898/2026.09.23.753733

**Authors:** Dominik Kusy, Michal Motyka, Gabriel Biffi, Elizabeth T. Arias, Fedor Ciampor, Eric Anton, Christopher E. Carlton, Michael S. Caterino, Hermes E. Escalona, Jiří Hájek, Brittany Owens, Mikko Pentinsaari, Adam Ślipiński, Andreas Zwick, Ladislav Bocak

**Author notes:** Correspondence |.

## Abstract

The controversy over Elateriformia relationships hampers understanding of the evolution of disparate soft-bodied and fully sclerotized phenotypes, bioluminescence, and the ontogenetic reprogramming that leads to larviform females. Elateriformia contains >50,000 species in 34 families, including fireflies, jewel, and click beetles. We sample 31 families and 195 ingroup taxa, each with up to 4,224 orthologs. Through explicit hypothesis testing, coalescent-based approaches, and evaluation of dataset properties, we resolve backbone relationships. We reconfirm the monophyly of Byrrhoidea *sensu lato*, rejecting the recent resurrection of Dryopoidea, and verify the reciprocally monophyletic lampyroid and elaterid clades. We characterize short internodes and gene-tree discordance, both of which are indicative of ancient rapid radiations. Counterintuitively, signals from moderately variable genes are crucial for resolving certain splits, suggesting that even subtle rate differences distinguish phylogenetically informative loci from overly conserved, less informative markers. Time-calibrated phylogenies indicate the early origin of Elateriformia (281–287 Mya), with superfamilies diverging at the Permian-Triassic boundary and diversifying throughout the Jurassic. We identify 15 independent origins of soft-bodiedness and neotenic modifications spanning the Mid Jurassic to the present. Our hypothesis provides a foundation for investigating the genetic mechanisms underlying ontogenetic reprogramming and bioluminescence.

## Introduction

Elateriformia contains >50,000 ecologically and morphologically diverse species (Hunt et al., 2007) and surpasses several vertebrate classes in species diversity. Even though these beetles are less conspicuous in nature than most vertebrates, Elateriformia includes many taxa well known to the public, such as fireflies, jewel, click, and soldier beetles. Some are well-sclerotized, with compact bodies and, in some cases, with a unique clicking mechanism, while others are soft-bodied, with loosely connected body segments, or have incompletely metamorphosed females (Figure 1). Their morphological disparity is reflected in the definitions of 34 modern families (Figure 2E) (Hunt et al., 2007; Lawrence & Newton, 1995; McKenna et al., 2019). The current concept of the infraorder emerged only gradually from the utter chaos of 19^th^ and early 20^th^-century classification schemes (Crowson, 1955, 1981; Lawrence & Newton, 1982). Some families were excluded from Elateriformia *sensu* Lawrence & Newton (1995) by Hunt et al. (2007), who identified five families, four of which were initially placed in Elateriformia, as the earliest branches of Polyphaga (Lawrence & Newton, 1995). Kusy et al. (2018b) excluded Rhinorhipidae from Elateroidea and proposed Rhinorhipoidea as a sister to other Elateriformia, and McKenna et al. (2019) placed them even deeper as a lineage following the earliest five polyphagan lineages. Six elateriform families were recently erected (Bocak et al., 2016; Grebennikov et al., 2026; Jäch et al., 2016; Kusy et al., 2021; Lawrence et al., 2024; Rosa et al., 2020) and five were downranked to subfamilies or tribes within earlier established families (Bocak et al., 2018; Kundrata & Bocak, 2011; Kundrata et al., 2019; Kusy et al., 2018a) (Text S1).

**Figure 1.**
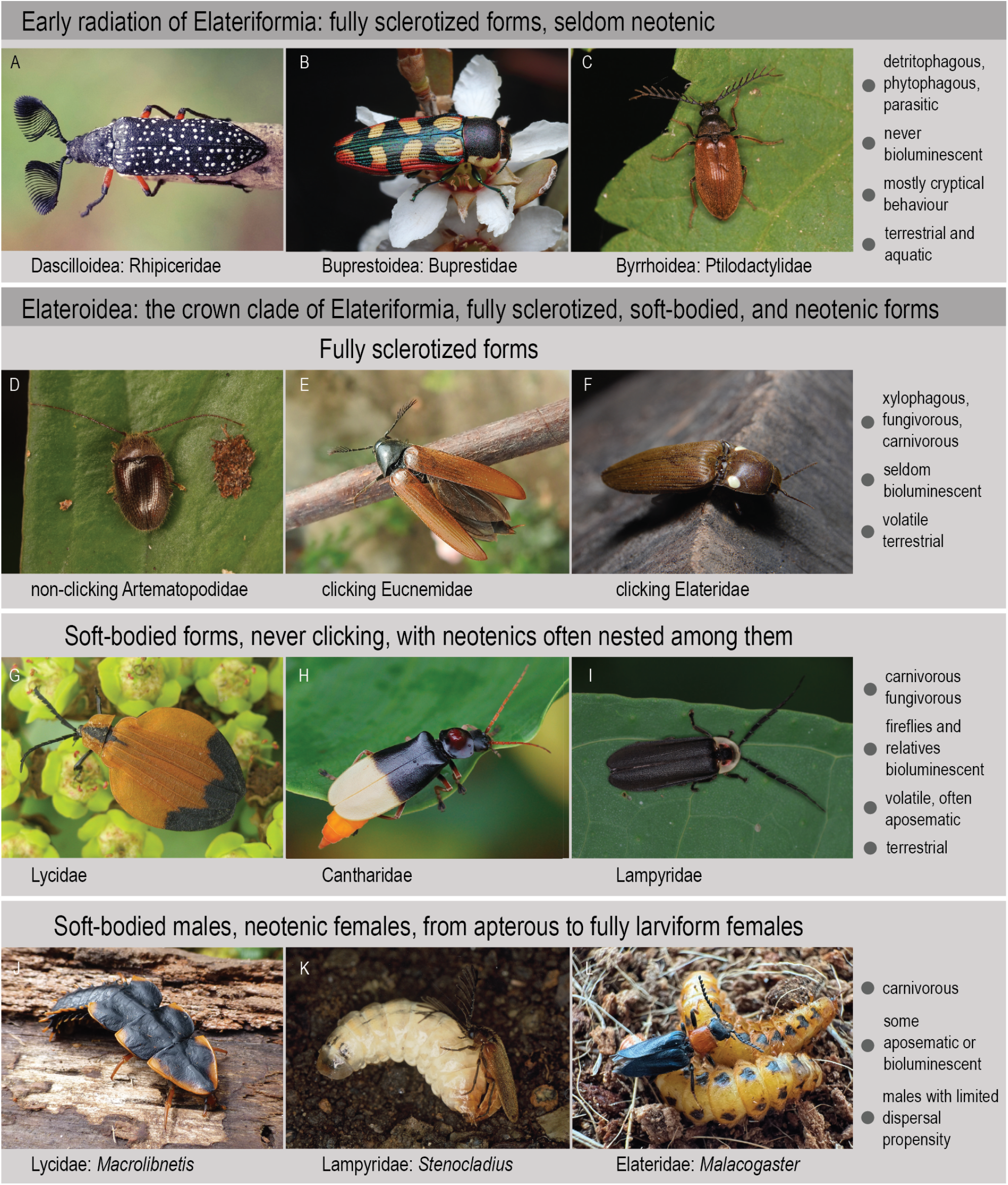
Selected representatives of Elateriformia. Characteristic traits and typical body plans are shown: A) male of *Rhipicera reichei*, Rhipiceridae (© Jennie Carruthers); B) *Castiarina pallidiventris*, Buprestidae (© Connor Margetts); C) *Anchycteis velutina*, Ptilodactylidae (© Chloe and Trevor Van Loon); D) *Artematopus* sp., Artematopodidae (© Thomas Oswald); E) *Phyllocerus flavipennis*, Eucnemidae (© Natalia Bulbulashvili); F) clicking and bioluminescent *Pyrophorus* sp., Elateridae (© Tomás Carranza Perales); G) *Lycus melanurus*, Lycidae (© T. Rulkens); H) female of *Chauliognathus* sp., Cantharidae; I) Lampyridae; J) mature female larva of *Macrolibnetis depressus*, Lycidae; K) mating female and male of *Stenocladius flavipennis*, Lampyridae (© Itsuro Kawashima); L) two females and a male of *Malacogaster* sp.; Elateridae.

**Figure 2.**
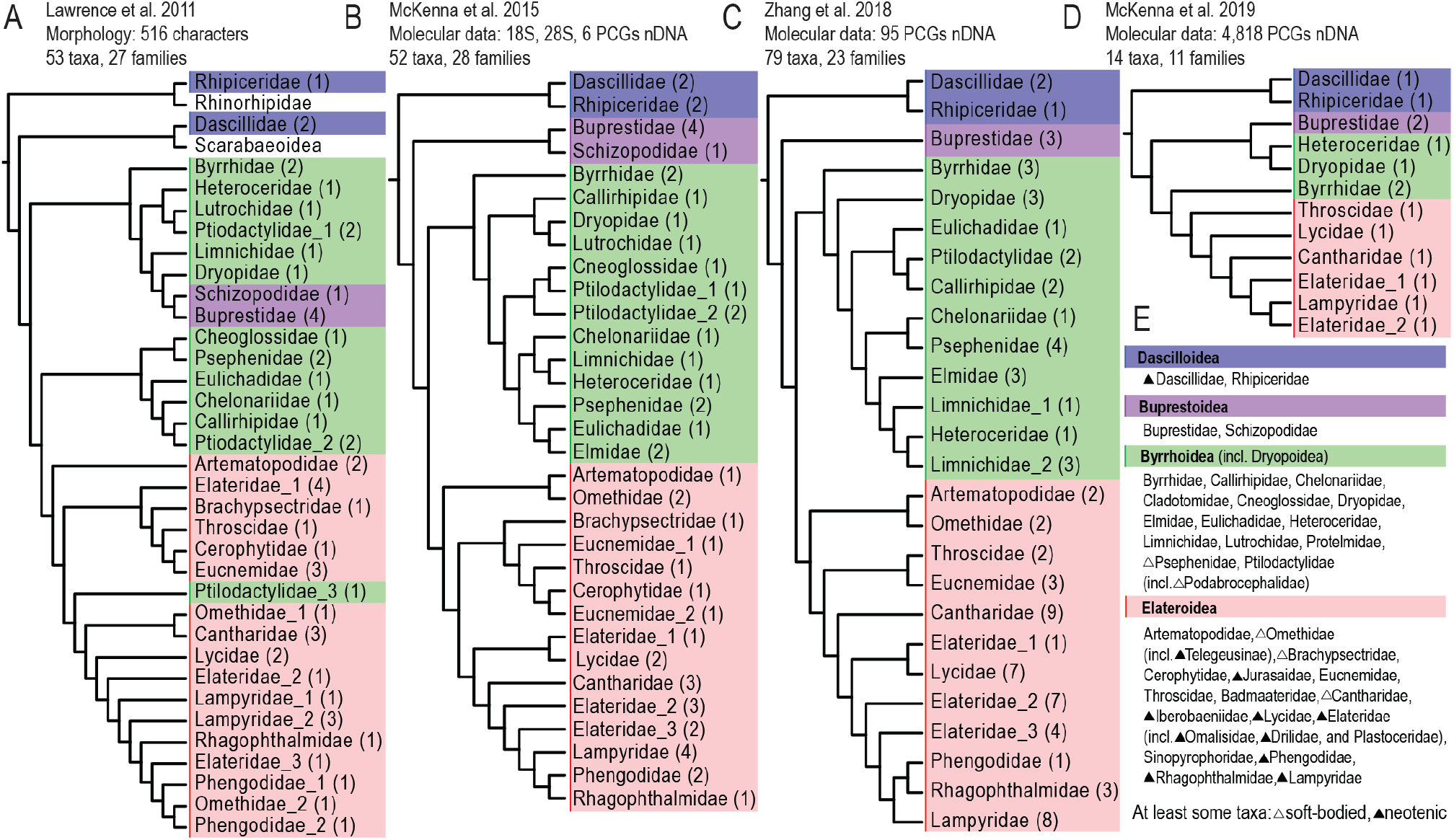
History of phylogenetic relationships among Elateriformia families from selected studies based on morphology and molecular data. Adapted to match currently recognized family classification. A) Lawrence et al., 2011 (Cladogram 3); B) McKenna et al., 2015 (Bayesian tree, Figs. 8–9; see also Maximum Likelihood tree); C) Zhang et al., 2018 (Fig. 2); D) McKenna et al., 2019 (Fig. 1); E) Currently recognized families. Only sections of respective trees are shown. Numbers in parentheses represent the number of species sampled in respective studies. PCGs, protein-coding genes; nDNA, nuclear DNA. See also Figure S1; Muona, 1995; Kundrata et al., 2014; Kusy et al., 2018a, b; Douglas et al., 2021; Hayashi et al., 2024 for further reference.

### Elateriform superfamilies

Four, sometimes five, constituent elateriform superfamilies are currently accepted (Figures 1, 2E) (Bocak et al., 2014; Zhang et al., 2018; Cai et al., 2022; Lawrence & Newton, 1995; Lawrence et al., 2011; McKenna et al., 2015, 2019). The concepts of Dascilloidea and Buprestoidea have not been questioned. Dascilloidea is an ancient superfamily, widely accepted as a sister to other elateriform groups, and contains Dascillidae and Rhipiceridae (Bocakova et al., 2007; Hunt et al., 2007; Lawrence & Newton, 1982, 1995). This group shares some morphological traits with Scarabaeiformia and Bostrichiformia (Crowson, 1971, 1978). Buprestoidea consists of two families, Schizopodidae and Buprestidae (Evans et al., 2015; Nelson & Bellamy, 1991). Morphological analyses sometimes hypothesize it as a sister of some byrrhoids (Lawrence, 1988; Lawrence et al., 2011), and similar relationships were proposed by some early molecular studies (Bocakova et al., 2007; Kundrata et al., 2014; Sagegami-Oba et al., 2007). The next branch in earlier topologies, Byrrhoidea, comprises the earlier-defined groups Byrrhoidea, Psephenoidea, and Dryopoidea (Crowson, 1978; Lawrence, 1988; Lawrence & Newton, 1982). Alternatively, Byrrhidae have recently been excluded from the Dryopoidea, forming a monotypic superfamily, and the Dryopoidea concept was resurrected (Cai et al., 2022; Hayashi et al., 2024). Analogously, the modern Elateroidea emerged from the fusion of Elateroidea, Cantharoidea, and Artematopodoidea (part) (Crowson, 1972, 1973; Lawrence, 1988; Lawrence & Newton, 1982, 1995). Cantharoidea was long considered monophyletic and placed either as a sister to Elateroidea (Crowson, 1972; Lawrence & Newton, 1982) or within Elateroidea as a cantharoid clade (Branham & Wenzel, 2003; Lawrence, 1988; Lawrence & Newton, 1995; Lawrence et al., 1995; Lawrence et al., 2011). Bocakova et al. (2007) and Sagegami-Oba et al. (2007) rejected the monophyly of cantharoids. However, even if most constituent families are clearly defined by synapomorphies, their relationships with one another remain ambiguous (Figures 1, 2, S1, S2).

All initial studies using limited Sanger data struggled with topological instability and low support for deep relationships (Bocakova et al., 2007; Hunt et al., 2007; Sagegami-Oba et al., 2007) (Figure 2, S1). Some trees suggested paraphyly, and sometimes polyphyly of Byrrhoidea and Elateroidea. Better-supported topologies became available when sampling was significantly expanded (Figure S1D, F) (Bocak et al., 2014; Kundrata et al., 2014). Complete mitogenomes and Sanger-produced nuclear protein-coding gene datasets were more informative and recovered the monophyly of all (or almost all) now-accepted superfamilies (Figure S1E, G, H) (Linard et al., 2018; McKenna et al., 2015; Timmermans et al., 2010, 2016; Zhang et al., 2018). However, support for relationships among superfamilies remained limited, and subsequent studies again called the early splits into question (Cai et al., 2022; McKenna et al., 2019; Hayashi et al., 2024).

A clear example of phylogenetic discordance is evident in studies of Byrrhoidea *sensu lato* (Figure S1G, I, K). McKenna et al. (McKenna et al., 2019), analysing ∼4,800 single-copy orthologs, found that Byrrhidae is sister to Elateroidea, with the remaining Byrrhoidea sister to Buprestidae. Zhang et al. (Zhang et al., 2018), analysing 95 nuclear coding genes, instead recovered Byrrhoidea as monophyletic and sister to Elateroidea in a relationship of (Buprestoidea, (Byrrhoidea, Elateroidea)). Cai et al. (Cai et al., 2022) later reanalysed Zhang et al.’s dataset (Zhang et al., 2018) and found a different arrangement: Byrrhidae as a sister to Buprestidae, with the remaining Byrrhoidea as a sister to Elateroidea. Following the latter results, Cai et al. (2022) resurrected Dryopoidea (Figure S1K), and this concept was adopted by Hayashi et al. (2024), although these authors showed a lack of clear support for the latest classification.

### Methodological challenges in resolving Elateriformia phylogenetics

The phylogenetic hypothesis is essential for subsequent investigation of the molecular mechanisms underlying diverse phenotypic traits (Jarvis et al., 2014; Chen et al., 2019; Song et al., 2020), including our main interest in the origins of soft-bodied and neotenic forms in Elateriformia. However, reconstructing some splits within the Elateriformia tree of life has always been challenging and mirrors broader issues in deep-time phylogenomics across other groups (Liu et al., 2019; Li et al., 2021c; Ballesteros et al., 2022; Parey et al., 2023; Schultz et al., 2023; Stiller et al., 2024; Chen & Zhang, 2025). First, reconstructing early splits in such an ancient group presents significant challenges due to the gradual erosion of phylogenetic signal over evolutionary time (Jeffroy et al., 2006; Sharma et al., 2014; Shen et al., 2017). Rapid radiations pose challenges for phylogenetic reconstruction, as rapid sequences of speciation events create short internal branches with limited accumulation of informative molecular changes (Irisarri & Meyer, 2016; Cloutier et al., 2019). Such scenarios may significantly affect Elateriformia, where some diversification events potentially coincided with the Early Jurassic biodiversity recovery (Montanaro et al., 2024; Sahney & Benton, 2008; Schoepfer et al., 2022). Furthermore, systematic biases such as compositional heterogeneity and long-branch attraction can complicate phylogenetic inference in groups with variable rates of molecular evolution (Ontano et al., 2021, 2022; Szánthó et al., 2023; Timmermans et al., 2016; Ballesteros & Sharma, 2019; Motyka et al., 2026b). Additionally, the issue is compounded by computational limitations, model inadequacies, and the inherent complexity of biological systems that can obscure true evolutionary relationships even with extensive datasets (Kapli et al., 2020, 2021; Boudinot et al., 2023a; Steenwyk et al., 2023; Vasilikopoulos et al., 2021).

We address these problems through several methodological decisions: data volume (thousands of single-copy orthologs, several million base pairs per sample), sampling density (near-complete sampling of families), sampling balance (proportional representation of major groups), dataset completeness, and advanced analytical approaches. We consider many factors because different analytical settings have produced conflicting topologies even when the same dataset was reanalysed, as shown by our earlier phylogenomic studies of Elateriformia (Kusy et al., 2021; Kusy, Motyka, Andujar, et al., 2018; Grebennikov, Kusy, Motyka, et al., 2026; Motyka, Kusy, et al., 2026) and whole-order phylogenies repeatedly inferred from the Zhang’s 95 protein coding gene dataset (Cai et al., 2022; McKenna et al., 2019; Zhang et al., 2018) (Figures 2C, S1J, K). Rather than relying on a single model and extensive data filtering, we focus on ambiguities and test alternative hypotheses (Boudinot et al., 2023a; Vasilikopoulos et al., 2021; Whelan & Halanych, 2017). We intend to compare multiple analyses and investigate topologies that conflict with other molecular studies and morphology-based delimitation of principal groups (*i*.*e*., Elateridae: Lissominae as a sister of Lycidae, the paraphyly of Byrrhoidea *s. l*., the Byrrhidae and Buprestoidea clade, lampyroids within Elateridae, etc.) (Figures 2A–D, S1A–L).

To fulfill the task, we significantly increase the sampling density and data volume compared to earlier Elateriformia-focused datasets (Bocak et al., 2016; Douglas et al., 2021; Kundrata et al., 2014; Kusy, Motyka, Bocek, et al., 2018) (Figure S1A–L). A high volume of data is needed to reliably address deep-time splits and assess the intrinsic structure of datasets (Kapli et al., 2020; Kjer et al., 2016; Young & Gillung, 2020; Rokas & Carroll, 2005; Koch, 2021).

High variability of substitution rates across lineages has been shown to negatively impact phylogenetic inference (Philippe et al., 2005). In Elateriformia specifically, slow molecular evolution of fully sclerotized elateroids and byrrhoids was apparent early in mtDNA- and rRNA-based studies (Bocak et al., 2014; Bocakova et al., 2007; Sagegami-Oba et al., 2007). In contrast, soft-bodied forms have much longer branches, and even higher evolutionary rates were observed in taxa with larviform females (Bocak et al., 2014; Bocak et al., 2008; Bocakova et al., 2007; Motyka et al., 2026b). Mitogenomic data also revealed heterogeneity in nucleotide composition as a source of systematic bias in Elateriformia (Timmermans et al., 2016). The authors applied various approaches to mitigate the tree’s instability. Subsequent studies aimed at expanding taxon sampling, but neither analysis produced a robust phylogenetic hypothesis (Bocak et al., 2014; Kundrata & Bocak, 2011; Kundrata et al., 2014) (Figure S1). A further dataset was compiled from a limited number of nuclear protein-coding markers and rRNA genes. Its analysis solved some problems, but suggested additional unexpected relationships (McKenna et al., 2015) (Figure S1E). Higher compatibility of proposed phylogenetic relationships with formal classification was recovered in analyses with expanded gene sampling (Zhang et al., 2018: 95 genes, 79 terminals; McKenna et al., 2019: 89 genes, 93 terminals), yet with some weakly supported and contradicting topologies such as the paraphyly of click beetles (Figure S1E, G). The latest reanalysis of Zhang et al.’s data recovered another inconsistent topology (Cai et al., 2022) (Figure S1K). Analyses of anchored hybrid enrichment (AHE) data were based on multiple genes. However, the analyses recovered lampyroid families, and sometimes even Lycidae, Cantharidae, Eucnemidae, Cerophytidae, and Throscidae, within Elateridae, in contrast to the accepted formal classification (Douglas et al., 2021). Similarly, Old-World families of Dryopoidea *sensu* Cai et al. (Cai et al., 2022) were analysed using ultraconserved elements (UCEs), and recovered Byrrhoidea *s. l*. as paraphyletic, but with low support and highly conflicting topologies (Hayashi et al., 2024).

The sequencing of transcriptomes and whole genomes produces datasets in which each sample was represented by several million base pairs. These datasets have played an increasing role in insect phylogenomics over the last decade (Misof et al., 2014) and have only recently entered beetle phylogenomics (McKenna et al., 2019) (Figures 2C, D, S1I). The recently produced datasets are orders of magnitude larger than those produced earlier, enabling more thorough analyses (Steenwyk et al., 2023). Phylogenomics using thousands of orthologs with variable rates of molecular evolution offers a new opportunity to resolve persisting ambiguities in Elateriformia. However, only eleven elateriform families were included in the whole-order beetle phylogeny, and results were partly incompatible with earlier hypotheses (McKenna et al., 2019) (Figure 2D). None of these studies focused on family-level relationships within Elateriformia, as they targeted Coleoptera as a whole, limiting sampling and precluding detailed evaluation of alternative topologies (Figures 2, S1G, J, L). Until now, no comprehensive phylogenomic dataset of Elateriformia has been assembled and thoroughly analysed with explicit consideration of analytical conflicts, and this study serves as the first step on this path.

### The timeframe of Elateriformia evolution

The timing of beetle diversification has long attracted research interest. Several Coleoptera-wide phylogenies have been time-calibrated, with successive analyses recovering progressively older ages for early divergences (Bocak et al., 2016; Cai et al., 2022; Hunt et al., 2007; McKenna et al., 2015, 2019; Toussaint et al., 2017; Zhang et al., 2018). The latest molecular studies strongly support Elateriformia as one of the oldest polyphagan lineages, originating close to the Permian extinction event (Cai et al., 2022; Kusy, Motyka, Andujar, et al., 2018; McKenna et al., 2019; Zhang et al., 2018; Beutel et al., 2024). Although the choice of calibration points was discussed in detail by all authors, even well-constrained calibrations cannot compensate for an uncertain phylogenetic framework resulting from limited sampling of Elateriformia and unexamined conflicting topologies. Alternative placements of jewel beetles, byrrhoids, and the lampyroid clade, as well as the paraphyly or polyphyly of click beetles (Cai et al., 2022; McKenna et al., 2019; Zhang et al., 2018), could substantially affect temporal scenarios regardless of the choice of calibration points.

The fossil record of Elateriformia compounds these challenges. Significant uncertainties include the Permian-Triassic origins of major polyphagan lineages (Beutel et al., 2024; Zheng et al., 2018; Montagna et al., 2024). Excluding fragmentary fossils, the Triassic fossil record of Elateriformia remains sparse, with *Elaterophanes vetustus* among the few described complete compression fossils (Handlirsch, 1906). Early fossils often exhibit typical elateriform body plans yet prove difficult to place phylogenetically. For example, Lasiosynidae were tentatively placed in Elateriformia based on elytral sculpture and thoracic interlocking mechanism (Kirejtshuk et al., 2010), but their phylogenetic position remains uncertain (Yan et al., 2014; Cai et al., 2022). Elateriform beetles are abundant in fossiliferous deposits since the Early Jurassic (Whalley, 1985; Dolin, 1973; Martin, 2010; Muona et al., 2020), and they flourished in the Mid to Late Jurassic. These deposits contain Dascillidae (Yan & Wang, 2010; Li et al., 2022), early byrrhoids (Yan et al., 2015), jewel beetles (Pan et al., 2011), click beetles (Dolin, 1980), Eucnemidae (Li, H., et al. 2023), Cerophytidae (Chang et al., 2011; Yu et al., 2019), and Artematopodidae (Cai et al., 2015; Oberprieler et al., 2016; Li et al., 2021a; Chang et al., 2009). To address these challenges, we employ an expanded set of well-constrained fossil calibrations based on a critical re-evaluation of the Elateriformia fossil record.

### Aims of the study

Indisputably, no Elateriformia phylogeny has achieved broad acceptance (Figures 2, S1), and several critical relationships require reinvestigation with proper representation of the group’s full diversity, including ontogenetically modified forms. Key outstanding questions include whether Byrrhoidea is monophyletic or paraphyletic, whether jewel beetles represent the second-earliest divergence within Elateriformia, are a sister to byrrhoids or part of them, the phylogenetic placement of enigmatic families such as Brachypsectridae, and whether lampyroids (*i*.*e*., Sinopyrophoridae, Lampyridae, Phengodidae, and Rhagophthalmidae) are sister to or nested within click beetles. Resolving these relationships is the principal goal of the present study. With a robustly supported backbone of Elateriformia, we can revisit earlier hypotheses proposing relationships of soft-bodied and neotenic elateroids. Although their monophyly was rejected by early molecular phylogenies (Bocakova et al., 2007; Evans et al., 2015; McKenna et al., 2019; McMahon & Hayward, 2016; Sagegami-Oba et al., 2007, but see Lawrence et al., 2011 for an alternative hypothesis), their detailed relationships remain hotly debated (Cai et al., 2022; Douglas et al., 2021; Kusy et al., 2021). Currently, the independent evolution of similar phenotypes is accepted, yet the number, timing, and phylogenetic distribution of these origins remain controversial, and some taxonomic changes have yet to gain wide acceptance. A robust, time-calibrated phylogenomic framework is therefore essential for identifying the independent origins of ontogenetic modifications, estimating their ages, and pinpointing their sister groups for future comparative genomic studies. The densely sampled phylogeny elucidates the origins of bioluminescence.

## Materials and Methods

### Sampling, DNA, and RNA extraction

Sampling covers almost the full family-level diversity of Elateriformia (31 of 34 extant families) and focuses specifically on taxa with modified body plans. Families Schizopodidae (7 spp.), Cladotomidae (20 spp.), and Badmaateridae (1 sp.) were unavailable or recognized after the dataset was assembled (Table S1, Figure S2). The sampling included 200 taxa, of which 195 belonged to the ingroup. In total, 125 samples were sequenced within the current Elateriformia project (60 elaterid samples were recently released by Motyka et al. (2026a) and are reused in this study; 65 samples are published for the first time: 55 samples were generated by the UPOL group, and 10 samples were provided by the CSIRO team). The remaining 75 samples were obtained from earlier studies (Kusy et al., 2018a, b, 2019; McKenna et al., 2019, and others; see Table S1, as available in January 2023). The outgroup was represented by beetle polyphagan taxa with reference genomes (Table S2).

The specimens were preserved in 96% ethanol, RNAlater, or snap-frozen in liquid nitrogen. Whole body or thoracic muscles were used for RNA and/or DNA extraction, depending on body size. Certain lineages, needed for the analyses due to their unsettled relationships or for representing unique morphological modifications, were unavailable in the recently collected material. Therefore, we included dry-mounted specimens in the sampling, allowing us to target additional taxa (Table S1). Rather than relying on genome reduction methods, we opted for low-coverage whole-genome sequencing (WGS) even for aged specimens. The dry-mounted specimens were first washed with 70% alcohol and briefly rehydrated with EDTA or 50% alcohol before processing for DNA isolation. When possible, only thoracic muscles were used to limit possible contamination from the body surface and digestive tract.

Genomic DNA from ethanol-preserved and dry-mounted specimens was extracted using Qiagen HMW DNA or DNeasy extraction kits; for dry specimens (both QIAGEN N.V., Venlo, The Netherlands), the lysis time was extended up to 24 hours. Total RNA was extracted from whole-body samples using TRIzol Reagent (Invitrogen, Thermo Fisher Scientific, Waltham, MA, USA) according to the manufacturer’s protocol. Concentrations of both DNA and RNA were evaluated using a Qubit fluorometer, Nanodrop ND-1000 spectrophotometer, and Agilent 2100 Bioanalyzer (Thermo Fisher Scientific, Waltham, MA, USA, and Agilent Technologies, Inc., Santa Clara, CA 95051, USA). The flowchart of sample and data processing is presented in Figure 3.

**Figure 3.**
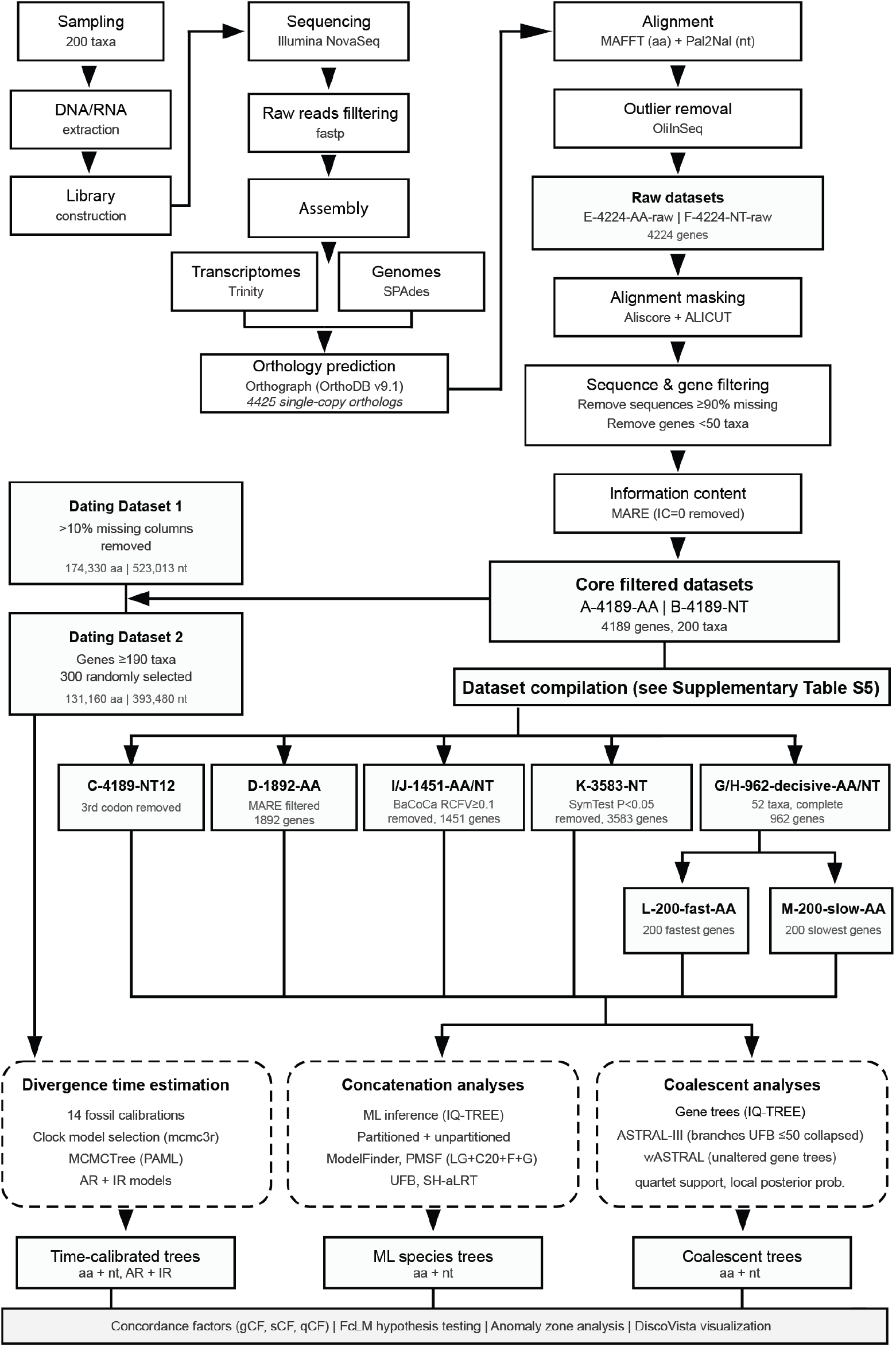
Flowchart of the phylogenomic data processing pipeline.

### Library construction, Sequencing, Data Processing, and Assembly

Sequencing was performed in multiple independent runs on Illumina flow cell lines during 2020–2023. All libraries for both stranded transcriptomes and whole genomes of UPOL samples were prepared by Novogene Co., Ltd. (Beijing, China) and sequenced on the NovaSeq platform (Illumina Inc, San Diego, USA) for 150 bp paired-end reads. Samples provided by the CSIRO team were sequenced by the Beijing Genomics Institute (Guangzhou, China) using the same technology and BGIseq 500, respectively.

#### Transcriptomes

Transcriptome libraries were sequenced to yield 6–8 Gbp of total data. The removal of low-quality reads and adaptor sequences was performed using fastp v.0.20.0 (Chen et al., 2018) with the following parameters: -q 5 -u 50 -l 50 -n 15. The quality was visualized with FastQC for both raw and filtered data (http://www.bioinformatics.babraham.ac.uk/projects/fastqc). All paired-end transcriptomic reads were assembled *de novo* using Trinity v.2.15 (Grabherr et al., 2011) with default settings and `--SS_lib_type RF` setting for strand-specific libraries or SOAPdenovo-Trans-31mer with default settings (Xie et al., 2014).

#### Whole Genomes

Genome libraries were shotgun-sequenced for 6–30 Gbp of total data, depending on genome size. Raw paired-end reads were filtered and quality checked as described above. The draft genomes were assembled using SPAdes v.3.15.3 (Bankevich et al., 2012), with k-mer sizes of 21, 33, 55, 77, 99, and 127. Resulting contigs were used to train ‘Augustus’ (Stanke & Waack, 2003) for species-specific gene models with BUSCO v.3 (Waterhouse et al., 2018). Predicted models were used for *ab initio* gene predictions with Augustus, and protein-coding sequences were used for subsequent analyses. The transcriptome and genome completeness were evaluated using BUSCO, targeting 2,442 Endopterygota single-copy orthologs. BUSCO quantifies completeness using evolutionary conserved expectations of gene content.

### Ortholog assignment

The single-copy ortholog set was collated by searching the OrthoDB v.9.1 database (Zdobnov et al., 2016) at the Coleoptera level with six Polyphaga reference taxa (Table S2). The search resulted in 4,425 single-copy orthologs. We carried out Orthograph v.0.6.3 searches (Petersen et al., 2017) on assembled transcriptomes and protein-coding gene sets, with the following settings: minimum-transcript-length = 30, frameshift-correction = 1, strict-search = 0, max-blast-hits = 100, max-blast-searches = 100, substitute-u-with = X. Data were summarized, and terminal stop codons were removed or masked using the Perl script summarize_orthograph_results.pl (Petersen et al., 2017).

### Multiple sequence alignment, outlier identification, and filtering

We aligned the amino acid (AA) sequences using Mafft v.7.407 with the L-INS-i algorithm (Katoh & Standley, 2013). Corresponding multiple sequence alignments of nucleotides (NT) were generated using Pal2Nal (Suyama et al., 2006). Outlier sequences were detected and masked using OliInSeq v0.9.5 (a part of the TEnriAn workflow; https://github.com/ZFMK/TEnriAn, https://github.com/cmayer/OliInSeq). The outlier search at the AA level used the following parameters: `--remove-gap-ambig-lowerCase-sites` `--windowsSize 20 --IQR-factor 2.0 -e 0.25`. We set searches for outlier sequences to a sliding window of size 20. Sequence segments identified as outliers were soft-masked with lowercase characters. When more than 25% of a sequence’s valid windows were identified as outliers, we removed the entire sequence from the corresponding gene alignment. OliInSeq automatically masks the outlier sequence segments in the corresponding NT alignments. Sequences excluded from the AA alignment were also removed from the NT alignments.

Moreover, we used Aliscore v.2.2 (Misof & Misof 2009; Kück et al., 2010) to identify ambiguous and randomly similar aligned sections in the AA alignments. Aliscore was invoked with a custom -r 10^27^ option to compare all sequence pairs within each sliding window of the default size, using a special scoring approach for gap-filled AA sites (option -e). Apart from these options, default parameters were used. After that, we used Alinuc.pl (Misof et al., 2014) to create a list of corresponding codons to be removed from NT alignments. Identified random or ambiguous similarities were masked using ALICUT v.2.3 (Kück et al., 2010). Additionally, to exclude short uninformative sequences in individual gene alignments, we removed sequences with ≥90% missing data that were calculated as a percentage of ‘-’ and ‘X’ in the AA alignments and ‘-’ and ‘N’ in the NT alignments using updated Python scripts TrimShortSeq.py (Zhang et al., 2020). To reduce the proportion of missing data, we removed individual gene alignments at both the AA and NT levels for which *<*25% (50) of taxa were present. We used MARE v.0.1.2-rc (Misof et al., 2013) to calculate the information content of each gene partition. Partitions with zero information content were removed, leaving 4,189 genes for analysis. Individual gene alignments were retained for coalescent analyses. We used AMAS (Borowiec, 2016) to calculate basic statistics for individual gene alignments (alignment length, GC content, % missing data, number of parsimony-informative sites, etc.). Additionally, using a custom Python script, we calculated basic statistics for unaligned data at the individual-taxon level, including the number of recovered single-copy orthologs, average length, N50, and the number of AA and NT positions for each species. These statistics were computed at three stages: raw data, outliers removed, and after Aliscore filtering.

### Dataset compilation

We designated diverse datasets to comprehensively test the sensitivity of our phylogenetic inferences to dataset properties and filtering approaches (Tables S3– 5). The concatenated datasets were generated using FasConCat-G v.1.4 (Kück & Longo, 2014). From the 4,189 genes, we generated the following datasets: A-4189-AA and B-4189-NT (designation: name-number of genes-amino acid or nucleotide level). To reduce the effect of possible saturation (Breinholt & Kawahara, 2013), we created dataset C-4189-NT12, excluding third-codon positions. To increase the information content of the dataset, we further reduced dataset A-4189-AA using MARE under default settings, and compiled dataset D-1892-AA-MARE. Moreover, to explore the possible effect of Aliscore alignment filtering, we used the original datasets after outlier removal (E-4224-AA-raw and F-4224-NT-raw). Next, to increase data decisiveness and reduce the computational burden for some analyses, we compiled additional datasets to cover all lineages of interest and address potential conflicts. For this purpose, we selected 52 taxa (one or two terminals per family, two outgroups, and 50 ingroups), choosing those with the most complete data (the highest number of genes and pairwise completeness). We kept only genes available for all 52 selected taxa. The dataset contained 962 genes (datasets G-962-AA-decisive and H-962-NT-decisive).

Additionally, to explore how topology could be affected by the rate of molecular evolution, we calculated positional conservation as the average pairwise identity across all informative alignment positions using a custom Python script. This column-wise approach maximized data utilization by treating gaps as missing data rather than excluding entire positions. Genes were ordered in reduced datasets from the slowest (i.e., most conserved) to the fastest evolving. We assembled two datasets containing the 200 fastest and 200 slowest genes at the AA level.

### Compositional heterogeneity tests and completeness evaluation

To explore the effect of among-species compositional heterogeneity and its possible effect on the tree reconstruction, we inspected the data at the AA level (dataset A-4189-AA) with BaCoCa v.1.105 (Kück and Struck, 2014) to identify the gene partitions that strongly deviated from compositional homogeneity using relative composition frequency variation value (RCFV). Following Vasilikopoulos et al. (2019), we considered compositional heterogeneity among species within a given partition to be high when RCFV ≥ 0.1. The heterogeneous partitions were excluded from the data to generate more compositionally homogeneous datasets (I-1451-AA-Bacoca). To enable direct comparisons, the same partitions were also removed at the NT level (J-1451-NT-Bacoca). Additionally, we used a matched-pairs test of symmetry (MPTS) (Naser-Khdour et al., 2019) implemented in IQ-TREE to identify the partitions that strongly deviated from compositional homogeneity directly at the NT level (p-value cutoff *<*0.05, `--symtest-pval 0.05`), and partitions below the threshold were excluded (`--symtest-remove-bad`). This step generated the dataset K-3583-NT-SymTest.

SymTest v.2.0.49 (https://github.com/ottmi/symtest) was used to calculate the overall deviation from stationarity, reversibility, and homogeneity (SRH) (Ababneh et al., 2006) for the AA and NT datasets. Heatmaps were generated to visualize the pairwise deviations from SRH conditions. The degree of missing data and overall pairwise completeness scores across all datasets were inspected using AliStat v.1.7 (https://github.com/thomaskf/AliStat), and heatmaps were generated. We used MARE to calculate matrix saturation and information content of each AA dataset. For a complete list of datasets and statistics, see Table S5.

### Phylogenetic analyses

Phylogenetic reconstructions were performed using the maximum likelihood (ML) criterion with IQ-TREE v.1.6.2 and v.2.4.0 (Minh et al., 2020a). First, we analysed all datasets using the initial gene partition boundary. The gene model selections were performed with ModelFinder (Chernomor et al., 2016; Kalyaanamoorthy et al., 2017) implemented in IQ-TREE (-MFP option). The GTR model was considered for NT supermatrices (additionally, all models were considered for the dataset B-4189-NT). For the AA supermatrices, the substitution models LG, DCMUT, JTT, JTTDCMUT, DAYHOFF, WAG, and free rate models LG4X and LG4M were tested. All possible combinations of modelling rate heterogeneity among sites were allowed (options: -mrate E,I,G,I + G,R -gmedian -merit BIC). We used the edge-linked partitioned model for tree reconstructions (-spp option), allowing each gene to have its own rate. Additionally, we performed unpartitioned analyses of all datasets, considering the same set of models for the model selection step as above. The optimized partition scheme was inferred in IQ-TREE v.2.4.0 for the datasets I-1451-AA-Bacoca and J-1451-NT-Bacoca using the - merge option and considering the same substitution models as above. The fast-relaxed clustering algorithm (Lanfear et al., 2017) was used. The top 10% of partition schemes `--rclusterf 10` and maximum partition pairs `--rcluster-max` were set to default (10 times the number of partitions). Moreover, for the dataset I-1451-AA-Bacoca, we used a heterogeneous posterior mean site frequency (PMSF) model with LG+C20+F+G as a rapid approximation to the time and memory-consuming profile mixture models, *e*.*g*., C10 to C120 (Wang et al., 2018), a variant of PhyloBayes’ CAT model. We used the tree from the unpartitioned ML analysis of the same dataset as the guide tree to calculate mixture model parameters and infer the site-specific frequency profile. Ultrafast bootstrap (UFB) (Hoang et al., 2018) and SH-like approximate likelihood ratio test (SH-aLRT) were calculated in IQ-TREE v.2.4.0 (options -bb 3000,-bnni, and -alrt 3000) to assess nodal supports. Additionally, we conducted five independent partitioned tree searches from random seeds for datasets A-4189-AA and B-4189-NT to avoid getting trapped in local optima.

### Coalescent and discordance analyses

To account for variation among gene trees owing to incomplete lineage sorting and to account for potential gene tree heterogeneity and discordance, the datasets A-4189-AA, B-4189-NT, E-4224-AA-raw, F-4224-NT-raw, decisive datasets F-962-AA-52, G-962-NT-52, and two rate-based subsets containing the 200 fastest-evolving and 200 slowest-evolving genes (derived from the G-962-AA-52 dataset) were also analysed using the coalescent-based species-tree method. For each gene partition, we calculated an ML gene tree in IQ-TREE with 3,000 ultrafast bootstrap (UFB) replicates (-bb option), implementing either the substitution models previously predicted by ModelFinder for the full partitioned analyses or models newly estimated for the reduced decisive dataset. For each gene tree, we calculated the average UFB and the average branch lengths.

For subsequent coalescent species tree estimation, we used the Accurate Species Tree Algorithm (ASTRAL-III MP v.5.15.5) (Sayyari & Mirarab, 2016; Zhang et al., 2018). To account for very poorly resolved branches on gene trees, we collapsed branches with ultrafast bootstrap values ≤50 using newick utilities v.1.6 (Junier & Zdobnov 2010) before running each ASTRAL analysis. Local posterior probabilities (lpp) (Sayyari & Mirarab 2016) and quartet frequencies of the internal branches (q1, q2, and q3) in every ASTRAL tree were calculated using the parameters ‘-t = 8 and 2’. We plotted pie charts representing quartet scores for the given topology and two alternatives to the resulting ASTRAL species trees in R using plot_Astral_trees (https://github.com/sidonieB/scripts/blob/master/plot_Astral_trees.R).

Recent publications have shown that ASTRAL accuracy is reduced by poorly resolved or noisy gene trees (Zhang & Mirarab, 2022; Zhang et al., 2025). Therefore, we also used Weighted ASTRAL (wASTRAL), a part of ASTER (https://github.com/chaoszhang/ASTER/) (Zhang & Mirarab, 2022; Zhang et al., 2025), on the set of gene trees without collapsed branches, using the default hybrid weighting mode. This approach improved phylogenetic accuracy by assigning weights to gene tree quartets based on both branch support values and branch lengths, mitigating the impact of noisy gene trees while preserving valuable phylogenetic signal.

Furthermore, we used DiscoVista v.1.0 (Sayyari et al., 2018) to visualize gene-tree quartet frequencies of three topologies around focal internal branches of the inferred species trees: ASTRAL trees from datasets A-4189-AA and B-4189-NT, and wASTRAL trees from datasets E-4224-AA-raw and F-4224-NT-raw. Here, various clades were considered to visualize the confounding signal at the level of individual genes.

To further evaluate the extent of phylogenetic incongruence, we employed the ASTRAL (B-4189-NT) tree as a reference species tree (-te) in IQ-TREE v.2.4.0 (Minh et al., 2020a, 2020b) to calculate both gene concordance factors (gCF, `--gcf`) and site concordance factors (sCF, `--scf`) using the enhanced likelihood calculation method (`--scfl 100`) as described by Mo et al. (2023), following the procedure described at https://iqtree.github.io/doc/recipes/concordance-vector. Next, we calculated concordance vectors using site, gene, and quartet measures to quantify topological variation at contentious nodes, following the framework introduced by Lanfear & Hahn (2024) (https://github.com/roblanf/concordance_vectors).

To determine whether the observed phylogenetic conflicts at deep internal nodes in the resulting congruent AA trees could be attributed to ancestral rapid radiations within the hypothetical anomaly zone, we applied the analytical framework proposed by Linkem et al. (2016). For pairs of consecutive parent-child internodes in our wASTRAL-derived species trees, we assessed whether they satisfied the anomaly zone condition using the equation from Degnan & Rosenberg (2006): 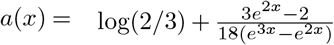, where *x* represents the length of the parent branch (in coalescent units) and *a(x)* defines the critical threshold. A pair of branches falls within the anomaly zone when the descendant branch length *y < a*(*x*), indicating conditions where gene tree discordance is expected even with error-free data. Calculations were performed using the script available at https://github.com/tkchafin/anomaly_zone.

### Alternative hypothesis testing

We used an explicit hypothesis-testing approach to interrogate contentious relationships along the backbone phylogeny. Using Four-cluster Likelihood Mapping (FcLM; Strimmer & von Haeseler, 1997) implemented in IQ-TREE v.2.4.0 (-lmap ALL, -lmclust, -wql), we evaluated alternative topologies for datasets A-4189-AA and B-4189-NT using the previously determined gene partition schemes and substitution models. This approach identifies incongruent or conflicting phylogenetic signals that may be obscured in concatenated ML analyses. We specifically tested several hypotheses at various hierarchical levels of relationships within Elateriformia, focusing on alternative placements of contentious clades revealed by our initial analyses. Detailed descriptions of all alternative topologies tested for each hypothesis are provided in Figure S32.

### Selection of fossil calibrations and divergence time estimation analysis

We re-evaluated fossils used in previous Coleoptera-wide studies (Toussaint et al. 2017; Cai et al. 2022; McKenna et al. 2015, 2019) and incorporated newly reported fossils (Table S6). We also reviewed Triassic-Jurassic fossil records that were not used as calibration points. In total, we employed 14 node calibrations primarily targeting deep phylogenetic splits (Figure S33; Table S6). Detailed justifications are provided in Text S2.

Divergence time analyses were conducted utilizing the ASTRAL topology derived from dataset B-4189-NT as the fixed reference tree. For the root calibration, we implemented a soft minimum constraint of 237 million years ago (Mya), derived from the unnamed Middle Triassic crown polyphagan fossil described by Zheng et al. (2018; fossil NIGP162053, dated to ∼238–237 Mya). The soft maximum constraint (293.69 Mya) is based on the oldest known beetle fossil *Coleopsis archaica* Kirejtshuk et al. (Kirejtshuk et al., 2014; Boudinot et al., 2023b; Schädel et al., 2022), and the age of the Sakmarian period (Schneider & Werneburg, 2006, 2012).

To reduce the computational burden, we applied two filtering methods. Method 1 used datasets A-4189-AA and B-4189-NT, which were filtered by removing all alignment columns with more than 10% missing data, yielding aligned datasets of 174,330 AA and 523,013 NT, respectively. For Method 2, using the same original datasets, we first excluded genes with fewer than 190 taxa (leaving 1,914 eligible genes), then we randomly selected 300 genes from this pool using a custom Python script (random_selection.py). This step produced reduced datasets containing 131,160 AA and 393,480 NT aligned positions, respectively.

Divergence time estimation was performed using the approximate likelihood calculation in MCMCTree implemented in PAML 4.10.7 (Yang, 2007), incorporating soft-bound minimum and maximum fossil calibrations on the nodes of the tree. Calibrated nodes were assigned uniform prior distributions bounded by minimum and maximum age constraints, with soft bounds allowing a 2.5% tail probability of ages falling outside these limits. The birth-death prior (BDparas, *λ* = *µ* = 1, *ρ* = 0.1), which approximates a uniform kernel, was used for uncalibrated nodes. To ensure our priors were appropriate, we ran MCMCTree without sequence data to verify that the effective priors were compatible with the specified priors. We constructed density plots for calibrated nodes comparing prior, effective prior, and posterior distributions. We used MCMCTree because of its computational efficiency when analysing large datasets, facilitating exploration of the impact of methodological variables. We first conducted Bayesian model selection using mcmc3r (https://github.com/dosreislab/mcmc3r) to determine the most appropriate clock model via marginal-likelihood calculation. MCMCTree implements three clock models: the geometric Brownian motion (GBM) model, the independent log-normal (ILN) model, and the strict clock (CLK) model (Rannala & Yang, 2007). Based on model selection results (Table S7), all analyses were run using both relaxed-clock models: GBM (autocorrelated rates, AR) and ILN (independent rates, IR). We aimed for 20,000 posterior samples with a sampling frequency of 1,000 and discarded 100,000 iterations as burn-in (2 *×* 10^7^ generations per chain). Other parameters were set as follows: cleandata = 0, BDparas = 1 1 0.1, kappa_-gamma = 6 2, alpha_gamma = 1 1, rgene_gamma = 2 40, sigma2_gamma = 1 10. As substitution models, we used HKY85+Γ5 for nucleotides and LG+Γ4 for amino acids, both with alpha = 0.5.

We used Tracer v1.7.2 to assess convergence, ensuring that effective sample sizes (ESS) exceeded the value 200 for all estimated parameters. We compared data from independent chains (either 2 or 4 chains, depending on the analysis) and plotted scatterplots of posterior mean divergence times and their 95% confidence intervals (CI) to assess: (1) consistency between individual chains within each run, and (2) consistency between separate runs. After confirming convergence, individual runs were merged with print = -1. We also created infinite-sites plots showing posterior mean times and 95% CI widths.

Ancestral-state reconstruction of morphological modifications leading to soft-bodiedness, incomplete metamorphosis, and fully larviform females was performed in BEAST v.1.10.4 (Suchard et al., 2018). The input topology was the dated tree obtained from the divergence-time analysis and was used as a fixed empirical chronogram. The character was coded as an ordered multistate trait with six states, ranging from 0, representing hard-bodied adults, to 5, representing the most strongly modified adult morphology. State reconstruction was estimated under a discrete continuous-time Markov model implemented in BEAST. Trees were logged using the standard BEAST tree-logging settings, and the initial burn-in was removed before summarizing posterior probabilities of ancestral states at internal nodes. We did not formally analyse the evolution of bioluminescence as the origin in the lampyroid clade, i.e., Sinopyrophoridae, Rhagophthalmidae, Phengodidae, and Lampyridae, was analyzed by Kusy et al. (2021), and the origins within Elateridae were analyzed by Motyka et al. (2023a). The latter study was based on a different dataset, with denser sampling, unavailable at the genomic level.

### Morphology

The investigation of the external morphology of elateriformian beetles was based on observations of the specimens used for sequencing and conspecific specimens collected in the field by the authors. Additionally, we examined material deposited in major natural history museums worldwide, including the Museum of Natural History, London, UK (C. O. Waterhouse collection); the Muséum national d’Histoire naturelle, Paris, France (M. Pic, J. Bourgeois, and H. S. Gorham collections); the Naturhistorisches Museum Basel, Switzerland (W. Wittmer collection); the Museum and Institute of Zoology, Polish Academy of Sciences, Warsaw, Poland (R. Kleine collection); the Museum of Comparative Zoology, Harvard University, Cambridge, Massachusetts, USA (J. L. LeConte collection); the American Museum of Natural History, New York City, USA (R. E. Blackwelder collection); and the Hokkaido University Museum, Sapporo, Japan (T. Nakane collection). We also examined material deposited in the senior author’s collection and specimens provided by colleagues, including R. Constantin, L. Dembický, P. Pacholátko, V. Kubáň, and the late V. Švihla, W. Büttiker, and W. Wittmer. The literature was surveyed for additional information on neotenic elateriform beetles and is referenced in the Discussion.

The specimens were collected worldwide between 1989 and 2025 under research permits for fieldwork held by the senior author or provided by colleagues. Ten specimens were sequenced as part of the Beetle Phylogenomics Project led by H. Escalona and A. Slipinski. Specimens collected in Europe, North America, and Japan originated from unprotected areas. No protected species were included in the analysis.

We examined the structure of the head, thorax, abdomen, and appendages, with particular attention to modifications of individual body parts. Examinations were performed using a stereomicroscope. Specimens were preserved in ethanol, dry-mounted, or macerated overnight in 10% KOH. When necessary, body parts were dissected for detailed examination. Some larviform females were unavailable and late instar large bodied larvae were examined (*Lyropaeus, Macrolibnetis*, and *Platerodrilus* species, Lycidae). The investigated specimens are deposited in the senior author’s collection. Standard taxonomic terminology was used to describe and compare the morphological characters of fully sclerotized species with those of species affected by incomplete sclerotization or retaining larval traits in sexually mature individuals.

## Results

The analysed dataset included 195 elateriform ingroup and five polyphagan outgroup taxa with an average of ∼3,800 orthologs per species. We recovered 2,642 (*Nyctophyxis ocellatus*) to 4,020 (*Drilaster* sp.) orthologs across the examined species. Protein sequence metrics showed N50 values ranging from 223 (*Cerophytum elateroides*) to 623 (*Agrilus zanthoxylumi* ), with total AA counts ranging from 575,941 (Elaterinae indet., G21027) to 1,988,355 (*Agrilus zanthoxylumi* ), and average sequence lengths from 189 (*Cerophytum elateroides*) to 499 AA (*Agrilus zanthoxylumi* ). Further details on the AA and NT datasets before and after outlier removal and Aliscore filtering are provided in Tables S3, S4, and Figures S3–S12. GC content varied considerably across Elateriformia, ranging from the highest values in Cerophytidae and Throscidae to the lowest in some Lycidae and Lampyridae, with variation observed across all three codon positions and the highest GC content at third codon positions, as expected (Figures S13, S14).

Our analyses used 13 phylogenomic datasets with varying properties (Table S5; Figures 4E, S3– 12). The primary datasets A-4189-AA and B-4189-NT contained 4,189 genes across 200 taxa, with completeness scores of 0.6615. Dataset A-4189-AA comprised 1,865,514 aligned AA positions and 1,133,846 parsimony-informative sites, while B-4189-NT included 5,596,542 NT aligned positions and 3,745,031 parsimony-informative sites. Raw unfiltered datasets E-4224-AA-raw and F-4224-NT-raw contained 3,383,300 AA and 10,149,900 NT aligned positions, respectively. Aliscore filtering reduced alignment lengths and missing data while maintaining similar numbers of parsimony informative sites and UFB support values between the filtered and raw datasets (Figure S12). Datasets addressing compositional heterogeneity, I-1451-AA-Bacoca, J-1451-NT-Bacoca, and K-3583-NT-

**Figure 4.**
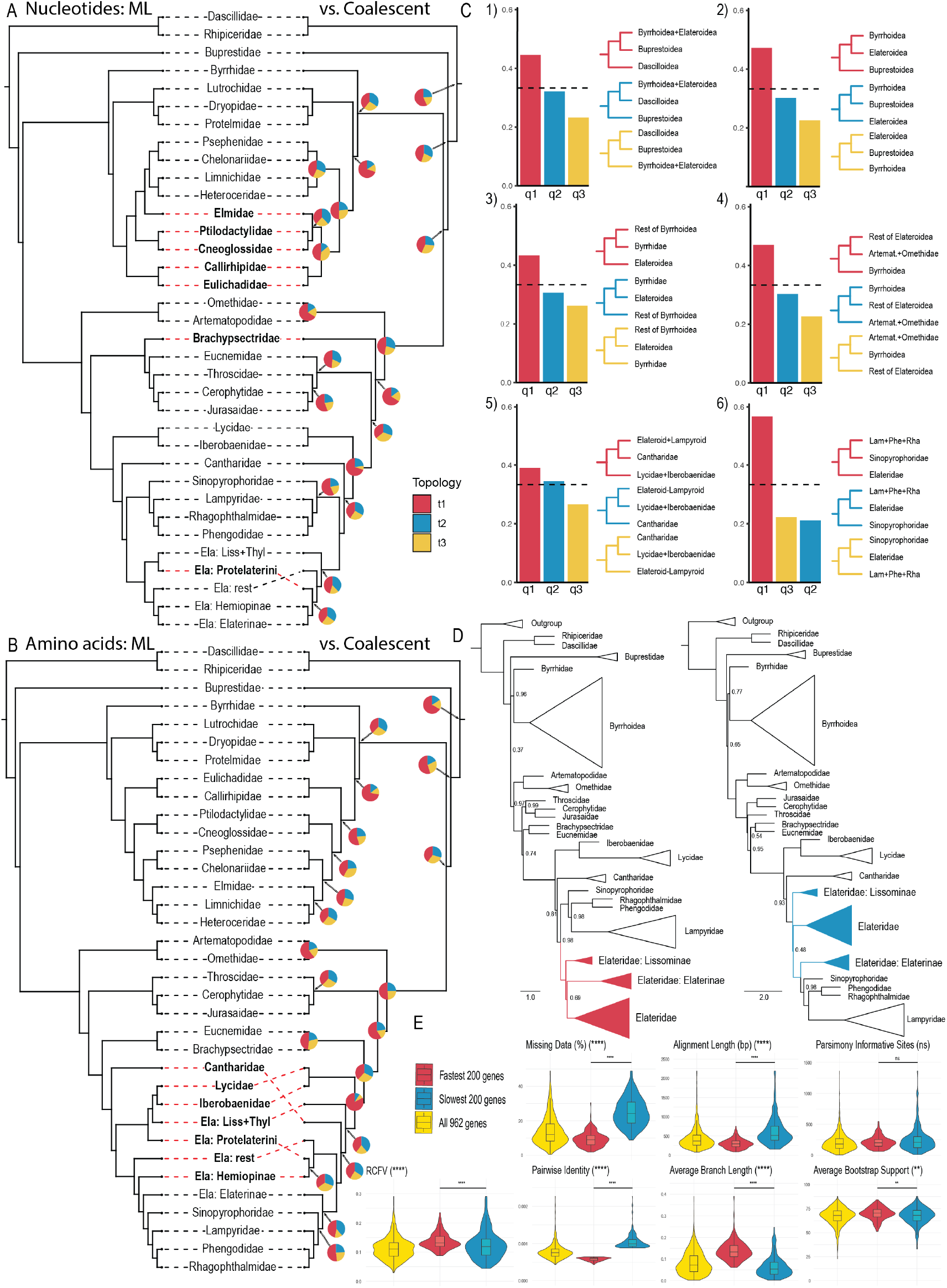
Results of phylogenomics analyses. Simplified tree comparisons: A) comparison between IQ-Tree maximum likelihood (ML) concatenated analysis partitioned by genes (left) and ASTRAL coalescent-based species tree methods (right) using NT sequences of dataset B-4189-NT and B) comparison between IQ-Tree ML concatenated analysis partitioned by genes (left) and ASTRAL coalescent-based species tree methods (right) using AA sequences of dataset A-4189-AA. Pie charts on nodes represent quartet concordance factors: red = q1 (proportion supporting the depicted topology), blue = q2 (proportion supporting the first alternative topology), yellow = q3 (proportion supporting the second alternative topology), see Figure S22 and S23 for full trees. C) DiscoVista visualization of quartet support for selected nodes across six different comparisons, showing the concordance vector (q1, q2, q3) for key clades, derived from individual gene trees of dataset B-4189-NT. D) ASTRAL coalescent species trees inferred from amino acid datasets partitioned by evolutionary rate: 200 fastest-evolving genes (left) and 200 slowest-evolving genes (right) derived from dataset G-962-AA-decisive. Blue and red shadings highlight the Elateridae clade. E) Violin plots comparing phylogenetic statistics between two evolutionary rate-based subsets containing the 200 fastest-evolving and 200 slowest-evolving genes (derived from the G-962-AA-decisive dataset).

SymTest, showed completeness scores of 0.651593– 0.652971. The 52-taxa datasets G-962-AA-decisive and H-962-NT-decisive had the highest completeness scores (both 0.83757). Subsets L-200-fast-AA-decisive and M-200-slow-AA-decisive contained 200 fastest- and slowest-evolving genes from G-962-AA-decisive, with 61,329 and 125,526 aligned AA positions, 42,397 and 51,144 parsimony-informative sites, and completeness scores of 0.902 and 0.747, respectively. MARE matrix saturation scores ranged from 0.833 to 0.935 across applicable AA datasets. Further details on all compiled datasets are provided in Table S5. An overview of the phylogenomic data processing pipeline is shown in Figure 3.

### Relationships within Elateriformia

A consistent backbone topology was recovered across most analyses (Figures 4, 5, S15–32), with approximately 95% of internal branches remaining congruent across data matrices and analytical approaches. Major topological conflicts were concentrated at nodes with short internal branch lengths (measured in coalescent units, mainly at the AA level; Figures 4A, B, S29–S32), indicative of ancient rapid radiations. Main conflicts are listed below, and the details are given in the section ‘Phylogenomic concordance and conflict analysis’.

**Figure 5.**
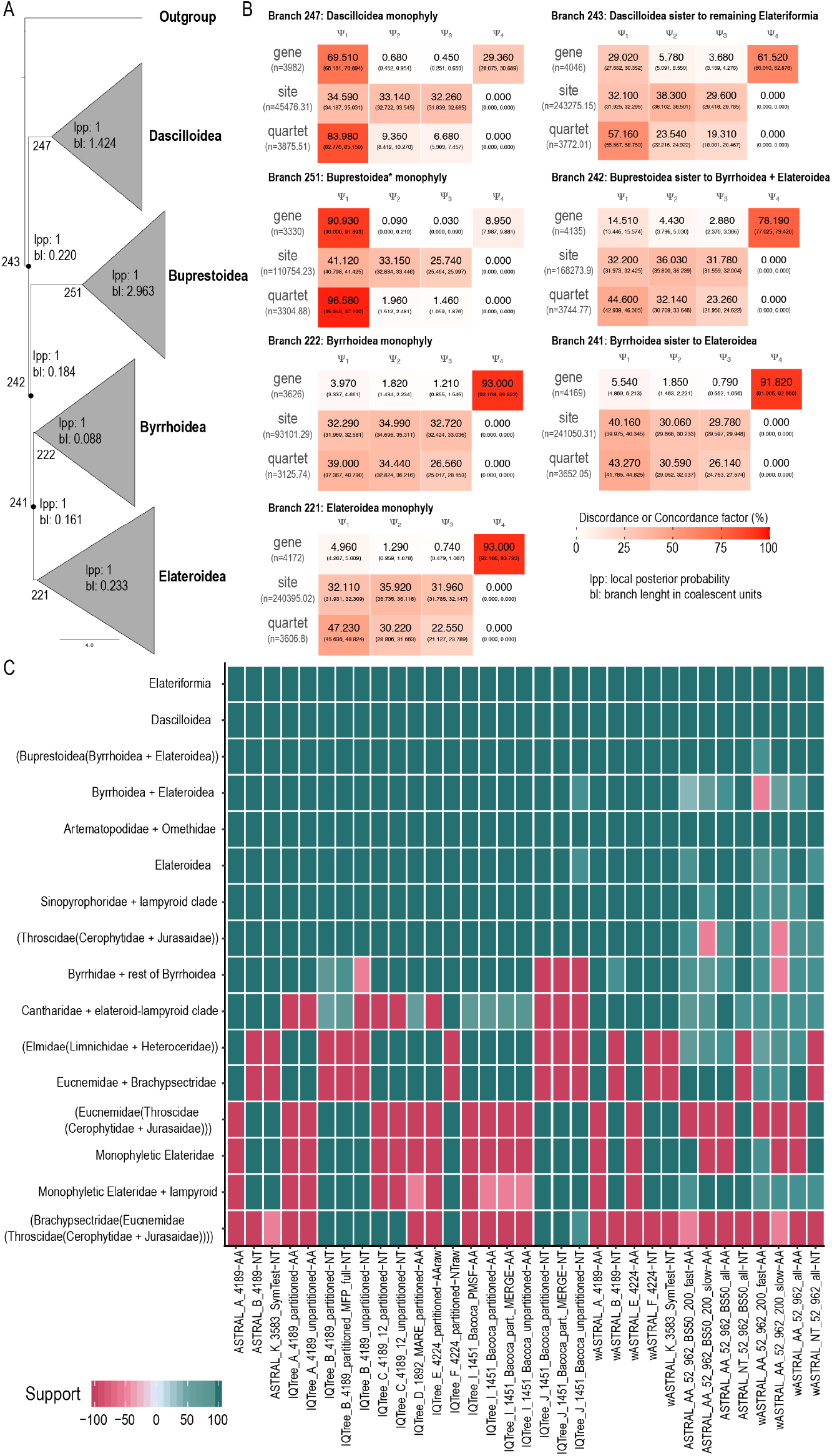
Concordance factors reveal extensive gene tree discordance across Elateriformia despite maximum statistical support in ML trees. A) Simplified ASTRAL phylogenetic relationships among four Elateriformia superfamilies based on Dataset B-4189-NT with local posterior probabilities (lpp) and branch lengths (bl) in coalescent units. All internal branches receive maximum statistical support (lpp = 1.00), but concordance vectors reveal dramatic variation in gene tree congruence. B) Heat maps show four concordance components: *ψ*_1_ (concordance factor: proportion of gene trees supporting the species tree topology), *ψ*_2_ and *ψ*_3_ (discordance factors: proportions of gene trees supporting the two alternative topologies obtained by nearest-neighbor interchanges), and *ψ*_4_ (proportion of gene trees matching all other discordant topologies not represented by *ψ*_2_ or *ψ*_3_ ), calculated from gene (gCF), site (sCF), and quartet (qCF) concordance factors. Values are percentages, with color intensity indicating magnitude. C) Heatmap showing support for select clades across 33 analyses. Analyses are grouped on the x-axis by method, dataset name (See Table S5) and parameters; clades are grouped on the y-axis. Color intensity indicates support values (pink = low support, teal = high support; see scale bar). Analysis abbreviations: ASTRAL = ASTRAL coalescent species tree analysis; wASTRAL = weighted ASTRAL; IQTree = IQ-TREE maximum likelihood analysis; AA = amino acid sequences; NT = nucleotide sequences; partitioned = partitioned by gene; unpartitioned = unpartitioned analysis; SymTest = partitions removed after SymTest analysis; MFP_full = all IQ-TREE models under -MFP were considered; Bacoca = partitions removed after Bacoca analysis; MERGE = optimized partition scheme search performed; PMSF = posterior mean site frequency model with LG+C20+F+G; BS50 = collapsed branches with ultrafast bootstrap values ≤ 50; 200_fast and 200_slow = 200 fastest-evolving and 200 slowest-evolving genes from the G-962-AA-decisive dataset.

All analyses consistently recovered Dascilloidea as the first-diverging lineage, sister to the remaining Elateriformia, followed by Buprestoidea as sister to Byrrhoidea + Elateroidea (Figures 4, 5A–C, S15–S32). The sole exception occurred in wASTRAL analyses of the 200 fastest-evolving genes from dataset G-962-AA-decisive, where Buprestidae was recovered as sister to a monophyletic Byrrhoidea (Figures 5C, S27C, D), with conflicting quartet support of q1: 36, q2: 33, and q3: 31, respectively.

Byrrhoidea *s. l*. was recovered as monophyletic in 28 of 33 analyses (Figures 4, 5A–C, S15–S32), i.e., with the clade of Byrrhidae and dryopoid families. Five analyses placed Byrrhidae as sister to Elateroidea: the ML analysis of dataset J-1451-NT-Bacoca (Figure S19), the unpartitioned ML analysis of dataset B-4189-NT (Figure S16B), and the coalescent wASTRAL analysis of 200 slowest-evolving genes from dataset G-962-AA-decisive (Figure S28E, F). However, these placements did not receive maximum support (aLRT/UFB: 99.9/100, 99.9/99, 99.9/97; 90.3/92) and quartet support (q1: 38, q2: 37, q3: 24), suggesting they may be artifacts. Within Byrrhoidea, Lutrochidae was consistently recovered as the sister group of Dryopidae + Protelmidae, representing the next divergence after Byrrhidae or the first split in the respective clade, in all 33 analyses. The subsequent split varied slightly: Eulichadidae + Callirhipidae formed the next diverging clade in most analyses, except in coalescent NT analyses, where they formed a monophylum with the clade (Elmidae, (Ptilodactylidae, Cneoglossidae)), with two distinct arrangements (Figures 4A, S22C–F, S23C–F, S24G, H, S25, S27, S28G, H), however always with a conflicting quartet support. Cneoglossidae was consistently recovered as the sister of Ptilodactylidae across all analyses. In all AA analyses, Ptilodactylidae + Cneoglossidae represented the next split, sister to a clade comprising ((Heteroceridae, Limnichidae), (Psephenidae, Chelonariidae)) (Figures 4A, B, S15–S32). The position of Elmidae varied across analyses: it was recovered as the sister of Limnichidae + Heteroceridae in ML and coalescent AA analyses, the sister of Ptilodactylidae + Cneoglossidae in ASTRAL coalescent NT analysis, the sister of Eulichadidae + Callirhipidae in wASTRAL coalescent NT analysis, or the sister of the entire ((Heteroceridae, Limnichidae), (Psephenidae, Chelonariidae)) clade in ML NT analysis (Figures 4A, B, 5, S15–S32). Monophyletic Elateroidea was recovered in all 33 analyses (Figures 4, 5C, S15–S32). Within Elateroidea, the clade comprising Artematopodidae and Omethidae, including Telegeusinae, was consistently recovered as the first-diverging lineage, i.e., the sister group to the remaining elateroids (Figures 4, 5C, S15–S32).

The position of Brachypsectridae showed discordance across analyses, with short internal branch lengths (Figures 4D, S22, S25). In ML NT analysis, Brachypsectridae was recovered as the sister to a clade (Eucnemidae, (Throscidae, (Cerophytidae, Jurasaidae))) (Figures 4A, S15B, C, 16B; S18B, S20, S21B) or the sister to Eucnemidae, forming a monophylum with (Throscidae, (Cerophytidae, Jurasaidae)) (Figure S17). In AA analyses, both ML and coalescent, a different branching order was recovered: (Throscidae, (Cerophytidae, Jurasaidae)) diverged first, followed by Eucnemidae + Brachypsectridae as the sister of the remaining families (Figures S15A, 16A, 18A, S19, S21A, S22– S28). Coalescent NT analyses showed a third topology, with Brachypsectridae as the sister group of all remaining families (excluding Artematopodidae and Omethidae) (Figures 4A, S22–S28). The clade (Eucnemidae, (Throscidae, (Cerophytidae, Jurasaidae))) was recovered across all NT ML and coalescent analyses (Figures 4A, S15–S28). In AA ML and coalescent analyses, (Throscidae, (Cerophytidae, Jurasaidae) diverged first, followed by Eucnemidae, but see the quartet supports at respective nodes (Figures 4B, S22–S29).

The crown Elateroidea always formed a clade containing Lycidae, Iberobaeniidae, Cantharidae, Elateridae, and the lampyroid branch represented by Sinopyrophoridae, Phengodidae, Rhagophthalmidae, and Lampyridae (Figures 4, 6, S15–S31). Within the clade, the relationships among Lycidae + Iberobaeniidae, Cantharidae, and the remaining families showed discordance between NT and AA analyses. In NT analyses, both ML and coalescent analyses, the branching order was consistent: Lycidae + Iberobaeniidae diverging first, followed by Cantharidae as the next split, and the terminal clade containing monophyletic Elateridae and the lampyroid clade (Figures 4, S15–S28). The sole exception to this pattern was the ML analysis of dataset C-4189-NT12.

**Figure 6.**
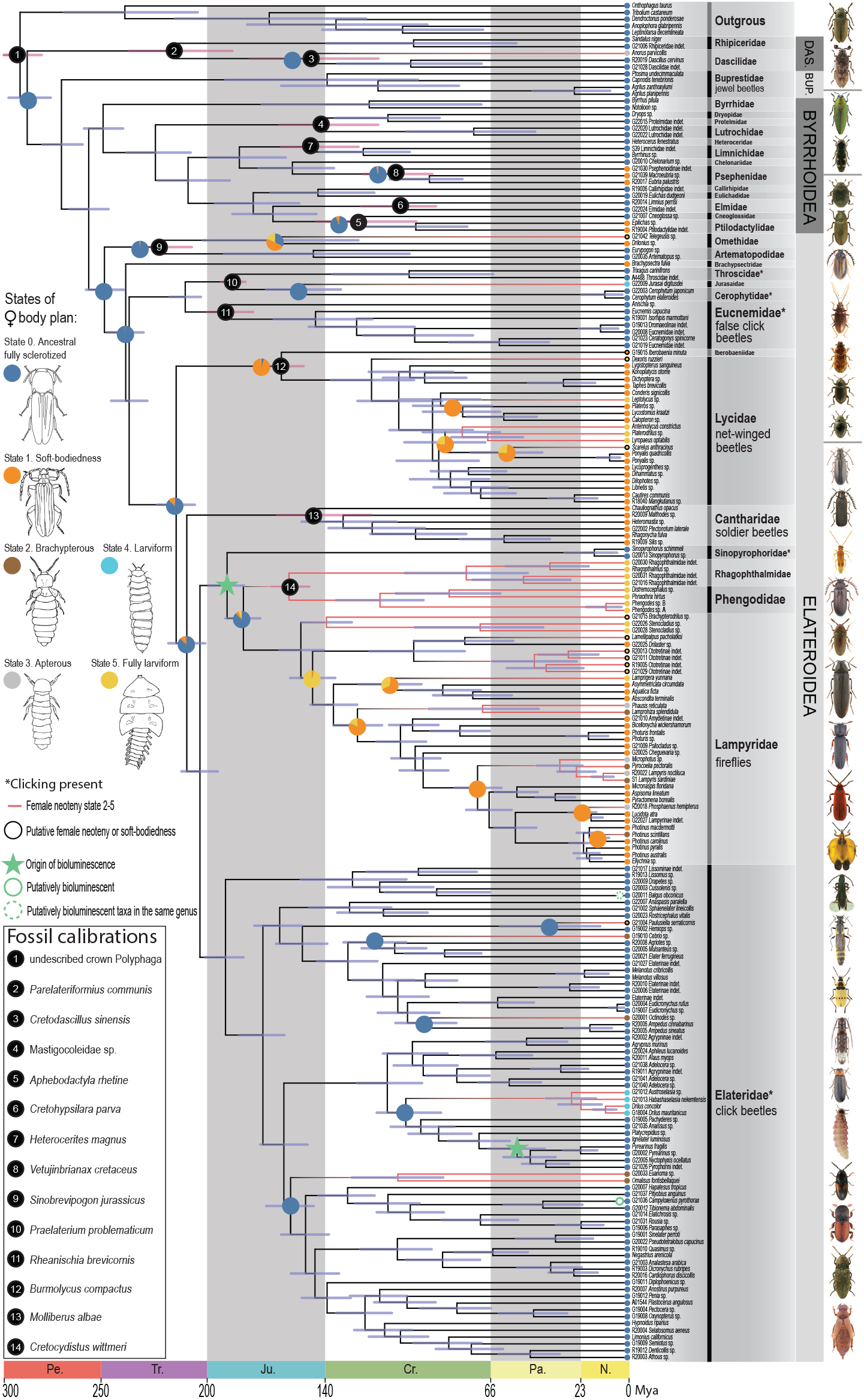
Divergence time estimation and trait reconstruction in Elateriformia. Time-calibrated phylogenetic tree based on ASTRAL species tree topology resulting from Dataset B-4189-NT and using Dataset A with nucleotide sequences under the autocorrelated rates (AR) clock model in MCMCtree. Node bars represent 95% credibility intervals (CI). Numbered black circles indicate fossil calibration points at corresponding nodes. Pink horizontal bars show the age uncertainty (95% CI) for calibrated nodes. Trait evolution reconstruction (6 states, only three of them reconstructed at nodes) is shown on the phylogeny: neotenic modifications of the female body plan types are illustrated on the left and shown in detail in Figure 8. The full trait-reconstruction tree is shown in Figure S54. Bioluminescence origins and presence are mapped without a formal analysis (see Methods). Family-level classification is shown on the right with representative beetle photographs. *Asterisks indicate families where clicking ability is present. Geological time scale shown at bottom: Pe = Permian, Tr = Triassic, Ju = Jurassic, Cr = Cretaceous, Pa = Paleogene, N = Neogene. Mya = Million years ago.

In contrast, AA analyses showed considerable topological variation. Coalescent AA analyses, excluding dataset G-962-AA-decisive and its subsets, recovered Lycidae + Iberobaeniidae as the sister group to all remaining lineages, with Cantharidae and the lampyroid clade nested within the polyphyletic Elateridae (Figures 4B, S22, S23, S25, S26). ML AA analyses exhibited two distinct patterns: some recovered Cantharidae as the first-diverging lineage (Figures 4B, S15–S17, S21A), while others recovered Lycidae + Iberobaeniidae as the first split (Figures S18, S19). In both ML AA patterns, Elateridae was rendered polyphyletic by the nested placement of Cantharidae and the lampyroid clade. Notably, dataset G-962-AA-decisive and its subsets recovered the same branching order as most NT analyses, including monophyletic Elateridae. All these conflicting nodes exhibited low quartet support and short internal branch lengths in coalescent units, suggesting these relationships fall within a hypothetical anomaly zone (Figures 4, S22–S31).

Analyses of the 200 fastest- and slowest-evolving genes from the reduced AA dataset (G-962-AA-decisive) both recovered Lycidae + Iberobaeniidae diverging first, followed by Cantharidae (Figure 4D). However, the monophyly of Elateridae differed between these analyses: the fastest-evolving genes recovered a monophyletic Elateridae (Figures 4D left, S24, S28), whereas the slowest-evolving genes recovered Elateridae as polyphyletic (Figures 4D right, S24, S28). The monophyly of Elateridae was thus recovered in all NT analyses and AA analyses of the fastest-evolving genes (Figure 5C). Within Elateridae, at the NT level, the positions of Elaterinae, Lissominae, and Protelaterini differed between ML and coalescent analyses.

The recently proposed family Sinopyrophoridae was always recovered as a sister to the soft-bodied lampyroids (Lampyridae, (Phengodidae, Rhagophthalmidae)) (Figures 4A, B, 5C, S15–S32). Within Lampyridae, we consistently recovered Ototretinae as paraphyletic. *Stenocladius* + *Brachypterodrilus* diverged first as the sister of other lampyrids, followed by the rest of Ototretinae as the next branch (Figure 6, S15–S28). Additional discordance within Lampyridae included the positions of Psilocladinae, Amydetinae, and Cheguevariinae, which differed across ML and coalescent analyses at the NT and AA levels (Figures 6, S15–28).

### Phylogenomic concordance and conflict analysis

We found gene, site, and quartet discordance along the tree backbone. Branches exhibiting the most severe topological conflicts were consistently characterized by short lengths (measured in coalescent units), likely reflecting rapid evolutionary radiations and/or incomplete lineage sorting which may have obscured phylogenetic signal (Figure 4A, B).

Having identified the primary areas of topological instability, we next investigated the underlying causes of these conflicts. High bootstrap support values alone cannot distinguish between well-resolved relationships and those affected by rapid radiations, incomplete lineage sorting, or systematic error (Thomson & Brown, 2022; Reddy et al., 2017), necessitating deeper investigation of the phylogenetic signal underlying these nodes through concordance analyses and anomaly zone assessments (Kapli et al., 2020; Steenwyk et al., 2023; Young & Gillung, 2020).

### Concordance and discordance patterns

Concordance factor (CF) analyses of the four major Elateriformia superfamilies revealed substantial variation in gene tree congruence despite uniformly maximum statistical support (lpp = 1.00) for all backbone splits in the ASTRAL and wASTRAL species trees (Figures 5A, S22, S25). Gene, site, and quartet CFs (gCF, sCF, and qCF) were calculated for all internal branches, with corresponding concordance vectors (CV) comprising *ψ*_1_ (concordance factor), *ψ*_2_, and *ψ*_3_ (the two primary discordance factors obtained by nearest-neighbour interchanges), and *ψ*_4_ (all other discordant topologies) (Figure 5B).

The split supporting Dascilloidea as the sister of remaining Elateriformia (branch 243: lpp = 1.00, bl = 0.220, branch length in coalescent units; see Figure 5A for branch numbers) showed gene concordance vector (gCV) (29.02%, 5.78%, 3.68%, 61.52%) and quartet concordance vectors (qCV) (57.16%, 23.54%, 19.31%, 0.00%) (Figure 5B). Dascilloidea monophyly (branch 247: lpp = 1.00, bl = 1.424) showed gCV (69.51%, 0.68%, 0.45%, 29.36%) and qCV (83.98%, 9.35%, 6.68%, 0.00%). The split placing Buprestoidea as sister to Byrrhoidea + Elateroidea (branch 242: lpp = 1.00, bl = 0.184) showed gCV (14.51%, 4.43%, 2.88%, 78.19%) and qCV (44.60%, 32.14%, 23.26%, 0.00%). Buprestoidea monophyly (branch 251: lpp = 1.00, bl = 2.963) showed gCV (90.93%, 0.09%, 0.03%, 8.95%) and qCV (96.58%, 1.96%, 1.46%, 0.00%). The split supporting Byrrhoidea monophyly (branch 222: lpp = 1.00, bl = 0.088) showed markedly lower concordance despite maximum statistical support, with gCV (3.97%, 1.82%, 1.21%, 93.00%) and qCV (39.00%, 34.44%, 26.56%, 0.00%). Similarly, the split supporting Elateroidea monophyly (branch 221: lpp = 1.00, bl = 0.233) showed gCV (4.96%, 1.29%, 0.74%, 93.00%) and qCV (47.23%, 30.22%, 22.55%, 0.00%). The split placing Byrrhoidea *s. l*. as the sister of Elateroidea (branch 241: lpp = 1.00, bl = 0.161) showed gCV (5.54%, 1.85%, 0.79%, 91.82%) and qCV (43.27%, 30.59%, 26.14%, 0.00%).

The gCV for branches with short internal lengths (0.088–0.233) showed substantial *ψ*_4_ values (78.19%– 93.00%), indicating that most gene trees lacked monophyly in one or more constituent clades and could not be classified as decisive for these branches (Minh et al., 2020b). In contrast, longer branches (1.424–2.963) showed lower *ψ*_4_ values (8.95%–29.36%).

Anomaly zone analyses of wASTRAL species trees from AA datasets A-4189-AA and E-4224-AA-raw identified 7 and 5 pairs of consecutive internal branches, respectively, that satisfied the anomaly zone criterion *y < a*(*x*) (Degnan & Rosenberg, 2006), where x represents the parent (*i*.*e*., deeper) branch length, y the child (shallower) branch length, and a(x) the anomaly zone threshold (Figure S31). In the species tree resulting from the analysis of dataset A-4189-AA, seven anomalous branch pairs were detected (Figure S31A, C): (1) consecutive internal branches at the divergence of Byrrhidae, where the ancestral branch within the Byrrhoidea + Elateroidea clade (x3 = 0.120, a(x) = 0.254) is followed by the branch leading to the Byrrhidae split (y3 = 0.079); (2) consecutive internal branches at the placement of Brachypsectridae, where the ancestral branch within the ((Brachypsectridae, Eucnemidae), ‘crown Elateroidea clade’)) (x4 = 0.058, a(x) = 0.589) is followed by the branch leading to the Brachypsectridae + Eucnemidae split (y4 = 0.257); (3) consecutive internal branches at the divergence of Lissominae + Thylacosterninae (Elateridae) from rest of the polyphyletic Elateridae, Cantharidae and the lampyroid clade, where the ancestral branch (x6 = 0.115, a(x) = 0.268) is followed by the branch leading to their split (y6 = 0.098); (4) consecutive internal branch leading to Cantharidae split, where the ancestral branch (x7 = 0.098, a(x) = 0.337) is followed by the branch leading to the rest of polyphyletic Elateridae + lampyroid split (y7 = 0.099); (5) consecutive internal branches at the divergence of the lampyroid clade + rest of polyphyletic Elateridae, where the ancestral branch (x8 = 0.099, a(x) = 0.331) is followed by the branch leading to their split (y8 = 0.218); (6) consecutive internal branches within Byrrhoidea, where the ancestral branch of Cneoglossidae + Ptilodactylidae (x9 = 0.135, a(x) = 0.209) is followed by the branch leading to the ((Chelonariidae, Psephenidae), (Elmidae, (Heteroceridae, Limnichidae))) split (y9 = 0.191); and (7) consecutive internal branches where the ancestral branch of Chelonariidae + Psephenidae (x10 = 0.191, a(x) = 0.093) is followed by the branch leading to the (Elmidae, (Heteroceridae, Limnichidae)) split (y10 = 0.091).

In dataset E-4224-AA-raw, five anomalous branch pairs were detected (Figure S31B, C) at the same phylogenetic positions as nodes 3, 4, 6, 7, and 8: (1) x3 = 0.122, a(x) = 0.248, y3 = 0.075; (2) x4 = 0.062, a(x) = 0.549, y4 = 0.266; (3) x6 = 0.127, a(x) = 0.232, y6 = 0.110; (4) x7 = 0.110, a(x) = 0.289, y7 = 0.104; (5) x8 = 0.104, a(x) = 0.310, y8 = 0.207. Non-anomalous branch pairs (1, 2, 5, and in E-4224-AA-raw also 9 and 10) corresponded to deeper backbone splits with longer parent branches (0.416–0.484 in coalescent units), resulting in negative a(x) values, or to cases where y exceeded the a(x) threshold (Figure S31C).

Gene-tree quartet frequencies for internal branches of the ASTRAL species trees were visualized using DiscoVista analyses (Figures 4C, S30A, B) and displayed as pie charts or quartet scores on tree nodes (Figures 4A, S23, S24, S27, S28). Quartet frequencies represent the proportion of gene trees supporting the primary topology (q1) and the two alternative topologies obtained by nearest-neighbour interchange (q2 and q3). We analysed dataset B-4189-NT using 4,189 gene trees with poorly supported branches collapsed (UFB ≤50) and dataset F-4224-NT-raw using 4,224 uncollapsed gene trees.

Topology frequencies were indecisive at several branches. Buprestoidea as sister to Byrrhoidea + Elateroidea showed q1 = 45%, with alternatives q2 = 23%, q3 = 32%. Byrrhoidea monophyly exhibited similar uncertainty (q1 = 39%, q2 = 27%, q3 = 34%), while Elateroidea monophyly showed slightly higher frequency for the preferred topology (q1 = 47%, q2 = 23%, q3 = 30%). Within Byrrhoidea, the placement of Elmidae (split from Ptilodactylidae + Cneoglossidae) showed high ambiguity (q1 = 38%, q2 = 24%, q3 = 38%). Within Elateroidea, Omethidae + Artematopodidae, as the first-diverging lineage, showed frequencies q1 = 47%, q2 = 23%, q3 = 30%, while Brachypsectridae, as sister to the remaining Elateroidea, was supported by a higher frequency q1 = 66%, compared to the alternatives (q2 = 19%, q3 = 15%). The position of the Eucnemidae-TJC clade relative to Brachypsectridae remained highly uncertain (q1 = 36%, q2 = 34%, q3 = 30%). Within the crown clade of Elateroidea, Lycidae + Iberobaeniidae as the sister to Cantharidae + (elaterid-lampyroid clade) showed q1 = 70%, q2 = 7%, and q3 = 23%, while the preferred placement of Cantharidae was based on a weaker signal (q1 = 41%, q2 = 24%, q3 = 35%). Elateridae monophyly showed values q1 = 45%, q2 = 17%, q3 = 38%. Within lampyroids, Sinopyrophoridae was the sister of the remaining families, with q1 = 57%, q2 = 21%, and q3 = 22%.

Amino acid analyses (datasets A-4189-AA and E-4224-AA-raw) revealed concordant or more pronounced patterns of gene tree discordance at these nodes (Figures 4B, S23, S24, S26, S28, S29). Similar patterns emerged across wASTRAL analyses using uncollapsed gene trees from datasets B-4189-NT and A-4189-AA (4,189 genes each), K-3583-NT-SymTest (3,583 genes), and raw datasets E-4224-AA-raw and F-4224-NT-raw (4,224 genes each) (Figures S26, S27). Analyses of decisive gene subsets (G-962-AA-decisive and H-962-NT-decisive, 962 genes each) and subsets of fastest- and slowest-evolving genes (L-200-fast-AA-decisive and M-200-slow-AA-decisive, 200 genes each) yielded consistent patterns of gene tree discordance (Figure S28).

Four-cluster likelihood mapping (FcLM) analyses were performed to evaluate the distribution of phylogenetic signal among alternative quartet topologies for six contentious relationships, using datasets A-4189-AA and B-4189-NT (Figure S32). For Hypothesis 1 (Buprestoidea sister to Byrrhoidea + Elateroidea), the AA data showed higher support, with 58.7% of quartets favouring this arrangement, compared to 45.1% at the NT level. Byrrhoidea monophyly (Hypothesis 2) received conflicting support: at the AA level, 48.1% of quartets supported monophyletic Byrrhoidea while 44.1% favoured the alternative topology with Byrrhidae sister to Elateroidea, whereas at the NT level, the alternative topology was more strongly supported (57.5%) compared to monophyly (26.7%).

The sister relationship between Psephenidae + Chelonariidae and Limnichidae + Heteroceridae (Hypothesis 3a) was supported by 63.9% of quartets at the AA level and 87.5% at the NT level. For Hypothesis 3b, Elmidae as sister to the clade ((Psephenidae, Chelonariidae), (Limnichidae, Heteroceridae)) received support from 98.8% of quartets at the AA level. However, 66.7% of quartets in the NT data suggested either Elmidae closer to Callirhipidae + Eulichadidae or, alternatively, the clade (Psephenidae, Chelonariidae, Limnichidae, Heteroceridae) as sister to Cneoglossidae + Ptilodactylidae, depending on root placement. The analyses favoured Brachypsectridae as the sister group of (Eucnemidae, (Throscidae, (Jurasaidae, Cerophytidae))) (Hypothesis 4) at both AA and NT levels, with 60.8% and 55.3% of quartets supporting this topology, respectively.

For Hypothesis 5, 73.6% of quartets at the AA level supported a closer relationship between Cantharidae and the elateroid-lampyroid clade, while at the NT level, the alternative topology with Lycidae + Iberobaeniidae closer to the elateroid-lampyroid clade was supported by 49.2%. Finally, Sinopyrophoridae as sister to the lampyroid clade (Hypothesis 6) received strong support at both the AA (85.8%) and NT (70.7%) levels.

### Sensitivity analyses and subsampling

A comparison of the 200 fastest-evolving (L-200-fast-AA-decisive) and 200 slowest-evolving genes (M-200-slow-AA-decisive) at the AA level, derived from dataset G-962-AA-decisive, revealed significant differences in gene properties (Figure 4E). The 200 fastest-evolving genes exhibited RCFV values of 0.010–0.015 and pairwise divergence of ∼0.0010, lower missing data, and longer average branch lengths, consistent with their higher evolutionary rates, while the 200 slowest-evolving genes showed RCFV values of 0.005–0.008 and pairwise divergence of ∼0.0005, with slightly higher alignment lengths and parsimony-informative sites. UFB support values remained high across all three subsets. Most coalescent analyses using the full set of 962 genes at both AA and NT levels, as well as both AA subsets, recovered the backbone relationships among the four superfamilies, monophyletic Byrrhoidea and the branching order of Lycidae + Iberobaeniidae diverging first, followed by Cantharidae, matching the topology consistently obtained from NT datasets (Figures 4D; S24A–H, S28A–H). However, the monophyly of Elateridae differed between subsets: analyses using the 200 fastest-evolving genes recovered monophyletic Elateridae (Figures 4D, right, S24C, D, S28C, D), while analyses using the 200 slowest-evolving genes recovered polyphyletic Elateridae with the lampyroid clade nested within Elateridae (Figures 4D, left, S24E, F, S28E, F).

### Evolutionary timeline of Elateriformia and origins of neoteny and bioluminescence

Model comparison using mmc3r analyses (Table S7) identified the autocorrelated rates (AR) clock model as the best-fitting model for all datasets except Dataset2 at the nucleotide level, where the independent rates (IR) clock model was preferred. Both AR and IR clock models, applied to two distinct datasets at the AA and NT levels, produced similar age estimates for major splits (Figures 6, 7, S43–S52), with only minor differences at deeper nodes. Examination of ESS values and a posteriori tree distribution across all chains demonstrated broad convergence of summary statistics (Figures S34– S39). The slope of the regression line in the infinite-sites plots was 0.2379–0.2889, indicating that for every 1 Mya of divergence, 0.2379–0.2889 Mya of uncertainty is added to the 95% confidence interval (Figures 7C, D, S40). We observed wider 95% confidence intervals at some nodes with extended stem branches preceding crown diversification. These nodes represent lineages with ancient origins but relatively young crown ages, a pattern that arises either from incomplete sampling of early-diverging lineages or from a delayed onset of diversification within these clades. Examples include Artematopodidae, Throscidae, Omethidae, and Byrrhidae (Figures 6, S43–S52).

**Figure 7.**
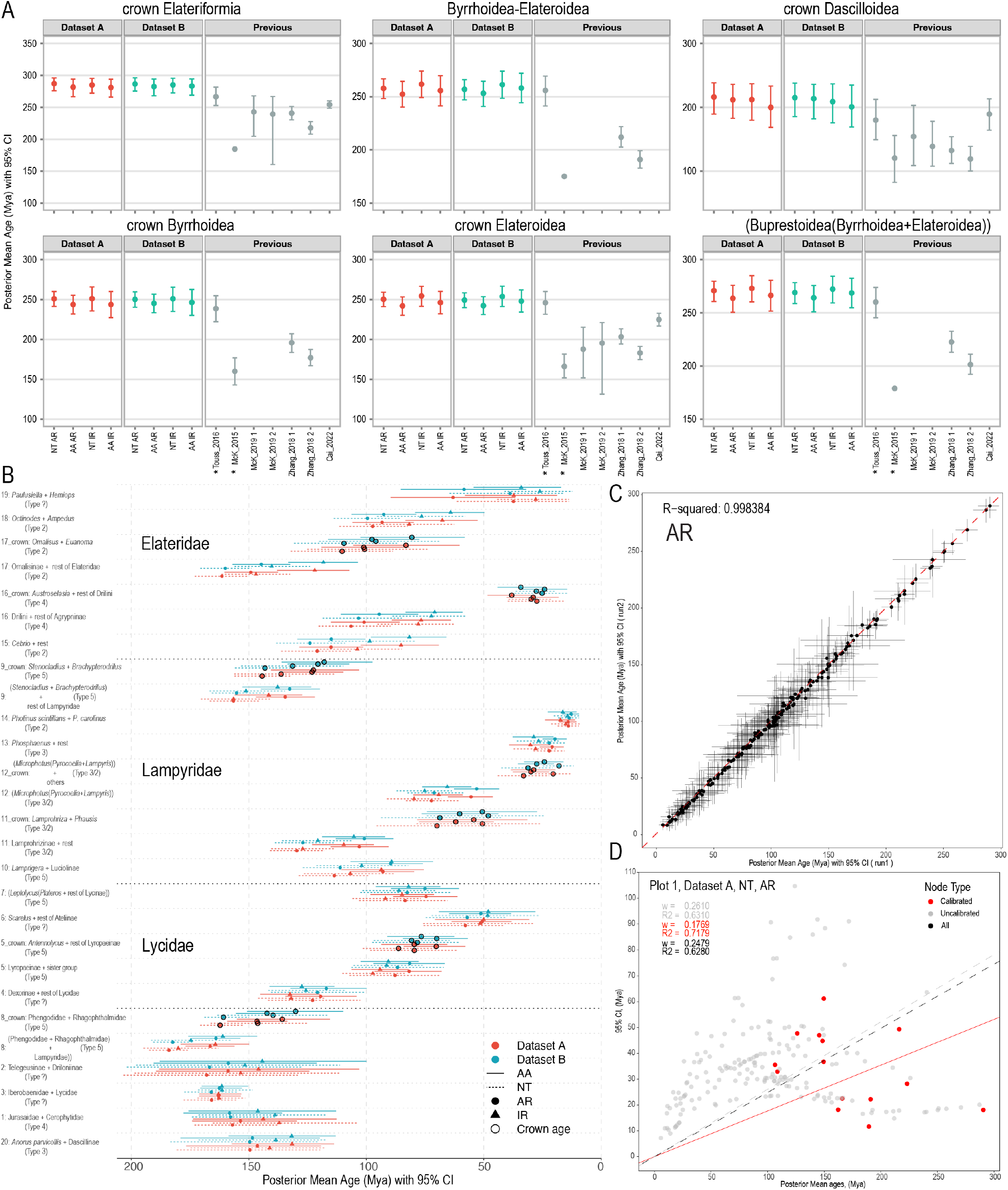
Divergence time estimation results. A) Comparison of age estimates for superfamilies (major clades) between the current study and previous works. Data are shown as means with 95% credibility intervals (CI) unless otherwise noted. *Asterisked studies (McKenna et al., 2015; Toussaint et al., 2017) show medians with 95% CI; some McKenna et al. (2015) values display medians only due to unreported 95% CI in the original study. B) Origin and divergence times of neotenic lineages. Age estimates are compared across datasets (A, B), data types (AA = amino acids, NT = nucleotides), and clock models (AR = autocorrelated rates, IR = independent rates) for different neotenic modification states (State 0–5). Crown ages (black circles) are shown for clades with multiple sampled representatives. Horizontal bars indicate 95% CI. C) Convergence plot between merged runs of Dataset A (n=4) and Dataset B (n=2) using NT sequences under the AR clock model. Posterior mean ages with 95% CI are compared (*R*^2^ = 0.998). D) Infinite-sites plot for Dataset A with NT sequences under the AR clock model. Posterior 95% CI width plotted against mean divergence times. Red dots represent calibrated nodes, grey dots represent uncalibrated nodes. Regression lines through the origin are shown for calibrated nodes (solid red), uncalibrated nodes (grey dashed), and all nodes (black dashed). For optimal readability, please refer to the PDF version of this figure.

Density plots of posterior age distributions for calibrated nodes demonstrated strong congruence across datasets and clock models (Figures S41, S42). Most nodes showed unimodal distributions with substantial overlap between the AA and NT datasets under both AR and IR models, indicating robust age estimates that are largely independent of the analytical strategy. Posterior distributions were generally narrow and tightly clustered around mean estimates for well-constrained nodes (e.g., crown Elateriformia, crown Elateroidea, Lycidae + Iberobaeniidae). Some deeper nodes exhibited broader posterior distributions with greater variation between methods (*e*.*g*., the branches Rhipiceridae + Dascillidae, Artematopodidae + Omethidae, and Limnichidae + Heteroceridae), reflecting increased temporal uncertainty at ancient divergences. Notably, effective priors (shown in blue) were consistently overridden by the phylogenetic data for all calibrated nodes, demonstrating that posterior age estimates were primarily data-driven rather than prior-constrained.

We consistently identified an Early Permian origin for Elateriformia, with crown mean ages ranging from 280.99 to 287.13 Mya across eight dating strategies (95% CI: 265.66–296.05 Mya, representing the lowest lower and highest upper bounds, respectively) (Figures 7A, S43–S52). The origin of all four major superfamilies followed during the Late Permian (Figures 6, 7A). The first internal split occurred in the Late Permian for Byrrhoidea (243.72–251.10 Mya; 95% CI: 227.26–265.75 Mya) and Elateroidea (242.07–254.45 Mya; 95% CI: 230.28–266.50 Mya), and later in the Late Triassic to Early Jurassic for Dascilloidea (199.95– 216.12 Mya; 95% CI: 168.51–238.57 Mya) (Figures 6, 7A, S43–S52).

The Jurassic period marked the origin of most extant families, with significant intrafamilial diversification occurring during the Cretaceous (Figures 6, S53). The latest splits between branches representing modern families are separations of Protelmidae + Dryopidae and Cneoglossidae + Ptilodactylidae (Figures 6, S43–S53). In contrast, the oldest splits lead to the modern Byrrhidae and Brachypsectridae. At the subfamily level, we identified some ancient lineages like Anischiinae (Eucnemidae) and Lissominae (Elateridae), and a delayed diversification of modern lycid and some elateroid subfamilies (Fig. 6). An increase in divergence events can be observed at 220–140 Mya at the family level and at 120–100 Mya in all divergence events (Figure S53).

We estimated ancestral states by allowing direct transitions between states. The selected results of the analysis are shown in the dated phylogenetic tree (Figure 6); the full tree is shown in Figure S54. We inferred a compact, well-sclerotized body with coadapted elytra and abdomen as the ancestral state (connected with the presence of a clicking mechanism in well-sclerotized Elateroidea except for Artematopodidae). State 1, soft-bodied forms with similar males and females, is derived and originated multiple times. The analysis suggested independent origin of soft-bodiedness in Lycidae + Iberobaeniidae and Cantharidae. The forms with substantial sexual dimorphism, leading to flightless and larviform females, originated in branches consisting of both well-sclerotized and soft-bodied forms. The shifts were mostly sudden, resulting in the uniform modification of all descendants. Only fireflies exhibit high variability, but the sample size is insufficient for a robust analysis.

Our analyses revealed that neotenic lineages first diverged from their non-neotenic sister groups during the Jurassic period (Figures 6, 7B, S43–S52; Table S9). The earliest split of the exclusively neotenic clade Phengodidae + Rhagophthalmidae is estimated to have occurred at ages ranging from 164.02–184.01 Mya across eight dating strategies (95% CI: 115.45–197.65 Mya). The crown age of this exclusively neotenic clade is only slightly younger (130.08–162.25 Mya; 95% CI: 109.84– 197.65 Mya). Other splits between modified lineages include *Stenocladius* + *Brachypterodrilus* in Lampyridae (117.94–144.35 Mya; 95% CI: 97.41–170.51 Mya) and *Omalisus* + *Euanoma* in Elateridae (80.61–110.31 Mya; 95% CI: 58.12–173.11 Mya). For other neotenic lineages represented by single taxa in our study (Iberobaeniidae, Jurasaidae, Telegeusinae, *Dexoris ruzzieri, Leptolycus* sp., *Lamprigera* sp., Drilini, *Paulusiella serraticornis*, and *Anorus parvicollis*), only stem divergence times from non-neotenic relatives can be reported, ranging from 26.12–167.95 Mya (Figures 6, 7; Table S9). Notably, we found no apparent correlation between lineage age and the degree of morphological modification among neotenic taxa.

Bioluminescence evolved independently in Elateriformia at least four times (Figure 6; Table S10). The earliest origin occurred in the lampyroid clade, which diverged from Elateridae at 181.85–204.73 Mya (95% CI: 155.10–217.78 Mya), with crown diversification of bioluminescent lampyroids (Sinopyrophoridae + remaining lampyroids) occurring at 169.45–191.79 Mya (95% CI: 155.10–217.78 Mya). Bioluminescence subsequently evolved independently within Pyrophorini (Elateridae), which split from Platycrepidiini at 68.79–73.79 Mya (95% CI: 53.10–87.09 Mya), with crown diversification at 23.96–53.41 Mya (95% CI: 17.16–86.16 Mya), in *Campyloxenus pyrothorax* (Elateridae), which split from *Tibionema abdominalis* at 30.22–41.97 Mya (95% CI: 15.83–59.69 Mya), and in the *Balgus* lineage containing the bioluminescent *B. schnusei* (not sampled here), which diverged from *Cussolenis* at 64.50–91.77 Mya (95% CI: 41.13–116.36 Mya).

### Phenotypic diversity across Elateriformia lineages

Based on our collection and material deposited in major museums (see Methods), we evaluated the phenotypes of all known families and important subfamilies of Elateriformia with a focus on taxa with proven or hypothesized ontogenetic modifications (Figures 1, 8; Text S2). Our comparative morphological approach documents substantial phenotypic variation. Examples and simplified information on phenotypic modifications are shown in Figure 8. An overview of the included modified lineages and their character states defining phenotypic categorization is provided in Table S9. The modifications are pronounced in females. Females lose wings and have shortened, vestigial, or absent elytra, but they also have modified legs, antennae, and mouthparts, are weakly sclerotized, and have a flexible, sometimes translucent cuticle. The modifications are more pronounced in posterior body parts, especially the abdomen, which resembles that of larvae, is flexible, and has a similar level of sclerotization in ventral and dorsal sclerites. Thoracic morphology includes the gradual loss of length differentiation of segments, simplification of sclerites, loss of sclerotization, and, finally, a larva-like thoracic morphology. Then, only the head is adult-like in the sexually mature individual. At the extreme, the females of some groups remain fully larviform, and only the sexual duct opens when they reach maturity. Males of the taxa having modified females are regularly soft-bodied, often with a modified cranium and miniaturized mouthparts, sometimes with shortened, vestigial, or lost elytra. The modified males, *e*.*g*., some net-winged beetles, resemble less modified females. For example, a similar modification is encountered in the female of *Omalisus* and the male of *Dexoris chome* in Figure 8.

**Figure 8.**
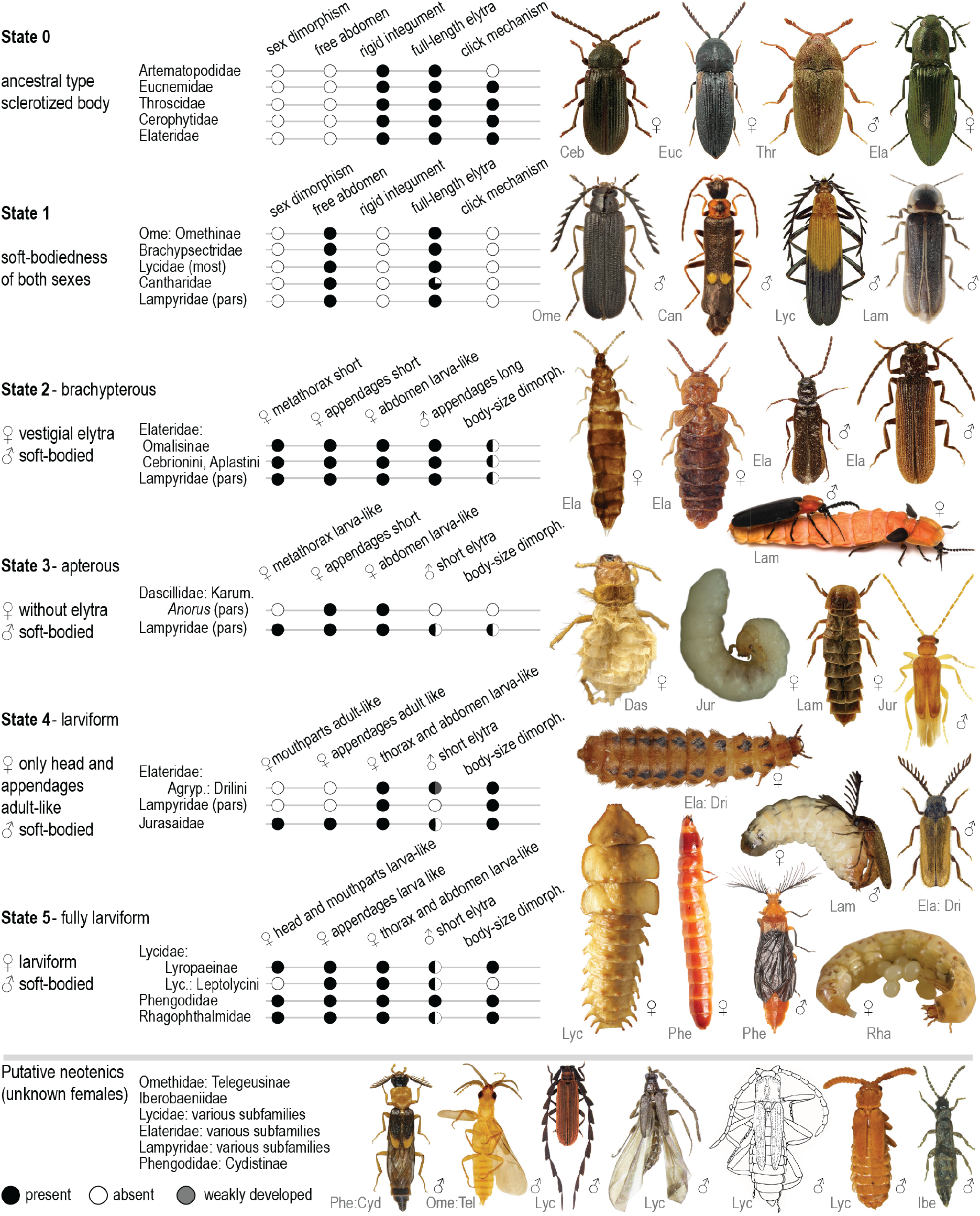
An overview of phenotypic variation in Elateriformia grouped by degree of neoteny. Six states (State 0 through State 5) represent a gradient from complete metamorphosis (State 0: fully sclerotized body with functional elytra in both sexes) to fully larviform adult females (State 5: larva-like morphology). The classification is based on sclerotization patterns, elytral development, and retention of larval morphology in sexually mature individuals. The upper section depicts groups with known males and females, while the bottom section shows examples of groups with putative neotenic females where only males are known.

Ontogenetic modifications are gradual yet can be categorized for the formal trait reconstruction (Figure 8). We propose levels based on sclerotization, elytral development, modification of the thorax, and the preservation of adult head morphology in sexually mature individuals (Figure 8). We designate hypothesized ancestral fully sclerotized forms with abdomens coadapted to elytra and connate abdominal ventrites as level 0. This phenotype resembles relatives from other sub- and infraorders of beetles and is dominant in Dascilloidea, Buprestoidea, and Byrrhoidea, and is also present in many elateroids (Artematopodidae and all clicking forms). Level 1 is designated as soft-bodiedness. At this level, both sexes are capable of flight and differ from level 0 in the flexible abdomen, which is often longer than the elytra. Abdominal ventrites are feebly sclerotized and never connate. Level 2 is defined by brachelytrous females with larva-like abdomens, lower meta- and mesothorax length ratios, and shortened appendages. Level 3 is defined by the complete loss of elytra, larva-like meso- and metathorax, but adult-like prothorax. The next level 4 is defined by larval thoracic and abdominal segments, with only the head and legs metamorphosing into an adult-like form. The most modified females, level 5, are larviform and have a non-sclerotized cuticle. Some shed the pupal cuticle with resulting changes in cuticular structure and coloration, while others retain it, forming a two-layered cuticle. The functional sexual ducts and observed copulation confirm that the individuals are mature.

## Discussion

### Elateriformia datasets

Large-scale phylogenomic datasets significantly contribute to resolving contentious relationships (Misof et al., 2014; McKenna et al., 2019; Zhu et al., 2019; Zuntini et al., 2024). Here, we present a study focused on one infraorder of beetles – Elateriformia (Table S1). Until now, the largest datasets used to infer its phylogeny have been limited to subsets in broader studies. These have contained 4,800 orthologs and 14 terminals (McKenna et al., 2019) and ∼2,000 orthologs and 13 terminals (Creedy et al., 2026). A comparable volume of data per sample was used only to resolve the relationships of Sinopyrophoridae and fireflies ( ∼4,000 orthologs and 25 terminals; Kusy et al., 2021) or net-winged beetle phylogeny ( ∼4,000 orthologs and 22 terminals; Kusy et al., 2019). The 95-ortholog Coleoptera dataset was produced by Zhang et al. (2018) and comprised 23 elateriform families (79 terminals, ∼71,000 aligned NT positions). The Zhang et al. dataset was pruned and reanalysed by Cai et al. (2022) and, in a slightly expanded form, by McKenna et al. (2019) and Kusy et al. (2018b) (Figures 2, S1). The dryopoid families were studied using the UCE dataset (75 terminals, up to 201,000 NT positions) (Hayashi et al., 2024). Until now, these studies have presented the largest datasets ever analysed to resolve the phylogeny of Elateriformia and have provided the backbone of Elateriformia phylogeny, yielding both decisive and ambiguous relationships. We add a high volume of new data. The raw, unfiltered NT dataset contains 195 ingroup terminals representing 31 families, 4,224 single-copy orthologs, and 10.1 million aligned positions per sample, *i*.*e*., a 2-billion-cell matrix (Tables S1, S3–S5; Figure S3). The dense sampling and volume of data provide new information on the newly sampled families and deeper insight into contentious relationships.

### The tree topology and implications for systematics

Our preferred topology (Figure 6) is based on explicit rejection of alternative arrangements through hypothesis testing (Figures 4, 5, S15–S32). Current results show a higher congruence with the classification accepted until recently (Zhang et al., 2018; McKenna et al., 2015; Bouchard et al., 2011; Figures 2B, C, S1) than with the relationships proposed by the most recent studies (Cai et al., 2022; Douglas et al., 2021; McKenna et al., 2019; Hayashi et al., 2024) (Figures 2D; S1). We show that monophyletic Elateriformia can be subdivided into four superfamilies (Figures 4–6, S15–S32).

Dascilloidea emerges as the earliest split, followed by Buprestoidea as the next branch. The crown is represented by the reciprocally monophyletic Byrrhoidea *s. l*. and Elateroidea (Figures 4–6, S15–S32). Our tests reject the paraphyly of Byrrhoidea (Douglas et al., 2021; McKenna et al., 2019; Cai et al., 2022; Hayashi et al., 2024). Consequently, the recent reclassification reintroducing Dryopoidea alongside Byrrhoidea (Cai et al., 2022) lacks empirical support (Boudinot et al., 2023a). There are 15 possible rooted topologies for Byrrhidae, Dryopoidea *sensu* Cai et al. (2022), Elateroidea, and Buprestoidea, and six of them have been recently recovered by the analyses of multigene datasets (Hayashi et al., 2024; Zhang et al., 2018; McKenna et al., 2019; Cai et al., 2022). The resurrection of Dryopoidea (Cai et al., 2022) was proposed based on the reanalysis of data that initially recovered a topology congruent with earlier superfamily delimitation and current results (Figure 2C) (Lawrence & Newton, 1995; Bouchard et al., 2011; Zhang et al., 2018). Notably, all competing topologies that suggest Byrrhoidea’s paraphyly garnered weak support (Hayashi et al., 2024), indicating a conflicting signal and very short branches, possibly resulting from rapid radiation and incomplete lineage sorting. If the paraphyly of Byrrhoidea and Dryopoidea *sensu* Cai et al. (2022) is rejected, there is no basis for the delimitation of two superfamilies. Both groups are small (430 and 3,935 spp., 1 and 13 families, respectively), and their morphological disparity does not warrant separate superfamilies (Hayashi et al., 2024; Bocak et al., 2014; Lawrence et al., 2011; Lawrence, 1988). Therefore, we synonymize Dryopoidea and return to Byrrhoidea *sensu lato* (Lawrence & Newton, 1995; Bouchard et al., 2011; Lawrence et al., 2011).

Furthermore, we reject competing hypotheses regarding jewel beetles’ close relationships with various byrrhoid families proposed by morphological analyses (Crowson, 1982; Lawrence, 1988; Lawrence et al., 2011) (Figure 2A) and early molecular studies using a few mtDNA and rRNA genes or mitogenomes (Bocakova et al., 2007; Bocak et al., 2014; Sagegami-Oba et al., 2007; Timmermans et al., 2016). The early multigene analyses also failed to resolve this ambiguity with strong support. Buprestidae was recovered as a sister of Byrrhoidea *s. l*. + Elateroidea in multigene analyses (Zhang et al., 2018), but different settings or datasets yielded the sister position to Byrrhidae (Cai et al., 2022) or Dryopidae + Heteroceridae (McKenna et al., 2019) (Figures 2D, S1). Our data, except for a single analysis (Figures 5C, S28C, D), support Buprestoidea as the second deepest branch in Elateriformia, and justify its superfamily rank (Figures 4–6, S15–S32).

With much denser sampling than in previous phylogenomic analyses, we also investigated the internal structure of superfamilies at the family and, to some extent, the subfamily levels. Dascilloidea was represented by both extant families, Dascillidae and Rhipiceridae, including *Anorus* (Karumiinae), which contains a species with incompletely metamorphosed, phenotypically distinct females that had been placed in conflicting relationships (Karumiidae *sensu* Crowson 1971; Figure 8). Buprestoidea was represented only by the hyperdiverse Buprestidae (15,000 spp.), as the small endemic family Schizopodidae (7 spp.) was unavailable. The Schizopodidae were earlier analysed using mitogenomes and multiple protein-coding DNA fragments produced by Sanger sequencing (McKenna et al., 2015; Timmermans et al., 2016; Evans et al., 2015) and recovered as a sister to Buprestidae, in agreement with morphology (Nelson & Bellamy, 1991; Kolibac, 2000; Lawrence et al., 2011). The fossil record suggests an early origin for the family, consistent with the early diversification of jewel beetles (Figure 6; Legalov, 2025). The internal structure of jewel beetles confirms the deep position of Polycestinae and the delayed origin of the hyperdiverse *Agrilus* (*>*3,000 spp., Agrilinae), the largest animal genus on Earth (Figures 6, S54; *>*3,000 spp., Jendek & Poláková, 2014; Kelnarova et al., 2019). This radiation is undoubtedly linked to the origin of angiosperms, as in other beetle super-radiations (*e*.*g*., Curculionidae and Chrysomelidae; McKenna et al., 2009, 2019; Shin et al., 2018; Evans et al., 2015).

The internal relationships within Byrrhoidea have been confounded by the inclusion of Buprestidae within the clade in most earlier studies and by repeated questions about the superfamily’s monophyly (see above). Additionally, the group’s radiation is ancient (Figure 6), and phylogenomic information may have been eroded since the group’s origin (Figure 5A, B). As a result, many mutually contradicting topologies have been proposed based not only on the morphology and mtDNA/rRNA fragments but also on multigene datasets (Lawrence et al., 2011; Kundrata et al., 2017; McKenna et al., 2015; Zhang et al., 2018; Cai et al., 2022). Hayashi et al. (2024) reported an unstable backbone, poorly supported deep splits, and numerous very short branches using UCE data. Unlike earlier studies (McKenna et al., 2019; Hayashi et al., 2024; Zhang et al., 2018), we analysed 13 families (Figures 2C, 4, 5), adding Lutrochidae, Protelmidae, and Cneoglossidae to multigene analyses for the first time (Cladotomidae *sensu* Lawrence et al., 2024 are missing, and therefore not discussed below). Additionally, we sequenced a much larger volume of data for each terminal than Hayashi et al. (2024) using the UCE approach and Zhang et al. (2018) using PCR-amplified genes. Despite the effort, some controversies persist, especially regarding the position of Elmidae. Although Byrrhoidea has fewer species than several elateroid families, it exhibits greater morphological disparity and a wider range of life strategies than those families. Therefore, future studies should be based on expanded sampling, especially by including ancient missing lineages such as Cladotomidae, additional ptilodactylids to confirm the sister relationship with Cneoglossidae (Figure 6; McKenna et al., 2015), and Psephenidae: Eubrianacinae.

We propose the early split of Byrrhidae followed by the clade of three families that can be designated as a dryopoid clade (Dryopidae, Protelmidae, and Lutrochidae). The similar position of these two early-splitting branches was previously suggested by Zhang et al. (2018), but only Byrrhidae and Dryopidae were sampled, and the statistical support was low. The topology was rejected by other recent studies (Figures 2D, S1). Further, we identified the Eulichadidae + Callirhipidae clade, as in some earlier analyses (McKenna et al., 2019; Cai et al., 2022). Most recent studies have suggested that the Eulichadidae + Callirhipidae clade is deeply rooted and congruent with most of the trees obtained by our analyses. Still, this topology is not supported by our coalescent analysis at the NT level (Figure 4A). We confirm, with little reservation due to some discordance at the NT-level coalescent analyses, the relationships of Cneoglossidae + Ptilodactylidae (McKenna et al., 2015) (Figures 2B, 4, 5) and their position as the subsequent serial split as recovered by some analyses that lacked Cneoglossidae (Hayashi et al., 2024; McKenna et al., 2019; but not Zhang et al., 2018). The terminal clade contains five families (Heteroceridae, Limnichidae, Chelonariidae, Psephenidae, and Elmidae; supported by AA-level analyses and ML NT analysis). The clade was repeatedly inferred from Zhang’s dataset (Zhang et al., 2018; McKenna et al., 2019; Cai et al., 2022) but was compromised by the presence of Ptilodactylidae and the paraphyly of Psephenidae under some settings (Hayashi et al., 2024). Although the clade could be potentially designated as a psephenoid group, it does not correspond with the earlier morphology-based Psephenoidea *sensu* Lawrence, 1988 (Psephenidae, Cneoglossidae, Chelonariidae, Ptilodactylidae, Eulichadidae, and Callirhipidae). The position of Elmidae remains contentious and would require further data to be robustly resolved.

Elateroidea stands out as the most diverse superfamily within Elateriformia, both in terms of species count and morphological disparity (*>*31,000 spp.; Figures 1, S2). The principal phylogenetic backbone was already recovered by the earliest molecular studies (Bocakova et al., 2007; Sagegami-Oba et al., 2007), challenging the morphology-based monophyly of cantharoids and, eventually, elateroids (Crowson, 1972; Lawrence, 1988; Lawrence & Newton, 1995; Lawrence et al., 2011). While our preferred topology (Figure 6) corroborates some recently proposed relationships (Kundrata et al., 2014; Bocak et al., 2016; McKenna et al., 2015, 2019; Zhang et al., 2018; Kusy et al., 2018a, b), it notably diverges from some recent studies that questioned the monophyly of Elateridae, sparking discussion of significant repercussions for family concepts (Cai et al., 2022; Douglas et al., 2021). Our analyses indicate that Elateroidea comprises three distinct clades: (1) the artematopodid clade, consisting of Artematopodidae and Omethidae, including Telegeusinae; (2) the eucnemid-cerophytid clade, housing Eucnemidae and the Throscidae + Jurasaidae + Cerophytidae clade (the TJC clade *sensu* Grebennikov et al., 2026), with Brachypsectridae as their sister or as a deeper split preceding the eucnemid-cerophytid clade; (3) the crown Elateroidea, with two sequential splits, Lycidae + Iberobaeniidae and Cantharidae and the elaterid-lampyroid clade marking the largest and latest radiation within Elateroidea. The latter clade comprises the monophyletic Elateridae and their sister, the lampyroid clade (Sinopyrophoridae, Phengodidae, Rhagophthalmidae, and Lampyridae; Figure 6), as suggested by earlier phylogenomic analyses (Kusy et al., 2021). These major lineages are well-supported, except for the position of Brachypsectridae, an ancient family with a wide distribution but low generic and species diversity (Figures 4, 5C, S15–S32; Costa et al., 2006).

Relationships within elateroid families would primarily require additional sampling to address these identified inconsistencies or mitigate the incompleteness of our dataset (Table S1). For example, the paraphyly of Ototretinae (Figure 6, S15–28) and the relationships among some lampyrid subfamilies, i.e., Psilocladinae, Amydetinae, and Cheguevariinae, need further investigation. Some alternative topologies were recovered within the monophyletic Elateridae (Motyka et al., 2026a). Further improvement will be achieved by including some of the rare species-poor and probably ancient lineages of Eucnemidae (Perothopinae, Phyllocerinae, Palaeoxeninae; Lawrence et al., 2007). Sequencing additional Brachypsectridae might help resolve the discordance over its position. Similarly, expanded sampling of Cantharidae might provide additional information on the Dysmorphocerinae and Heteromastiginae grade. Badmaateridae was discovered after we finished analysing our dataset (Grebennikov et al., 2026), and including it in future expanded analyses would be beneficial.

### Addressing incongruence in phylogenomics

Although phylogenomic datasets are expected to resolve contentious relationships (Misof et al., 2014; McKenna et al., 2019; Stiller et al., 2024; Koch et al., 2023; Van Damme et al., 2022), certain challenging nodes continue to yield contradictory results despite extensive hypothesis testing and increasingly large datasets (Liu et al., 2024; Steenwyk et al., 2023; Li et al., 2021c). Within Elateriformia, several areas of the backbone phylogeny have been difficult to resolve, including the monophyly of Elateridae and Byrrhoidea, the placement of Buprestoidea and Brachypsectridae, internal relationships within Byrrhoidea, and the contentious position of Lycidae + Iberobaeniidae and Cantharidae. Our analyses recovered monophyly of Elateridae and Byrrhoidea, resolved the position of Buprestoidea, and clarified the relationships of Lycidae + Iberobaeniidae and Cantharidae to the remaining Elateroidea. However, the placement of Brachypsectridae and the internal relationships of Byrrhoidea (particularly the positions of Elmidae and Ptilodactylidae + Cneoglossidae) remain uncertain.

The extremely short internodes characteristic of the Elateriformia backbone, with multiple nodes fulfilling the anomaly zone scenario in AA level analyses (Figures 5A, S31; Degnan & Rosenberg, 2006; Cloutier et al., 2019; Linkem et al., 2016), combined with high gene tree discordance at some nodes (Figures 4, 5B), strongly suggest rapid radiations concentrated in three distinct pulses dated to the Late Triassic (a diversification pulse within ∼220–200 Mya period), the Jurassic (~200–140 Mya), and the Early Cretaceous ( ∼140–100 Mya) (Figures S53, S54). The pattern of episodic diversification pulses, in which multiple lineages originate in relatively brief windows, poses substantial challenges for phylogenetic inference. Such evolutionary scenarios present analytical challenges, as traditional phylogenetic methods may fail to capture the limited signal deposited during brief cladogenesis events (Rokas & Carroll, 2005; Philippe et al., 2005, 2011).

These signal-based challenges are particularly acute in Elateriformia by a second, distinct problem. Many ancient lineages that are critical for resolving deep backbone relationships are often species-poor and difficult to sample adequately. For example, the byrrhoid families that diverged in the Jurassic comprise only ∼4,300 extant species across 14 families, with some represented by fewer than a dozen modern species. Similarly, the eucnemid-TJC clade (Eucnemidae, Throscidae, Jurasaidae, Cerophytidae) diversified in the Late Triassic but now comprises only ∼1,800 described species, of which ∼1,500 belong to just three Eucnemidae subfamilies, leaving the other three families and remaining Eucnemidae subfamilies extremely species-poor, with much of their historical diversity known only from fossils. In contrast, diversified clades such as Lycidae (∼ 4,250 species) and Elateridae (*>*14,600 species) contain enormous species diversity, classified into subfamilies that originated during the Cretaceous-Paleogene. Still, their signal is concentrated in more recent divergences rather than in the oldest backbone nodes, leaving the resolution of deep splits disproportionately dependent on the sparse representation of ancient, species-poor lineages.

Systematic error in large, concatenated datasets can be exacerbated when major model assumptions, such as base compositional stationarity, are violated (Philippe et al., 2005; Kapli et al., 2020), and gene-tree estimation error is known to mislead summary coalescent approaches (Degnan & Rosenberg, 2006). We tried to address these limitations through diverse analytical strategies and the depth of analyses. These challenges are often amplified by missing data in poorly sampled clades (Rokas & Carroll, 2005; Kjer et al., 2016) and likely require additional taxonomic sampling to achieve better resolution. Although our analyses provide the first phylogenomic data for many groups, there is undoubtedly room to improve the dataset.

Sophisticated analytical approaches, such as CATGTR, promise to account for compositional heterogeneity. However, their practical implementation often creates more problems than solutions. The CATGTR reanalysis of the Zhang et al. dataset was time-consuming, despite a relatively small dataset size, and produced a tree that challenged several long-held relationships (Cai et al., 2022; Zhang et al., 2018). Our analyses reject their principal conclusions about Elateriformia: paraphyletic Byrrhoidea, Byrrhidae + Buprestidae, lampyroids + Elaterinae, Lissominae + Lycidae, the Cantharidae + Agrypninae + Denticollinae clade, and the late origin of bioluminescence linked to visual-hunting predators like stem birds. Recent studies (Whelan & Halanych, 2017; Boudinot & Lieberman, 2025) have shown that CAT-GTR tends to overestimate the number of substitution categories, particularly in larger datasets, with the number of categories correlating more with dataset size than actual biological heterogeneity. The practice of reporting unconverged CATGTR analyses is statistically invalid and cannot be rationalized under assumptions of model performance. While we do not wish to imply that complex models are inherently problematic, complexity should not be confused with accuracy or biological realism, and any new substitution model should be thoroughly tested before wide-ranging conclusions about evolution are made (Boudinot et al., 2023a).

Comparisons of NT and AA datasets revealed significant differences in phylogenetic resolution (Figures 4, 5C, S15–32). NT-based analyses, using both ML and coalescent approaches, consistently recovered more robust and stable topologies with monophyletic groupings across Elateriformia lineages. In contrast, AA datasets lacked sufficient phylogenetic signal in several key groups, leading to paraphyly or polyphyly, particularly within Elateroidea (Figure 4). The superior performance of NT data likely stems from the capture of additional mutational variation that provides critical phylogenetic signal for resolving the extremely short internodes characteristic of ancient, rapid radiations in this group. We acknowledge that the utility of this signal decreases with node depth, as synonymous substitutions become increasingly saturated and susceptible to compositional biases. However, our compositional heterogeneity analyses, including consistently low RCFV values (∼ 0.010–0.015) and SRH tests, support the reliability of the NT signal across the depth of nodes examined here. The importance of moderately fast-evolving genes for resolving elaterid relationships challenges traditional phylogenetic approaches that prioritize slowly evolving characters. Our findings reveal that in Elateriformia, when analysing AA datasets, faster-evolving genes are essential for resolving certain clades, particularly Elateridae monophyly (Figure 4D, E), corroborating the near-equal split in gene-level AA support for competing Elateridae topologies reported by Kusy et al. (2018a). This counterintuitive result probably reflects the nature of molecular evolution in these beetles. Even our ‘fastest’ genes show remarkably low compositional heterogeneity (RCFV ∼0.010–0.015) and minimal sequence divergence (∼ 0.0010 pairwise divergence), while the slowest-evolving genes (RCFV ∼0.005–0.008, ∼0.0005 pairwise divergence) simply lack sufficient variation to resolve rapid diversification events. The narrow range of both RCFV values (0.005– 0.015) and pairwise divergence across our decisive datasets (Figure 4E) suggests that Elateriformia exhibits generally conservative molecular evolution, yet even these subtle differences have profound effects on phylogenetic resolution. The polyphyly of Elateridae in analyses using complete AA datasets or slower-evolving gene subsets likely reflects insufficient phylogenetic signal rather than biological discordance, as NT-based analyses consistently recovered monophyletic Elateridae across all datasets. Traditional approaches that automatically exclude faster-evolving markers or sites may inadvertently discard precisely those genes that capture synapomorphies during brief periods of cladogenesis (Superson & Battistuzzi, 2022). Although gene properties are routinely calculated by current phylogenomic pipelines, critical evaluation of these statistics in light of the specific clade’s evolutionary history remains essential (Young & Gillung, 2020). These results suggest that optimal gene selection strategies must be tailored to each group’s specific evolutionary history rather than following universal rules about evolutionary rates.

Despite our comprehensive sampling, some relationships remain uncertain and await resolution through targeted analyses. Rather than advocating for a single topology based on specific model assumptions, we explicitly present and discuss conflicting phylogenetic hypotheses. This approach provides a more comprehensive understanding of phylogenetic uncertainty and allows for critical evaluation of how different analytical choices affect tree inference. The presentation of alternative topologies, along with supporting evidence and associated model assumptions, provides greater transparency into the current state of knowledge and highlights areas requiring further investigation (Ballesteros et al., 2022; Young & Gillung, 2020; Steenwyk et al., 2023). This is particularly important when dealing with challenging phylogenetic relationships where different analytical approaches yield conflicting results (Liu et al., 2024; Simon et al., 2018). By acknowledging and examining these conflicts rather than favouring a single “best” tree, we can better understand the methodological and biological factors influencing phylogenetic inference and identify promising directions for future research (Li et al., 2021c; Stiller et al., 2024; Ballesteros et al., 2022). Moving forward, resolving the remaining ambiguities in Elateriformia phylogeny will require not just more data, but more thoughtful integration of diverse evidence types (Schultz et al., 2023; Steenwyk & King, 2024) and explicit modelling of the heterogeneous evolutionary processes that generated present-day diversity. Recent methodological advances have enabled the acquisition of chromosome-level genomes across diverse lineages (Darwin Tree of Life Consortium, ERGA). Therefore, genomic structural information may offer solutions for some of the conflicting deep splits in the Elateriformia tree, as it has been successfully applied in other phylogenomic studies (Steenwyk et al., 2023; Parey et al., 2023; Mirarab et al., 2024). This whole-evidence approach, uniting phylogenomics with comparisons of genome architecture, promises to illuminate the detailed topology of the tree of life.

### Divergence times and origins of neotenic modifications

Unlike the earliest dating analyses (Hunt et al., 2007; McKenna et al., 2015; Toussaint et al., 2017), recent studies rooted Elateriformia much deeper in the polyphagan phylogeny (Zhang et al., 2018; Kusy et al., 2018b; McKenna et al., 2019). The growing evidence that Polyphaga is the sister lineage to the other three suborders (McKenna et al., 2019; Beutel et al., 2024) suggests that Elateriformia is an ancient lineage. Although the early origin of Elateriformia within Polyphaga had not been formally recognized until recently, this finding is not surprising considering that some groups initially placed in Elateriformia are now widely accepted as the earliest polyphagan groups (Scirtiformia, Clambiformia, Rhinorhipiformia) (Hunt et al., 2007; Kusy et al., 2018b; McKenna et al., 2015, 2019; Zhang et al., 2018). Additionally, the rapidly accumulating information on extinct groups and the critical re-evaluation of earliest elateriform or elateroid records call for an updated, more precise calibration of Elateriformia phylogeny (Toussaint et al., 2017; McKenna et al., 2015; Yu et al., 2019).

Although we suggest an early origin of many groups (Figures 6, 7, S41–S52), the critical splits either confirm proposals by Toussaint et al. (2017) or suggest slightly older ages than those recovered by Zhang et al. (2018), McKenna et al. (2019), and Cai et al. (2022). Our analyses bring further evidence that some earlier analyses may have underestimated the ages of Elateriformia and some constituent lineages (e.g., major superfamilies and families). Possible sources of these differences include topologies that suggested splits of Staphyliniformia and Scarabaeoidea prior to the origin of Elateriformia (McKenna et al., 2015; Toussaint et al., 2017), paraphyly of Byrrhoidea, and polyphyly of Elateridae (McKenna et al., 2019; Cai et al., 2022), along with differences in model selection, limited fossil calibrations, lower data volume, and limited taxonomic sampling. Further, the age estimates for Lampyridae align with those reported by Höhna et al. (2025).

While our molecular estimates place the crown Elateriformia origin in the Permian (280.99–287.13 Mya), and the origin of all four superfamilies to the Late Permian-Early Triassic (Figures 6, 7A, S53), no Permian polyphagan fossils are currently known (Beutel et al., 2024). The earliest supposedly polyphagan compression fossil is an undescribed specimen from the Middle Triassic Tongchuan entomofauna (Zheng et al., 2018; fossil NIGP162053, ∼238–237 Mya), followed by the Late Triassic *†Elaterophanes vetustus* (Handlirsch, 1906). This substantial gap between molecular age estimates and the fossil record likely reflects taphonomic biases rather than true absence. Early polyphagan and elateriform beetles may have been small and possibly occupied ecological niches that disfavoured preservation, including cryptic habits in ephemeral terrestrial or aquatic microhabitats lacking fine-grained sediments suitable for compression fossils. Aquatic and detritivorous lineages, characteristic of early polyphagan lineages, including Byrrhoidea, would have been particularly underrepresented. The diversification of most families occurred after the Triassic–Jurassic boundary, coinciding with the recovery of land ecosystems following the end-Triassic mass extinction (Figures 6, 7, S53) (Wignall, 2015). The Jurassic marks the emergence of modern elateriform families, Dascillidae, Buprestidae, Artematopodidae, Eucnemidae, Cerophytidae, and Elateridae, in the fossil record, consistent with our molecular estimates. Most Cretaceous fossils represent lineages comparable to extant taxa, indicating that almost all currently recognized families of Elateriformia had originated by the end of the Jurassic (Figures 6, S43–54).

Moreover, the recently described Jurassic and Cretaceous families (Mysteriomorphidae, Cretophengodidae, and *†Anoeuma, incertae sedis* in Elateroidea) have unresolved phylogenetic positions, with some showing ontogenetic modifications. For example, males of *Anoeuma* exhibit numerous characteristics typical of lineages with larviform females. Cretophengodidae, a member of the lampyroid clade, was described from mid-Cretaceous amber (99 Mya) (Li et al., 2021b) when the split between related Phengodidae and Rhagophthalmidae was estimated only slightly earlier (Cai et al., 2022), at a similar time (McKenna et al., 2015), or much later (Zhang et al., 2018). Now, the origin of these families is estimated much earlier, at ∼160 Mya. The definition of Cretophengodidae is based solely on morphological disparity, which tells us little about relationships among ontogenetically modified groups (Lawrence et al., 2011).

It is worth noting that Jurasaidae likely represents a modified lineage of Cerophytidae, as certain cerophytid fossils (Yu et al., 2019) predate the split between Jurasaidae and extant Cerophytidae (Fig. 6; Grebennikov et al., 2026). The family was erected based on four-fragment analysis, and the timeframe of its origin has not been considered when the rank was proposed (Rosa et al., 2020).

### Morphology versus molecular phylogeny

Morphology- and DNA-based phylogenetic hypotheses of Elateriformia are irreconcilable (Figures 2, S1; Lawrence, 1988; Crowson, 1971, 1972; Branham & Wenzel, 2003; Lawrence et al., 2011; Muona, 1995). We suggest that discordance may be influenced by the highly diverse ecology, specific habitat adaptations, variable flight capability, and various anti-predatory strategies. These factors affect morphology and inevitably lead to the evolution of homoplastic characters, as seen in other beetles, e.g., the concepts of Hydradephaga and Geadephaga (Lawrence et al., 2011; McKenna et al., 2019; Shull et al., 2001). Additionally, some structures may be absent or simplified due to miniaturization, as observed in Badmaateridae and Eucnemidae: Anischiinae (Grebennikov et al., 2026; Grebennikov & Beutel, 2002; Polilov et al., 2019). Elateroidea is widely known for soft-bodiedness and ontogenetic modifications. These phenomena initially obscured morphology-based relationships of other beetles (*Micromalthinus, Thylodrias*, soft-bodied cleroids, i.e., ‘Malacodermata’, etc.; Crowson, 1981). However, most soft-bodied and ontogenetically modified beetles have been identified in Elateroidea (Figures 1, 6, 8).

### Soft-bodiedness and homoplasy

The convergent evolution of morphological traits associated with soft-bodiedness was initially unrecognized because morphological characters supporting the relationships of soft-bodied groups with their true relatives were absent (Lawrence et al., 2011). As a result, superficial morphological similarity has been reflected in the Linnean classification, which contained the superfamilies Elateroidea and Cantharoidea (Branham & Wenzel, 2003; Muona, 1995). The latter was defined by traits resulting from a hypothesized single loss of sclerotization and included Cantharidae, Lycidae, lampyroids, and lineages now placed in Elateridae or classified as close relatives of Artematopodidae (Crowson, 1955, 1972; Lawrence & Newton, 1982). Alternatively, the morphology-based analyses recovered the cantharoid clade and the grade of clicking elateroids (Lawrence, 1988; Lawrence et al., 1995, 2011).

Molecular analyses, including the present study, have clearly shown that soft-bodiedness has evolved multiple times. Therefore, we must accept that the loss of sclerotization and, in some elateroids, the loss of the clicking mechanism have a profound impact on multiple morphological characters, and that homoplasy misleads morphological analyses (Figures 1, 6, 8) (Gould, 1977, 2000; Bocakova et al., 2007; Sagegami-Oba et al., 2007; Motyka et al., 2023b; Kusy et al., 2018a, 2019; etc.). The interpretation of morphological phylogenetic signal is complicated by the diversity of these modifications across elateriform groups, with some only slightly modified (Brachypsectridae, Elateridae: *Plastocerus*, Elaterinae: Cebrionini) and others fully soft-bodied (e.g., Cantharidae, Lycidae, and most lampyroids) (Crowson, 1972; Lawrence et al., 2011; Kusy et al., 2018a; Bocak et al., 2008).

Our sampling includes twelve terminals or clades that were previously placed in Cantharoidea, i.e., in the present sense, within the arbitrary assemblage of soft-bodied elateroids, as recognized by various authors since Crowson’s natural classification of beetle families (Crowson, 1955, 1981; Lawrence & Newton, 1982, 1995; Lawrence et al., 2011). These include *Anorus*, now placed in Dascillidae: Karumiinae (Johnston & Gimmel, 2020); Paulusiellini and *Analestesa*, now in Elateridae, earlier placed in Karumiidae (Kusy et al., 2023); Cneoglossidae (now Byrrhoidea) (Lawrence & Newton, 1995); Omethidae, incl. earlier Telegeusidae (now a sister to Artematopodidae) (Bocakova et al., 2007); Brachypsectridae (McKenna et al., 2019); Lycidae, Cantharidae, lampyroids (Rhagophthalmidae, Phengodidae, and Lampyridae; all Elateroidea) (Kusy et al., 2021); and Drilidae, Plastoceridae, and Omalisidae (the latter three now placed within Elateridae; Kusy et al., 2018a; Motyka et al., 2026a). We can add to these taxa the recently described jurasaids with the cantharoid-like morphology of males (a sister to Cerophytidae or their part, see above; Figures 4, 6; Rosa et al., 2020), and separate branches representing Cebrionini and *Octinodes* (Elateridae, cebrionids were given the family and later subfamily rank) (Kundrata & Bocak, 2011; Motyka et al., 2026a). These raise the number of soft-bodied lineages in the current dataset to 15. Additionally, we may consider, here unsampled, taxa as Pleonominae, an incompletely sclerotized and sexually dimorphic group of click beetles with uncertain position, earlier Nyctorini, now either *Nyctor* as a valid genus or a synonym of *Cardiophorus* (Elateridae; Cate, 2007), and *Podabrocephalus*, displaying some traits that placed it as a sister to the cantharoid clade in Lawrence et al.’s tree (earlier Podabrocephalidae, now a synonym of Ptilodactylinae, Byrrhoidea) (Kundrata et al., 2019; Lawrence et al., 2011). Thirteen of these taxa are nested within fully sclerotized groups, and our analysis suggests multiple origins – no reversal scenario (Dollo’s law of irreversibility; Gould, 1977). The same scenario is supported by the presumed position of *Nyctor, Pleonomus*, and *Podabrocephalus* within Elateridae and Ptilodactylidae, respectively.

Cantharidae and Lycidae contain only soft-bodied species and were recovered as serial splits preceding the Elateridae-lampyroid clade (Figures 4, 6). The analysis of ancestral traits suggested a well-sclerotized ancestor (p=88 for both families). These splits retain a small probability of soft-bodiedness origin and reversal. The application of parsimony makes possible two scenarios: (1) a soft-bodied most recent common ancestor (mrca) of Lycidae, Cantharidae, Elateridae, and lampyroids and subsequent reappearance of complete sclerotization in the mrca of Elateridae + lampyroids or, alternatively, (2) the independent origins of soft-bodiedness in Lycidae and Cantharidae (p = 12 in our analysis). Although we do not have hard data to estimate the loss and regain probabilities, scenario 1 requires not only the reappearance of full sclerotization but also the reappearance of the finely tuned clicking mechanism, which depends on the precise fit of coevolving parts of the pro- and mesothorax and muscle apparatus (Bolmin et al., 2022; Johansson et al., 2012; Ribak & Weihs, 2011). Therefore, we consider the re-evolution of the complex clicking mechanism statistically improbable (Gould, 2002). The morphological uniformity of clicking elateroids may have contributed to the repeated recovery of these families in morphology-based analyses (Elateridae, Eucnemidae, Cerophytidae, and Throscidae) (Lawrence et al., 2011; Muona, 1995).

### Ontogenetic reprogramming in Elateroidea

Elateroidea contains not only soft-bodied beetles but also phenotypically highly variable females retaining larval traits into adulthood (Figures 1, 6 left, 8, S54; Table S9; Text S1). The general appearance of well-sclerotized elateroids resembles Dascilloidea, Buprestoidea, and the majority of Byrrhoidea. These phenotypes were recovered as ancestral within Elateroidea (Figures 6, S54) and are represented by Artematopodidae (non-clicking), Badmaateridae (possibly clicking), Eucnemidae, Throscidae, and Cerophytidae (all species sclerotized, resembling the clicking elateroids in general appearance. Eucnemidae are considered clicking; however, some species have a greatly reduced pro-mesosternal clicking mechanism, including Anischiinae and some Melasinae; no behavioural study has demonstrated that these species do not click. All species of Cerophytidae and Throscidae possess the well-developed clicking apparatus, although the strength and effectiveness of the click appear to vary. As in the case of Eucnemidae, no study has demonstrated that they do not click). These families were recovered as deeply rooted (Figures 4, 6, S54). The clicking Elateridae and Sinopyrophoridae are nested within the clade containing Cantharidae, Lycidae, and the lampyroids (Figures 4, 6, 8, S15–S32). The recovered probabilities in the ancestral traits analysis might be affected by oversampling of modified forms. The opposite phenotypic extreme is represented by fully larviform females known in net-winged beetles (Bocak et al., 2008; Wong, 1996; Makarov & Kazantsev, 2022; Masek et al., 2015; etc.). In between, there are almost all possible transitional phenotypes, represented by various degrees of retention of larval characters in the adult females. Morphological modifications of various body parts are often unlinked in different taxa. For example, most lycid neotenic males, including those unrelated, have hypognathous heads and miniaturized mouthparts except for *Platerodrilus* (Kusy et al., 2019; Masek et al., 2015), and this difference cannot be connected to either relationships or body miniaturization (Bocak et al., 2008; Kusy et al., 2019) (Figure 8).

We defined the character state reconstruction to include the well-sclerotized phenotype (state 0), soft-bodiedness (state 1), and four levels of ontogenetic modification (states 2, 3, 4, 5). These states are primarily based on the gradual loss of elytra and wings and modifications of the thorax (see Figure 8). However, the elytra/wings- and thorax-based definitions are correlated with loose connections between abdominal segments, shortened appendages, body miniaturization, and sometimes modifications of the cranium and mouthparts in males (Figures 6, 8). Apparently, a smooth gradient exists between weakly modified lineages that have brachelytrous, wingless females with adult-like thoracic segments (Elateridae: Omalisinae, some Lampyridae; Bocek et al., 2018) and fully larviform, large-bodied females in Lycidae (Figures 1, 8; Makarov & Kazantsev, 2022; Masek et al., 2015). Females are always more affected by ontogenetic reprogramming than conspecific males (females: states 2–5; Figures 6 left, 8). Analogous modifications sometimes lead to brachelytrous, weakly sclerotized miniaturized males that resemble females of weakly modified species (Figure 8, state 2), while the closely related taxa have large-bodied males (Bocak et al., 2008; Kazantsev, 1999, 2005; Bocak, Grebennikov, & Masek, 2013; Bocak, Grebennikov, & Sklenarova, 2014; Ferreira et al., 2023).

With changes categorized, we can discuss the origins of various modifications in the context of the preferred time-calibrated topology and based on the character-state reconstruction (Figures 6, 7, S54). The well-sclerotized body is an ancestral trait of Elateriformia. The soft-bodiedness originated multiple times and cannot be used as an apomorphy defining the ‘cantharoids’. Neotenics (states 2–5) originate within well-sclerotized, eventually clicking groups (state 0) as well as within soft-bodied groups (state 1). In contrast to the gradual nature of the changes, we never found less modified forms older and more advanced forms younger, or progressively modified taxa nested within clades consisting of less modified taxa (no shifts between characters, 2→ 3 → 4 → 5). Instead, the changes are often taxon-specific: the same phenotype is retained by all descendants after a shift to neoteny, and no further transformation is detected. This scenario is documented by several independently originating lineages: *Anorus* (Dascillidae: Karumiinae, state 3), Jurasaidae (state 4; Rosa et al., 2020), Drilini (Elateridae: Agrypninae; state 4; Kundrata & Bocak, 2019), all neotenic Lycidae (two origins; state 5; Bocak et al., 2008; Makarov & Kazantsev, 2022; Masek et al., 2014, 2015), Omalisinae (Elateridae, type 2; Bocek et al., 2018), the Rhagophthalmidae + Phengodidae clade (state 5, with rhagophthalmid and phengodid females having differently modified antennae), and Cebrionini and *Octinodes* (separate origins, both Elateridae: Elaterinae; state 2). Similar uniformity might be present in taxa with unknown females (Omethidae: Telegeusinae, Phengodidae: Cydistinae, Iberobaeniidae, Lycidae: Dexorinae, Ateliinae, *Cautires apterus* Bocak et al., 2013 (Metriorrhynchinae: Metriorrhynchini), and Paulusiellini (Elateridae: Hemiopinae). The observed patterns may suggest that modifications occur suddenly and are not driven by natural selection for further changes. Bocak et al. (2008) hypothesized that ecological traits arising from developmental changes are tolerated, whereas selective pressures shape reproductive features, such as large-bodied females, and eventually miniaturized males. As in the case of soft-bodiedness, a reversal to full metamorphosis has never been proposed within any clade that includes the modified taxa (Figures 6, 8, S54).

Fireflies differ from these cases and exhibit much higher phenotypic plasticity. Unfortunately, the sampling of Elateriformia phylogeny is inevitably insufficient for the reconstruction of ontogenetic modifications in fireflies. However, we can provide basic information about the group’s phenotypic diversity. Some fireflies have both sexes soft-bodied and fully winged: *Drilaster* (Ototretinae), most Luciolinae (state 2 in *Luciola filiformis*), most *Photinus* (Lampyrinae; but state 2 in *P. scintillans*), *Photuris* (Photurinae), and others. States 2–4 are commonly present in Lampyridae (Figure 8; South et al., 2011), with fully larviform females reported in *Lamprigera, Stenocladius*, and *Oculogryphus*; Jeng et al., 2021; Kawashima & Satou, 2004; Yiu & Jeng, 2018). A detailed discussion of the evolution of ontogenetic modifications in this group should be based on much denser sampling, as variability in these modifications has been reported even within genera (e.g., *Diaphanes* (see Cicero, 2008) and *Lampyris noctiluca* and *L. sardiniae*; observation on a breeding colony of authors).

Strepsiptera remains the closest group investigated for the molecular mechanisms of ontogenetic reprogramming (Erezyilmaz et al., 2014). Recently, molecular investigations have begun in Lampyridae, where Catalán et al. (2024) generated novel genomes for two firefly species with different degrees of sexual dimorphism (*Lamprohiza splendidula* and *Luciola italica*) and uncovered sex-biased gene expression patterns, X-chromosome dosage compensation, and higher nucleotide diversity in sex-biased genes, potentially maintained by sexual selection. These findings provide initial insights into the genetic architecture underlying sexually dimorphic traits in fireflies and represent a valuable resource for understanding the evolution of neotenic phenotypes. However, much remains unknown about the molecular mechanisms underlying soft-bodiedness and ontogenetic modifications in Elateriformia more broadly (Cicero, 1988; McMahon & Hayward, 2016). Therefore, a robust phylogeny that identifies the sister groups of modified lineages is crucial for further progress in studies of molecular-level ontogenetic modifications and for comparing transcriptomic changes across phylogeny.

### Bioluminescence

Bioluminescence is another phenotypic characteristic encountered in Elateroidea and widely discussed by evolutionary biologists (Fallon et al., 2018; Oba & Schultz, 2022; He et al., 2024; Carrasco-López et al., 2021; Oba et al., 2020). A single origin of bioluminescence has never been proposed, and only two questions have remained: how distant are bioluminescent lampyroids and elaterids, and how old is bioluminescence? Dissolution of cantharoids (Bocakova et al., 2007; Sagegami-Oba et al., 2007; Hunt et al., 2007) opened new possibilities for mutual relationships among bioluminescent taxa earlier placed to different superfamilies, including the recently proposed placement of lampyroids within Elateridae (Cai et al., 2022; Kundrata et al., 2014; McKenna et al., 2015, 2019; Timmermans et al., 2016; Zhang et al., 2018; Douglas et al., 2021; etc.) (Figures 2, S1). The placement of some bioluminescent lineages has recently been investigated by Kusy et al. (2021) and Motyka et al. (2023a, 2026a). The suggested topologies are confirmed here (Figure 6). Briefly, bioluminescence is limited to the clade consisting of Elateridae and the lampyroid clade (Kusy et al., 2021). The lampyroids include bioluminescent, clicking Sinopyrophoridae, which were previously considered to represent an independent origin of bioluminescence in click beetles (Bi et al., 2019). *Sinopyrophorus* marks the first split in the lampyroid clade that encompasses all soft-bodied bioluminescent elateroids (Lampyridae, Rhagophthalmidae, and Phengodidae). Separate origins of bioluminescence are identified within Elateridae: Pyrophorini and possibly Euplinthini (both surely bioluminescent; relationships of the latter should be confirmed by molecular analyses), *Campyloxenus pyrothorax* (unpublished video proving bioluminescence was reported by Motyka et al., 2023a, but the visible bioluminescence was not observed in the field when the species was recently collected, L. Bocak, personal observation), and *Balgus schnusei* (unconfirmed literature information; Costa, 1984). The concentration of bioluminescent forms in the crown clade of Elateroidea raises the question of whether active light production is driven by preadaptation (Vahtera et al., 2009; Fallon et al., 2018; Oba et al., 2020).

The Jurassic origin of the lampyroid clade and the much younger Cretaceous origin of Pyrophorini bioluminescence indicate that light production evolved independently, possibly under different ecological contexts (Figure 6; Table S10). Larval bioluminescence is considered ancestral in the lampyroid clade, likely serving an aposematic function, while adult bioluminescence for sexual communication evolved subsequently (Oba et al., 2020; Powell et al., 2022). However, the sinopyrophorid adult is bioluminescent, and larvae are unknown. The early origin of bioluminescence refutes the proposed role of bioluminescence as a response to predation pressure from stem birds as visual predators (Cai et al., 2022).

In Pyrophorini (Elateridae), larval bioluminescence may serve both aposematic and prey-luring functions (Redford, 1982), whereas adults use light production for both sexual communication and potentially aposematism, convergent with lampyrids (Stolz et al., 2003). Notably, in contrast with distant relationships, luciferase genes are duplicated in both Lampyridae, Phengodidae, and Pyrophorini, with distinct expression patterns reflecting this functional differentiation (He et al., 2024; Arnoldi et al., 2010; Stolz et al., 2003; Fallon et al., 2018; Oba et al., 2020). Chemical defense, indicating an aposematic function of active light production, evolved around the Cretaceous/Paleogene boundary within the Lampyrinae subfamily (Fig. 6; Zhu et al., 2024; Eisner et al., 1978), suggesting aposematic signaling preceded actual toxicity (but see Hosoe et al., 2014). Neophobia might have played a role in the early phases of the prey-predator interaction.

## Conclusion

Our phylogenomic analysis of Elateriformia reveals a robust backbone for major relationships (Figures 2, 6) and provides an essential framework for future investigations of the genetic mechanisms underlying the parallel evolution of ontogenetic modifications. Based on comprehensive tests, we return to the traditionally held concept of Byrrhoidea *s. l*., the deep rooting of Buprestoidea, the monophyletic Elateridae, and the lampyroid clade, including Sinopyrophoridae (Bouchard et al., 2011; Lawrence & Newton, 1995; Kusy et al., 2021). However, at some nodes, we uncovered a complex network of evolutionary histories rather than a single robustly supported tree. We show that different genomic regions tell slightly different evolutionary stories, particularly in lineages that rapidly radiated in deep time. These include the position of Elmidae, Brachypsectridae, and the radiation of the Eucnemidae-Cerophytidae clade. The preferred evolutionary trajectories have profound implications for understanding and dating the repeated evolution of soft-bodiedness, female neoteny, and bioluminescence. Some evolutionary scenarios are consistently suggested by the current analyses: Elateroidea and, to a lesser extent, Byrrhoidea are prone to the loss of a compact, well-sclerotized body with tightly coadapted body parts. As a result, phenotypically similar soft-bodied forms have evolved in distantly related well-sclerotized lineages from the Jurassic to the present. Female neoteny, similarly, evolved multiple times, but we do not observe gradual evolution. Instead, some taxa are nested among fully sclerotized relatives and exhibit significant modification, including the larviform females. Bioluminescence is limited to the crown clade of Elateroidea, with two origins well documented and an additional two taxa with bioluminescence observed only once.

To fully resolve remaining phylogenomic complexities, future research should embrace this complexity rather than seeking a single ‘true tree.’ We recommend additional sampling of underrepresented and absent families, and ontogenetically modified forms. Dating may be further improved by an objective revision of fossil placements that accounts for morphological convergence. Future integration of chromosome-level genomes with analyses of genomic structural features can contribute to a robust phylogeny of Elateriformia, as demonstrated by studies resolving other ancient radiations (Schultz et al., 2023; Parey et al., 2023; Mirarab et al., 2024; Stiller et al., 2024).

## Authors’ Contributions

L.B. conceived the study; D.K. designed and conducted bioinformatic analyses; D.K., L.B., M.M., G.B., F.C., and E.T.A. collected or supplied UPOL samples; CC, MSC, HE, JH, BO, and MP collected or supplied CSIRO samples; D.K. and M.M. generated UPOL data; AZ, HE, and AS generated CSIRO data; D.K. selected the fossils; D.K. and L.B. wrote the draft; other authors reviewed and edited it. All authors gave final approval for publication and agreed to be held accountable for the work performed therein.

## Acknowledgements

This research was funded by the Czech Science Foundation (GAČ R-22-35327S) for L.B., D.K., and M.M. The data, previously published by L.B., M.M., and D.K. and included in the current dataset, were generated under funding from research projects for L.B. (GAČ R P506/11/1757, 18-14942S). The Leverhulme Trust (F/00696/P), the European Social Fund, and the Ministry of Education of the Czech Republic (CZ.1.07/2.3.00/20.0166, CZ.1.07/2.3.00/30.0004) and the Japanese Society for the Promotion of Science (P02203) supported the LB’s field expeditions. We thank Renata Bílková for assistance with laboratory work. G.B. was supported by Vale Institute of Technology – Vale S/A (Projects R100603.CD.0X; R100603.CT.0X) and Fundação Guamá (0104/2025). Material provided by JH was produced under support from the Ministry of Culture of the Czech Republic (DKRVO 2024-2028/5.I.c, National Museum, 00023272). We are grateful to S. Vaz, M. Macedo, and R. Fernández for donating Jurasaidae specimens, B. Wang for providing information on fossil NIGP162053 (Zheng et al., 2018), L. Mazal, and K. Hirai for providing a specimen of *Cerophytum elateroides* and *C. japonicum*, respectively. We thank M. Bednarik, S. Bybee, V. Kuban, N. Lord, and A. Wild for providing additional valuable specimens.

Field research and export permits were obtained from the Sabah Biodiversity Council (JKNI/MBS.1000-2/2(1 10)); the Department of Environment, Government of Australia (PWS2014-AU-001866); University Los Banos, the Philippines (231/1991); Corporación Nacional Forestal, CONAF, Chile (39/2022); and the Government of Papua New Guinea (sda-2018-daf). We are obliged to landowners for allowing us to collect on private land. Specimens collected in the USA, Europe, and Japan were collected outside protected areas on public land or with permission from landowners.

## Data and Code Availability

Raw sequencing reads generated for this study have been deposited in the NCBI Sequence Read Archive under BioProject accession number PRJNA1420406.

Individual gene alignments, concatenated datasets (both amino acid and nucleotide), partition schemes, phylogenetic trees (ML and coalescent), and configuration files for all analyses are available from the Mendeley Data Repository (https://doi.org/10.17632/by48kddmvf.1). All code and software sources used in our study are listed in the “Methods” section with corresponding citations or links. Custom scripts used for data processing and analyses are available on GitHub (https://github.com/kusydominik/Elateriformia_phylogenomics).

