## Supplementary Material for "Time-calibrated phylogenomics reveals relationships and parallel evolution of neoteny within Elateriformia (Coleoptera) despite pervasive homoplasy"

### Supplementary Tables

Supplementary Table S1. List of taxa used in analyses, indicating data type (transcriptome or genome), specimen preservation (transcriptomes: RNAlater; genomes: fresh material in ethanol or dry museum specimens), and data origin (current study or publicly available data).

Supplementary Table S2. Overview of gene sets of six reference species used for ortholog assessment, including the source, version and number of genes. URLs for downloading the data are given. The number of genes corresponds with OrthoDB 9.1.

Supplementary Table S3. Summary statistics by taxon across data processing stages (raw data, outlier removal, and AliScore filtering) at the amino acid level.

Supplementary Table S4. Summary statistics by taxon across data processing stages (raw data, outlier removal, and AliScore filtering) at the nucleotide level.

Supplementary Table S5. Overview of compiled datasets, statistics and descriptions.

Supplementary Table S6. Overview of selected fossil taxa used for node calibrations, with their corresponding soft minimum and soft maximum age constraints. Detailed fossil placement and justifications are provided in Supplementary Figure S32 and Supplementary Text S1.

Supplementary Table S7. Estimated marginal likelihood of relaxed-clock models with MCMCTree and mcmc3r.

Supplementary Table S8. Comparison of divergence time estimates (Mya) for major Elateriformia lineages between this study and previous studies.

Supplementary Table S9. Divergence time estimates (Mya) for neotenic and putatively neotenic lineages, indicating type of neotenic modification.

Supplementary Table S10. Divergence time estimates (Mya) for the origins of bioluminescent lineages in Elateriformia.

### Supplementary Figures

Supplementary Figure S1. History of phylogenetic relationships among Elateriformia families from selected studies based on morphology and molecular data.

Supplementary Figure S2. Numbers of described species in Elateriformia.

Supplementary Figure S3. Number of genes vs. total number of amino acids A) and nucleotides B) recovered for each of the categories. Three processing stages are shown: Raw (initial Orthograph output), Outliers (after filtering outlier sequences), and Aliscore (after outlier removal and ambiguous alignment filtering with Aliscore).

Supplementary Figure S4. AliStat heat maps showing pairwise alignment completeness for all species included in supermatrices: A) A-4189-AA and B) B-4189-NT.

Supplementary Figure S5. AliStat heat maps showing pairwise alignment completeness for all species included in supermatrices: A) I-1451-AA-Bacoca and B) J-1451-NT-Bacoca.

Supplementary Figure S6. AliStat heat maps showing pairwise alignment completeness for all species included in supermatrices: A) F-4224-NT-raw and B) E-4224-AA-raw.

Supplementary Figure S7. AliStat heat maps showing pairwise alignment completeness for all species included in supermatrices: A) K-3583-NT-SymTest and B) D-1892-AA-MARE.

Supplementary Figure S8. AliStat heat maps showing pairwise alignment completeness for all species included in supermatrices: A) G-962-AA-decisive and B) H-962-NT-decisive.

Supplementary Figure S9. Heat maps calculated with SymTest showing p-values for the pairwise Bowker's tests in supermatrices: A) B-4189-NT all codon positions, B) B-4189-NT first and second codon position, C) B-4189-NT first codon position, D) B-4189-NT second codon position and E) B-4189-NT third codon position.

Supplementary Figure S10. Heat maps calculated with SymTest showing p-values for the pairwise Bowker's tests in supermatrices: A) A-4189-AA, B) I-1451-AA-Bacoca, C) J-1451-NT-Bacoca, D) E-4224-AA-raw and E) F-4224-NT-raw.

Supplementary Figure S11. Heat maps calculated with SymTest showing p-values for the pairwise Bowker's tests in supermatrices: A) K-3583-NT-SymTest, B) D-1892-AA-MARE, C) G-962-AA-decisive and D) H-962-NT-decisive.

Supplementary Figure S12. Comparison of characteristics between datasets 4189 (coral) and 4224 (teal) for amino acid A) and nucleotide B) sequence alignments. Violin plots display distributions of missing data percentage, alignment length, parsimony informative sites, average branch length, average bootstrap support, and number of taxa. Statistical significance determined by Wilcoxon rank-sum tests: *** $p < 0.001$, **** $p < 0.0001$. All metrics showed significant differences between datasets. Grey points represent outliers identified using the IQR method (values beyond Q1 - 1.5×IQR or Q3 + 1.5×IQR).

Supplementary Figure S13. GC content plot for supermatrix B-4189-NT A) first codon position; B) second codon position; C) third codon position and D) all three codon positions.

Supplementary Figure S14. GC content plot for supermatrix F-4224-NT-raw A) first codon position; B) second codon position; C) third codon position and D) all three codon positions.

Supplementary Figure S15. Maximum likelihood trees from IQ-TREE analyses of the following supermatrices: A) A-4189-AA supermatrix partitioned by genes; B) B-4189-NT supermatrix partitioned by genes with only GTR model considered in model selection step and C) B-4189-NT supermatrix partitioned by genes with all available models considered in model selection step -m MFP. The depicted branch support values are based on SH-aLRT and ultrafast bootstrap (3000 replicates each).

Supplementary Figure S16. Maximum likelihood trees from IQ-TREE analyses of the following supermatrices: A) A-4189-AA supermatrix analysed as single partition and B) B-4189-NT supermatrix analysed as single partition with only GTR model considered in model selection step. The depicted branch support values are based on SH-aLRT and ultrafast bootstrap (3000 replicates each).

Supplementary Figure S17. Maximum likelihood trees from IQ-TREE analyses of the supermatrix C-4189-NT12 A) partitioned by genes and B) analysed as a single partition. Only the GTR model was considered in the model selection step. The depicted branch support values are based on SH-aLRT and ultrafast bootstrap (3000 replicates each).

Supplementary Figure S18. Maximum likelihood trees from IQ-TREE analyses of the following supermatrices: A) D-1892-AA-MARE supermatrix partitioned by genes and B) K-3583-NT-SymTest supermatrix partitioned by genes with only GTR model considered in model selection step. The depicted branch support values are based on SH-aLRT and ultrafast bootstrap (3000 replicates each).

Supplementary Figure S19. Maximum likelihood trees from IQ-TREE analyses of the following supermatrices: A) I-1451-AA-Bacoca supermatrix analysed as single partition; B) I-1451-AA-Bacoca supermatrix partitioned by genes; C) I-1451-AA-Bacoca supermatrix with optimalized partitioning scheme options: –merge –rclusterf 10 and –rcluster-max set to default; D) I-1451-AA-Bacoca supermatrix analysed under heterogeneous posterior mean site frequency (PMSF) model.The depicted branch support values are based on SH-aLRT and ultrafast bootstrap (3000 replicates each).

Supplementary Figure S20. Maximum likelihood trees from IQ-TREE analyses of the following supermatrices: A) J-1451-NT-Bacoca supermatrix partitioned by genes; B) J-1451-NT-Bacoca supermatrix analysed as single partition; C) J-1451-NT-Bacoca supermatrix with optimalized partitioning scheme options: –merge –rclusterf 10 and –rcluster-max set to default. The depicted branch support values are based on SH-aLRT and ultrafast bootstrap (3000 replicates each).

Supplementary Figure S21. Maximum likelihood trees from IQ-TREE analyses of the following supermatrices: A) E-4224-AA-raw supermatrix partitioned by genes; B) F-4224-NT-raw supermatrix partitioned by genes with only GTR model considered in model selection step. The depicted branch support values are based on SH-aLRT and ultrafast bootstrap (3000 replicates each).

Supplementary Figure S22. Coalescent species trees inferred by ASTRAL from individual gene trees. Maximum likelihood gene trees were constructed using IQ-TREE with datasets A) A-4189-AA; B) B-4189-NT and C) K-3583-NT-SymTest. Branches with ultra-fast bootstrap support ≤50 were collapsed prior to coalescent analysis. Branch lengths are shown in coalescent units, and node support values represent local posterior probabilities.

Supplementary Figure S23. Coalescent species trees inferred by ASTRAL from individual gene trees. Maximum likelihood gene trees were constructed using IQ-TREE with datasets A), B) A-4189-AA; C), D) B-4189-NT and E), F) K-3583-NT-SymTest. Branches with ultra-fast bootstrap support ≤50 were collapsed prior to coalescent analysis. Branch lengths are in coalescent units. Node support is shown as quartet frequencies (q1: main topology; q2, q3: alternative topologies) displayed as percentages in numerical or pie chart format.

Supplementary Figure S24. Coalescent species trees inferred by ASTRAL from subsets of individual gene trees. Maximum likelihood gene trees were constructed using IQ-TREE with A), B) 200 slowest-evolving genes from dataset G-962-AA-decisive; C), D) 200 fastest-evolving genes from dataset G-962-AA-decisive; E), F) all genes from dataset G-962-AA-decisive and G), H) all genes from dataset H-962-NT-decisive. Branches with ultra-fast bootstrap support ≤50 were collapsed prior to coalescent analysis. Branch lengths are in coalescent units. Node support is shown as quartet frequencies (q1: main topology; q2, q3: alternative topologies) displayed as percentages in numerical or pie chart format.

Supplementary Figure S25. Coalescent species trees inferred by wASTRAL from individual gene trees. Maximum likelihood gene trees were constructed using IQ-TREE with datasets A) F-4224-NT-raw; B) B-4189-NT; C) K-3583-NT-SymTest; D) E-4224-AA-raw and E) A-4189-AA. Gene trees were used without collapsing poorly supported branches. Branch lengths are shown in coalescent units, and node support values represent local posterior probabilities.

Supplementary Figure S26. Coalescent species trees inferred by wASTRAL from individual gene trees. Maximum likelihood gene trees were constructed using IQ-TREE with datasets A), B) E-4224-AA-raw and C), D) A-4189-AA. Gene trees were used without collapsing poorly supported branches. Branch lengths are in coalescent units. Node support is shown as quartet frequencies (q1: main topology; q2, q3: alternative topologies) displayed as percentages in numerical or pie chart format.

Supplementary Figure S27. Coalescent species trees inferred by wASTRAL from individual gene trees. Maximum likelihood gene trees were constructed using IQ-TREE with datasets A), B) F-4224-NT-raw; C), D) B-4189-NT and E), F) K-3583-NT-SymTest. Gene trees were used without collapsing poorly supported branches. Branch lengths are in coalescent units. Node support is shown as quartet frequencies (q1: main topology; q2, q3: alternative topologies) displayed as percentages in numerical or pie chart format.

Supplementary Figure S28. Coalescent species trees inferred by wASTRAL from subsets of individual gene trees. Maximum likelihood gene trees were constructed using IQ-TREE with A), B) all genes from dataset G-962-AA-decisive; C), D) 200 fastest-evolving genes from dataset G-962-AA-decisive; E), F) 200 slowest-evolving genes from dataset G-962-AA-decisive and G), H) all genes from dataset H-962-NT-decisive. Gene trees were used without collapsing poorly supported branches. Branch lengths are in coalescent units. Node support is shown as quartet frequencies (q1: main topology; q2, q3: alternative topologies) displayed as percentages in numerical or pie chart format.

Supplementary Figure S29. DiscoVista relative frequency analyses of datasets: A) A-4189-AA branches with ultra-fast bootstrap supports ≤50 were collapsed in individual gene trees; B) E-4224-AA-raw gene trees were used without collapsing poorly supported branches.

Supplementary Figure S30. DiscoVista relative frequency analyses of datasets: A) B-4189-NT branches with ultra-fast bootstrap supports ≤50 were collapsed in individual gene trees; B) F-4224-NT-raw gene trees were used without collapsing poorly supported branches.

Supplementary Figure S31. Pairs of short successive internal branches create conditions predicted by anomaly zone theory. wASTRAL species tree topologies for datasets A) A-4189-AA and B) E-4224-AA-raw show internal branch lengths in coalescent units, derived from individual maximum-likelihood gene tree reconstructions by IQTree. Branch length values for all consecutive branch pairs (x and y) are provided in the data table C), with values a(x) computed following Degnan and Rosenberg (2006) Equation 4. Gene tree incongruence is predicted under anomaly zone conditions when $y < a(x)$. Branch pairs satisfying this anomaly zone condition are marked in the topologies by red arrows.

Supplementary Figure S32. Results of Four cluster likelihood mapping (FcLM) analyses for a selection of phylogenetic hypotheses applied at the amino acid and nucleotide level for datasets: A) A-4189-AA and B) B-4189-NT.

Supplementary Figure S33. A) The cladogram with preferred topology (Fig. 5) with fossil calibration points (marked by arrows) used for molecular dating. Pink ovals and dotted lines indicate soft minimum and maximum age constraints, respectively. Numbered fossils (1–14) are listed with their placements; see Supplementary Text S1 for detailed fossil justifications. B) Same tree as (A) but with node numbers displayed.

Supplementary Figure S34. Convergence assessment of four independent MCMCTREE runs for dataset A at the amino acid level under the autocorrelated rates (AR) clock model. Scatter plots compare posterior mean ages and 95% credibility intervals (CI) between pairs of independent runs. Each plot shows one run on the x-axis versus another on the y-axis. High $R^2$ values (>0.999) across all pairwise comparisons indicate convergence among chains.

Supplementary Figure S35. Convergence assessment of four independent MCMCTREE runs for dataset A at the amino acid level under the independent rates (IR) clock model. Scatter plots compare posterior mean ages and 95% credibility intervals (CI) between pairs of independent runs. Each plot shows one run on the x-axis versus another on the y-axis. High $R^2$ values (>0.999) across all pairwise comparisons indicate convergence among chains.

Supplementary Figure S36. Convergence assessment of four independent MCMCTREE runs for dataset A at the nucleotide level under the autocorrelated rates (AR) clock model. Scatter plots compare posterior mean ages and 95% credibility intervals (CI) between pairs of independent runs. Each plot shows one run on the x-axis versus another on the y-axis. High $R^2$ values (>0.999) across all pairwise comparisons indicate convergence among chains.

Supplementary Figure S37. Convergence assessment of four independent MCMCTREE runs for dataset A at the nucleotide level under the independent rates (IR) clock model. Scatter plots compare posterior mean ages and 95% credibility intervals (CI) between pairs of independent runs. Each plot shows one run on the x-axis versus another on the y-axis. High $R^2$ values (>0.999) across all pairwise comparisons indicate convergence among chains.

Supplementary Figure S38. Convergence assessment of two independent MCMCTREE runs for dataset B under different data types and clock models. Scatter plots compare posterior mean ages and 95% credibility intervals (CI) between the two runs for: A) amino acid sequences under the autocorrelated rates (AR) clock model, B) amino acid sequences under the independent rates (IR) clock model, C) nucleotide sequences under the autocorrelated rates (AR) clock model, and D) nucleotide sequences under the independent rates (IR) clock model. High $R^2$ values (>0.999) across all comparisons indicate convergence between runs.

Supplementary Figure S39. Convergence assessment comparing merged MCMCTREE runs between datasets A and B under different data types and clock models. Scatter plots compare posterior mean ages and 95% credibility intervals (CI) between dataset A (four merged independent runs) and dataset B (two merged independent runs) for: A) nucleotide sequences under the independent rates (IR) clock model, B) nucleotide sequences under the autocorrelated rates (AR) clock model, C) amino acid sequences under the independent rates (IR) clock model, and D) amino acid sequences under the autocorrelated rates (AR) clock model. High $R^2$ values (>0.998) across all comparisons indicate strong agreement in age estimates between the two datasets despite differences in data composition and number of runs.

Supplementary Figure S40. Infinite-sites plots. Posterior CI width is plotted against mean posterior divergence times, dataset A and B on both NT = nucleotides and AA = amino acid level under AR = autocorrelated rates and IR = independent rates clock models. Red dots represent nodes with fossil calibrations, grey dots represent the rest of nodes. Solid red line: regression through the origin fitted to the red dots. Grey dashed line: regression through the origin fitted to the grey dots. Black dashed line: regression through the origin fitted to all dots.

Supplementary Figure S41. Density plots of age estimates of calibrated nodes across all time tree analyses performed. Datasets A and B on amino acid level, AR = autocorrelated rates and IR = independent rates clock models. Color legend: solid black line = prior age; blue = effective priors; red = posterior Dataset A, autocorrelated rates; yellow = posterior Dataset B, autocorrelated rates; green = posterior Dataset B, independent rates; grey = posterior Dataset A, independent rates.

Supplementary Figure S42. Density plots of age estimates of calibrated nodes across all time tree analyses performed. Datasets A and B on nucleotide level, AR = autocorrelated rates and IR = independent rates clock models. Color legend: solid black line = prior age; blue = effective priors; red = posterior Dataset A, autocorrelated rates; yellow = posterior Dataset B, autocorrelated rates; green = posterior Dataset B, independent rates; grey = posterior Dataset A, independent rates.

Supplementary Figure S43. Divergence time estimation results. Preferred topology analyzed with Dataset A at the amino acid level under autocorrelated rates. A) Blue horizontal bars represent 95% confidence intervals with lower and upper bound ages labeled at bar endpoints. B) Mean node ages given in units of 100 Mya (e.g., 1.2511 = 125.11 million years ago).

Supplementary Figure S44. Divergence time estimation results. Preferred topology analyzed with Dataset B at the amino acid level under autocorrelated rates. A) Blue horizontal bars represent 95% confidence intervals with lower and upper bound ages labeled at bar endpoints. B) Mean node ages given in units of 100 Mya (e.g., 1.2511 = 125.11 million years ago).

Supplementary Figure S45. Divergence time estimation results. Preferred topology analyzed with Dataset A at the amino acid level under independent rates. A) Blue horizontal bars represent 95% confidence intervals with lower and upper bound ages labeled at bar endpoints. B) Mean node ages given in units of 100 Mya (e.g., 1.2511 = 125.11 million years ago).

Supplementary Figure S46. Divergence time estimation results. Preferred topology analyzed with Dataset B at the amino acid level under independent rates. A) Blue horizontal bars represent 95% confidence intervals with lower and upper bound ages labeled at bar endpoints. B) Mean node ages given in units of 100 Mya (e.g., 1.2511 = 125.11 million years ago).

Supplementary Figure S47. Divergence time estimation results. Preferred topology analyzed with Dataset A at the nucleotide level under autocorrelated rates. A) Blue horizontal bars represent 95% confidence intervals with lower and upper bound ages labeled at bar endpoints. B) Mean node ages given in units of 100 Mya (e.g., 1.2511 = 125.11 million years ago).

Supplementary Figure S48. Divergence time estimation results. Preferred topology analyzed with Dataset B at the nucleotide level under autocorrelated rates. A) Blue horizontal bars represent 95% confidence intervals with lower and upper bound ages labeled at bar endpoints. B) Mean node ages given in units of 100 Mya (e.g., 1.2511 = 125.11 million years ago).

Supplementary Figure S49. Divergence time estimation results. Preferred topology analyzed with Dataset A at the nucleotide level under independent rates. A) Blue horizontal bars represent 95% confidence intervals with lower and upper bound ages labeled at bar endpoints. B) Mean node ages given in units of 100 Mya (e.g., 1.2511 = 125.11 million years ago).

Supplementary Figure S50. Divergence time estimation results. Preferred topology analyzed with Dataset B at the nucleotide level under independent rates. A) Blue horizontal bars represent 95% confidence intervals with lower and upper bound ages labeled at bar endpoints. B) Mean node ages given in units of 100 Mya (e.g., 1.2511 = 125.11 million years ago).

Supplementary Figure S51. Divergence time estimation results. Preferred topology analyzed with alternative Lycidae fossil calibration placement under two relaxed clock models (autocorrelated rates and independent rates) and two datasets (Dataset A and Dataset B) analysed at amino acid level. For each analysis, left panels (A, C, E, G) display 95% confidence intervals as blue horizontal bars with lower and upper bound ages labeled at bar endpoints. Right panels (B, D, F, H) show mean node ages given in units of 100 Mya (e.g., 1.2511 = 125.11 million years ago). A–B) Dataset A under autocorrelated rates. C–D) Dataset B under autocorrelated rates. E–F) Dataset A under independent rates. G–H) Dataset B under independent rates.

Supplementary Figure S52. Divergence time estimation results. Preferred topology analyzed with alternative Lycidae fossil calibration placement under two relaxed clock models (autocorrelated rates and independent rates) and two datasets (Dataset A and Dataset B) analysed at nucleotide level. For each analysis, left panels (A, C, E, G) display 95% confidence intervals as blue horizontal bars with lower and upper bound ages labeled at bar endpoints. Right panels (B, D, F, H) show mean node ages given in units of 100 Mya (e.g., 1.2511 = 125.11 million years ago). A–B) Dataset A under autocorrelated rates. C–D) Dataset B under autocorrelated rates. E–F) Dataset A under independent rates. G–H) Dataset B under independent rates.

Supplementary Figure S53. Family-Level Divergence Events by 20 Mya Intervals (n=22); All divergence Events (Node Ages) by 20 Mya Intervals (n=199). Based on results analyzed with Dataset A at the nucleotide level under autocorrelated rates.

Supplementary Figure S54. Ancestral state reconstruction of neoteny using Bayesian inference (BEAST). Trees A) and B) (left) display the estimated states and their probability values for the internal nodes. Tree C) (right) represents the collation of these data, with the color-coded nodes corresponding to the reconstructed female body states (see "Female body state" legend).

### Supplementary Texts

Supplementary Text S1. List of currently recognized families and subfamilies of Elateriformia, with indication of neoteny, soft-bodied forms, and extinct taxa.

Supplementary Text S2. Detailed justifications of selected fossil calibrations.

**Supplementary Table S1.** List of taxa used in analyses, indicating data type (transcriptome or genome), specimen preservation (transcriptomes: RNAlater; genomes: fresh material in ethanol or dry museum specimens), and data origin (current study or publicly available data).

| | Species | Superfamily | Family | Subfamily | Data type | Source | Voucher UPOL collection | Reference | Locality |
|---|---|---|---|---|---|---|---|---|---|
| 1 | Anorus parvicollis | Dascilloidea | Dascillidae | | TSA | 1kite, Sharkey et al. 2017, McKenna et al. 2019 | | Sharkey et al. 2017, McKenna et al. 2019 | |
| 2 | Dascillus cervinus | Dascilloidea | Dascillidae | | TSA | This study (UPOL) | R20019 | | Czech Republic |
| 3 | Dascillidae indet. | Dascilloidea | Dascillidae | | WGS | This study (UPOL) | G21028 | | Dominican Republic |
| 4 | Sandalus niger | Dascilloidea | Rhipiceridae | | TSA | 1kite, McKenna et al. 2019 | | McKenna et al. 2019 | |
| 5 | Rhipiceridae indet. | Dascilloidea | Rhipiceridae | | WGS | This study (UPOL) | G21006 | | Zambia, 2009 |
| 6 | Cneoglossa sp. | Byrrhoidea | Cneoglossidae | | WGS | This study (UPOL) | G21007 | | Nicaragua |
| 7 | Callirhipidae indet. | Byrrhoidea | Callirhipidae | | TSA | This study (UPOL) | R19006_Cali6 | | New Guinea, 2017 |
| 8 | Epilichas sp. | Byrrhoidea | Ptilodactylidae | | WGS | This study (UPOL) | | | |
| 9 | Ptilodactylidae indet. | Byrrhoidea | Ptilodactylidae | | TSA | This study (UPOL) | R19004_A3755_Ptil4 | | Dominican Republic, 2019 |
| 10 | Byrrhus pilula | Byrrhoidea | Byrrhidae | | TSA | 1kite, McKenna et al. 2019 | | McKenna et al. 2019 | |
| 11 | Notolioon sp. | Byrrhoidea | Byrrhidae | | TSA | 1kite, McKenna et al. 2019 | | McKenna et al. 2019 | |
| 12 | Dryops sp. | Byrrhoidea | Dryopidae | | TSA | 1kite, Sharkey et al. 2017, McKenna et al. 2019 | | Sharkey et al. 2017, McKenna et al. 2019 | |
| 13 | Heterocerus fenestratus | Byrrhoidea | Heteroceridae | | TSA | 1kite, McKenna et al. 2019 | | McKenna et al. 2019 | |
| 14 | Elmidae indet. | Byrrhoidea | Elmidae | | WGS | This study (UPOL) | G22024 | | Australia, 2019 |
| 15 | Limnius perrisi | Byrrhoidea | Elmidae | | TSA | This study (UPOL) | R20014 | | Czech Republic, 2020 |
| 16 | Byrrhinus sp. | Byrrhoidea | Limnichidae | | WGS | This study (UPOL) | | | |
| 17 | Limnichidae indet. | Byrrhoidea | Limnichidae | | TSA | This study (UPOL) | S39_PNG2019-39 | | Papua New Guinea, 2019 |
| 18 | Chelonarium sp. | Byrrhoidea | Chelonariidae | | WGS | This study (UPOL) | G20010 | | Indonesia, 2017 |
| 19 | Eulichas dudgeoni | Byrrhoidea | Eulichadidae | | WGS | This study (UPOL) | G20019 | | China, 2010 |
| 20 | Eubria palustris | Byrrhoidea | Psephenidae | Eubriinae | TSA | This study (UPOL) | R20017_CZ65 | | Czech Republic, 2020 |
| 21 | *Macroeubria* sp. | Byrrhoidea | Psephenidae | Eubriinae | WGS | This study (UPOL) | G21039 | | Sumatra |
| 22 | Psephenoidinae indet. | Byrrhoidea | Psephenidae | Psephenoidinae | WGS | This study (UPOL) | G21030 | | Borneo, 2013 |
| 23 | Lutrochidae indet. | Byrrhoidea | Lutrochidae | | WGS | This study (UPOL) | G22022 | | Ecuador, 2013 |
| 24 | Lutrochidae indet. | Byrrhoidea | Lutrochidae | | WGS | This study (UPOL) | G22020 | | Ecuador, 2013 |
| 25 | Protelmidae indet. | Byrrhoidea | Protelmidae | | WGS | This study (UPOL) | G22015 | | Venezuela, 2010 |
| 26 | Agrilus zanthoxylumi | Buprestoidea | Buprestidae | Agrilinae | TSA | Yang et al. 2019 | | https://www.ncbi.nlm.nih.gov/sra/?term=SRR10867869 | |
| 27 | Agrilus planipennis | Buprestoidea | Buprestidae | Agrilinae | WGS | i5k, Poelchau et al. 2014 | | Poelchau et al. 2014 | |
| 28 | Capnodis tenebrionis | Buprestoidea | Buprestidae | Chrysochroinae | TSA | Bari et al. 2019 | | Bari et al. 2019 | |
| 29 | Ptosima undecimmaculata | Buprestoidea | Buprestidae | Polycestinae | TSA | 1kite, McKenna et al. 2019 | | McKenna et al. 2019 | |
| 30 | Plectonotum laterale | Elateroidea | Cantharidae | Dysmorphocerinae | WGS | This study (UPOL) | G22002 | | Brazil |
| 31 | Rhagonycha fulva | Elateroidea | Cantharidae | Cantharinae | TSA | 1kite, McKenna et al. 2019 | | McKenna et al. 2019 | |
| 32 | Silis sp. | Elateroidea | Cantharidae | Silinae | TSA | This study (UPOL) | R19009_Sil9 | | Dominican Republic, 2019 |
| 33 | Heteromastix sp. | Elateroidea | Cantharidae | Dysmorphocerinae | TSA | This study (CSIRO) | | | ? |
| 34 | Malthodes sp. | Elateroidea | Cantharidae | Malthininae | TSA | This study (UPOL) | R20009 | | Georgia |
| 35 | Chauliognathus opacus | Elateroidea | Cantharidae | Chauliognathinae | TSA | Amaral et al. 2017 | | Amaral et al. 2017 | Brazil |
| 36 | Telegeusis sp. | Elateroidea | Omethidae | Telegeusinae | WGS | This study (UPOL) | G21042 | | USA |
| 37 | Drilonius sp. | Elateroidea | Omethidae | Driloniinae | TSA | This study (CSIRO) | | | |
| 38 | Eurypogon sp. | Elateroidea | Artematopodidae | | TSA | This study (CSIRO) | | | |
| 39 | Artematopus sp. | Elateroidea | Artematopodidae | | WGS | This study (UPOL) | G20035 | | Ecuador, 2010 |
| 40 | Aulonothroscus constrictor | Elateroidea | Throscidae | | WGS | This study (UPOL) | A4488 | | USA |
| 41 | Trixagus carinifrons | Elateroidea | Throscidae | | TSA | 1kite, McKenna et al. 2019 | | McKenna et al. 2019 | |
| 42 | Jurasai digitusdei | Elateroidea | Jurasaidae | | WGS | This study (UPOL) | G22009 | | Brazil |
| 43 | Cerophytum elateroides | Elateroidea | Cerophytidae | | WGS | This study (UPOL) | Cero14 | | Slovakia |
| 44 | Cerophytum japonicum | Elateroidea | Cerophytidae | | WGS | This study (UPOL) | G22003 | | Japan, 2021 |
| 45 | Brachypsectra fulva | Elateroidea | Brachypsectridae | | TSA | This study (CSIRO) | | | USA |
| 46 | Anischia sp. | Elateroidea | Eucnemidae | Anischinae | WGS | This study (UPOL) | | | New Caledonia |
| 47 | Isorhipis marmottani | Elateroidea | Eucnemidae | Melasinae | TSA | This study (UPOL) | R19001_Isor01 | | Czech Republic |
| 48 | Eucnemis capucina | Elateroidea | Eucnemidae | Eucneminae | TSA | This study (CSIRO) | | | ? |
| 49 | Dromaeolinae indet. | Elateroidea | Eucnemidae | Dromaeolinae | WGS | This study (UPOL) | G19013_Elat13 | | Indonesia |
| 50 | Eucnemidae indet. | Elateroidea | Eucnemidae | | WGS | This study (UPOL) | G20008 | | Papua New Guinea, 2019 |
| 51 | Ceratogonys spinicorne | Elateroidea | Eucnemidae | Dromaeolinae | WGS | This study (UPOL) | G21023 | | French Guiana, 2016 |
| 52 | Eucnemidae indet. | Elateroidea | Eucnemidae | | WGS | This study (UPOL) | G21019 | | French Guiana, 2021 |
| 53 | Rhagophthalmidae indet. | Elateroidea | Rhagophthalmidae | | WGS | This study (UPOL) | G20030 | | Borneo |
| 54 | Rhagophthalmidae indet. | Elateroidea | Rhagophthalmidae | | WGS | This study (UPOL) | G20031 | | Thailand |
| 55 | Rhagophthalmus sp. | Elateroidea | Rhagophthalmidae | | TSA | Wang et al. 2017 | | Wang et al. 2017 | |
| 56 | Rhagophthalmidae indet. | Elateroidea | Rhagophthalmidae | | WGS | This study (UPOL) | G21016 | | China, 2014 |
| 57 | Phengodes sp. | Elateroidea | Phengodidae | Phengodinae | WGS | Timothy Fallon, unpublished | | | |
| 58 | Phengodes sp. | Elateroidea | Phengodidae | Phengodinae | TSA | This study (CSIRO) | | | ? |
| 59 | Phrixothrix hirtus | Elateroidea | Phengodidae | Mastinocerinae | TSA | Amaral et al. 2017 | | Amaral et al. 2017 | |
| 60 | Distremocephalus sp. | Elateroidea | Phengodidae | Mastinocerinae | TSA | This study (CSIRO) | | | ? |
| 61 | Sinopyrophorus schimmeli | Elateroidea | Sinopyrophoridae | | WGS | He et al. 2019 | | | |
| 62 | Sinopyrophorus sp. | Elateroidea | Sinopyrophoridae | | WGS | This study (UPOL) | G20013 | | Vietnam |
| 63 | Brachypterodrilus sp. | Elateroidea | Lampyridae | Ototretinae | WGS | This study (UPOL) | G21015 | | Malaysia, 2013 |
| 64 | Stenocladius sp. | Elateroidea | Lampyridae | Ototretinae | WGS | This study (UPOL) | G20028 | | China, 2011 |
| 65 | Stenocladius sp. | Elateroidea | Lampyridae | Ototretinae | WGS | This study (UPOL) | G22026 | | Sumatra |
| 66 | Drilaster sp. | Elateroidea | Lampyridae | Ototretinae | WGS | This study (UPOL) | G22025 | | Japan |
| 67 | Lamellipalpus pacholatkoi | Elateroidea | Lampyridae | Ototretinae | WGS | This study (UPOL) | Lam_379 | | ? |
| 68 | Ototretinae indet. | Elateroidea | Lampyridae | Ototretinae | TSA | This study (UPOL) | R19005_Otot05 | | Indonesia, 2017 |
| 69 | Ototretinae indet. | Elateroidea | Lampyridae | Ototretinae | TSA | This study (UPOL) | R20013_A3673 | | Papua New Guinea, 2019 |
| 70 | Ototretinae indet. | Elateroidea | Lampyridae | Ototretinae | WGS | This study (UPOL) | G21011 | | Malaysia, 2016 |
| 71 | Ototretinae indet. | Elateroidea | Lampyridae | Ototretinae | WGS | This study (UPOL) | G21029 | | Malaysia |
| 72 | Psilocladus sp. | Elateroidea | Lampyridae | Psilocladinae | WGS | This study (UPOL) | G21009 | | Nicaragua |
| 73 | Amydetinae indet. | Elateroidea | Lampyridae | Amydetinae | WGS | This study (UPOL) | G21010 | | Brazil |
| 74 | Cheguevaria sp. | Elateroidea | Lampyridae | Cheguevarinae | WGS | This study (UPOL) | G20025 | | ? |
| 75 | Asymmetricata circumdata | Elateroidea | Lampyridae | Luciolinae | TSA | Wang et al. 2017 | | Wang et al. 2017 | |
| 76 | Aquatica ficta | Elateroidea | Lampyridae | Luciolinae | TSA | Wang et al. 2017 | | Wang et al. 2017 | |
| 77 | Abscondita terminalis | Elateroidea | Lampyridae | Luciolinae | WGS | Zhang et al. 2020 | | | |
| 78 | Lamprigera yunnana | Elateroidea | Lampyridae | Luciolinae | WGS | Zhang et al. 2020 | | | |
| 79 | Lamprohiza splendidula | Elateroidea | Lampyridae | Lamprohizinae | TSA | 1kite, McKenna et al. 2019 | | McKenna et al. 2019 | |
| 80 | Phausis reticulata | Elateroidea | Lampyridae | Lamprohizinae | TSA | Sander and Hall 2015 | | Sander and Hall 2015 | |
| 81 | Lampyrinae indet. | Elateroidea | Lampyridae | Lampyrinae | WGS | This study (UPOL) | G22027 | | Borneo |
| 82 | Microphotus sp. | Elateroidea | Lampyridae | Lampyrinae | TSA | Gavin et al. 2021 | | Gavin et al. 2021 | |
| 83 | Ellychnia sp. | Elateroidea | Lampyridae | Lampyrinae | TSA | Gavin et al. 2021 | | Gavin et al. 2021 | |
| 84 | Micronaspis floridana | Elateroidea | Lampyridae | Lampyrinae | TSA | Gavin et al. 2021 | | Gavin et al. 2021 | |
| 85 | Photinus carolinus | Elateroidea | Lampyridae | Lampyrinae | TSA | Sander and Hall 2015 | | | |
| 86 | Photinus australis | Elateroidea | Lampyridae | Lampyrinae | TSA | Sander and Hall 2015 | | | |
| 87 | Photinus pyralis | Elateroidea | Lampyridae | Lampyrinae | TSA | Fallon et al. 2018 | | | |
| 88 | Photinus scintillans | Elateroidea | Lampyridae | Lampyrinae | TSA | Sander and Hall 2015 | | Sander and Hall 2015 | |
| 89 | Photinus macdermotti | Elateroidea | Lampyridae | Lampyrinae | TSA | Sander and Hall 2015 | | | |
| 90 | Pyrocoelia pectoralis | Elateroidea | Lampyridae | Lampyrinae | TSA | Wang et al. 2017 | | Wang et al. 2017 | |
| 91 | Lampyris noctiluca | Elateroidea | Lampyridae | Lampyrinae | TSA | This study (UPOL) | R20022 | | Czech Republic, 2020 |
| 92 | Pyractomena borealis | Elateroidea | Lampyridae | Lampyrinae | TSA | Sander and Hall 2015 | | Sander and Hall 2015 | |
| 93 | Aspisoma lineatum | Elateroidea | Lampyridae | Lampyrinae | TSA | Amaral et al. 2017 | | Amaral et al. 2017 | |
| 94 | Lucidota atra | Elateroidea | Lampyridae | Lampyrinae | TSA | Sander and Hall 2015 | | Sander and Hall 2015 | |
| 95 | Phosphaenus hemipterus | Elateroidea | Lampyridae | Lampyrinae | TSA | This study (UPOL) | R20018 | | Czech Republic, 2020 |
| 96 | Lampyris sardiniae | Elateroidea | Lampyridae | Lampyrinae | TSA | This study (UPOL) | S1 | | Sardinia |
| 97 | Photuris frontalis | Elateroidea | Lampyridae | Photurinae | TSA | Sander and Hall 2015 | | Sander and Hall 2015 | |
| 98 | Photuris sp. | Elateroidea | Lampyridae | Photurinae | TSA | Sander and Hall 2015 | | Sander and Hall 2015 | |
| 99 | Bicellonycha wickershamorum | Elateroidea | Lampyridae | Photurinae | TSA | Gavin et al. 2021 | | Gavin et al. 2021 | |
| 100 | Iberobaenia minuta | Elateroidea | Iberobaeniidae | | WGS | This study (UPOL) | G19015_Iber15 | | Spain |
| 101 | Dexoris ruzzieri | Elateroidea | Lycidae | Dexorinae | WGS | UPOL, Kusy et al. 2019 | | Kusy et al. 2019 | |
| 102 | Lygistopterus sanguineus | Elateroidea | Lycidae | Calochrominae | TSA | UPOL, Kusy et al. 2019 | | Kusy et al. 2019 | |
| 103 | Konoplatycis otome | Elateroidea | Lycidae | Erotinae | TSA | UPOL, Kusy et al. 2019 | | Kusy et al. 2019 | |
| 104 | Dictyoptera sp. | Elateroidea | Lycidae | Erotinae | TSA | UPOL, Kusy et al. 2019 | | Kusy et al. 2019 | |
| 105 | Taphes brevicollis | Elateroidea | Lycidae | Erotinae | TSA | UPOL, Kusy et al. 2019 | | Kusy et al. 2019 | |
| 106 | Scarelus anthracinus | Elateroidea | Lycidae | Atelinae | TSA | UPOL, Kusy et al. 2019 | | Kusy et al. 2019 | |
| 107 | Ponyalis_sp | Elateroidea | Lycidae | Atelinae | TSA | UPOL, Kusy et al. 2019 | | Kusy et al. 2019 | |
| 108 | Ponyalis quadricollis | Elateroidea | Lycidae | Atelinae | TSA | UPOL, Kusy et al. 2019 | | Kusy et al. 2019 | |
| 109 | Conderis signicollis | Elateroidea | Lycidae | Lycinae | TSA | UPOL, Kusy et al. 2019 | | Kusy et al. 2019 | |
| 110 | Leptolycus sp. | Elateroidea | Lycidae | Lycinae | TSA | UPOL, Kusy et al. 2019 | | Kusy et al. 2019 | |
| 111 | Calopteron sp. | Elateroidea | Lycidae | Lycinae | TSA | UPOL, Kusy et al. 2019 | | Kusy et al. 2019 | |
| 112 | Lycostomus kraatzi | Elateroidea | Lycidae | Lycinae | TSA | UPOL, Kusy et al. 2019 | | Kusy et al. 2019 | |
| 113 | Plateros sp. | Elateroidea | Lycidae | Lycinae | TSA | UPOL, Kusy et al. 2019 | | Kusy et al. 2019 | |
| 114 | Antennolycus constrictus | Elateroidea | Lycidae | Lyropaeinae | TSA | UPOL, Kusy et al. 2019 | | Kusy et al. 2019 | |
| 115 | Lyropaeus optabilis | Elateroidea | Lycidae | Lyropaeinae | TSA | UPOL, Kusy et al. 2019 | | Kusy et al. 2019 | |
| 116 | Platerodrilus sp. | Elateroidea | Lycidae | Lyropaeinae | TSA | UPOL, Kusy et al. 2019 | | Kusy et al. 2019 | |
| 117 | Dihammatus sp. | Elateroidea | Lycidae | Metriorrhynchinae | TSA | UPOL, Kusy et al. 2019 | | Kusy et al. 2019 | |
| 118 | Lycoprogenthes sp. | Elateroidea | Lycidae | Metriorrhynchinae | TSA | UPOL, Kusy et al. 2019 | | Kusy et al. 2019 | |
| 119 | Libnetis sp. | Elateroidea | Lycidae | Metriorrhynchinae | TSA | UPOL, Kusy et al. 2019 | | Kusy et al. 2019 | |
| 120 | Dilophotes sp. | Elateroidea | Lycidae | Metriorrhynchinae | TSA | UPOL, Kusy et al. 2019 | | Kusy et al. 2019 | |
| 121 | Cautires communis | Elateroidea | Lycidae | Metriorrhynchinae | WGS | UPOL, Kusy et al. 2019 | | Kusy et al. 2019 | |
| 122 | Mangkutanus sp. | Elateroidea | Lycidae | Metriorrhynchinae | TSA | UPOL, Kusy et al. 2019 | R18040 | | |
| 123 | Anaspasis paralella | Elateroidea | Elateridae | Lissominae | WGS | UPOL, Motyka et al. 2025 | G22007 | | Chile, 2016 |
| 124 | Rostricephalus vitalis | Elateroidea | Elateridae | Lissominae | WGS | UPOL, Motyka et al. 2025 | G20023 | | Vietnam |
| 125 | Sphaenelater lineicollis | Elateroidea | Elateridae | Lissominae | WGS | UPOL, Motyka et al. 2025 | G21002 | | New Zealand, 2020 |
| 126 | Lissomus sp. | Elateroidea | Elateridae | Lissominae | TSA | UPOL, Motyka et al. 2025 | R19013_Liss13_A3758 | | Dominican Republic |
| 127 | Lissominae indet. | Elateroidea | Elateridae | Lissominae | WGS | UPOL, Motyka et al. 2025 | G21017 | | Tanzania, 2014 |
| 128 | Drapetes sp. | Elateroidea | Elateridae | Lissominae | WGS | UPOL, Motyka et al. 2025 | G20009 | | Papua New Guinea |
| 129 | Balgus obconicus | Elateroidea | Elateridae | Thylacosterninae | WGS | UPOL, Motyka et al. 2025 | G20011 | | French Guiana |
| 130 | Cussolenis sp. | Elateroidea | Elateridae | Thylacosterninae | WGS | UPOL, Motyka et al. 2025 | G20003 | | Philippines |
| 131 | Cebrio sp. | Elateroidea | Elateridae | Elaterinae | WGS | UPOL, Motyka et al. 2025 | G19010_Cebr10 | | Italy |
| 132 | Agriotes sp. | Elateroidea | Elateridae | Elaterinae | TSA | UPOL, Motyka et al. 2025 | R20008 | | Slovakia |
| 133 | Elater ferrugineus | Elateroidea | Elateridae | Elaterinae | WGS | UPOL, Motyka et al. 2025 | G20021 | | Czech Republic, 2020 |
| 134 | Mulsanteus_sp. | Elateroidea | Elateridae | Elaterinae | WGS | UPOL, Motyka et al. 2025 | G20005 | | ? |
| 135 | Melanotus cribricollis | Elateroidea | Elateridae | Elaterinae | TSA | Ye et al. 2018 | | Ye et al. 2018 | China |
| 136 | Melanotus villosus | Elateroidea | Elateridae | Elaterinae | TSA | Mckenna et al. 2019 | | Mckenna et al. 2019 | United Kingdom |
| 137 | Ampedus sineatus | Elateroidea | Elateridae | Elaterinae | TSA | UPOL, Motyka et al. 2025 | R20005 | | Czech Republic, 2020 |
| 138 | Ampedus cinnabarinus | Elateroidea | Elateridae | Elaterinae | TSA | UPOL, Motyka et al. 2025 | R20006 | | Czech Republic, 2020 |
| 139 | Elaterinae indet. | Elateroidea | Elateridae | Elaterinae | WGS | UPOL, Motyka et al. 2025 | G21027 | | French Guiana, 2021 |
| 140 | Elaterinae indet. | Elateroidea | Elateridae | Elaterinae | TSA | UPOL, Motyka et al. 2025 | R20010 | | Papua New Guinea |
| 141 | Elaterinae indet. | Elateroidea | Elateridae | Elaterinae | WGS | UPOL, Motyka et al. 2025 | G20006 | | Papua New Guinea |
| 142 | Octinodes sp. | Elateroidea | Elateridae | Elaterinae | WGS | UPOL, Motyka et al. 2025 | G20001 | | Panama |
| 143 | Eudicronychus rufus | Elateroidea | Elateridae | Elaterinae | WGS | UPOL, Motyka et al. 2025 | G20004 | | Zambia |
| 144 | Eudicronychus sp. | Elateroidea | Elateridae | Elaterinae | WGS | UPOL, Motyka et al. 2025 | G19007_Eudi07 | | Zambia |
| 145 | Elaterinae indet. | Elateroidea | Elateridae | Elaterinae | TSA | Amaral et al. 2018 | | | |
| 146 | Hemiops sp. | Elateroidea | Elateridae | Hemiopinae | WGS | UPOL, Motyka et al. 2025 | G19002_Hemi02 | | Malaysia |
| 147 | Paulusiella serraticornis | Elateroidea | Elateridae | Hemiopinae | WGS | UPOL, Motyka et al. 2025 | G21004 | | Iran, 2018 |
| 148 | Diplophoenicus sp. | Elateroidea | Elateridae | Morostomatinae | WGS | UPOL, Motyka et al. 2025 | G19011_Moro11 | | Madagascar |
| 149 | Elatichrosis sp. | Elateroidea | Elateridae | Pityobiinae | WGS | UPOL, Motyka et al. 2025 | G21014 | | Australia, 2019 |
| 150 | Parasaphes sp. | Elateroidea | Elateridae | Pityobiinae | WGS | UPOL, Motyka et al. 2025 | G19006_Para06 | | Australia |
| 151 | Rousia sp. | Elateroidea | Elateridae | Pityobiinae | WGS | UPOL, Motyka et al. 2025 | G21031 | | Australia, 2014 |
| 152 | Tibionema abdominalis | Elateroidea | Elateridae | Pityobiinae | WGS | UPOL, Motyka et al. 2025 | G20012 | | Chile |
| 153 | Pityobius anguinus | Elateroidea | Elateridae | Pityobiinae | WGS | UPOL, Motyka et al. 2025 | G21037_G21037b | | Chile, 1992 |
| 154 | Campyloxenus pyrothorax | Elateroidea | Elateridae | Pityobiinae | WGS | UPOL, Motyka et al. 2025 | G21036 | | Chile, 2001 |
| 155 | Hapatesus tropicus | Elateroidea | Elateridae | Hapatesinae | WGS | UPOL, Motyka et al. 2025 | G20007 | | Papua |
| 156 | Hypnoidus riparius | Elateroidea | Elateridae | Dendrometrinae | TSA | This study (CSIRO) | | | ? |
| 157 | Denticollis sp. | Elateroidea | Elateridae | Dendrometrinae | TSA | UPOL, Motyka et al. 2025 | R19012_Dent12_A3752 | | Japan |
| 158 | Semiotus sp. | Elateroidea | Elateridae | Dendrometrinae | WGS | UPOL, Motyka et al. 2025 | G19009_Semi09 | | Ecuador |
| 159 | Athous sp. | Elateroidea | Elateridae | Dendrometrinae | TSA | UPOL, Motyka et al. 2025 | R20003 | | Czech Republic, 2020 |
| 160 | Selatosomus aeneus | Elateroidea | Elateridae | Dendrometrinae | TSA | UPOL, Motyka et al. 2025 | R20004 | | Czech Republic, 2020 |
| 161 | Limonius californicus | Elateroidea | Elateridae | Dendrometrinae | WGS | Andrews et al. 2020 | | | |
| 162 | Anostirus purpureus | Elateroidea | Elateridae | Dendrometrinae | TSA | UPOL, Motyka et al. 2025 | R20007 | | Slovakia, 2020 |
| 163 | Penia sp. | Elateroidea | Elateridae | Dendrometrinae | WGS | UPOL, Motyka et al. 2025 | G19012_Elat12 | | China |
| 164 | Pectocera sp. | Elateroidea | Elateridae | Dendrometrinae | WGS | UPOL, Motyka et al. 2025 | G19004_Pect04 | | China |
| 165 | Oxynopterus sp. | Elateroidea | Elateridae | Dendrometrinae | WGS | UPOL, Motyka et al. 2025 | G19008_Oxyn08 | | Philippines |
| 166 | Plastocerus angulosus | Elateroidea | Elateridae | Dendrometrinae | WGS | UPOL, Kusy et al. 2018 | A01544 | Kusy et al. 2018 | Turkey |
| 167 | Negastrius arenicola | Elateroidea | Elateridae | Negastriinae | TSA | This study (CSIRO) | | | ? |
| 168 | Quasimus sp. | Elateroidea | Elateridae | Negastriinae | TSA | UPOL, Motyka et al. 2025 | R19010_Elat10_A3757 | | Japan |
| 169 | Dicronychus rubripes | Elateroidea | Elateridae | Cardiophorinae | TSA | UPOL, Motyka et al. 2025 | R19003_Card3_A3754 | | Slovakia |
| 170 | Cardiophorus discicollis | Elateroidea | Elateridae | Cardiophorinae | TSA | UPOL, Motyka et al. 2025 | R20016 | | Czech Republic |
| 171 | Analastesa arabica | Elateroidea | Elateridae | Cardiophorinae | WGS | UPOL, Motyka et al. 2025 | G21003 | | Saudi Arabia, 1975 |
| 172 | Pseudotetralobus capucinus | Elateroidea | Elateridae | Tetralobinae | WGS | UPOL, Motyka et al. 2025 | G20022 | | Australia |
| 173 | Sinelater perroti | Elateroidea | Elateridae | Tetralobinae | WGS | UPOL, Motyka et al. 2025 | G19001_Tetr01 | | China |
| 174 | Euanoma sp. | Elateroidea | Elateridae | Omalisinae | WGS | UPOL, Motyka et al. 2025 | G20033 | | Georgia |
| 175 | Omalisus fontisbellaquei | Elateroidea | Elateridae | Omalisinae | WGS | UPOL, Kusy et al. 2018 | G18001 | Kusy et al. 2018 | Czech Republic |
| 176 | Drilus concolor | Elateroidea | Elateridae | Agrypninae | TSA | 1kite, Mckenna et al. 2019 | | Mckenna et al. 2019 | |
| 177 | Drilus mauritanicus | Elateroidea | Elateridae | Agrypninae | WGS | UPOL, Kusy et al. 2018 | G18004 | Kusy et al. 2018 | Turkey |
| 178 | Austroselasia sp. | Elateroidea | Elateridae | Agrypninae | WGS | UPOL, Motyka et al. 2025 | G21012 | | Zambia, 2007 |
| 179 | Habashaselasia nekemtensis | Elateroidea | Elateridae | Agrypninae | WGS | UPOL, Motyka et al. 2025 | G21013 | | Ethiopia, 2009 |
| 180 | Pachyderes sp. | Elateroidea | Elateridae | Agrypninae | WGS | UPOL, Motyka et al. 2025 | G19005_Pach05 | | Malaysia |
| 181 | Anaissus sp. | Elateroidea | Elateridae | Agrypninae | WGS | UPOL, Motyka et al. 2025 | G21035 | | Panama |
| 182 | Platycrepidius sp. | Elateroidea | Elateridae | Agrypninae | WGS | UPOL, Motyka et al. 2025 | G19060 | | Panama |
| 183 | Ignelater luminosus | Elateroidea | Elateridae | Agrypninae | WGS | Fallon et al. 2018 | | Fallon et al. 2018 | |
| 184 | Pyrearinus fragilis | Elateroidea | Elateridae | Agrypninae | TSA | Amaral et al. 2019 | | Amaral et al. 2019 | |
| 185 | Pyrophorini indet. | Elateroidea | Elateridae | Agrypninae | WGS | UPOL, Motyka et al. 2025 | G21026 | | French Guiana, 2021 |
| 186 | Pyrearinus sp. | Elateroidea | Elateridae | Agrypninae | WGS | UPOL, Motyka et al. 2025 | G20002 | | Bolivia |
| 187 | Agrypnus murinus | Elateroidea | Elateridae | Agrypninae | TSA | This study (CSIRO) | | | ? |
| 188 | Agrypninae indet. | Elateroidea | Elateridae | Agrypninae | TSA | UPOL, Motyka et al. 2025 | R20002 | | Papua New Guinea |
| 189 | Alaus myops | Elateroidea | Elateridae | Agrypninae | TSA | UPOL, Motyka et al. 2025 | R20011 | | USA |
| 190 | Aphileus lucanoides | Elateroidea | Elateridae | Agrypninae | WGS | UPOL, Motyka et al. 2025 | G20024 | | Australia |
| 191 | Agrypninae indet. | Elateroidea | Elateridae | Agrypninae | TSA | UPOL, Motyka et al. 2025 | R19011_Elat11_A3756 | | Japan, 2015 |
| 192 | Adelocera sp. | Elateroidea | Elateridae | Agrypninae | WGS | UPOL, Motyka et al. 2025 | G21038 | | Philippines, 2019 |
| 193 | Adelocera sp. | Elateroidea | Elateridae | Agrypninae | WGS | UPOL, Motyka et al. 2025 | G21041 | | Vietnam, 2020 |
| 194 | Adelocera sp. | Elateroidea | Elateridae | Agrypninae | WGS | UPOL, Motyka et al. 2025 | G21040 | | Vietnam, 2020 |
| 195 | Nyctophyxis ocellatus | Elateroidea | Elateridae | Agrypninae | WGS | UPOL, Motyka et al. 2025 | G22005 | | Chile, 2019 |
| 196 | Tribolium castaneum | Tenebrionoidea | Tenebrionidae | | WGS | Reference, outgroup | | See Supplementary Table S2 | |
| 197 | Dendroctonus ponderosae | Curculionoidea | Curculionidae | | WGS | Reference, outgroup | | See Supplementary Table S2 | |
| 198 | Onthophagus taurus | Scarabaeoidea | Scarabeidae | | WGS | Reference, outgroup | | See Supplementary Table S2 | |
| 199 | Anoplophora glabripennis | Chrysomeloidea | Cerambycidae | | WGS | Reference, outgroup | | See Supplementary Table S2 | |
| 200 | Leptinotarsa decemlineata | Chrysomeloidea | Chrysomelidae | | WGS | Reference, outgroup | | See Supplementary Table S2 | |

**Supplementary Table S2.** Overview of gene sets of six reference species used for ortholog assessment, including the source, version and number of genes. URLs for downloading the data are given. The number of genes corresponds with OrthoDB 9.1.

| Species | Accession | # of genes | Version | Source | Download Date | Reference |
|---|---|---|---|---|---|---|
| *Onthophagus taurus* | PRJNA167478 | 17483 | 0.5.3 | i5K | 5.3.2017 | 1 |
| *Tribolium castaneum* | PRJNA12540 | 16631 | 5.2 | iBeetle | 5.3.2017 | 2,3 |
| *Dendroctonus ponderosae* | PRJNA360270 | 13088 | 1.0 | ENS Metazoa | 5.3.2017 | 4 |
| *Anoplophora glabripennis* | PRJNA167479 | 22035 | 0.5.3 | i5K | 5.3.2017 | 5 |
| *Leptinotarsa decemlineata* | PRJNA171749 | 24671 | 0.5.3 | i5K | 5.3.2017 | 1 |
| *Agrilus planipennis* | PRJNA230921 | 15497 | 0.5.3 | i5K | 5.3.2017 | 1 |

***Tribolium castaneum***
http://bioinf.uni-greifswald.de/tcas/genes/tcas5_annotation/Tcas5.2_GenBank.corrected_v5.renamed.aa
http://bioinf.uni-greifswald.de/tcas/genes/tcas5_annotation/Tcas5.2_GenBank.corrected_v5.renamed.codingseq
***Onthophagus taurus***
https://i5k.nal.usda.gov/data/Arthropoda/onttau-%28Onthophagus_taurus%29/Current%20Genome%20Assembly/2.Official%20or%20Primary%20Gene%20Set/BCM_version_0.5.3/consensus_gene_set/OTAU.faa
https://i5k.nal.usda.gov/data/Arthropoda/onttau-%28Onthophagus_taurus%29/Current%20Genome%20Assembly/2.Official%20or%20Primary%20Gene%20Set/BCM_version_0.5.3/consensus_gene_set/OTAU.CDS.fna
***Anoplophora glabripennis***

https://i5k.nal.usda.gov/data/Arthropoda/anogla-%28Anoplophora_glabripennis%29/Current%20Genome%20Assembly/3.Additional%20Gene%20Sets%20and%20Annotation%20Projects/BCM_version_0.5.3-Primary_Gene_Set/primary_gene_set/AGLA.CDS.fna.gz
https://i5k.nal.usda.gov/data/Arthropoda/anogla-%28Anoplophora_glabripennis%29/Current%20Genome%20Assembly/3.Additional%20Gene%20Sets%20and%20Annotation%20Projects/BCM_version_0.5.3-Primary_Gene_Set/primary_gene_set/AGLA.faa.gz

***Agrilus planipennis***

https://i5k.nal.usda.gov/data/Arthropoda/agrpla-%28Agrilus_planipennis%29/Current%20Genome%20Assembly/2.Official%20or%20Primary%20Gene%20Set/BCM_version_0.5.3/consensus_gene_set/APLA.faa
https://i5k.nal.usda.gov/data/Arthropoda/agrpla-%28Agrilus_planipennis%29/Current%20Genome%20Assembly/2.Official%20or%20Primary%20Gene%20Set/BCM_version_0.5.3/consensus_gene_set/APLA.CDS.fna

***Dendroctonus ponderosae***

ftp://ftp.ensemblgenomes.org/pub/metazoa/release-37/fasta/dendroctonus_ponderosae/cds/Dendroctonus_ponderosae.DendPond_male_1.0.cds.all.fa.gz
ftp://ftp.ensemblgenomes.org/pub/metazoa/release-37/fasta/dendroctonus_ponderosae/pep/Dendroctonus_ponderosae.DendPond_male_1.0.pep.all.fa.gz

***Leptinotarsa decemlineata***

https://i5k.nal.usda.gov/data/Arthropoda/lepdec-%28Leptinotarsa_decemlineata%29/Current%20Genome%20Assembly/2.Official%20or%20Primary%20Gene%20Set/BCM_version_0.5.3-Primary_Gene_Set/primary_gene_set/LDEC.faa.gz
https://i5k.nal.usda.gov/data/Arthropoda/lepdec-%28Leptinotarsa_decemlineata%29/Current%20Genome%20Assembly/2.Official%20or%20Primary%20Gene%20Set/BCM_version_0.5.3-Primary_Gene_Set/primary_gene_set/LDEC.CDS.fna.gz

**Supplementary Table S3.** Summary statistics by taxon across data processing stages (raw data, outlier removal, and AliScore filtering) at the amino acid level.

| Taxon _name_Family | AA raw data | | | | | | | |
|---|---|---|---|---|---|---|---|---|
| | no_genes | total_no_aa | N50 | percent_determined | average_length | median_length | max_length | min_length |
| A4488_Throscidae_indet_Throscidae | 3839 | 1155214 | 358 | 37.16 | 300.9 | 250 | 5615 | 6 |
| Abscondita_terminalis_Lampyridae | 3756 | 1643592 | 528 | 54.75 | 437.6 | 356 | 5504 | 29 |
| Agrilus_planipennis_Buprestidae | 4224 | 2338433 | 706 | 67.52 | 553.6 | 422 | 9009 | 54 |
| Agrilus_zanthoxylumi_Buprestidae | 3981 | 1942274 | 611 | 60.84 | 487.9 | 388 | 5701 | 15 |
| Agrypnus_murinus_Elateridae | 3666 | 1415530 | 472 | 48.26 | 386.1 | 327 | 4104 | 24 |
| Anoplophora_glabripennis | 4224 | 2266122 | 665 | 65.43 | 536.5 | 408 | 19530 | 59 |
| Anorus_parvicollis_Dascillidae | 3719 | 1455139 | 477 | 49.23 | 391.3 | 326 | 7875 | 7 |
| Antennolycus_constrictus_Lycidae | 3739 | 1556417 | 511 | 52.2 | 416.3 | 341 | 5028 | 31 |
| Aquatica_ficta_Lampyridae | 3703 | 1431279 | 477 | 48.38 | 386.5 | 324 | 4749 | 24 |
| Asymmetricata_circumdata_Lampyridae | 3603 | 1461460 | 493 | 51.28 | 405.6 | 339 | 5726 | 12 |
| Aspisoma_lineatum_Lampyridae | 3908 | 1715219 | 532 | 54.72 | 438.9 | 358 | 7036 | 16 |
| Bicellonycha_wickershamorum_Lampyridae | 3622 | 1364334 | 465 | 46.98 | 376.7 | 317 | 5362 | 6 |
| Brachypsectra_fulva_Brachypsectridae | 3586 | 1261695 | 433 | 44.5 | 351.8 | 315 | 3308 | 7 |
| Byrrhus_pilula_Byrrhidae | 3349 | 1181416 | 432 | 44.61 | 352.8 | 306 | 3614 | 30 |
| R19006_Callirhipidae_indet_Callirhipidae | 3700 | 1519799 | 507 | 51.59 | 410.8 | 333.5 | 7663 | 12 |
| Calopteron_sp_Lycidae | 3668 | 1574663 | 521 | 53.8 | 429.3 | 353.5 | 5603 | 8 |
| Capnodis_tenebrionis_Buprestidae | 3777 | 1658014 | 535 | 54.16 | 439 | 362 | 5917 | 16 |
| R19003_Dicronychus_rubripes_Elateridae | 3885 | 1725210 | 535 | 55.88 | 444.1 | 364 | 8322 | 37 |
| Cautires_communis_Lycidae | 3780 | 1069710 | 331 | 34.4 | 283 | 240 | 3254 | 11 |
| G19010_Cebrio_sp_Elateridae | 3535 | 895518 | 304 | 29.91 | 253.3 | 217 | 3373 | 6 |
| Cerophytum_elateroides_Cerophytiae | 3250 | 610441 | 221 | 22.59 | 187.8 | 157 | 1359 | 7 |
| Chauliognathus_opacus_Cantharidae | 3669 | 948903 | 311 | 32.24 | 258.6 | 212 | 5468 | 8 |
| Conderis_signicollis_Lycidae | 3678 | 1422221 | 477 | 48.07 | 386.7 | 318.5 | 7547 | 35 |
| Elaterinae_indet_Elateridae | 3907 | 1803747 | 554 | 57.53 | 461.7 | 378 | 6136 | 9 |
| Dendroctonus_ponderosae | 4224 | 2242670 | 662 | 64.75 | 530.9 | 415 | 7334 | 50 |
| R19012_Denticollis_sp_Elateridae | 3908 | 1706815 | 526 | 54.94 | 436.7 | 364 | 4323 | 16 |
| Dexoris_ruzzieri_Lycidae | 3897 | 1026773 | 309 | 31.98 | 263.5 | 224 | 3495 | 12 |
| Dictyoptera_sp_Lycidae | 3401 | 873345 | 328 | 32.44 | 256.8 | 210 | 3363 | 7 |
| Dihammatus_sp_Lycidae | 3731 | 1533411 | 505 | 50.91 | 411 | 342 | 7122 | 10 |
| Dilophotes_sp_Lycidae | 3715 | 1542067 | 504 | 51.84 | 415.1 | 346 | 3641 | 7 |
| Distremocephalus_sp_Phengodidae | 3648 | 1374234 | 460 | 47.92 | 376.7 | 328 | 2853 | 11 |
| Drilonius_sp_Omethidae | 3870 | 1691498 | 529 | 54.48 | 437.1 | 361 | 6144 | 20 |
| G18004_Drilus_mauritanicus_Elateridae | 3978 | 1535735 | 469 | 47.73 | 386.1 | 317 | 6557 | 7 |
| Drilus_concolor_Elateridae | 3030 | 865385 | 356 | 36.11 | 285.6 | 241 | 3350 | 19 |
| Dryops_sp_Dryopidae | 3696 | 1508774 | 500 | 51.44 | 408.2 | 343 | 3112 | 17 |
| R19010_Quasimus_sp_Elateridae | 3909 | 1707394 | 528 | 54.73 | 436.8 | 359 | 5064 | 6 |
| R19011_Agrypninae_indet_Elateridae | 3843 | 1678495 | 532 | 54.63 | 436.8 | 364 | 7423 | 8 |
| G19012_Penia_sp_Elateridae | 3935 | 1427446 | 435 | 44.6 | 362.8 | 309 | 4774 | 6 |
| G19013_Dromaeolinae indet_Eucnemidae | 3866 | 1449084 | 459 | 45.81 | 374.8 | 307 | 4896 | 13 |
| Ellychnia_sp_Lampyridae | 3734 | 1496580 | 493 | 49.38 | 400.8 | 332 | 4657 | 8 |
| Eucnemis_capucina_Eucnemidae | 3928 | 1637848 | 508 | 51.66 | 417 | 340.5 | 4781 | 16 |
| G19007_Eudicronychus_sp_Elateridae | 3511 | 883158 | 304 | 29.51 | 251.5 | 211 | 2219 | 10 |
| Eurypogon_sp_Artematopodidae | 3653 | 1230644 | 404 | 42.46 | 336.9 | 313 | 1444 | 12 |
| G20001_Octinodes_sp_Elateridae | 3932 | 1417691 | 438 | 44.3 | 360.6 | 294 | 3635 | 31 |
| G20002_Pyrearinus_sp_Elateridae | 3978 | 1631799 | 497 | 50.53 | 410.2 | 339 | 4726 | 21 |
| G20003_Cussolenis_sp_Elateridae | 3853 | 1364437 | 440 | 43.08 | 354.1 | 283 | 8623 | 12 |
| G20004_Eudicronychus_rufus_Elateridae | 3458 | 828213 | 291 | 27.91 | 239.5 | 202.5 | 1470 | 7 |
| G20005_Mulsanteus_sp_Elateridae | 3948 | 1327925 | 399 | 41.58 | 336.4 | 280 | 3361 | 7 |
| G20006_Elaterinae_indet_Elateridae | 3964 | 1499010 | 459 | 46.59 | 378.2 | 306.5 | 3479 | 9 |
| G20007_Hapatesus_tropicus_Elateridae | 3844 | 1142966 | 350 | 36.27 | 297.3 | 251 | 2684 | 6 |
| G20008_Eucnemidae_indet_Eucnemidae | 3606 | 1042709 | 359 | 34.42 | 289.2 | 227 | 7544 | 12 |
| G20009_Drapetes_sp_Elateridae | 3822 | 1223326 | 391 | 38.89 | 320.1 | 260 | 3650 | 13 |
| G20010_Chelonarium_sp_Chelonaridae | 3844 | 1021700 | 328 | 32.42 | 265.8 | 212 | 3161 | 14 |
| G20011_Balgus_obconicus_Elateridae | 3743 | 1130979 | 368 | 36.39 | 302.2 | 257 | 2974 | 17 |
| G20012_Tibionema_abdominalis_Elateridae | 3224 | 677752 | 252 | 24.76 | 210.2 | 182 | 2043 | 6 |
| G20019_Eulichas_dudgeoni_Eulichadidae | 3970 | 1457630 | 445 | 44.96 | 367.2 | 295 | 5609 | 20 |
| G20021_Elater_ferrugineus_Elateridae | 3892 | 1500953 | 476 | 47.49 | 385.7 | 315 | 4522 | 10 |
| G20022_Pseudotetralobus_capucinus_Elateridae | 3915 | 1332315 | 413 | 41.83 | 340.3 | 280 | 3439 | 6 |
| G20023_Rostricephalus_vitalis_Elateridae | 3748 | 1042299 | 327 | 33.98 | 278.1 | 237 | 3508 | 14 |
| G20024_Aphileus_lucanoides_Elateridae | 3905 | 1292656 | 397 | 40.45 | 331 | 280 | 3778 | 6 |
| G20025_Cheguevaria_sp_Lampyridae | 3824 | 1073585 | 326 | 33.94 | 280.7 | 240 | 2212 | 12 |
| G20028_Stenocladius_sp_Lampyridae | 3826 | 1227674 | 393 | 38.96 | 320.9 | 269 | 3232 | 6 |
| G20030_Rhagophthalmidae_indet_Rhagophthalmidae | 3625 | 993374 | 331 | 32.4 | 274 | 236 | 2166 | 6 |
| G20031_Rhagophthalmidae_indet_Rhagophthalmidae | 3809 | 1134824 | 355 | 36.76 | 297.9 | 255 | 2178 | 7 |
| G20033_Euanoma_sp_Elateridae | 3926 | 1509918 | 461 | 46.95 | 384.6 | 316.5 | 9517 | 19 |
| G20035_Artematopus_sp_Artematopodidae | 3949 | 1542960 | 480 | 48.26 | 390.7 | 313 | 5727 | 13 |
| G21002_Sphaenelater_lineicollis_Elateridae | 3850 | 1225154 | 382 | 38.76 | 318.2 | 275 | 2052 | 10 |
| G21003_Analastesa_arabica_Elateridae | 3132 | 627974 | 238 | 22.52 | 200.5 | 168.5 | 1894 | 8 |
| G21004_Paulusiella_serraticornis_Elateridae | 3977 | 1498171 | 452 | 46.33 | 376.7 | 307 | 5507 | 8 |
| G21006_Rhipiceridae_indet_Rhipiceridae | 3826 | 1616983 | 512 | 51.62 | 422.6 | 343 | 7658 | 25 |
| G21007_Cneoglossa_sp_Cneoglossidae | 3927 | 1691725 | 531 | 53.52 | 430.8 | 348 | 4449 | 7 |
| G21009_Psilocladus_sp_Lampyridae | 3861 | 1235028 | 386 | 39.21 | 319.9 | 271 | 2226 | 20 |
| G21010_Amydetinae_indet_Lampyridae | 3988 | 1382504 | 413 | 42.66 | 346.7 | 304 | 3604 | 7 |
| G21011_Ototretinae_indet_Lampyridae | 3964 | 1362356 | 404 | 42.17 | 343.7 | 293 | 3270 | 6 |
| G21012_Austroselasia_sp_Elateridae | 3966 | 1309007 | 397 | 40.9 | 330.1 | 276 | 3450 | 10 |
| G21013_Habashaselasia_nekemtensis_Elateridae | 3946 | 1277632 | 387 | 39.81 | 323.8 | 274.5 | 2901 | 7 |
| G21014_Elatichrosis_sp_Elateridae | 3897 | 1266316 | 386 | 39.83 | 324.9 | 270 | 3944 | 10 |
| G21015_Brachypterodrilus_sp_Lampyridae | 3850 | 1151292 | 357 | 35.98 | 299 | 254 | 3531 | 8 |
| G21016_Rhagophthalmidae_indet_Rhagophthalmidae | 3951 | 1394282 | 419 | 43.39 | 352.9 | 298 | 3664 | 7 |
| G21017_Lissominae_indet_Elateridae | 3863 | 1703094 | 540 | 54.07 | 440.9 | 356 | 6018 | 8 |
| G21019_Eucnemidae_indet_Eucnemidae | 3233 | 771479 | 297 | 26.8 | 238.6 | 196 | 2734 | 7 |
| G21023_Ceratogonys_spinicorne_Eucnemidae | 3843 | 1312480 | 420 | 41.27 | 341.5 | 286 | 2681 | 9 |
| G21026_Pyrophorini_indet_Elateridae | 3872 | 1293236 | 400 | 40.76 | 334 | 283 | 3787 | 9 |
| G21027_Elaterinae_indet_Elateridae | 2882 | 571415 | 232 | 21.65 | 198.3 | 168 | 1915 | 12 |
| G21028_Dascilidae_indet_Dascillidae | 3917 | 1723285 | 532 | 54.46 | 440 | 359 | 7997 | 6 |
| G21029_Drilaster_sp_Lampyridae | 3569 | 877254 | 295 | 28.7 | 245.8 | 209 | 1946 | 6 |
| G21030_Psephenidae_indet_Psephenidae | 3948 | 1576569 | 483 | 48.97 | 399.3 | 329 | 7591 | 7 |
| G21031_Rousia_sp_Elateridae | 3955 | 1719519 | 524 | 53.66 | 434.8 | 357 | 6559 | 14 |
| G21035_Anaissus_sp_Elateridae | 3950 | 1459943 | 448 | 45.62 | 369.6 | 310 | 3023 | 6 |
| G21036_Campyloxenus_pyrothorax_Elateridae | 3855 | 1283174 | 400 | 41.26 | 332.9 | 275 | 8664 | 7 |
| G21037_Pityobius_anguinus_Elateridae | 3632 | 1044361 | 349 | 34.61 | 287.5 | 240 | 1852 | 10 |
| G21038_Adelocera_sp_Elateridae | 3836 | 1216992 | 376 | 38.57 | 317.3 | 265 | 2647 | 6 |
| G21039_Psephenidae_indet_Psephenidae | 3967 | 1683348 | 523 | 52.61 | 424.3 | 343 | 4949 | 13 |
| G21040_Adelocera_sp_Elateridae | 3864 | 1123940 | 342 | 35.56 | 290.9 | 247 | 2906 | 9 |
| G21041_Adelocera_sp_Elateridae | 3996 | 1731117 | 519 | 53.68 | 433.2 | 353 | 5886 | 8 |
| G21042_Telegeusis_sp_Omethidae | 3857 | 1304391 | 418 | 41.43 | 338.2 | 277 | 4031 | 6 |
| G22002_Plectonotum_laterale_Cantharidae | 3878 | 1265094 | 386 | 40.12 | 326.2 | 264 | 4411 | 9 |
| G22003_Cerophytum_japonicum_Cerophytiae | 3387 | 696170 | 240 | 25.34 | 205.5 | 175 | 1855 | 9 |
| G22005_Nyctophyxis_ocellatus_Elateridae | 2642 | 979750 | 466 | 41.78 | 370.8 | 314 | 3440 | 10 |
| G22007_Anaspasis_paralella_Elateridae | 2979 | 590340 | 231 | 21.82 | 198.2 | 174 | 1824 | 7 |
| G22009_Jurasai_digitusdei_Jurasaidae | 3726 | 1204511 | 383 | 39.21 | 323.3 | 270 | 4045 | 8 |
| G22015_Protelmidae_indet_Protelmidae | 3892 | 1391051 | 439 | 43.9 | 357.4 | 290 | 2611 | 7 |
| G22020_Lutrochidae_indet_Lutrochidae | 3936 | 1557171 | 489 | 48.93 | 395.6 | 320 | 5678 | 10 |
| G22022_Lutrochidae_indet_Lutrochidae | 3966 | 1716433 | 532 | 53.35 | 432.8 | 354.5 | 6731 | 6 |
| G22024_Elmidae_indet_Elmidae | 3438 | 1443109 | 515 | 50.37 | 419.8 | 338 | 7599 | 8 |
| G22025_Ototretinae_indet_Lampyridae | 4020 | 1500197 | 436 | 46.09 | 373.2 | 320 | 6556 | 6 |
| G22026_Stenocladius_sp_Lampyridae | 3664 | 959132 | 314 | 31.45 | 261.8 | 220 | 2860 | 7 |
| G22027_Lampyrinae_indet_Lampyridae | 3868 | 1491406 | 462 | 47.04 | 385.6 | 314 | 5511 | 9 |
| G19002_Hemiops_sp_Elateridae | 3767 | 998599 | 318 | 31.84 | 265.1 | 221 | 2745 | 6 |
| Heterocerus_fenestratus_Heteroceridae | 3365 | 774647 | 270 | 29.4 | 230.2 | 214 | 933 | 10 |
| Heteromastix_sp_Cantharidae | 3909 | 1687931 | 529 | 53.61 | 431.8 | 356 | 3018 | 13 |
| Hypnoidus_riparius_Elateridae | 3878 | 1724153 | 541 | 55.49 | 444.6 | 367 | 4123 | 7 |
| G19015_Iberobaenia_minuta_Iberobaeniidae | 3751 | 993161 | 309 | 32.06 | 264.8 | 223 | 3434 | 6 |
| Ignelater_luminosus_Elateridae | 3869 | 1724141 | 542 | 54.84 | 445.6 | 365 | 5502 | 15 |
| R19001_Isorhipis_marmottani_Eucnemidae | 3922 | 1688440 | 525 | 54.11 | 430.5 | 353 | 5819 | 10 |
| Konoplatycis_otome_Lycidae | 3516 | 1073278 | 373 | 38.6 | 305.3 | 262 | 7521 | 14 |
| Lamprigera_yunnana_Lampyridae | 3757 | 1682752 | 549 | 54.76 | 447.9 | 359 | 5608 | 41 |
| Lamprohiza_splendidula_Lampyridae | 3673 | 1377826 | 453 | 47.09 | 375.1 | 320 | 3497 | 10 |
| S1_Lampyris_sardiniae_Lampyridae | 3782 | 1639917 | 526 | 54.18 | 433.6 | 357 | 6746 | 22 |
| Leptinotarsa_decemlineata | 4224 | 1943146 | 564 | 56.1 | 460 | 364 | 7360 | 47 |
| Leptolycus_sp_Lycidae | 3797 | 964879 | 303 | 30.55 | 254.1 | 210 | 3637 | 7 |
| Libnetis_sp_Lycidae | 3732 | 1524413 | 498 | 51.37 | 408.5 | 342 | 3467 | 27 |
| Limonius_californicus_Elateridae | 3942 | 1335615 | 393 | 41.73 | 338.8 | 283 | 6560 | 10 |
| R19013_Lissomus_sp_Elateridae | 3841 | 1684384 | 533 | 54.98 | 438.5 | 365 | 3586 | 31 |
| Lucidota_atra_Lampyridae | 3761 | 1567532 | 509 | 52.46 | 416.8 | 346 | 5937 | 6 |
| Lycoprogenthes_sp_Lycidae | 3686 | 1559932 | 518 | 53.22 | 423.2 | 346.5 | 3823 | 9 |
| Lycostomus_kraatzi_Lycidae | 3501 | 1410794 | 494 | 50.87 | 403 | 336 | 4050 | 7 |
| Lygistopterus_sanguineus_Lycidae | 3694 | 1550303 | 506 | 52.47 | 419.7 | 348 | 5712 | 19 |
| Lyropaeus_optabilis_Lycidae | 3529 | 1386414 | 479 | 49.58 | 392.9 | 326 | 5666 | 13 |
| Ponyalis_sp_Lycidae | 3656 | 1532736 | 513 | 52.48 | 419.2 | 349 | 4473 | 9 |
| R20009_Malthodes_sp_Cantharidae | 3864 | 1587345 | 493 | 51.78 | 410.8 | 351 | 3053 | 12 |
| Melanotus_cribricollis_Elateridae | 3683 | 1659300 | 550 | 56.34 | 450.5 | 365 | 5505 | 28 |
| Melanotus_villosus_Elateridae | 3706 | 1586669 | 517 | 53.67 | 428.1 | 359.5 | 4252 | 29 |
| Micronaspis_floridana_Lampyridae | 3555 | 1254726 | 435 | 44.46 | 352.9 | 299 | 2584 | 6 |
| Microphotus_sp_Lampyridae | 3616 | 1422091 | 484 | 48.77 | 393.3 | 326 | 4903 | 6 |
| G19011_Diplophoenicus_sp_Elateridae | 3957 | 1649448 | 504 | 51.57 | 416.8 | 346 | 6417 | 7 |
| Negastrius_arenicola_Elateridae | 3956 | 1740659 | 533 | 55.54 | 440 | 364 | 4372 | 10 |
| Notolioon_sp_Byrrhidae | 3256 | 1184383 | 455 | 47.48 | 363.8 | 313 | 2275 | 8 |
| Omalisus_fontisbellaquei_Elateridae | 3918 | 1423666 | 429 | 44.77 | 363.4 | 308 | 7584 | 8 |
| Onthophagus_taurus | 4224 | 2342918 | 687 | 67.65 | 554.7 | 423 | 24150 | 55 |
| R19005_Ototretinae_indet_Lampyridae | 3852 | 1687405 | 533 | 55.02 | 438.1 | 358 | 8354 | 8 |
| R20013_Ototretinae_indet_Lampyridae | 3897 | 1671957 | 525 | 53.68 | 429 | 356 | 3122 | 9 |
| G19008_Oxynopterus_sp_Elateridae | 3957 | 1654983 | 510 | 51.63 | 418.2 | 345 | 4617 | 6 |
| G19005_Pachyderes_sp_Elateridae | 3959 | 1429498 | 432 | 44.76 | 361.1 | 308 | 4707 | 6 |
| G19006_Parasaphes_sp_Elateridae | 3941 | 1601702 | 492 | 50.25 | 406.4 | 336 | 5241 | 6 |
| G19004_Pectocera_sp_Elateridae | 3984 | 1428198 | 429 | 44.25 | 358.5 | 302 | 3552 | 15 |
| Phausis_reticulata_Lampyridae | 3733 | 1550112 | 503 | 52.38 | 415.2 | 346 | 5029 | 5 |
| Phengodes_sp_B_Phengodidae | 3675 | 1519924 | 504 | 52.08 | 413.6 | 342 | 6458 | 12 |
| Phengodes_sp_A_Phengodidae | 3928 | 1372647 | 419 | 42.9 | 349.5 | 295 | 4373 | 13 |
| Photinus_australis_Lampyridae | 3640 | 1398721 | 477 | 48.05 | 384.3 | 328.5 | 3758 | 7 |
| Photinus_carolinus_Lampyridae | 3657 | 1422259 | 481 | 49 | 388.9 | 326 | 4899 | 7 |
| Photinus_macdermotti_Lampyridae | 3521 | 1410974 | 511 | 49.28 | 400.7 | 329 | 5622 | 6 |
| Photinus_pyralis_Lampyridae | 3894 | 1723963 | 533 | 55.4 | 442.7 | 360 | 7758 | 27 |
| Photinus_scintillans_Lampyridae | 3796 | 1629491 | 525 | 54.1 | 429.3 | 352 | 4902 | 8 |
| Photuris_frontalis_Lampyridae | 3733 | 1579266 | 517 | 53.07 | 423.1 | 352 | 6558 | 9 |
| Photuris_sp_Lampyridae | 3612 | 1482333 | 517 | 51.39 | 410.4 | 339 | 5972 | 6 |
| Phrixothrix_hirtus_Phengodidae | 2878 | 769634 | 352 | 35.08 | 267.4 | 217 | 4278 | 23 |
| A01544_Plastocerus_angulosus_Elateridae | 3960 | 1578007 | 485 | 49.35 | 398.5 | 328 | 7720 | 6 |
| Platerodrilus_sp_Lycidae | 3283 | 1248803 | 462 | 48.52 | 380.4 | 318 | 5550 | 8 |
| Plateros_sp_Lycidae | 3661 | 1576875 | 523 | 53.78 | 430.7 | 353 | 3576 | 6 |
| Ponyalis_quadricollis_Lycidae | 3563 | 1438693 | 487 | 50.41 | 403.8 | 336 | 4139 | 37 |
| R19004_Ptilodactylidae_indet_Ptilodactylidae | 3978 | 1792156 | 549 | 56.83 | 450.5 | 371 | 6210 | 7 |
| Ptosima_undecimmaculata_Buprestidae | 3445 | 1308534 | 461 | 48.06 | 379.8 | 318 | 6634 | 12 |
| Pyractomena_borealis_Lampyridae | 3752 | 1587762 | 518 | 53.18 | 423.2 | 351 | 6771 | 13 |
| Pyrearinus_fragilis_Elateridae | 3880 | 1783896 | 558 | 57.71 | 459.8 | 375.5 | 6699 | 11 |
| Pyrocoelia_pectoralis_Lampyridae | 3767 | 1528898 | 499 | 50.66 | 405.9 | 336 | 6917 | 9 |
| R18040_Mangkutanus_sp_Lycidae | 3775 | 1578968 | 503 | 52.1 | 418.3 | 346 | 7628 | 7 |
| R20002_Agrypninae_indet_Elateridae | 3780 | 1585525 | 510 | 52.88 | 419.5 | 358 | 2355 | 33 |
| R20003_Athous_sp_Elateridae | 3945 | 1725229 | 527 | 54.41 | 437.3 | 362 | 3644 | 16 |
| R20004_Selatosomus_aeneus_Elateridae | 3701 | 1442970 | 474 | 49.36 | 389.9 | 343 | 2208 | 6 |
| R20005_Ampedus_sineatus_Elateridae | 3863 | 1726848 | 544 | 56.24 | 447 | 372 | 3793 | 14 |
| R20006_Ampedus_cinnabarinu_Elateridaes | 3898 | 1737698 | 537 | 55.76 | 445.8 | 371 | 3152 | 8 |
| R20007_Anostirus_purpureus_Elateridae | 3877 | 1708896 | 534 | 55.23 | 440.8 | 365 | 7082 | 10 |
| R20008_Agriotes_sp_Elateridae | 3915 | 1735006 | 536 | 55.05 | 443.2 | 373 | 5512 | 8 |
| R20010_Elaterinae_indet_Elateridae | 3898 | 1676223 | 519 | 53.38 | 430 | 359 | 3258 | 11 |
| R20011_Alaus_myops_Elateridae | 3361 | 1103552 | 398 | 42.68 | 328.3 | 283 | 5145 | 10 |
| R20014_Limnius_perrisi_Elmidae | 3948 | 1765563 | 543 | 56.22 | 447.2 | 372 | 2660 | 11 |
| R20016_Cardiophorus_discicollis_Elateridae | 3931 | 1706500 | 523 | 54.29 | 434.1 | 358 | 3155 | 6 |
| R20017_Eubria_palustris_Psephenidae | 3938 | 1739194 | 535 | 55.58 | 441.6 | 365 | 3579 | 8 |
| R20018_Phosphaenus_hemipterus_Lampyridae | 3898 | 1679238 | 519 | 53.88 | 430.8 | 354 | 3844 | 6 |
| R20019_Dascillus_cervinus_Dascillidae | 3959 | 1720122 | 529 | 54.55 | 434.5 | 360 | 2914 | 9 |
| R20022_Lampyris_noctiluca_Lampyridae | 3914 | 1686003 | 524 | 53.72 | 430.8 | 360.5 | 3035 | 31 |
| Anischia_sp_Eucnemidae | 3939 | 1549723 | 476 | 48.94 | 393.4 | 322 | 6243 | 31 |
| Platycrepidius_sp_Elateridae | 3996 | 1713414 | 523 | 53.1 | 428.8 | 353 | 8611 | 9 |
| Lamellipalpus_pacholatkoi_Lampyridae | 4006 | 1495885 | 435 | 46.13 | 373.4 | 320 | 3437 | 5 |
| Byrrhinus_sp_Limnichidae | 3949 | 1765721 | 550 | 55.77 | 447.1 | 362 | 5954 | 11 |
| Epilichas_sp_Ptilodactylidae | 3615 | 908241 | 304 | 30.29 | 251.2 | 203 | 3438 | 12 |
| Rhagonycha_fulva_Cantharidae | 3538 | 1410255 | 485 | 50.11 | 398.6 | 342 | 3374 | 7 |
| Rhagopthalmus_sp_Rhagophthalmidae | 3769 | 1659542 | 536 | 55.29 | 440.3 | 358 | 8290 | 8 |
| S39_Limnichidae_indet_Limnichidae | 3740 | 1529483 | 504 | 51.09 | 409 | 341 | 5081 | 6 |
| Sandalus_niger_Rhipiceridae | 3641 | 1460667 | 487 | 50.31 | 401.2 | 337 | 7792 | 6 |
| Scarelus_anthracinus_Lycidae | 3744 | 1571772 | 507 | 52.28 | 419.8 | 344 | 7145 | 24 |
| G19009_Semiotus_sp_Elateridae | 3994 | 1626793 | 497 | 50.21 | 407.3 | 327.5 | 5940 | 8 |
| R19009_Silis_sp_Cantharidae | 3947 | 1742656 | 535 | 55.33 | 441.5 | 365 | 5718 | 8 |
| Sinopyrophorus_schimmeli_Sinopyrophoridae | 3845 | 1315367 | 424 | 42.44 | 342.1 | 277 | 2675 | 10 |
| G20013_Sinopyrophorus_sp_Sinopyrophoridae | 3854 | 1245249 | 403 | 39.57 | 323.1 | 259 | 3582 | 7 |
| Taphes_brevicollis_Lycidae | 3710 | 1566805 | 518 | 52.97 | 422.3 | 351 | 5725 | 26 |
| G19001_Sinelater_perroti_Elateridae | 3905 | 1379108 | 430 | 43.77 | 353.2 | 290 | 4703 | 9 |
| Tribolium_castaneum | 4224 | 2479551 | 746 | 71.59 | 587 | 452.5 | 8743 | 64 |
| Trixagus_carinifrons_Throscidae | 3592 | 1147846 | 384 | 40.06 | 319.6 | 295 | 2586 | 10 |

| Taxon _name_Family | AA outliers removed | | | | | | | |
|---|---|---|---|---|---|---|---|---|
| | no_genes | total_no_aa | N50 | percent_determined | average_length | median_length | max_length | min_length |
| A4488_Throscidae_indet_Throscidae | 3782 | 1140068 | 359 | 38.33 | 301.4 | 250 | 5615 | 6 |
| Abscondita_terminalis_Lampyridae | 3729 | 1636896 | 529 | 56.12 | 439 | 357 | 5504 | 29 |
| Agrilus_planipennis_Buprestidae | 4137 | 2306923 | 711 | 69.11 | 557.6 | 424 | 9009 | 54 |
| Agrilus_zanthoxylumi_Buprestidae | 3937 | 1928361 | 613 | 62.53 | 489.8 | 389 | 5701 | 15 |
| Agrypnus_murinus_Elateridae | 3640 | 1407228 | 474 | 49.8 | 386.6 | 328 | 4104 | 24 |
| Anoplophora_glabripennis | 4152 | 2240709 | 669 | 66.92 | 539.7 | 411 | 19530 | 59 |
| Anorus_parvicollis_Dascillidae | 3690 | 1447963 | 478 | 50.63 | 392.4 | 326.5 | 7875 | 7 |
| Antennolycus_constrictus_Lycidae | 3724 | 1551832 | 511 | 53.64 | 416.7 | 341 | 5007 | 31 |
| Aquatica_ficta_Lampyridae | 3690 | 1428169 | 478 | 49.69 | 387 | 324 | 4749 | 25 |
| Asymmetricata_circumdata_Lampyridae | 3589 | 1457625 | 493 | 52.61 | 406.1 | 339 | 5726 | 12 |
| Aspisoma_lineatum_Lampyridae | 3896 | 1711080 | 532 | 56.22 | 439.2 | 359 | 7029 | 16 |
| Bicellonycha_wickershamorum_Lampyridae | 3578 | 1351489 | 466 | 48.63 | 377.7 | 318 | 5286 | 6 |
| Brachypsectra_fulva_Brachypsectridae | 3548 | 1251881 | 434 | 46.28 | 352.8 | 316 | 3293 | 7 |
| Byrrhus_pilula_Byrrhidae | 3299 | 1169248 | 433 | 46.21 | 354.4 | 307 | 3614 | 30 |
| R19006_Callirhipidae_indet_Callirhipidae | 3676 | 1514059 | 509 | 53.12 | 411.9 | 334.5 | 7663 | 31 |
| Calopteron_sp_Lycidae | 3652 | 1570872 | 522 | 55.28 | 430.1 | 354 | 5603 | 8 |
| Capnodis_tenebrionis_Buprestidae | 3744 | 1650122 | 537 | 55.67 | 440.7 | 363 | 5917 | 16 |
| R19003_Dicronychus_rubripes_Elateridae | 3879 | 1723191 | 535 | 57.47 | 444.2 | 364 | 8322 | 37 |
| Cautires_communis_Lycidae | 3696 | 1044441 | 330 | 35.24 | 282.6 | 239.5 | 3254 | 11 |
| G19010_Cebrio_sp_Elateridae | 3458 | 879617 | 306 | 30.9 | 254.4 | 218 | 3373 | 6 |
| Cerophytum_elateroides_Cerophytiae | 3110 | 587356 | 223 | 23.66 | 188.9 | 158 | 1359 | 7 |
| Chauliognathus_opacus_Cantharidae | 3571 | 929758 | 312 | 33.13 | 260.4 | 213 | 5468 | 8 |
| Conderis_signicollis_Lycidae | 3653 | 1415954 | 478 | 49.54 | 387.6 | 320 | 7547 | 35 |
| Elaterinae_indet_Elateridae | 3902 | 1802073 | 555 | 58.94 | 461.8 | 378 | 6080 | 9 |
| Dendroctonus_ponderosae | 3989 | 2158659 | 673 | 66.36 | 541.2 | 425 | 7334 | 50 |
| R19012_Denticollis_sp_Elateridae | 3895 | 1703701 | 526 | 56.45 | 437.4 | 365 | 4323 | 40 |
| Dexoris_ruzzieri_Lycidae | 3773 | 1001200 | 311 | 32.78 | 265.4 | 226 | 3495 | 20 |
| Dictyoptera_sp_Lycidae | 3336 | 862857 | 328 | 33.85 | 258.7 | 211 | 3363 | 7 |
| Dihammatus_sp_Lycidae | 3711 | 1525974 | 505 | 52.69 | 411.2 | 342 | 7122 | 10 |
| Dilophotes_sp_Lycidae | 3701 | 1537663 | 504 | 53.18 | 415.5 | 346 | 3641 | 7 |
| Distremocephalus_sp_Phengodidae | 3610 | 1365023 | 461 | 49.55 | 378.1 | 330 | 2853 | 11 |
| Drilonius_sp_Omethidae | 3850 | 1686057 | 529 | 55.95 | 437.9 | 362 | 6144 | 20 |
| G18004_Drilus_mauritanicus_Elateridae | 3957 | 1529927 | 469 | 49 | 386.6 | 318 | 6557 | 7 |
| Drilus_concolor_Elateridae | 2974 | 851810 | 357 | 37.5 | 286.4 | 242 | 3350 | 19 |
| Dryops_sp_Dryopidae | 3676 | 1502511 | 500 | 52.88 | 408.7 | 343 | 3112 | 17 |
| R19010_Quasimus_sp_Elateridae | 3898 | 1701714 | 528 | 56.2 | 436.6 | 359 | 5064 | 6 |
| R19011_Agrypninae_indet_Elateridae | 3836 | 1676651 | 533 | 56.13 | 437.1 | 365 | 7423 | 8 |
| G19012_Penia_sp_Elateridae | 3912 | 1420906 | 436 | 45.79 | 363.2 | 309 | 4774 | 6 |
| G19013_Dromaeolinae indet_Eucnemidae | 3826 | 1437183 | 459 | 47 | 375.6 | 307 | 4896 | 21 |
| Ellychnia_sp_Lampyridae | 3702 | 1489615 | 494 | 51 | 402.4 | 333 | 4657 | 8 |
| Eucnemis_capucina_Eucnemidae | 3866 | 1622404 | 512 | 53.07 | 419.7 | 342 | 4781 | 28 |
| G19007_Eudicronychus_sp_Elateridae | 3401 | 860004 | 307 | 30.43 | 252.9 | 212 | 2219 | 10 |
| Eurypogon_sp_Artematopodidae | 3610 | 1219700 | 404 | 44.1 | 337.9 | 314 | 1444 | 12 |
| G20001_Octinodes_sp_Elateridae | 3907 | 1412327 | 439 | 45.51 | 361.5 | 295 | 3634 | 31 |
| G20002_Pyrearinus_sp_Elateridae | 3969 | 1629009 | 497 | 51.75 | 410.4 | 339 | 4726 | 21 |
| G20003_Cussolenis_sp_Elateridae | 3824 | 1356809 | 440 | 44.36 | 354.8 | 284 | 8623 | 12 |
| G20004_Eudicronychus_rufus_Elateridae | 3354 | 805519 | 292 | 28.81 | 240.2 | 203 | 1470 | 7 |
| G20005_Mulsanteus_sp_Elateridae | 3923 | 1321378 | 399 | 42.69 | 336.8 | 280 | 3361 | 7 |
| G20006_Elaterinae_indet_Elateridae | 3935 | 1491250 | 460 | 47.82 | 379 | 308 | 3479 | 9 |
| G20007_Hapatesus_tropicus_Elateridae | 3807 | 1135565 | 350 | 37.39 | 298.3 | 252 | 2684 | 6 |
| G20008_Eucnemidae_indet_Eucnemidae | 3507 | 1020317 | 360 | 35.45 | 290.9 | 229 | 7544 | 12 |
| G20009_Drapetes_sp_Elateridae | 3789 | 1215338 | 393 | 39.94 | 320.8 | 260 | 3650 | 13 |
| G20010_Chelonarium_sp_Chelonaridae | 3768 | 1007137 | 330 | 33.52 | 267.3 | 213 | 3161 | 14 |
| G20011_Balgus_obconicus_Elateridae | 3676 | 1113614 | 368 | 37.44 | 302.9 | 258 | 2968 | 17 |
| G20012_Tibionema_abdominalis_Elateridae | 3043 | 645530 | 254 | 25.74 | 212.1 | 184 | 2043 | 6 |
| G20019_Eulichas_dudgeoni_Eulichadidae | 3947 | 1449694 | 445 | 46.14 | 367.3 | 295 | 5609 | 20 |
| G20021_Elater_ferrugineus_Elateridae | 3863 | 1495120 | 477 | 48.77 | 387 | 316 | 4522 | 10 |
| G20022_Pseudotetralobus_capucinus_Elateridae | 3878 | 1322376 | 414 | 42.94 | 341 | 281 | 3439 | 6 |
| G20023_Rostricephalus_vitalis_Elateridae | 3671 | 1026295 | 328 | 35.13 | 279.6 | 239 | 3508 | 15 |
| G20024_Aphileus_lucanoides_Elateridae | 3866 | 1282626 | 399 | 41.53 | 331.8 | 280 | 3778 | 6 |
| G20025_Cheguevaria_sp_Lampyridae | 3792 | 1066629 | 326 | 34.88 | 281.3 | 240 | 2212 | 25 |
| G20028_Stenocladius_sp_Lampyridae | 3783 | 1218715 | 394 | 39.96 | 322.2 | 270 | 3232 | 6 |
| G20030_Rhagophthalmidae_indet_Rhagophthalmidae | 3568 | 981195 | 331 | 33.43 | 275 | 236 | 2165 | 6 |
| G20031_Rhagophthalmidae_indet_Rhagophthalmidae | 3759 | 1123194 | 356 | 37.92 | 298.8 | 255 | 2178 | 7 |
| G20033_Euanoma_sp_Elateridae | 3892 | 1500707 | 461 | 48.29 | 385.6 | 317 | 9517 | 19 |
| G20035_Artematopus_sp_Artematopodidae | 3917 | 1535921 | 480 | 49.46 | 392.1 | 313 | 5727 | 24 |
| G21002_Sphaenelater_lineicollis_Elateridae | 3810 | 1215530 | 382 | 39.93 | 319 | 276 | 2052 | 10 |
| G21003_Analastesa_arabica_Elateridae | 2869 | 571451 | 236 | 22.71 | 199.2 | 168 | 1787 | 8 |
| G21004_Paulusiella_serraticornis_Elateridae | 3950 | 1488077 | 453 | 47.45 | 376.7 | 307 | 5507 | 8 |
| G21006_Rhipiceridae_indet_Rhipiceridae | 3769 | 1600314 | 515 | 53.14 | 424.6 | 345 | 7658 | 25 |
| G21007_Cneoglossa_sp_Cneoglossidae | 3901 | 1685601 | 532 | 54.87 | 432.1 | 349 | 4449 | 7 |
| G21009_Psilocladus_sp_Lampyridae | 3829 | 1225460 | 387 | 40.23 | 320 | 271 | 2226 | 28 |
| G21010_Amydetinae_indet_Lampyridae | 3960 | 1375233 | 413 | 43.85 | 347.3 | 305 | 3604 | 7 |
| G21011_Ototretinae_indet_Lampyridae | 3944 | 1357074 | 404 | 43.26 | 344.1 | 294 | 3270 | 6 |
| G21012_Austroselasia_sp_Elateridae | 3928 | 1296717 | 397 | 41.98 | 330.1 | 276.5 | 3450 | 10 |
| G21013_Habashaselasia_nekemtensis_Elateridae | 3910 | 1267010 | 387 | 40.81 | 324 | 275 | 2901 | 7 |
| G21014_Elatichrosis_sp_Elateridae | 3870 | 1258913 | 387 | 40.86 | 325.3 | 270 | 3944 | 10 |
| G21015_Brachypterodrilus_sp_Lampyridae | 3798 | 1139155 | 357 | 36.91 | 299.9 | 255 | 3531 | 8 |
| G21016_Rhagophthalmidae_indet_Rhagophthalmidae | 3931 | 1388536 | 419 | 44.57 | 353.2 | 299 | 3664 | 7 |
| G21017_Lissominae_indet_Elateridae | 3842 | 1696945 | 540 | 55.51 | 441.7 | 357 | 6018 | 8 |
| G21019_Eucnemidae_indet_Eucnemidae | 3025 | 723309 | 299 | 27.24 | 239.1 | 196 | 2733 | 7 |
| G21023_Ceratogonys_spinicorne_Eucnemidae | 3778 | 1297361 | 422 | 42.41 | 343.4 | 289 | 2681 | 9 |
| G21026_Pyrophorini_indet_Elateridae | 3832 | 1281763 | 401 | 41.83 | 334.5 | 283 | 3787 | 9 |
| G21027_Elaterinae_indet_Elateridae | 2665 | 527902 | 232 | 22.19 | 198.1 | 169 | 1915 | 12 |
| G21028_Dascilidae_indet_Dascillidae | 3894 | 1717686 | 532 | 55.87 | 441.1 | 360 | 7997 | 6 |
| G21029_Drilaster_sp_Lampyridae | 3463 | 850270 | 293 | 29.35 | 245.5 | 209 | 1944 | 6 |
| G21030_Psephenidae_indet_Psephenidae | 3906 | 1565209 | 484 | 50.38 | 400.7 | 331.5 | 7591 | 7 |
| G21031_Rousia_sp_Elateridae | 3935 | 1715624 | 524 | 55.1 | 436 | 358 | 6559 | 37 |
| G21035_Anaissus_sp_Elateridae | 3924 | 1451857 | 448 | 46.89 | 370 | 310 | 3023 | 6 |
| G21036_Campyloxenus_pyrothorax_Elateridae | 3814 | 1273100 | 400 | 42.38 | 333.8 | 276 | 8664 | 7 |
| G21037_Pityobius_anguinus_Elateridae | 3486 | 1007787 | 349 | 35.74 | 289.1 | 241 | 1852 | 10 |
| G21038_Adelocera_sp_Elateridae | 3790 | 1203924 | 378 | 39.6 | 317.7 | 265 | 2647 | 6 |
| G21039_Psephenidae_indet_Psephenidae | 3937 | 1675659 | 523 | 54.05 | 425.6 | 345 | 4949 | 13 |
| G21040_Adelocera_sp_Elateridae | 3810 | 1109856 | 342 | 36.66 | 291.3 | 247 | 2906 | 9 |
| G21041_Adelocera_sp_Elateridae | 3982 | 1726874 | 519 | 55.05 | 433.7 | 354 | 5872 | 8 |
| G21042_Telegeusis_sp_Omethidae | 3804 | 1289375 | 418 | 42.58 | 339 | 277.5 | 4031 | 6 |
| G22002_Plectonotum_laterale_Cantharidae | 3817 | 1251515 | 388 | 41.24 | 327.9 | 265 | 4411 | 9 |
| G22003_Cerophytum_japonicum_Cerophytiae | 3260 | 674752 | 242 | 26.21 | 207 | 177 | 1855 | 9 |
| G22005_Nyctophyxis_ocellatus_Elateridae | 2578 | 962841 | 470 | 43.18 | 373.5 | 315.5 | 3440 | 10 |
| G22007_Anaspasis_paralella_Elateridae | 2810 | 559475 | 232 | 22.72 | 199.1 | 174 | 1823 | 7 |
| G22009_Jurasai_digitusdei_Jurasaidae | 3637 | 1181868 | 385 | 40.29 | 325 | 272 | 4045 | 8 |
| G22015_Protelmidae_indet_Protelmidae | 3845 | 1379092 | 440 | 45.01 | 358.7 | 290 | 2611 | 7 |
| G22020_Lutrochidae_indet_Lutrochidae | 3908 | 1551515 | 490 | 50.2 | 397 | 321 | 5678 | 10 |
| G22022_Lutrochidae_indet_Lutrochidae | 3937 | 1709261 | 532 | 54.86 | 434.2 | 355 | 6731 | 6 |
| G22024_Elmidae_indet_Elmidae | 3411 | 1437139 | 517 | 51.77 | 421.3 | 340 | 7599 | 8 |
| G22025_Ototretinae_indet_Lampyridae | 4004 | 1495746 | 436 | 47.36 | 373.6 | 320.5 | 6556 | 6 |
| G22026_Stenocladius_sp_Lampyridae | 3579 | 937675 | 313 | 32.22 | 262 | 220 | 2860 | 7 |
| G22027_Lampyrinae_indet_Lampyridae | 3844 | 1483552 | 461 | 48.24 | 385.9 | 314 | 5511 | 9 |
| G19002_Hemiops_sp_Elateridae | 3689 | 981824 | 319 | 32.86 | 266.1 | 222 | 2745 | 6 |
| Heterocerus_fenestratus_Heteroceridae | 3246 | 753580 | 273 | 30.92 | 232.2 | 215 | 933 | 10 |
| Heteromastix_sp_Cantharidae | 3897 | 1684018 | 529 | 54.93 | 432.1 | 356 | 3018 | 13 |
| Hypnoidus_riparius_Elateridae | 3869 | 1722331 | 541 | 56.98 | 445.2 | 367 | 4112 | 7 |
| G19015_Iberobaenia_minuta_Iberobaeniidae | 3676 | 978176 | 310 | 33.08 | 266.1 | 224 | 3434 | 6 |
| Ignelater_luminosus_Elateridae | 3855 | 1720747 | 544 | 56.37 | 446.4 | 365 | 5502 | 15 |
| R19001_Isorhipis_marmottani_Eucnemidae | 3896 | 1680670 | 527 | 55.62 | 431.4 | 354 | 5819 | 10 |
| Konoplatycis_otome_Lycidae | 3477 | 1065994 | 374 | 39.9 | 306.6 | 264 | 7521 | 14 |
| Lamprigera_yunnana_Lampyridae | 3737 | 1678807 | 549 | 56.14 | 449.2 | 362 | 5608 | 41 |
| Lamprohiza_splendidula_Lampyridae | 3654 | 1373345 | 453 | 48.41 | 375.8 | 320 | 3497 | 10 |
| S1_Lampyris_sardiniae_Lampyridae | 3764 | 1634178 | 527 | 55.85 | 434.2 | 358 | 6746 | 22 |
| Leptinotarsa_decemlineata | 4042 | 1888004 | 570 | 57.47 | 467.1 | 369 | 7360 | 47 |
| Leptolycus_sp_Lycidae | 3730 | 949224 | 303 | 31.41 | 254.5 | 210 | 3637 | 7 |
| Libnetis_sp_Lycidae | 3710 | 1519255 | 498 | 52.82 | 409.5 | 343 | 3450 | 27 |
| Limonius_californicus_Elateridae | 3916 | 1327711 | 393 | 42.93 | 339 | 284 | 6560 | 10 |
| R19013_Lissomus_sp_Elateridae | 3827 | 1679935 | 534 | 56.5 | 439 | 365 | 3574 | 38 |
| Lucidota_atra_Lampyridae | 3753 | 1565213 | 509 | 53.75 | 417.1 | 346 | 5937 | 6 |
| Lycoprogenthes_sp_Lycidae | 3678 | 1556469 | 517 | 54.56 | 423.2 | 346.5 | 3823 | 9 |
| Lycostomus_kraatzi_Lycidae | 3459 | 1399812 | 494 | 52.57 | 404.7 | 338 | 4050 | 7 |
| Lygistopterus_sanguineus_Lycidae | 3678 | 1546075 | 506 | 53.88 | 420.4 | 348.5 | 5711 | 19 |
| Lyropaeus_optabilis_Lycidae | 3503 | 1381270 | 481 | 51.16 | 394.3 | 328 | 5666 | 13 |
| Ponyalis_sp_Lycidae | 3640 | 1527392 | 513 | 54.05 | 419.6 | 349 | 4473 | 9 |
| R20009_Malthodes_sp_Cantharidae | 3840 | 1580267 | 493 | 53.2 | 411.5 | 351 | 3053 | 12 |
| Melanotus_cribricollis_Elateridae | 3673 | 1656067 | 550 | 57.8 | 450.9 | 366 | 5505 | 28 |
| Melanotus_villosus_Elateridae | 3690 | 1580880 | 517 | 55.29 | 428.4 | 360 | 4252 | 29 |
| Micronaspis_floridana_Lampyridae | 3532 | 1249511 | 436 | 45.82 | 353.8 | 300 | 2584 | 6 |
| Microphotus_sp_Lampyridae | 3580 | 1414150 | 485 | 50.44 | 395 | 327 | 4903 | 6 |
| G19011_Diplophoenicus_sp_Elateridae | 3939 | 1644699 | 504 | 53.08 | 417.5 | 347 | 6417 | 7 |
| Negastrius_arenicola_Elateridae | 3952 | 1739673 | 533 | 57.06 | 440.2 | 364.5 | 4370 | 10 |
| Notolioon_sp_Byrrhidae | 3218 | 1173852 | 457 | 49.46 | 364.8 | 314 | 2275 | 8 |
| Omalisus_fontisbellaquei_Elateridae | 3890 | 1416260 | 429 | 45.89 | 364.1 | 309 | 7584 | 8 |
| Onthophagus_taurus | 4114 | 2298140 | 691 | 69.17 | 558.6 | 425 | 24150 | 55 |
| R19005_Ototretinae_indet_Lampyridae | 3836 | 1683952 | 534 | 56.6 | 439 | 358.5 | 8354 | 8 |
| R20013_Ototretinae_indet_Lampyridae | 3890 | 1668677 | 525 | 55.04 | 429 | 355.5 | 3122 | 9 |
| G19008_Oxynopterus_sp_Elateridae | 3934 | 1649336 | 511 | 53.04 | 419.3 | 345.5 | 4617 | 6 |
| G19005_Pachyderes_sp_Elateridae | 3934 | 1423224 | 432 | 45.96 | 361.8 | 308 | 4707 | 6 |
| G19006_Parasaphes_sp_Elateridae | 3921 | 1597693 | 492 | 51.64 | 407.5 | 337 | 5239 | 6 |
| G19004_Pectocera_sp_Elateridae | 3951 | 1418613 | 430 | 45.55 | 359.1 | 303 | 3552 | 31 |
| Phausis_reticulata_Lampyridae | 3713 | 1544535 | 504 | 53.79 | 416 | 346 | 5029 | 5 |
| Phengodes_sp_B_Phengodidae | 3662 | 1516105 | 505 | 53.63 | 414 | 342 | 6458 | 12 |
| Phengodes_sp_A_Phengodidae | 3885 | 1359535 | 419 | 44.05 | 349.9 | 295 | 4373 | 13 |
| Photinus_australis_Lampyridae | 3606 | 1392992 | 478 | 49.9 | 386.3 | 330 | 3758 | 7 |
| Photinus_carolinus_Lampyridae | 3634 | 1417515 | 482 | 50.55 | 390.1 | 326.5 | 4899 | 7 |
| Photinus_macdermotti_Lampyridae | 3473 | 1401982 | 511 | 51.31 | 403.7 | 331 | 5622 | 6 |
| Photinus_pyralis_Lampyridae | 3886 | 1721254 | 533 | 56.8 | 442.9 | 360 | 7758 | 27 |
| Photinus_scintillans_Lampyridae | 3789 | 1624993 | 525 | 55.51 | 428.9 | 352 | 4902 | 8 |
| Photuris_frontalis_Lampyridae | 3722 | 1576366 | 517 | 54.52 | 423.5 | 352 | 6558 | 9 |
| Photuris_sp_Lampyridae | 3586 | 1477303 | 517 | 52.96 | 412 | 341 | 5972 | 6 |
| Phrixothrix_hirtus_Phengodidae | 2828 | 759138 | 354 | 36.56 | 268.4 | 217 | 4278 | 23 |
| A01544_Plastocerus_angulosus_Elateridae | 3943 | 1572643 | 485 | 50.58 | 398.8 | 328 | 7720 | 6 |
| Platerodrilus_sp_Lycidae | 3252 | 1239402 | 463 | 49.94 | 381.1 | 318 | 5548 | 8 |
| Plateros_sp_Lycidae | 3646 | 1572819 | 524 | 55.31 | 431.4 | 353 | 3576 | 6 |
| Ponyalis_quadricollis_Lycidae | 3546 | 1433140 | 488 | 52.09 | 404.2 | 336 | 4139 | 37 |
| R19004_Ptilodactylidae_indet_Ptilodactylidae | 3964 | 1787219 | 550 | 58.34 | 450.9 | 372 | 6210 | 7 |
| Ptosima_undecimmaculata_Buprestidae | 3388 | 1294231 | 465 | 49.77 | 382 | 320 | 6634 | 12 |
| Pyractomena_borealis_Lampyridae | 3743 | 1585280 | 518 | 54.51 | 423.5 | 351 | 6771 | 13 |
| Pyrearinus_fragilis_Elateridae | 3871 | 1781721 | 558 | 59.24 | 460.3 | 376 | 6699 | 24 |
| Pyrocoelia_pectoralis_Lampyridae | 3755 | 1526082 | 499 | 52.08 | 406.4 | 336 | 6909 | 9 |
| R18040_Mangkutanus_sp_Lycidae | 3749 | 1570104 | 504 | 53.57 | 418.8 | 346 | 7628 | 7 |
| R20002_Agrypninae_indet_Elateridae | 3763 | 1580240 | 511 | 54.6 | 419.9 | 359 | 2355 | 33 |
| R20003_Athous_sp_Elateridae | 3925 | 1718654 | 527 | 56.23 | 437.9 | 362 | 3643 | 23 |
| R20004_Selatosomus_aeneus_Elateridae | 3672 | 1434651 | 475 | 51.07 | 390.7 | 344 | 2208 | 6 |
| R20005_Ampedus_sineatus_Elateridae | 3857 | 1725256 | 544 | 57.69 | 447.3 | 372 | 3793 | 20 |
| R20006_Ampedus_cinnabarinu_Elateridaes | 3884 | 1733974 | 537 | 57.19 | 446.4 | 372 | 3152 | 8 |
| R20007_Anostirus_purpureus_Elateridae | 3864 | 1704518 | 534 | 56.96 | 441.1 | 365 | 7078 | 34 |
| R20008_Agriotes_sp_Elateridae | 3899 | 1729070 | 536 | 56.64 | 443.5 | 373 | 5512 | 8 |
| R20010_Elaterinae_indet_Elateridae | 3884 | 1673022 | 519 | 54.77 | 430.7 | 360 | 3258 | 11 |
| R20011_Alaus_myops_Elateridae | 3319 | 1094510 | 400 | 44.51 | 329.8 | 284 | 5145 | 10 |
| R20014_Limnius_perrisi_Elmidae | 3932 | 1761955 | 544 | 57.78 | 448.1 | 373 | 2660 | 20 |
| R20016_Cardiophorus_discicollis_Elateridae | 3922 | 1703110 | 523 | 55.73 | 434.2 | 358 | 3155 | 6 |
| R20017_Eubria_palustris_Psephenidae | 3923 | 1735629 | 535 | 56.98 | 442.4 | 365 | 3578 | 8 |
| R20018_Phosphaenus_hemipterus_Lampyridae | 3882 | 1677007 | 520 | 55.36 | 432 | 355 | 3844 | 6 |
| R20019_Dascillus_cervinus_Dascillidae | 3945 | 1716513 | 529 | 56.2 | 435.1 | 361 | 2914 | 9 |
| R20022_Lampyris_noctiluca_Lampyridae | 3906 | 1682987 | 524 | 55.15 | 430.9 | 361 | 3035 | 31 |
| Anischia_sp_Eucnemidae | 3884 | 1536639 | 479 | 50.25 | 395.6 | 324 | 6243 | 31 |
| Platycrepidius_sp_Elateridae | 3979 | 1708980 | 525 | 54.43 | 429.5 | 353 | 8611 | 9 |
| Lamellipalpus_pacholatkoi_Lampyridae | 3995 | 1491034 | 435 | 47.24 | 373.2 | 320 | 3437 | 5 |
| Byrrhinus_sp_Limnichidae | 3924 | 1759804 | 551 | 57.21 | 448.5 | 363 | 5954 | 11 |
| Epilichas_sp_Ptilodactylidae | 3503 | 882665 | 304 | 31.19 | 252 | 203 | 3428 | 12 |
| Rhagonycha_fulva_Cantharidae | 3480 | 1394773 | 487 | 51.63 | 400.8 | 344 | 3374 | 7 |
| Rhagopthalmus_sp_Rhagophthalmidae | 3759 | 1657354 | 536 | 56.73 | 440.9 | 359 | 8290 | 8 |
| S39_Limnichidae_indet_Limnichidae | 3700 | 1519176 | 507 | 52.59 | 410.6 | 343 | 5081 | 6 |
| Sandalus_niger_Rhipiceridae | 3582 | 1444217 | 488 | 52.01 | 403.2 | 339 | 7792 | 6 |
| Scarelus_anthracinus_Lycidae | 3727 | 1565674 | 507 | 53.94 | 420.1 | 344 | 7145 | 33 |
| G19009_Semiotus_sp_Elateridae | 3978 | 1620841 | 497 | 51.55 | 407.5 | 328 | 5940 | 8 |
| R19009_Silis_sp_Cantharidae | 3895 | 1725059 | 537 | 56.78 | 442.9 | 367 | 5718 | 8 |
| Sinopyrophorus_schimmeli_Sinopyrophoridae | 3817 | 1309593 | 426 | 43.55 | 343.1 | 278 | 2675 | 10 |
| G20013_Sinopyrophorus_sp_Sinopyrophoridae | 3812 | 1234280 | 403 | 40.71 | 323.8 | 260 | 3582 | 7 |
| Taphes_brevicollis_Lycidae | 3697 | 1563002 | 518 | 54.32 | 422.8 | 351 | 5725 | 26 |
| G19001_Sinelater_perroti_Elateridae | 3860 | 1365121 | 430 | 45.02 | 353.7 | 290 | 4703 | 9 |
| Tribolium_castaneum | 4197 | 2467205 | 747 | 73.52 | 587.8 | 453 | 8743 | 64 |
| Trixagus_carinifrons_Throscidae | 3505 | 1129285 | 386 | 41.93 | 322.2 | 298 | 2559 | 10 |

| Taxon _name_Family | AA after AliScore filtering | | | | | | | |
|---|---|---|---|---|---|---|---|---|
| | no_genes | total_no_aa | N50 | percent_determined | average_length | median_length | max_length | min_length |
| A4488_Throscidae_indet_Throscidae | 3741 | 1002776 | 320 | 60.19 | 268.1 | 223 | 4712 | 13 |
| Abscondita_terminalis_Lampyridae | 3723 | 1436880 | 463 | 87.22 | 385.9 | 319 | 5382 | 23 |
| Agrilus_planipennis_Buprestidae | 4102 | 1663441 | 485 | 90.42 | 405.5 | 329 | 7111 | 46 |
| Agrilus_zanthoxylumi_Buprestidae | 3910 | 1537960 | 476 | 88.55 | 393.3 | 324 | 5570 | 13 |
| Agrypnus_murinus_Elateridae | 3604 | 1232906 | 420 | 77.77 | 342.1 | 292.5 | 4032 | 21 |
| Anoplophora_glabripennis | 4116 | 1625471 | 470 | 88.6 | 394.9 | 326 | 4729 | 56 |
| Anorus_parvicollis_Dascillidae | 3657 | 1263518 | 422 | 78.47 | 345.5 | 291 | 7117 | 22 |
| Antennolycus_constrictus_Lycidae | 3710 | 1380613 | 450 | 84.01 | 372.1 | 310 | 4942 | 19 |
| Aquatica_ficta_Lampyridae | 3655 | 1265981 | 429 | 78.61 | 346.4 | 292 | 4542 | 16 |
| Asymmetricata_circumdata_Lampyridae | 3551 | 1293331 | 443 | 82.67 | 364.2 | 306 | 5382 | 12 |
| Aspisoma_lineatum_Lampyridae | 3889 | 1512249 | 470 | 87.83 | 388.9 | 320 | 6533 | 24 |
| Bicellonycha_wickershamorum_Lampyridae | 3528 | 1195356 | 421 | 77.27 | 338.8 | 287 | 4934 | 21 |
| Brachypsectra_fulva_Brachypsectridae | 3493 | 1100582 | 385 | 72.34 | 315.1 | 282 | 3173 | 23 |
| Byrrhus_pilula_Byrrhidae | 3237 | 1014810 | 386 | 72.97 | 313.5 | 272 | 3517 | 22 |
| R19006_Callirhipidae_indet_Callirhipidae | 3645 | 1321513 | 444 | 82.6 | 362.6 | 301 | 6794 | 17 |
| Calopteron_sp_Lycidae | 3642 | 1401415 | 462 | 86.42 | 384.8 | 320 | 5528 | 12 |
| Capnodis_tenebrionis_Buprestidae | 3703 | 1383319 | 450 | 83.83 | 373.6 | 312 | 5570 | 19 |
| R19003_Dicronychus_rubripes_Elateridae | 3874 | 1510170 | 470 | 88.43 | 389.8 | 324 | 7117 | 28 |
| Cautires_communis_Lycidae | 3638 | 927188 | 301 | 56.78 | 254.9 | 217 | 3214 | 25 |
| G19010_Cebrio_sp_Elateridae | 3386 | 757998 | 268 | 48.9 | 223.9 | 192 | 2672 | 13 |
| Cerophytum_elateroides_Cerophytiae | 2977 | 516314 | 203 | 39.63 | 173.4 | 144 | 1359 | 17 |
| Chauliognathus_opacus_Cantharidae | 3489 | 836092 | 292 | 54.18 | 239.6 | 197 | 5344 | 16 |
| Conderis_signicollis_Lycidae | 3631 | 1269492 | 431 | 78.74 | 349.6 | 293 | 6783 | 19 |
| Elaterinae_indet_Elateridae | 3889 | 1566031 | 483 | 90.61 | 402.7 | 333 | 5939 | 30 |
| Dendroctonus_ponderosae | 3952 | 1612240 | 485 | 90.16 | 408 | 333 | 7099 | 44 |
| R19012_Denticollis_sp_Elateridae | 3885 | 1489155 | 464 | 87.18 | 383.3 | 323 | 4249 | 16 |
| Dexoris_ruzzieri_Lycidae | 3710 | 886401 | 282 | 53.3 | 238.9 | 203 | 2893 | 20 |
| Dictyoptera_sp_Lycidae | 3183 | 771805 | 307 | 56.64 | 242.5 | 201 | 3273 | 20 |
| Dihammatus_sp_Lycidae | 3685 | 1356364 | 447 | 83 | 368.1 | 308 | 6051 | 26 |
| Dilophotes_sp_Lycidae | 3680 | 1375319 | 452 | 84.07 | 373.7 | 314 | 3526 | 28 |
| Distremocephalus_sp_Phengodidae | 3571 | 1208506 | 411 | 77.44 | 338.4 | 294 | 2426 | 30 |
| Drilonius_sp_Omethidae | 3837 | 1478801 | 464 | 86.73 | 385.4 | 321 | 6029 | 23 |
| G18004_Drilus_mauritanicus_Elateridae | 3937 | 1349429 | 416 | 77.03 | 342.8 | 285 | 6426 | 24 |
| Drilus_concolor_Elateridae | 2876 | 740980 | 318 | 60.43 | 257.6 | 216.5 | 2561 | 20 |
| Dryops_sp_Dryopidae | 3655 | 1321529 | 440 | 82 | 361.6 | 309 | 3046 | 22 |
| R19010_Quasimus_sp_Elateridae | 3886 | 1496961 | 463 | 87.55 | 385.2 | 321 | 4986 | 20 |
| R19011_Agrypninae_indet_Elateridae | 3814 | 1461487 | 462 | 86.71 | 383.2 | 322 | 6822 | 24 |
| G19012_Penia_sp_Elateridae | 3875 | 1234350 | 386 | 71.74 | 318.5 | 273 | 4565 | 20 |
| G19013_Dromaeolinae indet_Eucnemidae | 3802 | 1259515 | 407 | 73.81 | 331.3 | 277 | 4693 | 19 |
| Ellychnia_sp_Lampyridae | 3649 | 1325516 | 448 | 81.74 | 363.3 | 303 | 4197 | 16 |
| Eucnemis_capucina_Eucnemidae | 3850 | 1431888 | 453 | 83.97 | 371.9 | 307 | 4700 | 28 |
| G19007_Eudicronychus_sp_Elateridae | 3316 | 744169 | 271 | 49.42 | 224.4 | 190 | 2105 | 16 |
| Eurypogon_sp_Artematopodidae | 3530 | 1066182 | 362 | 69.27 | 302 | 279.5 | 1308 | 24 |
| G20001_Octinodes_sp_Elateridae | 3878 | 1228125 | 388 | 70.74 | 316.7 | 258 | 3359 | 16 |
| G20002_Pyrearinus_sp_Elateridae | 3951 | 1414813 | 435 | 80.42 | 358.1 | 299 | 4531 | 27 |
| G20003_Cussolenis_sp_Elateridae | 3788 | 1180762 | 390 | 69.3 | 311.7 | 253 | 7119 | 27 |
| G20004_Eudicronychus_rufus_Elateridae | 3248 | 694488 | 264 | 46.93 | 213.8 | 178 | 1347 | 20 |
| G20005_Mulsanteus_sp_Elateridae | 3884 | 1152882 | 354 | 66.89 | 296.8 | 250 | 3315 | 14 |
| G20006_Elaterinae_indet_Elateridae | 3917 | 1290198 | 402 | 74.08 | 329.4 | 269 | 3426 | 22 |
| G20007_Hapatesus_tropicus_Elateridae | 3752 | 984707 | 311 | 58.45 | 262.4 | 225 | 2623 | 21 |
| G20008_Eucnemidae_indet_Eucnemidae | 3444 | 891093 | 324 | 56.99 | 258.7 | 204 | 6787 | 17 |
| G20009_Drapetes_sp_Elateridae | 3745 | 1067186 | 350 | 63.36 | 285 | 232 | 3049 | 19 |
| G20010_Chelonarium_sp_Chelonaridae | 3710 | 863278 | 287 | 51.99 | 232.7 | 186.5 | 2375 | 20 |
| G20011_Balgus_obconicus_Elateridae | 3608 | 976576 | 332 | 59.91 | 270.7 | 227.5 | 2894 | 13 |
| G20012_Tibionema_abdominalis_Elateridae | 2904 | 558379 | 231 | 42.59 | 192.3 | 166 | 1874 | 15 |
| G20019_Eulichas_dudgeoni_Eulichadidae | 3926 | 1237007 | 380 | 70.96 | 315.1 | 258 | 5477 | 23 |
| G20021_Elater_ferrugineus_Elateridae | 3834 | 1290044 | 415 | 75.61 | 336.5 | 281 | 4371 | 22 |
| G20022_Pseudotetralobus_capucinus_Elateridae | 3837 | 1152828 | 369 | 67.55 | 300.5 | 248 | 3420 | 20 |
| G20023_Rostricephalus_vitalis_Elateridae | 3590 | 890614 | 298 | 55.61 | 248.1 | 213 | 3435 | 21 |
| G20024_Aphileus_lucanoides_Elateridae | 3814 | 1116646 | 354 | 65.28 | 292.8 | 246 | 3408 | 14 |
| G20025_Cheguevaria_sp_Lampyridae | 3734 | 946074 | 297 | 56.51 | 253.4 | 217 | 1841 | 20 |
| G20028_Stenocladius_sp_Lampyridae | 3732 | 1077278 | 350 | 63.96 | 288.7 | 244 | 3224 | 16 |
| G20030_Rhagophthalmidae_indet_Rhagophthalmidae | 3498 | 862378 | 299 | 54.21 | 246.5 | 211.5 | 2006 | 22 |
| G20031_Rhagophthalmidae_indet_Rhagophthalmidae | 3708 | 990075 | 321 | 60.27 | 267 | 228 | 2027 | 23 |
| G20033_Euanoma_sp_Elateridae | 3867 | 1306300 | 406 | 75.6 | 337.8 | 280 | 7119 | 21 |
| G20035_Artematopus_sp_Artematopodidae | 3891 | 1320382 | 416 | 76.23 | 339.3 | 276 | 4735 | 18 |
| G21002_Sphaenelater_lineicollis_Elateridae | 3755 | 1055442 | 336 | 62.57 | 281.1 | 244 | 2004 | 20 |
| G21003_Analastesa_arabica_Elateridae | 2718 | 490501 | 214 | 38.38 | 180.5 | 153.5 | 1688 | 18 |
| G21004_Paulusiella_serraticornis_Elateridae | 3922 | 1289211 | 398 | 73.78 | 328.7 | 270 | 5377 | 20 |
| G21006_Rhipiceridae_indet_Rhipiceridae | 3753 | 1401611 | 448 | 82.6 | 373.5 | 308 | 7100 | 22 |
| G21007_Cneoglossa_sp_Cneoglossidae | 3889 | 1454289 | 460 | 84.19 | 373.9 | 308 | 4032 | 17 |
| G21009_Psilocladus_sp_Lampyridae | 3785 | 1087638 | 347 | 64.35 | 287.4 | 244 | 2043 | 16 |
| G21010_Amydetinae_indet_Lampyridae | 3924 | 1219228 | 376 | 69.84 | 310.7 | 271 | 3427 | 20 |
| G21011_Ototretinae_indet_Lampyridae | 3904 | 1199288 | 363 | 68.88 | 307.2 | 264 | 3262 | 28 |
| G21012_Austroselasia_sp_Elateridae | 3889 | 1140363 | 356 | 66.5 | 293.2 | 245 | 3420 | 26 |
| G21013_Habashaselasia_nekemtensis_Elateridae | 3875 | 1110849 | 343 | 64.32 | 286.7 | 242 | 2893 | 28 |
| G21014_Elatichrosis_sp_Elateridae | 3828 | 1092778 | 338 | 63.79 | 285.5 | 236 | 3740 | 22 |
| G21015_Brachypterodrilus_sp_Lampyridae | 3747 | 1013088 | 324 | 59.96 | 270.4 | 230 | 3005 | 9 |
| G21016_Rhagophthalmidae_indet_Rhagophthalmidae | 3889 | 1221070 | 376 | 70.66 | 314 | 265 | 3426 | 26 |
| G21017_Lissominae_indet_Elateridae | 3832 | 1476760 | 464 | 85.83 | 385.4 | 317 | 5891 | 31 |
| G21019_Eucnemidae_indet_Eucnemidae | 2938 | 633175 | 268 | 44.54 | 215.5 | 178 | 2715 | 16 |
| G21023_Ceratogonys_spinicorne_Eucnemidae | 3741 | 1138209 | 371 | 67.37 | 304.3 | 255 | 2595 | 16 |
| G21026_Pyrophorini_indet_Elateridae | 3786 | 1112993 | 354 | 65.4 | 294 | 250 | 3425 | 11 |
| G21027_Elaterinae_indet_Elateridae | 2518 | 450730 | 211 | 37.39 | 179 | 150 | 1656 | 17 |
| G21028_Dascilidae_indet_Dascillidae | 3884 | 1494666 | 462 | 86.42 | 384.8 | 317 | 7117 | 24 |
| G21029_Drilaster_sp_Lampyridae | 3392 | 749251 | 266 | 47.93 | 220.9 | 187 | 1906 | 15 |
| G21030_Psephenidae_indet_Psephenidae | 3879 | 1371509 | 427 | 79.02 | 353.6 | 297 | 6786 | 27 |
| G21031_Rousia_sp_Elateridae | 3926 | 1486680 | 452 | 85.07 | 378.7 | 316 | 6426 | 25 |
| G21035_Anaissus_sp_Elateridae | 3890 | 1264568 | 398 | 73.25 | 325.1 | 275 | 2958 | 24 |
| G21036_Campyloxenus_pyrothorax_Elateridae | 3779 | 1103791 | 350 | 66.09 | 292.1 | 246 | 7119 | 11 |
| G21037_Pityobius_anguinus_Elateridae | 3419 | 874914 | 310 | 56.43 | 255.9 | 216 | 1589 | 18 |
| G21038_Adelocera_sp_Elateridae | 3735 | 1044892 | 334 | 62.03 | 279.8 | 233 | 2416 | 19 |
| G21039_Psephenidae_indet_Psephenidae | 3917 | 1459505 | 452 | 83.88 | 372.6 | 309 | 4713 | 20 |
| G21040_Adelocera_sp_Elateridae | 3758 | 961955 | 305 | 57.12 | 256 | 218 | 2333 | 20 |
| G21041_Adelocera_sp_Elateridae | 3969 | 1492449 | 450 | 84.71 | 376 | 312 | 5097 | 9 |
| G21042_Telegeusis_sp_Omethidae | 3756 | 1126342 | 367 | 66.93 | 299.9 | 247 | 3737 | 17 |
| G22002_Plectonotum_laterale_Cantharidae | 3772 | 1095035 | 346 | 65.02 | 290.3 | 237 | 3424 | 23 |
| G22003_Cerophytum_japonicum_Cerophytiae | 3157 | 597183 | 222 | 43.4 | 189.2 | 159 | 1480 | 15 |
| G22005_Nyctophyxis_ocellatus_Elateridae | 2536 | 827998 | 413 | 69.03 | 326.5 | 278 | 3420 | 13 |
| G22007_Anaspasis_paralella_Elateridae | 2667 | 479186 | 210 | 38.52 | 179.7 | 155 | 1714 | 21 |
| G22009_Jurasai_digitusdei_Jurasaidae | 3593 | 1040676 | 343 | 64.24 | 289.6 | 243 | 3701 | 15 |
| G22015_Protelmidae_indet_Protelmidae | 3816 | 1195462 | 387 | 70.05 | 313.3 | 257 | 2514 | 16 |
| G22020_Lutrochidae_indet_Lutrochidae | 3883 | 1349773 | 429 | 78.15 | 347.6 | 285 | 5323 | 15 |
| G22022_Lutrochidae_indet_Lutrochidae | 3923 | 1485519 | 459 | 84.97 | 378.7 | 314 | 6424 | 23 |
| G22024_Elmidae_indet_Elmidae | 3376 | 1226560 | 443 | 79.75 | 363.3 | 298 | 6787 | 27 |
| G22025_Ototretinae_indet_Lampyridae | 3978 | 1323403 | 393 | 75.06 | 332.7 | 288 | 6419 | 16 |
| G22026_Stenocladius_sp_Lampyridae | 3492 | 821722 | 285 | 52.13 | 235.3 | 200 | 2144 | 19 |
| G22027_Lampyrinae_indet_Lampyridae | 3829 | 1319366 | 411 | 76.49 | 344.6 | 281 | 5381 | 20 |
| G19002_Hemiops_sp_Elateridae | 3615 | 845844 | 283 | 52.19 | 234 | 196 | 2357 | 24 |
| Heterocerus_fenestratus_Heteroceridae | 3082 | 661962 | 252 | 52.3 | 214.8 | 198 | 777 | 12 |
| Heteromastix_sp_Cantharidae | 3876 | 1480162 | 463 | 86.7 | 381.9 | 318 | 2839 | 16 |
| Hypnoidus_riparius_Elateridae | 3849 | 1502133 | 470 | 88.31 | 390.3 | 327 | 4026 | 30 |
| G19015_Iberobaenia_minuta_Iberobaeniidae | 3601 | 861557 | 284 | 53.16 | 239.3 | 202 | 3074 | 21 |
| Ignelater_luminosus_Elateridae | 3834 | 1492845 | 472 | 87.23 | 389.4 | 322 | 5377 | 20 |
| R19001_Isorhipis_marmottani_Eucnemidae | 3876 | 1478933 | 462 | 86.7 | 381.6 | 320 | 5474 | 30 |
| Konoplatycis_otome_Lycidae | 3381 | 955964 | 344 | 65.7 | 282.7 | 244 | 6317 | 22 |
| Lamprigera_yunnana_Lampyridae | 3725 | 1472146 | 479 | 88.21 | 395.2 | 324 | 5382 | 16 |
| Lamprohiza_splendidula_Lampyridae | 3623 | 1207533 | 401 | 75.53 | 333.3 | 285 | 3444 | 26 |
| S1_Lampyris_sardiniae_Lampyridae | 3744 | 1448966 | 470 | 87.36 | 387 | 322 | 6598 | 11 |
| Leptinotarsa_decemlineata | 4007 | 1405872 | 417 | 77.88 | 350.9 | 294 | 4506 | 33 |
| Leptolycus_sp_Lycidae | 3678 | 839150 | 275 | 50.45 | 228.2 | 190 | 3262 | 22 |
| Libnetis_sp_Lycidae | 3686 | 1355115 | 445 | 83.25 | 367.6 | 311 | 3248 | 28 |
| Limonius_californicus_Elateridae | 3884 | 1156365 | 344 | 67.04 | 297.7 | 250 | 6426 | 19 |
| R19013_Lissomus_sp_Elateridae | 3815 | 1470736 | 464 | 87.64 | 385.5 | 325 | 3498 | 37 |
| Lucidota_atra_Lampyridae | 3730 | 1391560 | 453 | 84.05 | 373.1 | 313 | 5381 | 25 |
| Lycoprogenthes_sp_Lycidae | 3664 | 1387503 | 459 | 85.7 | 378.7 | 316 | 3645 | 31 |
| Lycostomus_kraatzi_Lycidae | 3430 | 1246325 | 440 | 83.06 | 363.4 | 306 | 3820 | 34 |
| Lygistopterus_sanguineus_Lycidae | 3662 | 1374969 | 450 | 84.68 | 375.5 | 314 | 5359 | 28 |
| Lyropaeus_optabilis_Lycidae | 3472 | 1239256 | 435 | 80.22 | 356.9 | 300.5 | 5593 | 19 |
| Ponyalis_sp_Lycidae | 3625 | 1357731 | 454 | 84.67 | 374.5 | 315 | 4395 | 15 |
| R20009_Malthodes_sp_Cantharidae | 3812 | 1396125 | 439 | 83.2 | 366.2 | 316 | 2581 | 27 |
| Melanotus_cribricollis_Elateridae | 3663 | 1444342 | 477 | 88.7 | 394.3 | 325 | 5383 | 32 |
| Melanotus_villosus_Elateridae | 3666 | 1376048 | 452 | 85.23 | 375.4 | 320.5 | 4175 | 30 |
| Micronaspis_floridana_Lampyridae | 3471 | 1115479 | 396 | 73.52 | 321.4 | 274 | 2558 | 16 |
| Microphotus_sp_Lampyridae | 3534 | 1252292 | 432 | 79.89 | 354.4 | 297 | 4432 | 26 |
| G19011_Diplophoenicus_sp_Elateridae | 3921 | 1423405 | 438 | 81.9 | 363 | 308 | 5382 | 23 |
| Negastrius_arenicola_Elateridae | 3938 | 1528793 | 472 | 88.52 | 388.2 | 324 | 4292 | 16 |
| Notolioon_sp_Byrrhidae | 3157 | 1033372 | 406 | 76.44 | 327.3 | 281 | 2223 | 24 |
| Omalisus_fontisbellaquei_Elateridae | 3858 | 1240747 | 383 | 72.1 | 321.6 | 275 | 7119 | 20 |
| Onthophagus_taurus | 4079 | 1656119 | 481 | 90.68 | 406 | 334 | 7002 | 43 |
| R19005_Ototretinae_indet_Lampyridae | 3823 | 1483093 | 472 | 87.87 | 387.9 | 321 | 6790 | 32 |
| R20013_Ototretinae_indet_Lampyridae | 3875 | 1471731 | 460 | 86.13 | 379.8 | 320 | 2887 | 11 |
| G19008_Oxynopterus_sp_Elateridae | 3919 | 1427311 | 444 | 82.07 | 364.2 | 305 | 4451 | 27 |
| G19005_Pachyderes_sp_Elateridae | 3901 | 1237875 | 381 | 71.34 | 317.3 | 273 | 4519 | 22 |
| G19006_Parasaphes_sp_Elateridae | 3907 | 1392013 | 430 | 80.12 | 356.3 | 299 | 5125 | 22 |
| G19004_Pectocera_sp_Elateridae | 3915 | 1227936 | 377 | 71.08 | 313.6 | 268 | 3415 | 18 |
| Phausis_reticulata_Lampyridae | 3688 | 1361533 | 447 | 83.69 | 369.2 | 313 | 4542 | 11 |
| Phengodes_sp_B_Phengodidae | 3635 | 1339712 | 447 | 83.29 | 368.6 | 308 | 6325 | 28 |
| Phengodes_sp_A_Phengodidae | 3858 | 1189928 | 372 | 68.92 | 308.4 | 260 | 3707 | 28 |
| Photinus_australis_Lampyridae | 3533 | 1217065 | 425 | 78.72 | 344.5 | 297 | 3469 | 16 |
| Photinus_carolinus_Lampyridae | 3566 | 1236581 | 428 | 78.61 | 346.8 | 294 | 4431 | 15 |
| Photinus_macdermotti_Lampyridae | 3396 | 1222625 | 448 | 81.52 | 360 | 298 | 5276 | 14 |
| Photinus_pyralis_Lampyridae | 3874 | 1516269 | 471 | 88.47 | 391.4 | 323 | 7117 | 18 |
| Photinus_scintillans_Lampyridae | 3775 | 1440109 | 462 | 86.53 | 381.5 | 317 | 4432 | 26 |
| Photuris_frontalis_Lampyridae | 3698 | 1394041 | 461 | 85.49 | 377 | 315 | 6425 | 22 |
| Photuris_sp_Lampyridae | 3531 | 1285401 | 451 | 82.08 | 364 | 306 | 5382 | 17 |
| Phrixothrix_hirtus_Phengodidae | 2654 | 676935 | 325 | 62.96 | 255.1 | 206 | 4205 | 17 |
| A01544_Plastocerus_angulosus_Elateridae | 3918 | 1361116 | 423 | 78.34 | 347.4 | 289 | 7104 | 11 |
| Platerodrilus_sp_Lycidae | 3218 | 1112182 | 421 | 78.68 | 345.6 | 291 | 5470 | 22 |
| Plateros_sp_Lycidae | 3633 | 1394296 | 464 | 86.31 | 383.8 | 320 | 3392 | 20 |
| Ponyalis_quadricollis_Lycidae | 3527 | 1275376 | 438 | 82.25 | 361.6 | 307 | 3288 | 29 |
| R19004_Ptilodactylidae_indet_Ptilodactylidae | 3951 | 1545324 | 477 | 89.03 | 391.1 | 329 | 6093 | 27 |
| Ptosima_undecimmaculata_Buprestidae | 3331 | 1100732 | 404 | 76.23 | 330.5 | 281 | 5560 | 21 |
| Pyractomena_borealis_Lampyridae | 3727 | 1408040 | 462 | 85.25 | 377.8 | 316 | 6424 | 23 |
| Pyrearinus_fragilis_Elateridae | 3857 | 1541735 | 481 | 90.34 | 399.7 | 332 | 6541 | 31 |
| Pyrocoelia_pectoralis_Lampyridae | 3735 | 1348123 | 445 | 81.79 | 360.9 | 301 | 6531 | 21 |
| R18040_Mangkutanus_sp_Lycidae | 3726 | 1394284 | 452 | 84.94 | 374.2 | 311 | 6787 | 16 |
| R20002_Agrypninae_indet_Elateridae | 3741 | 1377418 | 446 | 83.53 | 368.2 | 316 | 2269 | 20 |
| R20003_Athous_sp_Elateridae | 3910 | 1501810 | 462 | 86.9 | 384.1 | 320 | 3572 | 29 |
| R20004_Selatosomus_aeneus_Elateridae | 3611 | 1256143 | 424 | 80.84 | 347.9 | 306 | 2102 | 24 |
| R20005_Ampedus_sineatus_Elateridae | 3841 | 1498722 | 472 | 88.47 | 390.2 | 328 | 3719 | 19 |
| R20006_Ampedus_cinnabarinu_Elateridaes | 3873 | 1510326 | 471 | 88.28 | 390 | 327 | 2844 | 26 |
| R20007_Anostirus_purpureus_Elateridae | 3842 | 1482063 | 467 | 87.4 | 385.8 | 323 | 6582 | 27 |
| R20008_Agriotes_sp_Elateridae | 3877 | 1508716 | 472 | 87.96 | 389.1 | 328 | 5382 | 31 |
| R20010_Elaterinae_indet_Elateridae | 3861 | 1463145 | 460 | 85.42 | 379 | 319 | 3193 | 20 |
| R20011_Alaus_myops_Elateridae | 3220 | 969328 | 363 | 71.31 | 301 | 259 | 5060 | 22 |
| R20014_Limnius_perrisi_Elmidae | 3914 | 1518600 | 467 | 88.24 | 388 | 324.5 | 2400 | 17 |
| R20016_Cardiophorus_discicollis_Elateridae | 3906 | 1498878 | 463 | 86.76 | 383.7 | 319 | 3025 | 20 |
| R20017_Eubria_palustris_Psephenidae | 3911 | 1521238 | 474 | 88.23 | 389 | 322 | 3524 | 32 |
| R20018_Phosphaenus_hemipterus_Lampyridae | 3861 | 1482758 | 464 | 87.06 | 384 | 319 | 3505 | 30 |
| R20019_Dascillus_cervinus_Dascillidae | 3930 | 1498445 | 462 | 86.6 | 381.3 | 319.5 | 2688 | 24 |
| R20022_Lampyris_noctiluca_Lampyridae | 3891 | 1488813 | 462 | 86.61 | 382.6 | 323 | 2973 | 16 |
| Anischia_sp_Eucnemidae | 3858 | 1351586 | 427 | 78.39 | 350.3 | 287 | 5386 | 24 |
| Platycrepidius_sp_Elateridae | 3961 | 1478041 | 454 | 83.96 | 373.1 | 310 | 7119 | 9 |
| Lamellipalpus_pacholatkoi_Lampyridae | 3973 | 1320858 | 391 | 74.97 | 332.5 | 286 | 3418 | 27 |
| Byrrhinus_sp_Limnichidae | 3913 | 1528680 | 474 | 88.22 | 390.7 | 322 | 5281 | 34 |
| Epilichas_sp_Ptilodactylidae | 3414 | 752353 | 265 | 48.97 | 220.4 | 178 | 2094 | 16 |
| Rhagonycha_fulva_Cantharidae | 3455 | 1230562 | 431 | 80.67 | 356.2 | 310 | 3311 | 32 |
| Rhagopthalmus_sp_Rhagophthalmidae | 3740 | 1450880 | 467 | 88.01 | 387.9 | 322 | 7118 | 31 |
| S39_Limnichidae_indet_Limnichidae | 3666 | 1333168 | 449 | 82.5 | 363.7 | 306 | 4994 | 26 |
| Sandalus_niger_Rhipiceridae | 3549 | 1276236 | 435 | 81.24 | 359.6 | 304 | 6793 | 16 |
| Scarelus_anthracinus_Lycidae | 3711 | 1394069 | 455 | 84.9 | 375.7 | 314 | 6441 | 20 |
| G19009_Semiotus_sp_Elateridae | 3956 | 1407994 | 431 | 80.06 | 355.9 | 293 | 5382 | 32 |
| R19009_Silis_sp_Cantharidae | 3885 | 1516875 | 469 | 88.47 | 390.4 | 324 | 5382 | 22 |
| Sinopyrophorus_schimmeli_Sinopyrophoridae | 3788 | 1143793 | 370 | 68.51 | 302 | 249.5 | 2232 | 18 |
| G20013_Sinopyrophorus_sp_Sinopyrophoridae | 3765 | 1082323 | 353 | 64.74 | 287.5 | 235 | 3190 | 14 |
| Taphes_brevicollis_Lycidae | 3684 | 1394266 | 462 | 85.24 | 378.5 | 317 | 5619 | 19 |
| G19001_Sinelater_perroti_Elateridae | 3828 | 1194734 | 377 | 70.51 | 312.1 | 257 | 4509 | 15 |
| Tribolium_castaneum | 4163 | 1796992 | 514 | 97.02 | 431.7 | 352 | 7119 | 52 |
| Trixagus_carinifrons_Throscidae | 3409 | 992887 | 351 | 68.17 | 291.3 | 270 | 1099 | 12 |

**Supplementary Table S4.** Summary statistics by taxon across data processing stages (raw data, outlier removal, and AliScore filtering) at the nucleotide level.

**NT raw data**

| Taxon _name_Family | no_genes | total_no_nt | N50 | percent_determined | average_length | median_length | max_length | min_length |
|---|---|---|---|---|---|---|---|---|
| A4488_Throscidae_indet_Throscidae | 3839 | 3465643 | 1074 | 37.16 | 902.7 | 750 | 16845 | 18 |
| Abscondita_terminalis_Lampyridae | 3756 | 4930776 | 1584 | 54.75 | 1312.8 | 1068 | 16512 | 87 |
| Agrilus_planipennis_Buprestidae | 4224 | 7015396 | 2118 | 67.52 | 1660.8 | 1266 | 27027 | 162 |
| Agrilus_zanthoxylumi_Buprestidae | 3981 | 5826822 | 1833 | 60.84 | 1463.7 | 1164 | 17103 | 45 |
| Agrypnus_murinus_Elateridae | 3666 | 4246613 | 1416 | 48.26 | 1158.4 | 981 | 12312 | 72 |
| Anoplophora_glabripennis | 4224 | 6798372 | 1995 | 65.43 | 1609.5 | 1224 | 58590 | 177 |
| Anorus_parvicollis_Dascillidae | 3719 | 4365434 | 1431 | 49.23 | 1173.8 | 978 | 23625 | 21 |
| Antennolycus_constrictus_Lycidae | 3739 | 4669290 | 1533 | 52.2 | 1248.8 | 1023 | 15084 | 93 |
| Aquatica_ficta_Lampyridae | 3703 | 4293852 | 1431 | 48.38 | 1159.6 | 972 | 14247 | 72 |
| Asymmetricata_circumdata_Lampyridae | 3603 | 4384390 | 1479 | 51.28 | 1216.9 | 1017 | 17178 | 36 |
| Aspisoma_lineatum_Lampyridae | 3908 | 5145713 | 1596 | 54.73 | 1316.7 | 1074 | 21108 | 48 |
| Bicellonycha_wickershamorum_Lampyridae | 3622 | 4093002 | 1395 | 46.98 | 1130 | 951 | 16086 | 18 |
| Brachypsectra_fulva_Brachypsectridae | 3586 | 3785123 | 1299 | 44.5 | 1055.5 | 945 | 9924 | 21 |
| Byrrhus_pilula_Byrrhidae | 3349 | 3544270 | 1296 | 44.61 | 1058.3 | 918 | 10842 | 90 |
| R19006_Callirhipidae_indet_Callirhipidae | 3700 | 4559397 | 1521 | 51.59 | 1232.3 | 1000.5 | 22989 | 36 |
| Calopteron_sp_Lycidae | 3668 | 4724015 | 1563 | 53.8 | 1287.9 | 1060.5 | 16809 | 24 |
| Capnodis_tenebrionis_Buprestidae | 3777 | 4974042 | 1605 | 54.16 | 1316.9 | 1086 | 17751 | 48 |
| R19003_Dicronychus_rubripes_Elateridae | 3885 | 5175630 | 1605 | 55.88 | 1332.2 | 1092 | 24966 | 111 |
| Cautires_communis_Lycidae | 3780 | 3209130 | 993 | 34.4 | 849 | 720 | 9762 | 33 |
| G19010_Cebrio_sp_Elateridae | 3535 | 2686604 | 912 | 29.91 | 760 | 651 | 10119 | 18 |
| Cerophytum_elateroides_Cerophytiae | 3250 | 1831373 | 663 | 22.59 | 563.5 | 471 | 4077 | 21 |
| Chauliognathus_opacus_Cantharidae | 3669 | 2846709 | 933 | 32.24 | 775.9 | 636 | 16404 | 24 |
| Conderis_signicollis_Lycidae | 3678 | 4266720 | 1431 | 48.07 | 1160.1 | 955.5 | 22641 | 105 |
| Elaterinae_indet_Elateridae | 3907 | 5411260 | 1662 | 57.53 | 1385 | 1134 | 18408 | 27 |
| Dendroctonus_ponderosae | 4224 | 6728070 | 1986 | 64.75 | 1592.8 | 1245 | 22002 | 150 |
| R19012_Denticollis_sp_Elateridae | 3908 | 5120445 | 1578 | 54.94 | 1310.2 | 1092 | 12969 | 48 |
| Dexoris_ruzzieri_Lycidae | 3897 | 3080319 | 927 | 31.98 | 790.4 | 672 | 10485 | 36 |
| Dictyoptera_sp_Lycidae | 3401 | 2620240 | 984 | 32.45 | 770.4 | 630 | 10089 | 21 |
| Dihammatus_sp_Lycidae | 3731 | 4600263 | 1515 | 50.91 | 1233 | 1026 | 21366 | 30 |
| Dilophotes_sp_Lycidae | 3715 | 4626266 | 1512 | 51.84 | 1245.3 | 1038 | 10923 | 21 |
| Distremocephalus_sp_Phengodidae | 3648 | 4122740 | 1380 | 47.92 | 1130.1 | 984 | 8559 | 33 |
| Drilonius_sp_Omethidae | 3870 | 5074603 | 1587 | 54.48 | 1311.3 | 1083 | 18432 | 60 |
| G18004_Drilus_mauritanicus_Elateridae | 3978 | 4607205 | 1407 | 47.73 | 1158.2 | 951 | 19671 | 21 |
| Drilus_concolor_Elateridae | 3030 | 2596157 | 1068 | 36.11 | 856.8 | 723 | 10050 | 57 |
| Dryops_sp_Dryopidae | 3696 | 4526334 | 1500 | 51.44 | 1224.7 | 1029 | 9336 | 51 |
| R19010_Quasimus_sp_Elateridae | 3909 | 5122182 | 1584 | 54.73 | 1310.4 | 1077 | 15192 | 18 |
| R19011_Agrypninae_indet_Elateridae | 3843 | 5035485 | 1596 | 54.63 | 1310.3 | 1092 | 22269 | 24 |
| G19012_Penia_sp_Elateridae | 3935 | 4282346 | 1305 | 44.6 | 1088.3 | 927 | 14322 | 18 |
| G19013_Dromaeolinae indet_Eucnemidae | 3866 | 4347260 | 1377 | 45.81 | 1124.5 | 921 | 14688 | 39 |
| Ellychnia_sp_Lampyridae | 3734 | 4489740 | 1479 | 49.38 | 1202.4 | 996 | 13971 | 24 |
| Eucnemis_capucina_Eucnemidae | 3928 | 4913616 | 1524 | 51.66 | 1250.9 | 1021.5 | 14343 | 48 |
| G19007_Eudicronychus_sp_Elateridae | 3511 | 2649521 | 912 | 29.51 | 754.6 | 633 | 6657 | 30 |
| Eurypogon_sp_Artematopodidae | 3653 | 3691979 | 1212 | 42.46 | 1010.7 | 939 | 4332 | 36 |
| G20001_Octinodes_sp_Elateridae | 3932 | 4253079 | 1314 | 44.3 | 1081.7 | 882 | 10905 | 93 |
| G20002_Pyrearinus_sp_Elateridae | 3978 | 4895399 | 1491 | 50.53 | 1230.6 | 1017 | 14178 | 63 |
| G20003_Cussolenis_sp_Elateridae | 3853 | 4093311 | 1320 | 43.08 | 1062.4 | 849 | 25869 | 36 |
| G20004_Eudicronychus_rufus_Elateridae | 3458 | 2484642 | 873 | 27.91 | 718.5 | 607.5 | 4410 | 21 |
| G20005_Mulsanteus_sp_Elateridae | 3948 | 3983777 | 1197 | 41.58 | 1009.1 | 840 | 10083 | 21 |
| G20006_Elaterinae_indet_Elateridae | 3964 | 4497034 | 1377 | 46.59 | 1134.5 | 919.5 | 10437 | 27 |
| G20007_Hapatesus_tropicus_Elateridae | 3844 | 3428905 | 1050 | 36.27 | 892 | 753 | 8052 | 18 |
| G20008_Eucnemidae_indet_Eucnemidae | 3606 | 3128127 | 1077 | 34.42 | 867.5 | 681 | 22632 | 36 |
| G20009_Drapetes_sp_Elateridae | 3822 | 3669981 | 1173 | 38.89 | 960.2 | 780 | 10950 | 39 |
| G20010_Chelonarium_sp_Chelonaridae | 3844 | 3065102 | 984 | 32.42 | 797.4 | 636 | 9483 | 42 |
| G20011_Balgus_obconicus_Elateridae | 3743 | 3392941 | 1104 | 36.39 | 906.5 | 771 | 8922 | 51 |
| G20012_Tibionema_abdominalis_Elateridae | 3224 | 2033258 | 756 | 24.76 | 630.7 | 546 | 6129 | 18 |
| G20019_Eulichas_dudgeoni_Eulichadidae | 3970 | 4372912 | 1335 | 44.96 | 1101.5 | 885 | 16827 | 60 |
| G20021_Elater_ferrugineus_Elateridae | 3892 | 4502875 | 1428 | 47.49 | 1157 | 945 | 13566 | 30 |
| G20022_Pseudotetralobus_capucinus_Elateridae | 3915 | 3996948 | 1239 | 41.83 | 1020.9 | 840 | 10317 | 18 |
| G20023_Rostricephalus_vitalis_Elateridae | 3748 | 3126899 | 981 | 33.98 | 834.3 | 711 | 10524 | 42 |
| G20024_Aphileus_lucanoides_Elateridae | 3905 | 3877995 | 1191 | 40.45 | 993.1 | 840 | 11334 | 18 |
| G20025_Cheguevaria_sp_Lampyridae | 3824 | 3220794 | 978 | 33.94 | 842.3 | 720 | 6636 | 36 |
| G20028_Stenocladius_sp_Lampyridae | 3826 | 3683030 | 1179 | 38.96 | 962.6 | 807 | 9696 | 18 |
| G20030_Rhagophthalmidae_indet_Rhagophthalmidae | 3625 | 2980140 | 993 | 32.4 | 822.1 | 708 | 6498 | 18 |
| G20031_Rhagophthalmidae_indet_Rhagophthalmidae | 3809 | 3404500 | 1065 | 36.76 | 893.8 | 765 | 6534 | 21 |
| G20033_Euanoma_sp_Elateridae | 3926 | 4529753 | 1383 | 46.95 | 1153.8 | 949.5 | 28551 | 57 |
| G20035_Artematopus_sp_Artematopodidae | 3949 | 4628881 | 1440 | 48.26 | 1172.2 | 939 | 17181 | 39 |
| G21002_Sphaenelater_lineicollis_Elateridae | 3850 | 3675462 | 1146 | 38.76 | 954.7 | 825 | 6156 | 30 |
| G21003_Analastesa_arabica_Elateridae | 3132 | 1883921 | 714 | 22.52 | 601.5 | 505.5 | 5682 | 24 |
| G21004_Paulusiella_serraticornis_Elateridae | 3977 | 4494513 | 1356 | 46.33 | 1130.1 | 921 | 16521 | 24 |
| G21006_Rhipiceridae_indet_Rhipiceridae | 3826 | 4850949 | 1536 | 51.62 | 1267.9 | 1029 | 22974 | 75 |
| G21007_Cneoglossa_sp_Cneoglossidae | 3927 | 5075172 | 1593 | 53.52 | 1292.4 | 1044 | 13347 | 21 |
| G21009_Psilocladus_sp_Lampyridae | 3861 | 3705090 | 1158 | 39.21 | 959.6 | 813 | 6678 | 60 |
| G21010_Amydetinae_indet_Lampyridae | 3988 | 4147517 | 1239 | 42.66 | 1040 | 912 | 10812 | 21 |
| G21011_Ototretinae_indet_Lampyridae | 3964 | 4087103 | 1212 | 42.17 | 1031.1 | 879 | 9810 | 18 |
| G21012_Austroselasia_sp_Elateridae | 3966 | 3927037 | 1191 | 40.9 | 990.2 | 828 | 10350 | 30 |
| G21013_Habashaselasia_nekemtensis_Elateridae | 3946 | 3832898 | 1161 | 39.81 | 971.3 | 823.5 | 8703 | 21 |
| G21014_Elatichrosis_sp_Elateridae | 3897 | 3796952 | 1158 | 39.83 | 974.8 | 810 | 11832 | 30 |
| G21015_Brachypterodrilus_sp_Lampyridae | 3850 | 3453884 | 1071 | 35.98 | 897.1 | 762 | 10593 | 24 |
| G21016_Rhagophthalmidae_indet_Rhagophthalmidae | 3951 | 4182864 | 1257 | 43.39 | 1058.7 | 894 | 10992 | 21 |
| G21017_Lissominae_indet_Elateridae | 3863 | 5109282 | 1620 | 54.07 | 1322.6 | 1068 | 18054 | 24 |
| G21019_Eucnemidae_indet_Eucnemidae | 3233 | 2314446 | 891 | 26.8 | 715.9 | 588 | 8202 | 21 |
| G21023_Ceratogonys_spinicorne_Eucnemidae | 3843 | 3937449 | 1260 | 41.27 | 1024.6 | 858 | 8043 | 27 |
| G21026_Pyrophorini_indet_Elateridae | 3872 | 3879753 | 1200 | 40.76 | 1002 | 849 | 11361 | 27 |
| G21027_Elaterinae_indet_Elateridae | 2882 | 1714261 | 696 | 21.65 | 594.8 | 504 | 5745 | 36 |
| G21028_Dascillidae_indet_Dascillidae | 3917 | 5169855 | 1596 | 54.46 | 1319.9 | 1077 | 23991 | 18 |
| G21029_Drilaster_sp_Lampyridae | 3569 | 2631776 | 885 | 28.7 | 737.4 | 627 | 5838 | 18 |
| G21030_Psephenidae_indet_Psephenidae | 3948 | 4729707 | 1449 | 48.97 | 1198 | 987 | 22773 | 21 |
| G21031_Rousia_sp_Elateridae | 3955 | 5158559 | 1572 | 53.66 | 1304.3 | 1071 | 19677 | 42 |
| G21035_Anaissus_sp_Elateridae | 3950 | 4379852 | 1341 | 45.62 | 1108.8 | 930 | 9069 | 18 |
| G21036_Campyloxenus_pyrothorax_Elateridae | 3855 | 3849522 | 1200 | 41.26 | 998.6 | 825 | 25992 | 21 |
| G21037_Pityobius_anguinus_Elateridae | 3632 | 3133083 | 1047 | 34.61 | 862.6 | 720 | 5556 | 30 |
| G21038_Adelocera_sp_Elateridae | 3836 | 3650993 | 1128 | 38.57 | 951.8 | 795 | 7941 | 18 |
| G21039_Psephenidae_indet_Psephenidae | 3967 | 5050044 | 1569 | 52.61 | 1273 | 1029 | 14847 | 39 |
| G21040_Adelocera_sp_Elateridae | 3864 | 3371832 | 1026 | 35.56 | 872.6 | 741 | 8718 | 27 |
| G21041_Adelocera_sp_Elateridae | 3996 | 5193353 | 1557 | 53.68 | 1299.6 | 1059 | 17658 | 24 |
| G21042_Telegeusis_sp_Omethidae | 3857 | 3913173 | 1254 | 41.43 | 1014.6 | 831 | 12093 | 18 |
| G22002_Plectonotum_laterale_Cantharidae | 3878 | 3795282 | 1158 | 40.12 | 978.7 | 792 | 13233 | 27 |
| G22003_Cerophytum_japonicum_Cerophytiae | 3387 | 2088518 | 720 | 25.34 | 616.6 | 525 | 5565 | 27 |
| G22005_Nyctophyxis_ocellatus_Elateridae | 2642 | 2939250 | 1398 | 41.78 | 1112.5 | 942 | 10320 | 30 |
| G22007_Anaspasis_paralella_Elateridae | 2979 | 1771020 | 693 | 21.82 | 594.5 | 522 | 5472 | 21 |
| G22009_Jurasai_digitusdei_Jurasaidae | 3726 | 3613533 | 1149 | 39.21 | 969.8 | 810 | 12135 | 24 |
| G22015_Protelmidae_indet_Protelmidae | 3892 | 4173155 | 1317 | 43.9 | 1072.2 | 870 | 7833 | 21 |
| G22020_Lutrochidae_indet_Lutrochidae | 3936 | 4671513 | 1467 | 48.93 | 1186.9 | 960 | 17034 | 30 |
| G22022_Lutrochidae_indet_Lutrochidae | 3966 | 5149299 | 1596 | 53.35 | 1298.4 | 1063.5 | 20193 | 18 |
| G22024_Elmidae_indet_Elmidae | 3438 | 4329327 | 1545 | 50.37 | 1259.3 | 1014 | 22797 | 24 |
| G22025_Ototretinae_indet_Lampyridae | 4020 | 4500591 | 1308 | 46.09 | 1119.5 | 960 | 19668 | 18 |
| G22026_Stenocladius_sp_Lampyridae | 3664 | 2877409 | 942 | 31.45 | 785.3 | 660 | 8580 | 21 |
| G22027_Lampyrinae_indet_Lampyridae | 3868 | 4474218 | 1386 | 47.04 | 1156.7 | 942 | 16533 | 27 |
| G19002_Hemiops_sp_Elateridae | 3767 | 2995824 | 954 | 31.84 | 795.3 | 663 | 8235 | 18 |
| Heterocerus_fenestratus_Heteroceridae | 3365 | 2323993 | 810 | 29.4 | 690.6 | 642 | 2799 | 30 |
| Heteromastix_sp_Cantharidae | 3909 | 5063887 | 1587 | 53.61 | 1295.4 | 1068 | 9054 | 39 |
| Hypnoidus_riparius_Elateridae | 3878 | 5172513 | 1623 | 55.49 | 1333.8 | 1101 | 12369 | 21 |
| G19015_Iberobaenia_minuta_Iberobaeniidae | 3751 | 2979515 | 927 | 32.06 | 794.3 | 669 | 10302 | 18 |
| Ignelater_luminosus_Elateridae | 3869 | 5172422 | 1626 | 54.84 | 1336.9 | 1095 | 16506 | 45 |
| R19001_Isorhipis_marmottani_Eucnemidae | 3922 | 5065320 | 1575 | 54.11 | 1291.5 | 1059 | 17457 | 30 |
| Konoplatycis_otome_Lycidae | 3516 | 3219945 | 1119 | 38.61 | 915.8 | 786 | 22563 | 42 |
| Lamprigera_yunnana_Lampyridae | 3757 | 5048256 | 1647 | 54.76 | 1343.7 | 1077 | 16824 | 123 |
| Lamprohiza_splendidula_Lampyridae | 3673 | 4133499 | 1359 | 47.09 | 1125.4 | 960 | 10491 | 30 |
| S1_Lampyris_sardiniae_Lampyridae | 3782 | 4919751 | 1578 | 54.18 | 1300.8 | 1071 | 20238 | 66 |
| Leptinotarsa_decemlineata | 4224 | 5829499 | 1692 | 56.1 | 1380.1 | 1092 | 22080 | 141 |
| Leptolycus_sp_Lycidae | 3797 | 2894637 | 909 | 30.55 | 762.3 | 630 | 10911 | 21 |
| Libnetis_sp_Lycidae | 3732 | 4573291 | 1494 | 51.37 | 1225.4 | 1026 | 10401 | 81 |
| Limonius_californicus_Elateridae | 3942 | 4006845 | 1179 | 41.73 | 1016.4 | 849 | 19680 | 30 |
| R19013_Lissomus_sp_Elateridae | 3841 | 5053152 | 1599 | 54.98 | 1315.6 | 1095 | 10758 | 93 |
| Lucidota_atra_Lampyridae | 3761 | 4702596 | 1527 | 52.46 | 1250.4 | 1038 | 17811 | 18 |
| Lycoprogenthes_sp_Lycidae | 3686 | 4679925 | 1553 | 53.22 | 1269.6 | 1039.5 | 11469 | 27 |
| Lycostomus_kraatzi_Lycidae | 3501 | 4232410 | 1482 | 50.87 | 1208.9 | 1008 | 12150 | 21 |
| Lygistopterus_sanguineus_Lycidae | 3694 | 4650965 | 1518 | 52.47 | 1259.1 | 1044 | 17136 | 57 |
| Lyropaeus_optabilis_Lycidae | 3529 | 4159410 | 1437 | 49.58 | 1178.6 | 978 | 16998 | 39 |
| Ponyalis_sp_Lycidae | 3656 | 4598300 | 1539 | 52.49 | 1257.7 | 1047 | 13419 | 27 |
| R20009_Malthodes_sp_Cantharidae | 3864 | 4762035 | 1479 | 51.78 | 1232.4 | 1053 | 9159 | 36 |
| Melanotus_cribricollis_Elateridae | 3683 | 4977915 | 1650 | 56.34 | 1351.6 | 1095 | 16515 | 84 |
| Melanotus_villosus_Elateridae | 3706 | 4760030 | 1551 | 53.67 | 1284.4 | 1078.5 | 12756 | 87 |
| Micronaspis_floridana_Lampyridae | 3555 | 3764178 | 1305 | 44.46 | 1058.8 | 897 | 7752 | 18 |
| Microphotus_sp_Lampyridae | 3616 | 4266273 | 1452 | 48.77 | 1179.8 | 978 | 14709 | 18 |
| G19011_Diplophoenicus_sp_Elateridae | 3957 | 4948343 | 1512 | 51.57 | 1250.5 | 1038 | 19251 | 21 |
| Negastrius_arenicola_Elateridae | 3956 | 5222051 | 1599 | 55.54 | 1320 | 1093 | 13116 | 30 |
| Notolioon_sp_Byrrhidae | 3256 | 3553218 | 1365 | 47.48 | 1091.3 | 939 | 6825 | 24 |
| Omalisus_fontisbellaquei_Elateridae | 3918 | 4270998 | 1287 | 44.77 | 1090.1 | 924 | 22752 | 24 |
| Onthophagus_taurus | 4224 | 7028760 | 2061 | 67.65 | 1664 | 1269 | 72450 | 165 |
| R19005_Ototretinae_indet_Lampyridae | 3852 | 5062215 | 1599 | 55.02 | 1314.2 | 1074 | 25062 | 24 |
| R20013_Ototretinae_indet_Lampyridae | 3897 | 5015871 | 1575 | 53.68 | 1287.1 | 1068 | 9366 | 27 |
| G19008_Oxynopterus_sp_Elateridae | 3957 | 4964960 | 1530 | 51.63 | 1254.7 | 1035 | 13851 | 18 |
| G19005_Pachyderes_sp_Elateridae | 3959 | 4288501 | 1296 | 44.76 | 1083.2 | 924 | 14121 | 18 |
| G19006_Parasaphes_sp_Elateridae | 3941 | 4805112 | 1476 | 50.25 | 1219.3 | 1008 | 15723 | 18 |
| G19004_Pectocera_sp_Elateridae | 3984 | 4284607 | 1287 | 44.25 | 1075.5 | 906 | 10656 | 45 |
| Phausis_reticulata_Lampyridae | 3733 | 4650336 | 1509 | 52.38 | 1245.7 | 1038 | 15087 | 15 |
| Phengodes_sp_B_Phengodidae | 3675 | 4559814 | 1512 | 52.08 | 1240.8 | 1026 | 19374 | 36 |
| Phengodes_sp_A_Phengodidae | 3928 | 4117941 | 1257 | 42.9 | 1048.4 | 885 | 13119 | 39 |
| Photinus_australis_Lampyridae | 3640 | 4196163 | 1431 | 48.05 | 1152.8 | 985.5 | 11274 | 21 |
| Photinus_carolinus_Lampyridae | 3657 | 4266777 | 1443 | 49 | 1166.7 | 978 | 14697 | 21 |
| Photinus_macdermotti_Lampyridae | 3521 | 4232922 | 1533 | 49.28 | 1202.2 | 987 | 16866 | 18 |
| Photinus_pyralis_Lampyridae | 3894 | 5171889 | 1599 | 55.4 | 1328.2 | 1080 | 23274 | 81 |
| Photinus_scintillans_Lampyridae | 3796 | 4888473 | 1575 | 54.1 | 1287.8 | 1056 | 14706 | 24 |
| Photuris_frontalis_Lampyridae | 3733 | 4737798 | 1551 | 53.07 | 1269.2 | 1056 | 19674 | 27 |
| Photuris_sp_Lampyridae | 3612 | 4446999 | 1551 | 51.39 | 1231.2 | 1017 | 17916 | 18 |
| Phrixothrix_hirtus_Phengodidae | 2878 | 2308903 | 1056 | 35.08 | 802.3 | 651 | 12834 | 69 |
| A01544_Plastocerus_angulosus_Elateridae | 3960 | 4734021 | 1455 | 49.35 | 1195.5 | 984 | 23160 | 18 |
| Platerodrilus_sp_Lycidae | 3283 | 3746477 | 1386 | 48.52 | 1141.2 | 954 | 16650 | 24 |
| Plateros_sp_Lycidae | 3661 | 4730701 | 1569 | 53.78 | 1292.2 | 1059 | 10728 | 18 |
| Ponyalis_quadricollis_Lycidae | 3563 | 4316139 | 1461 | 50.42 | 1211.4 | 1008 | 12417 | 111 |
| R19004_Ptilodactylidae_indet_Ptilodactylidae | 3978 | 5376468 | 1647 | 56.83 | 1351.6 | 1113 | 18630 | 21 |
| Ptosima_undecimmaculata_Buprestidae | 3445 | 3925619 | 1383 | 48.06 | 1139.5 | 954 | 19902 | 36 |
| Pyractomena_borealis_Lampyridae | 3752 | 4763286 | 1554 | 53.18 | 1269.5 | 1053 | 20313 | 39 |
| Pyrearinus_fragilis_Elateridae | 3880 | 5351733 | 1674 | 57.72 | 1379.3 | 1126.5 | 20097 | 33 |
| Pyrocoelia_pectoralis_Lampyridae | 3767 | 4586709 | 1497 | 50.66 | 1217.6 | 1008 | 20751 | 27 |
| R18040_Mangkutanus_sp_Lycidae | 3775 | 4736910 | 1509 | 52.1 | 1254.8 | 1038 | 22884 | 21 |
| R20002_Agrypninae_indet_Elateridae | 3780 | 4756575 | 1530 | 52.88 | 1258.4 | 1074 | 7065 | 99 |
| R20003_Athous_sp_Elateridae | 3945 | 5175687 | 1581 | 54.41 | 1312 | 1086 | 10932 | 48 |
| R20004_Selatosomus_aeneus_Elateridae | 3701 | 4328910 | 1422 | 49.36 | 1169.7 | 1029 | 6624 | 18 |
| R20005_Ampedus_sineatus_Elateridae | 3863 | 5180544 | 1632 | 56.24 | 1341.1 | 1116 | 11379 | 42 |
| R20006_Ampedus_cinnabarinu_Elateridaes | 3898 | 5213094 | 1611 | 55.76 | 1337.4 | 1113 | 9456 | 24 |
| R20007_Anostirus_purpureus_Elateridae | 3877 | 5126688 | 1602 | 55.23 | 1322.3 | 1095 | 21246 | 30 |
| R20008_Agriotes_sp_Elateridae | 3915 | 5205018 | 1608 | 55.05 | 1329.5 | 1119 | 16536 | 24 |
| R20010_Elaterinae_indet_Elateridae | 3898 | 5028669 | 1557 | 53.38 | 1290.1 | 1077 | 9774 | 33 |
| R20011_Alaus_myops_Elateridae | 3361 | 3310656 | 1194 | 42.68 | 985 | 849 | 15435 | 30 |
| R20014_Limnius_perrisi_Elmidae | 3948 | 5296689 | 1629 | 56.22 | 1341.6 | 1116 | 7980 | 33 |
| R20016_Cardiophorus_discicollis_Elateridae | 3931 | 5119500 | 1569 | 54.29 | 1302.3 | 1074 | 9465 | 18 |
| R20017_Eubria_palustris_Psephenidae | 3938 | 5217582 | 1605 | 55.58 | 1324.9 | 1095 | 10737 | 24 |
| R20018_Phosphaenus_hemipterus_Lampyridae | 3898 | 5037714 | 1557 | 53.88 | 1292.4 | 1062 | 11532 | 18 |
| R20019_Dascillus_cervinus_Dascillidae | 3959 | 5160366 | 1587 | 54.55 | 1303.5 | 1080 | 8742 | 27 |
| R20022_Lampyris_noctiluca_Lampyridae | 3914 | 5058009 | 1572 | 53.72 | 1292.3 | 1081.5 | 9105 | 93 |
| Anischia_sp_Eucnemidae | 3939 | 4649169 | 1428 | 48.94 | 1180.3 | 966 | 18729 | 93 |
| Platycrepidius_sp_Elateridae | 3996 | 5140245 | 1569 | 53.1 | 1286.3 | 1059 | 25833 | 27 |
| Lamellipalpus_pacholatkoi_Lampyridae | 4006 | 4487657 | 1305 | 46.13 | 1120.2 | 960 | 10311 | 15 |
| Byrrhinus_sp_Limnichidae | 3949 | 5297163 | 1650 | 55.77 | 1341.4 | 1086 | 17862 | 33 |
| Epilichas_sp_Ptilodactylidae | 3615 | 2724727 | 912 | 30.29 | 753.7 | 609 | 10314 | 36 |
| Rhagonycha_fulva_Cantharidae | 3538 | 4230782 | 1455 | 50.11 | 1195.8 | 1026 | 10122 | 21 |
| Rhagopthalmus_sp_Rhagophthalmidae | 3769 | 4978643 | 1608 | 55.29 | 1320.9 | 1074 | 24870 | 24 |
| S39_Limnichidae_indet_Limnichidae | 3740 | 4588449 | 1512 | 51.09 | 1226.9 | 1023 | 15243 | 18 |
| Sandalus_niger_Rhipiceridae | 3641 | 4382024 | 1461 | 50.31 | 1203.5 | 1011 | 23376 | 18 |
| Scarelus_anthracinus_Lycidae | 3744 | 4715405 | 1521 | 52.28 | 1259.5 | 1032 | 21435 | 72 |
| G19009_Semiotus_sp_Elateridae | 3994 | 4880378 | 1491 | 50.21 | 1221.9 | 982.5 | 17820 | 24 |
| R19009_Silis_sp_Cantharidae | 3947 | 5227968 | 1605 | 55.33 | 1324.5 | 1095 | 17154 | 24 |
| Sinopyrophorus_schimmeli_Sinopyrophoridae | 3845 | 3946086 | 1272 | 42.44 | 1026.3 | 831 | 8025 | 30 |
| G20013_Sinopyrophorus_sp_Sinopyrophoridae | 3854 | 3735747 | 1209 | 39.57 | 969.3 | 777 | 10746 | 21 |
| Taphes_brevicollis_Lycidae | 3710 | 4700466 | 1554 | 52.97 | 1267 | 1053 | 17175 | 78 |
| G19001_Sinelater_perroti_Elateridae | 3905 | 4137345 | 1290 | 43.77 | 1059.5 | 870 | 14109 | 27 |
| Tribolium_castaneum | 4224 | 7438663 | 2238 | 71.59 | 1761 | 1357.5 | 26229 | 192 |
| Trixagus_carinifrons_Throscidae | 3592 | 3443544 | 1152 | 40.06 | 958.7 | 885 | 7758 | 30 |

**NT outliers removed**

| Taxon _name_Family | no_genes | total_no_nt | N50 | percent_determined | average_length | median_length | max_length | min_length |
|---|---|---|---|---|---|---|---|---|
| A4488_Throscidae_indet_Throscidae | 3782 | 3420204 | 1077 | 38.33 | 904.3 | 750 | 16845 | 18 |
| Abscondita_terminalis_Lampyridae | 3729 | 4910688 | 1587 | 56.12 | 1316.9 | 1071 | 16512 | 87 |
| Agrilus_planipennis_Buprestidae | 4137 | 6920836 | 2133 | 69.11 | 1672.9 | 1272 | 27027 | 162 |
| Agrilus_zanthoxylumi_Buprestidae | 3937 | 5785083 | 1839 | 62.53 | 1469.4 | 1167 | 17103 | 45 |
| Agrypnus_murinus_Elateridae | 3640 | 4221705 | 1422 | 49.8 | 1159.8 | 984 | 12312 | 72 |
| Anoplophora_glabripennis | 4152 | 6722131 | 2007 | 66.92 | 1619 | 1233 | 58590 | 177 |
| Anorus_parvicollis_Dascillidae | 3690 | 4343906 | 1434 | 50.63 | 1177.2 | 979.5 | 23625 | 21 |
| Antennolycus_constrictus_Lycidae | 3724 | 4655533 | 1533 | 53.64 | 1250.1 | 1023 | 15021 | 93 |
| Aquatica_ficta_Lampyridae | 3690 | 4284522 | 1434 | 49.69 | 1161.1 | 972 | 14247 | 75 |
| Asymmetricata_circumdata_Lampyridae | 3589 | 4372885 | 1479 | 52.61 | 1218.4 | 1017 | 17178 | 36 |
| Aspisoma_lineatum_Lampyridae | 3896 | 5133292 | 1596 | 56.23 | 1317.6 | 1077 | 21087 | 48 |
| Bicellonycha_wickershamorum_Lampyridae | 3578 | 4054467 | 1398 | 48.63 | 1133.2 | 954 | 15858 | 18 |
| Brachypsectra_fulva_Brachypsectridae | 3548 | 3755681 | 1302 | 46.28 | 1058.5 | 948 | 9879 | 21 |
| Byrrhus_pilula_Byrrhidae | 3299 | 3507766 | 1299 | 46.21 | 1063.3 | 921 | 10842 | 90 |
| R19006_Callirhipidae_indet_Callirhipidae | 3676 | 4542177 | 1527 | 53.12 | 1235.6 | 1003.5 | 22989 | 93 |
| Calopteron_sp_Lycidae | 3652 | 4712642 | 1566 | 55.28 | 1290.4 | 1062 | 16809 | 24 |
| Capnodis_tenebrionis_Buprestidae | 3744 | 4950366 | 1611 | 55.67 | 1322.2 | 1089 | 17751 | 48 |
| R19003_Dicronychus_rubripes_Elateridae | 3879 | 5169573 | 1605 | 57.47 | 1332.7 | 1092 | 24966 | 111 |
| Cautires_communis_Lycidae | 3696 | 3133323 | 990 | 35.24 | 847.8 | 718.5 | 9762 | 33 |
| G19010_Cebrio_sp_Elateridae | 3458 | 2638893 | 918 | 30.9 | 763.1 | 654 | 10119 | 18 |
| Cerophytum_elateroides_Cerophytiae | 3110 | 1762105 | 669 | 23.66 | 566.6 | 474 | 4077 | 21 |
| Chauliognathus_opacus_Cantharidae | 3571 | 2789274 | 936 | 33.13 | 781.1 | 639 | 16404 | 24 |
| Conderis_signicollis_Lycidae | 3653 | 4247919 | 1434 | 49.54 | 1162.9 | 960 | 22641 | 105 |
| Elaterinae_indet_Elateridae | 3902 | 5406238 | 1665 | 58.94 | 1385.5 | 1134 | 18240 | 27 |
| Dendroctonus_ponderosae | 3989 | 6476023 | 2019 | 66.36 | 1623.5 | 1275 | 22002 | 150 |
| R19012_Denticollis_sp_Elateridae | 3895 | 5111103 | 1578 | 56.45 | 1312.2 | 1095 | 12969 | 120 |
| Dexoris_ruzzieri_Lycidae | 3773 | 3003600 | 933 | 32.78 | 796.1 | 678 | 10485 | 60 |
| Dictyoptera_sp_Lycidae | 3336 | 2588771 | 984 | 33.85 | 776 | 633 | 10089 | 21 |
| Dihammatus_sp_Lycidae | 3711 | 4577946 | 1515 | 52.69 | 1233.6 | 1026 | 21366 | 30 |
| Dilophotes_sp_Lycidae | 3701 | 4613055 | 1512 | 53.18 | 1246.4 | 1038 | 10923 | 21 |
| Distremocephalus_sp_Phengodidae | 3610 | 4095107 | 1383 | 49.55 | 1134.4 | 990 | 8559 | 33 |
| Drilonius_sp_Omethidae | 3850 | 5058280 | 1587 | 55.95 | 1313.8 | 1086 | 18432 | 60 |
| G18004_Drilus_mauritanicus_Elateridae | 3957 | 4589781 | 1407 | 49 | 1159.9 | 954 | 19671 | 21 |
| Drilus_concolor_Elateridae | 2974 | 2555432 | 1071 | 37.5 | 859.3 | 726 | 10050 | 57 |
| Dryops_sp_Dryopidae | 3676 | 4507545 | 1500 | 52.88 | 1226.2 | 1029 | 9336 | 51 |
| R19010_Quasimus_sp_Elateridae | 3898 | 5105142 | 1584 | 56.2 | 1309.7 | 1077 | 15192 | 18 |
| R19011_Agrypninae_indet_Elateridae | 3836 | 5029953 | 1599 | 56.13 | 1311.2 | 1095 | 22269 | 24 |
| G19012_Penia_sp_Elateridae | 3912 | 4262726 | 1308 | 45.79 | 1089.7 | 927 | 14322 | 18 |
| G19013_Dromaeolinae indet_Eucnemidae | 3826 | 4311557 | 1377 | 47 | 1126.9 | 921 | 14688 | 63 |
| Ellychnia_sp_Lampyridae | 3702 | 4468845 | 1482 | 51 | 1207.1 | 999 | 13971 | 24 |
| Eucnemis_capucina_Eucnemidae | 3866 | 4867282 | 1536 | 53.07 | 1259 | 1026 | 14343 | 84 |
| G19007_Eudicronychus_sp_Elateridae | 3401 | 2580045 | 921 | 30.43 | 758.6 | 636 | 6657 | 30 |
| Eurypogon_sp_Artematopodidae | 3610 | 3659146 | 1212 | 44.1 | 1013.6 | 942 | 4332 | 36 |
| G20001_Octinodes_sp_Elateridae | 3907 | 4236987 | 1317 | 45.51 | 1084.5 | 885 | 10902 | 93 |
| G20002_Pyrearinus_sp_Elateridae | 3969 | 4887029 | 1491 | 51.75 | 1231.3 | 1017 | 14178 | 63 |
| G20003_Cussolenis_sp_Elateridae | 3824 | 4070427 | 1320 | 44.36 | 1064.4 | 852 | 25869 | 36 |
| G20004_Eudicronychus_rufus_Elateridae | 3354 | 2416560 | 876 | 28.81 | 720.5 | 609 | 4410 | 21 |
| G20005_Mulsanteus_sp_Elateridae | 3923 | 3964134 | 1197 | 42.69 | 1010.5 | 840 | 10083 | 21 |
| G20006_Elaterinae_indet_Elateridae | 3935 | 4473754 | 1380 | 47.82 | 1136.9 | 924 | 10437 | 27 |
| G20007_Hapatesus_tropicus_Elateridae | 3807 | 3406698 | 1050 | 37.39 | 894.9 | 756 | 8052 | 18 |
| G20008_Eucnemidae_indet_Eucnemidae | 3507 | 3060951 | 1080 | 35.45 | 872.8 | 687 | 22632 | 36 |
| G20009_Drapetes_sp_Elateridae | 3789 | 3646017 | 1179 | 39.94 | 962.3 | 780 | 10950 | 39 |
| G20010_Chelonarium_sp_Chelonaridae | 3768 | 3021413 | 990 | 33.52 | 801.9 | 639 | 9483 | 42 |
| G20011_Balgus_obconicus_Elateridae | 3676 | 3340846 | 1104 | 37.44 | 908.8 | 774 | 8904 | 51 |
| G20012_Tibionema_abdominalis_Elateridae | 3043 | 1936592 | 762 | 25.74 | 636.4 | 552 | 6129 | 18 |
| G20019_Eulichas_dudgeoni_Eulichadidae | 3947 | 4349100 | 1335 | 46.14 | 1101.9 | 885 | 16827 | 60 |
| G20021_Elater_ferrugineus_Elateridae | 3863 | 4485373 | 1431 | 48.77 | 1161.1 | 948 | 13566 | 30 |
| G20022_Pseudotetralobus_capucinus_Elateridae | 3878 | 3967131 | 1242 | 42.94 | 1023 | 843 | 10317 | 18 |
| G20023_Rostricephalus_vitalis_Elateridae | 3671 | 3078886 | 984 | 35.13 | 838.7 | 717 | 10524 | 45 |
| G20024_Aphileus_lucanoides_Elateridae | 3866 | 3847895 | 1197 | 41.53 | 995.3 | 840 | 11334 | 18 |
| G20025_Cheguevaria_sp_Lampyridae | 3792 | 3199917 | 978 | 34.88 | 843.9 | 720 | 6636 | 75 |
| G20028_Stenocladius_sp_Lampyridae | 3783 | 3656152 | 1182 | 39.96 | 966.5 | 810 | 9696 | 18 |
| G20030_Rhagophthalmidae_indet_Rhagophthalmidae | 3568 | 2943599 | 993 | 33.43 | 825 | 708 | 6495 | 18 |
| G20031_Rhagophthalmidae_indet_Rhagophthalmidae | 3759 | 3369604 | 1068 | 37.92 | 896.4 | 765 | 6534 | 21 |
| G20033_Euanoma_sp_Elateridae | 3892 | 4502120 | 1383 | 48.29 | 1156.8 | 951 | 28551 | 57 |
| G20035_Artematopus_sp_Artematopodidae | 3917 | 4607762 | 1440 | 49.46 | 1176.3 | 939 | 17181 | 72 |
| G21002_Sphaenelater_lineicollis_Elateridae | 3810 | 3646590 | 1146 | 39.93 | 957.1 | 828 | 6156 | 30 |
| G21003_Analastesa_arabica_Elateridae | 2869 | 1714352 | 708 | 22.71 | 597.5 | 504 | 5361 | 24 |
| G21004_Paulusiella_serraticornis_Elateridae | 3950 | 4464231 | 1359 | 47.45 | 1130.2 | 921 | 16521 | 24 |
| G21006_Rhipiceridae_indet_Rhipiceridae | 3769 | 4800942 | 1545 | 53.14 | 1273.8 | 1035 | 22974 | 75 |
| G21007_Cneoglossa_sp_Cneoglossidae | 3901 | 5056800 | 1596 | 54.87 | 1296.3 | 1047 | 13347 | 21 |
| G21009_Psilocladus_sp_Lampyridae | 3829 | 3676383 | 1161 | 40.23 | 960.1 | 813 | 6678 | 84 |
| G21010_Amydetinae_indet_Lampyridae | 3960 | 4125702 | 1239 | 43.85 | 1041.8 | 915 | 10812 | 21 |
| G21011_Ototretinae_indet_Lampyridae | 3944 | 4071248 | 1212 | 43.26 | 1032.3 | 882 | 9810 | 18 |
| G21012_Austroselasia_sp_Elateridae | 3928 | 3890164 | 1191 | 41.98 | 990.4 | 829.5 | 10350 | 30 |
| G21013_Habashaselasia_nekemtensis_Elateridae | 3910 | 3801031 | 1161 | 40.81 | 972.1 | 825 | 8703 | 21 |
| G21014_Elatichrosis_sp_Elateridae | 3870 | 3776739 | 1161 | 40.86 | 975.9 | 810 | 11832 | 30 |
| G21015_Brachypterodrilus_sp_Lampyridae | 3798 | 3417468 | 1071 | 36.91 | 899.8 | 765 | 10593 | 24 |
| G21016_Rhagophthalmidae_indet_Rhagophthalmidae | 3931 | 4165620 | 1257 | 44.57 | 1059.7 | 897 | 10992 | 21 |
| G21017_Lissominae_indet_Elateridae | 3842 | 5090835 | 1620 | 55.51 | 1325 | 1071 | 18054 | 24 |
| G21019_Eucnemidae_indet_Eucnemidae | 3025 | 2169935 | 897 | 27.24 | 717.3 | 588 | 8199 | 21 |
| G21023_Ceratogonys_spinicorne_Eucnemidae | 3778 | 3892087 | 1266 | 42.41 | 1030.2 | 867 | 8043 | 27 |
| G21026_Pyrophorini_indet_Elateridae | 3832 | 3845325 | 1203 | 41.83 | 1003.5 | 849 | 11361 | 27 |
| G21027_Elaterinae_indet_Elateridae | 2665 | 1583721 | 696 | 22.19 | 594.3 | 507 | 5745 | 36 |
| G21028_Dascillidae_indet_Dascillidae | 3894 | 5153058 | 1596 | 55.87 | 1323.3 | 1080 | 23991 | 18 |
| G21029_Drilaster_sp_Lampyridae | 3463 | 2550821 | 879 | 29.35 | 736.6 | 627 | 5832 | 18 |
| G21030_Psephenidae_indet_Psephenidae | 3906 | 4695627 | 1452 | 50.38 | 1202.2 | 994.5 | 22773 | 21 |
| G21031_Rousia_sp_Elateridae | 3935 | 5146874 | 1572 | 55.1 | 1308 | 1074 | 19677 | 111 |
| G21035_Anaissus_sp_Elateridae | 3924 | 4355586 | 1344 | 46.89 | 1110 | 930 | 9069 | 18 |
| G21036_Campyloxenus_pyrothorax_Elateridae | 3814 | 3819300 | 1200 | 42.38 | 1001.4 | 828 | 25992 | 21 |
| G21037_Pityobius_anguinus_Elateridae | 3486 | 3023361 | 1047 | 35.74 | 867.3 | 723 | 5556 | 30 |
| G21038_Adelocera_sp_Elateridae | 3790 | 3611785 | 1134 | 39.6 | 953 | 795 | 7941 | 18 |
| G21039_Psephenidae_indet_Psephenidae | 3937 | 5026977 | 1569 | 54.05 | 1276.9 | 1035 | 14847 | 39 |
| G21040_Adelocera_sp_Elateridae | 3810 | 3329576 | 1026 | 36.66 | 873.9 | 741 | 8718 | 27 |
| G21041_Adelocera_sp_Elateridae | 3982 | 5180624 | 1557 | 55.05 | 1301 | 1062 | 17616 | 24 |
| G21042_Telegeusis_sp_Omethidae | 3804 | 3868125 | 1254 | 42.58 | 1016.9 | 832.5 | 12093 | 18 |
| G22002_Plectonotum_laterale_Cantharidae | 3817 | 3754545 | 1164 | 41.24 | 983.6 | 795 | 13233 | 27 |
| G22003_Cerophytum_japonicum_Cerophytiae | 3260 | 2024262 | 726 | 26.21 | 620.9 | 531 | 5565 | 27 |
| G22005_Nyctophyxis_ocellatus_Elateridae | 2578 | 2888523 | 1410 | 43.18 | 1120.5 | 946.5 | 10320 | 30 |
| G22007_Anaspasis_paralella_Elateridae | 2810 | 1678425 | 696 | 22.72 | 597.3 | 522 | 5469 | 21 |
| G22009_Jurasai_digitusdei_Jurasaidae | 3637 | 3545604 | 1155 | 40.29 | 974.9 | 816 | 12135 | 24 |
| G22015_Protelmidae_indet_Protelmidae | 3845 | 4137278 | 1320 | 45.01 | 1076 | 870 | 7833 | 21 |
| G22020_Lutrochidae_indet_Lutrochidae | 3908 | 4654545 | 1470 | 50.2 | 1191 | 963 | 17034 | 30 |
| G22022_Lutrochidae_indet_Lutrochidae | 3937 | 5127783 | 1596 | 54.86 | 1302.5 | 1065 | 20193 | 18 |
| G22024_Elmidae_indet_Elmidae | 3411 | 4311417 | 1551 | 51.77 | 1264 | 1020 | 22797 | 24 |
| G22025_Ototretinae_indet_Lampyridae | 4004 | 4487238 | 1308 | 47.36 | 1120.7 | 961.5 | 19668 | 18 |
| G22026_Stenocladius_sp_Lampyridae | 3579 | 2813036 | 939 | 32.22 | 786 | 660 | 8580 | 21 |
| G22027_Lampyrinae_indet_Lampyridae | 3844 | 4450656 | 1383 | 48.24 | 1157.8 | 942 | 16533 | 27 |
| G19002_Hemiops_sp_Elateridae | 3689 | 2945491 | 957 | 32.86 | 798.5 | 666 | 8235 | 18 |
| Heterocerus_fenestratus_Heteroceridae | 3246 | 2260787 | 819 | 30.92 | 696.5 | 645 | 2799 | 30 |
| Heteromastix_sp_Cantharidae | 3897 | 5052148 | 1587 | 54.93 | 1296.4 | 1068 | 9054 | 39 |
| Hypnoidus_riparius_Elateridae | 3869 | 5167045 | 1623 | 56.98 | 1335.5 | 1101 | 12336 | 21 |
| G19015_Iberobaenia_minuta_Iberobaeniidae | 3676 | 2934552 | 930 | 33.08 | 798.3 | 672 | 10302 | 18 |
| Ignelater_luminosus_Elateridae | 3855 | 5162240 | 1632 | 56.37 | 1339.1 | 1095 | 16506 | 45 |
| R19001_Isorhipis_marmottani_Eucnemidae | 3896 | 5042010 | 1581 | 55.62 | 1294.2 | 1062 | 17457 | 30 |
| Konoplatycis_otome_Lycidae | 3477 | 3198089 | 1122 | 39.9 | 919.8 | 792 | 22563 | 42 |
| Lamprigera_yunnana_Lampyridae | 3737 | 5036421 | 1647 | 56.14 | 1347.7 | 1086 | 16824 | 123 |
| Lamprohiza_splendidula_Lampyridae | 3654 | 4120056 | 1359 | 48.41 | 1127.5 | 960 | 10491 | 30 |
| S1_Lampyris_sardiniae_Lampyridae | 3764 | 4902534 | 1581 | 55.85 | 1302.5 | 1074 | 20238 | 66 |
| Leptinotarsa_decemlineata | 4042 | 5664069 | 1710 | 57.47 | 1401.3 | 1107 | 22080 | 141 |
| Leptolycus_sp_Lycidae | 3730 | 2847672 | 909 | 31.41 | 763.5 | 630 | 10911 | 21 |
| Libnetis_sp_Lycidae | 3710 | 4557815 | 1494 | 52.82 | 1228.5 | 1029 | 10350 | 81 |
| Limonius_californicus_Elateridae | 3916 | 3983133 | 1179 | 42.93 | 1017.1 | 852 | 19680 | 30 |
| R19013_Lissomus_sp_Elateridae | 3827 | 5039805 | 1602 | 56.5 | 1316.9 | 1095 | 10722 | 114 |
| Lucidota_atra_Lampyridae | 3753 | 4695639 | 1527 | 53.75 | 1251.2 | 1038 | 17811 | 18 |
| Lycoprogenthes_sp_Lycidae | 3678 | 4669536 | 1551 | 54.57 | 1269.6 | 1039.5 | 11469 | 27 |
| Lycostomus_kraatzi_Lycidae | 3459 | 4199462 | 1482 | 52.57 | 1214.1 | 1014 | 12150 | 21 |
| Lygistopterus_sanguineus_Lycidae | 3678 | 4638277 | 1518 | 53.88 | 1261.1 | 1045.5 | 17133 | 57 |
| Lyropaeus_optabilis_Lycidae | 3503 | 4143977 | 1443 | 51.16 | 1183 | 984 | 16998 | 39 |
| Ponyalis_sp_Lycidae | 3640 | 4582268 | 1539 | 54.06 | 1258.9 | 1047 | 13419 | 27 |
| R20009_Malthodes_sp_Cantharidae | 3840 | 4740801 | 1479 | 53.2 | 1234.6 | 1053 | 9159 | 36 |
| Melanotus_cribricollis_Elateridae | 3673 | 4968212 | 1650 | 57.8 | 1352.6 | 1098 | 16515 | 84 |
| Melanotus_villosus_Elateridae | 3690 | 4742661 | 1551 | 55.29 | 1285.3 | 1080 | 12756 | 87 |
| Micronaspis_floridana_Lampyridae | 3532 | 3748533 | 1308 | 45.82 | 1061.3 | 900 | 7752 | 18 |
| Microphotus_sp_Lampyridae | 3580 | 4242450 | 1455 | 50.44 | 1185 | 981 | 14709 | 18 |
| G19011_Diplophoenicus_sp_Elateridae | 3939 | 4934096 | 1512 | 53.08 | 1252.6 | 1041 | 19251 | 21 |
| Negastrius_arenicola_Elateridae | 3952 | 5219093 | 1599 | 57.06 | 1320.6 | 1094.5 | 13110 | 30 |
| Notolioon_sp_Byrrhidae | 3218 | 3521621 | 1371 | 49.46 | 1094.4 | 942 | 6825 | 24 |
| Omalisus_fontisbellaquei_Elateridae | 3890 | 4248780 | 1287 | 45.89 | 1092.2 | 927 | 22752 | 24 |
| Onthophagus_taurus | 4114 | 6894426 | 2073 | 69.17 | 1675.8 | 1275 | 72450 | 165 |
| R19005_Ototretinae_indet_Lampyridae | 3836 | 5051856 | 1602 | 56.6 | 1317 | 1075.5 | 25062 | 24 |
| R20013_Ototretinae_indet_Lampyridae | 3890 | 5006031 | 1575 | 55.04 | 1286.9 | 1066.5 | 9366 | 27 |
| G19008_Oxynopterus_sp_Elateridae | 3934 | 4948016 | 1533 | 53.04 | 1257.8 | 1036.5 | 13851 | 18 |
| G19005_Pachyderes_sp_Elateridae | 3934 | 4269677 | 1296 | 45.96 | 1085.3 | 924 | 14121 | 18 |
| G19006_Parasaphes_sp_Elateridae | 3921 | 4793085 | 1476 | 51.64 | 1222.4 | 1011 | 15717 | 18 |
| G19004_Pectocera_sp_Elateridae | 3951 | 4255847 | 1290 | 45.55 | 1077.2 | 909 | 10656 | 93 |
| Phausis_reticulata_Lampyridae | 3713 | 4633605 | 1512 | 53.79 | 1247.9 | 1038 | 15087 | 15 |
| Phengodes_sp_B_Phengodidae | 3662 | 4548357 | 1515 | 53.63 | 1242 | 1026 | 19374 | 36 |
| Phengodes_sp_A_Phengodidae | 3885 | 4078605 | 1257 | 44.05 | 1049.8 | 885 | 13119 | 39 |
| Photinus_australis_Lampyridae | 3606 | 4178976 | 1434 | 49.9 | 1158.9 | 990 | 11274 | 21 |
| Photinus_carolinus_Lampyridae | 3634 | 4252545 | 1446 | 50.55 | 1170.2 | 979.5 | 14697 | 21 |
| Photinus_macdermotti_Lampyridae | 3473 | 4205946 | 1533 | 51.31 | 1211 | 993 | 16866 | 18 |
| Photinus_pyralis_Lampyridae | 3886 | 5163762 | 1599 | 56.8 | 1328.8 | 1080 | 23274 | 81 |
| Photinus_scintillans_Lampyridae | 3789 | 4874979 | 1575 | 55.51 | 1286.6 | 1056 | 14706 | 24 |
| Photuris_frontalis_Lampyridae | 3722 | 4729098 | 1551 | 54.52 | 1270.6 | 1056 | 19674 | 27 |
| Photuris_sp_Lampyridae | 3586 | 4431909 | 1551 | 52.96 | 1235.9 | 1023 | 17916 | 18 |
| Phrixothrix_hirtus_Phengodidae | 2828 | 2277415 | 1062 | 36.56 | 805.3 | 651 | 12834 | 69 |
| A01544_Plastocerus_angulosus_Elateridae | 3943 | 4717929 | 1455 | 50.58 | 1196.5 | 984 | 23160 | 18 |
| Platerodrilus_sp_Lycidae | 3252 | 3718274 | 1389 | 49.94 | 1143.4 | 954 | 16644 | 24 |
| Plateros_sp_Lycidae | 3646 | 4718531 | 1572 | 55.31 | 1294.2 | 1059 | 10728 | 18 |
| Ponyalis_quadricollis_Lycidae | 3546 | 4299480 | 1464 | 52.09 | 1212.5 | 1008 | 12417 | 111 |
| R19004_Ptilodactylidae_indet_Ptilodactylidae | 3964 | 5361657 | 1650 | 58.34 | 1352.6 | 1116 | 18630 | 21 |
| Ptosima_undecimmaculata_Buprestidae | 3388 | 3882710 | 1395 | 49.77 | 1146 | 960 | 19902 | 36 |
| Pyractomena_borealis_Lampyridae | 3743 | 4755840 | 1554 | 54.51 | 1270.6 | 1053 | 20313 | 39 |
| Pyrearinus_fragilis_Elateridae | 3871 | 5345208 | 1674 | 59.24 | 1380.8 | 1128 | 20097 | 72 |
| Pyrocoelia_pectoralis_Lampyridae | 3755 | 4578261 | 1497 | 52.08 | 1219.2 | 1008 | 20727 | 27 |
| R18040_Mangkutanus_sp_Lycidae | 3749 | 4710316 | 1512 | 53.57 | 1256.4 | 1038 | 22884 | 21 |
| R20002_Agrypninae_indet_Elateridae | 3763 | 4740720 | 1533 | 54.6 | 1259.8 | 1077 | 7065 | 99 |
| R20003_Athous_sp_Elateridae | 3925 | 5155962 | 1581 | 56.23 | 1313.6 | 1086 | 10929 | 69 |
| R20004_Selatosomus_aeneus_Elateridae | 3672 | 4303953 | 1425 | 51.07 | 1172.1 | 1032 | 6624 | 18 |
| R20005_Ampedus_sineatus_Elateridae | 3857 | 5175768 | 1632 | 57.69 | 1341.9 | 1116 | 11379 | 60 |
| R20006_Ampedus_cinnabarinu_Elateridaes | 3884 | 5201922 | 1611 | 57.19 | 1339.3 | 1116 | 9456 | 24 |
| R20007_Anostirus_purpureus_Elateridae | 3864 | 5113554 | 1602 | 56.96 | 1323.4 | 1095 | 21234 | 102 |
| R20008_Agriotes_sp_Elateridae | 3899 | 5187210 | 1608 | 56.64 | 1330.4 | 1119 | 16536 | 24 |
| R20010_Elaterinae_indet_Elateridae | 3884 | 5019066 | 1557 | 54.77 | 1292.2 | 1080 | 9774 | 33 |
| R20011_Alaus_myops_Elateridae | 3319 | 3283530 | 1200 | 44.51 | 989.3 | 852 | 15435 | 30 |
| R20014_Limnius_perrisi_Elmidae | 3932 | 5285865 | 1632 | 57.78 | 1344.3 | 1119 | 7980 | 60 |
| R20016_Cardiophorus_discicollis_Elateridae | 3922 | 5109330 | 1569 | 55.73 | 1302.7 | 1074 | 9465 | 18 |
| R20017_Eubria_palustris_Psephenidae | 3923 | 5206887 | 1605 | 56.98 | 1327.3 | 1095 | 10734 | 24 |
| R20018_Phosphaenus_hemipterus_Lampyridae | 3882 | 5031021 | 1560 | 55.36 | 1296 | 1065 | 11532 | 18 |
| R20019_Dascillus_cervinus_Dascillidae | 3945 | 5149539 | 1587 | 56.2 | 1305.3 | 1083 | 8742 | 27 |
| R20022_Lampyris_noctiluca_Lampyridae | 3906 | 5048961 | 1572 | 55.15 | 1292.6 | 1083 | 9105 | 93 |
| Anischia_sp_Eucnemidae | 3884 | 4609917 | 1437 | 50.25 | 1186.9 | 972 | 18729 | 93 |
| Platycrepidius_sp_Elateridae | 3979 | 5126943 | 1575 | 54.43 | 1288.5 | 1059 | 25833 | 27 |
| Lamellipalpus_pacholatkoi_Lampyridae | 3995 | 4473103 | 1305 | 47.24 | 1119.7 | 960 | 10311 | 15 |
| Byrrhinus_sp_Limnichidae | 3924 | 5279412 | 1653 | 57.21 | 1345.4 | 1089 | 17862 | 33 |
| Epilichas_sp_Ptilodactylidae | 3503 | 2647999 | 912 | 31.19 | 755.9 | 609 | 10284 | 36 |
| Rhagonycha_fulva_Cantharidae | 3480 | 4184336 | 1461 | 51.63 | 1202.4 | 1032 | 10122 | 21 |
| Rhagopthalmus_sp_Rhagophthalmidae | 3759 | 4972079 | 1608 | 56.73 | 1322.7 | 1077 | 24870 | 24 |
| S39_Limnichidae_indet_Limnichidae | 3700 | 4557528 | 1521 | 52.59 | 1231.8 | 1029 | 15243 | 18 |
| Sandalus_niger_Rhipiceridae | 3582 | 4332674 | 1464 | 52.01 | 1209.6 | 1017 | 23376 | 18 |
| Scarelus_anthracinus_Lycidae | 3727 | 4697103 | 1521 | 53.94 | 1260.3 | 1032 | 21435 | 99 |
| G19009_Semiotus_sp_Elateridae | 3978 | 4862522 | 1491 | 51.55 | 1222.4 | 984 | 17820 | 24 |
| R19009_Silis_sp_Cantharidae | 3895 | 5175177 | 1611 | 56.78 | 1328.7 | 1101 | 17154 | 24 |
| Sinopyrophorus_schimmeli_Sinopyrophoridae | 3817 | 3928765 | 1278 | 43.55 | 1029.3 | 834 | 8025 | 30 |
| G20013_Sinopyrophorus_sp_Sinopyrophoridae | 3812 | 3702840 | 1209 | 40.71 | 971.4 | 780 | 10746 | 21 |
| Taphes_brevicollis_Lycidae | 3697 | 4689055 | 1554 | 54.32 | 1268.3 | 1053 | 17175 | 78 |
| G19001_Sinelater_perroti_Elateridae | 3860 | 4095379 | 1290 | 45.02 | 1061 | 870 | 14109 | 27 |
| Tribolium_castaneum | 4197 | 7401620 | 2241 | 73.52 | 1763.6 | 1359 | 26229 | 192 |
| Trixagus_carinifrons_Throscidae | 3505 | 3387861 | 1158 | 41.93 | 966.6 | 894 | 7677 | 30 |

**NT after AliScore filtering**

| Taxon _name_Family | no_genes | total_no_nt | N50 | percent_determined | average_length | median_length | max_length | min_length |
|---|---|---|---|---|---|---|---|---|
| A4488_Throscidae_indet_Throscidae | 3741 | 3008328 | 960 | 60.19 | 804.2 | 669 | 14136 | 39 |
| Abscondita_terminalis_Lampyridae | 3723 | 4310640 | 1389 | 87.22 | 1157.8 | 957 | 16146 | 69 |
| Agrilus_planipennis_Buprestidae | 4102 | 4990372 | 1455 | 90.42 | 1216.6 | 987 | 21333 | 138 |
| Agrilus_zanthoxylumi_Buprestidae | 3910 | 4613880 | 1428 | 88.55 | 1180 | 972 | 16710 | 39 |
| Agrypnus_murinus_Elateridae | 3604 | 3698733 | 1260 | 77.77 | 1026.3 | 877.5 | 12096 | 63 |
| Anoplophora_glabripennis | 4116 | 4876415 | 1410 | 88.6 | 1184.7 | 978 | 14187 | 168 |
| Anorus_parvicollis_Dascillidae | 3657 | 3790567 | 1266 | 78.47 | 1036.5 | 873 | 21351 | 66 |
| Antennolycus_constrictus_Lycidae | 3710 | 4141874 | 1350 | 84.01 | 1116.4 | 930 | 14826 | 57 |
| Aquatica_ficta_Lampyridae | 3655 | 3797954 | 1287 | 78.61 | 1039.1 | 876 | 13626 | 48 |
| Asymmetricata_circumdata_Lampyridae | 3551 | 3880003 | 1329 | 82.67 | 1092.7 | 918 | 16146 | 36 |
| Aspisoma_lineatum_Lampyridae | 3889 | 4536792 | 1410 | 87.83 | 1166.6 | 960 | 19599 | 72 |
| Bicellonycha_wickershamorum_Lampyridae | 3528 | 3586068 | 1263 | 77.27 | 1016.5 | 861 | 14802 | 63 |
| Brachypsectra_fulva_Brachypsectridae | 3493 | 3301776 | 1155 | 72.34 | 945.3 | 846 | 9519 | 69 |
| Byrrhus_pilula_Byrrhidae | 3237 | 3044453 | 1158 | 72.97 | 940.5 | 816 | 10551 | 66 |
| R19006_Callirhipidae_indet_Callirhipidae | 3645 | 3964539 | 1332 | 82.6 | 1087.7 | 903 | 20382 | 51 |
| Calopteron_sp_Lycidae | 3642 | 4204264 | 1386 | 86.42 | 1154.4 | 960 | 16584 | 36 |
| Capnodis_tenebrionis_Buprestidae | 3703 | 4149957 | 1350 | 83.83 | 1120.7 | 936 | 16710 | 57 |
| R19003_Dicronychus_rubripes_Elateridae | 3874 | 4530510 | 1410 | 88.43 | 1169.5 | 972 | 21351 | 84 |
| Cautires_communis_Lycidae | 3638 | 2781564 | 903 | 56.78 | 764.6 | 651 | 9642 | 75 |
| G19010_Cebrio_sp_Elateridae | 3386 | 2274031 | 804 | 48.91 | 671.6 | 576 | 8016 | 39 |
| Cerophytum_elateroides_Cerophytiae | 2977 | 1548974 | 609 | 39.63 | 520.3 | 432 | 4077 | 51 |
| Chauliognathus_opacus_Cantharidae | 3489 | 2508276 | 876 | 54.18 | 718.9 | 591 | 16032 | 48 |
| Conderis_signicollis_Lycidae | 3631 | 3808524 | 1293 | 78.74 | 1048.9 | 879 | 20349 | 57 |
| Elaterinae_indet_Elateridae | 3889 | 4698109 | 1449 | 90.61 | 1208.1 | 999 | 17817 | 90 |
| Dendroctonus_ponderosae | 3952 | 4836753 | 1455 | 90.16 | 1223.9 | 999 | 21297 | 132 |
| R19012_Denticollis_sp_Elateridae | 3885 | 4467465 | 1392 | 87.18 | 1149.9 | 969 | 12747 | 48 |
| Dexoris_ruzzieri_Lycidae | 3710 | 2659203 | 846 | 53.3 | 716.8 | 609 | 8679 | 60 |
| Dictyoptera_sp_Lycidae | 3183 | 2315577 | 921 | 56.65 | 727.5 | 603 | 9819 | 60 |
| Dihammatus_sp_Lycidae | 3685 | 4069115 | 1341 | 83 | 1104.2 | 924 | 18153 | 78 |
| Dilophotes_sp_Lycidae | 3680 | 4126013 | 1356 | 84.07 | 1121.2 | 942 | 10578 | 84 |
| Distremocephalus_sp_Phengodidae | 3571 | 3625555 | 1233 | 77.44 | 1015.3 | 882 | 7278 | 90 |
| Drilonius_sp_Omethidae | 3837 | 4436502 | 1392 | 86.73 | 1156.2 | 963 | 18087 | 69 |
| G18004_Drilus_mauritanicus_Elateridae | 3937 | 4048287 | 1248 | 77.03 | 1028.3 | 855 | 19278 | 72 |
| Drilus_concolor_Elateridae | 2876 | 2222942 | 954 | 60.43 | 772.9 | 649.5 | 7683 | 60 |
| Dryops_sp_Dryopidae | 3655 | 3964597 | 1320 | 82 | 1084.7 | 927 | 9138 | 66 |
| R19010_Quasimus_sp_Elateridae | 3886 | 4490883 | 1389 | 87.55 | 1155.7 | 963 | 14958 | 60 |
| R19011_Agrypninae_indet_Elateridae | 3814 | 4384461 | 1386 | 86.71 | 1149.6 | 966 | 20466 | 72 |
| G19012_Penia_sp_Elateridae | 3875 | 3703055 | 1158 | 71.74 | 955.6 | 819 | 13695 | 60 |
| G19013_Dromaeolinae indet_Eucnemidae | 3802 | 3778550 | 1221 | 73.81 | 993.8 | 831 | 14079 | 57 |
| Ellychnia_sp_Lampyridae | 3649 | 3976548 | 1344 | 81.74 | 1089.8 | 909 | 12591 | 48 |
| Eucnemis_capucina_Eucnemidae | 3850 | 4295723 | 1359 | 83.97 | 1115.8 | 921 | 14100 | 84 |
| G19007_Eudicronychus_sp_Elateridae | 3316 | 2232527 | 813 | 49.42 | 673.3 | 570 | 6315 | 48 |
| Eurypogon_sp_Artematopodidae | 3530 | 3198578 | 1086 | 69.27 | 906.1 | 839.5 | 3924 | 72 |
| G20001_Octinodes_sp_Elateridae | 3878 | 3684381 | 1164 | 70.74 | 950.1 | 774 | 10077 | 48 |
| G20002_Pyrearinus_sp_Elateridae | 3951 | 4244441 | 1305 | 80.42 | 1074.3 | 897 | 13593 | 81 |
| G20003_Cussolenis_sp_Elateridae | 3788 | 3542286 | 1170 | 69.3 | 935.1 | 759 | 21357 | 81 |
| G20004_Eudicronychus_rufus_Elateridae | 3248 | 2083468 | 792 | 46.93 | 641.5 | 534 | 4041 | 60 |
| G20005_Mulsanteus_sp_Elateridae | 3884 | 3458646 | 1062 | 66.89 | 890.5 | 750 | 9945 | 42 |
| G20006_Elaterinae_indet_Elateridae | 3917 | 3870598 | 1206 | 74.08 | 988.2 | 807 | 10278 | 66 |
| G20007_Hapatesus_tropicus_Elateridae | 3752 | 2954124 | 933 | 58.45 | 787.3 | 675 | 7869 | 63 |
| G20008_Eucnemidae_indet_Eucnemidae | 3444 | 2673279 | 972 | 56.99 | 776.2 | 612 | 20361 | 51 |
| G20009_Drapetes_sp_Elateridae | 3745 | 3201560 | 1050 | 63.36 | 854.9 | 696 | 9147 | 57 |
| G20010_Chelonarium_sp_Chelonaridae | 3710 | 2589836 | 861 | 51.99 | 698.1 | 559.5 | 7125 | 60 |
| G20011_Balgus_obconicus_Elateridae | 3608 | 2929730 | 996 | 59.91 | 812 | 682.5 | 8682 | 39 |
| G20012_Tibionema_abdominalis_Elateridae | 2904 | 1675139 | 693 | 42.59 | 576.8 | 498 | 5622 | 45 |
| G20019_Eulichas_dudgeoni_Eulichadidae | 3926 | 3711031 | 1140 | 70.96 | 945.2 | 774 | 16431 | 69 |
| G20021_Elater_ferrugineus_Elateridae | 3834 | 3870145 | 1245 | 75.61 | 1009.4 | 843 | 13113 | 66 |
| G20022_Pseudotetralobus_capucinus_Elateridae | 3837 | 3458487 | 1107 | 67.55 | 901.4 | 744 | 10260 | 60 |
| G20023_Rostricephalus_vitalis_Elateridae | 3590 | 2671843 | 894 | 55.61 | 744.2 | 639 | 10305 | 63 |
| G20024_Aphileus_lucanoides_Elateridae | 3814 | 3349953 | 1062 | 65.28 | 878.3 | 738 | 10224 | 42 |
| G20025_Cheguevaria_sp_Lampyridae | 3734 | 2838247 | 891 | 56.51 | 760.1 | 651 | 5523 | 60 |
| G20028_Stenocladius_sp_Lampyridae | 3732 | 3231838 | 1050 | 63.96 | 866 | 732 | 9672 | 48 |
| G20030_Rhagophthalmidae_indet_Rhagophthalmidae | 3498 | 2587145 | 897 | 54.21 | 739.6 | 634.5 | 6018 | 66 |
| G20031_Rhagophthalmidae_indet_Rhagophthalmidae | 3708 | 2970244 | 963 | 60.27 | 801 | 684 | 6081 | 69 |
| G20033_Euanoma_sp_Elateridae | 3867 | 3918899 | 1218 | 75.6 | 1013.4 | 840 | 21357 | 63 |
| G20035_Artematopus_sp_Artematopodidae | 3891 | 3961146 | 1248 | 76.23 | 1018 | 828 | 14205 | 54 |
| G21002_Sphaenelater_lineicollis_Elateridae | 3755 | 3166326 | 1008 | 62.57 | 843.2 | 732 | 6012 | 60 |
| G21003_Analastesa_arabica_Elateridae | 2718 | 1471502 | 642 | 38.38 | 541.4 | 460.5 | 5064 | 54 |
| G21004_Paulusiella_serraticornis_Elateridae | 3922 | 3867633 | 1194 | 73.78 | 986.1 | 810 | 16131 | 60 |
| G21006_Rhipiceridae_indet_Rhipiceridae | 3753 | 4204833 | 1344 | 82.6 | 1120.4 | 924 | 21300 | 66 |
| G21007_Cneoglossa_sp_Cneoglossidae | 3889 | 4362865 | 1380 | 84.19 | 1121.8 | 924 | 12096 | 51 |
| G21009_Psilocladus_sp_Lampyridae | 3785 | 3262915 | 1041 | 64.35 | 862.1 | 732 | 6129 | 48 |
| G21010_Amydetinae_indet_Lampyridae | 3924 | 3657687 | 1128 | 69.84 | 932.1 | 813 | 10281 | 60 |
| G21011_Ototretinae_indet_Lampyridae | 3904 | 3597888 | 1089 | 68.88 | 921.6 | 792 | 9786 | 84 |
| G21012_Austroselasia_sp_Elateridae | 3889 | 3421100 | 1068 | 66.5 | 879.7 | 735 | 10260 | 78 |
| G21013_Habashaselasia_nekemtensis_Elateridae | 3875 | 3332547 | 1029 | 64.32 | 860 | 726 | 8679 | 84 |
| G21014_Elatichrosis_sp_Elateridae | 3828 | 3278334 | 1014 | 63.79 | 856.4 | 708 | 11220 | 66 |
| G21015_Brachypterodrilus_sp_Lampyridae | 3747 | 3039266 | 972 | 59.96 | 811.1 | 690 | 9015 | 27 |
| G21016_Rhagophthalmidae_indet_Rhagophthalmidae | 3889 | 3663222 | 1128 | 70.66 | 941.9 | 795 | 10278 | 78 |
| G21017_Lissominae_indet_Elateridae | 3832 | 4430280 | 1392 | 85.83 | 1156.1 | 951 | 17673 | 93 |
| G21019_Eucnemidae_indet_Eucnemidae | 2938 | 1899533 | 804 | 44.54 | 646.5 | 534 | 8145 | 48 |
| G21023_Ceratogonys_spinicorne_Eucnemidae | 3741 | 3414631 | 1113 | 67.37 | 912.8 | 765 | 7785 | 48 |
| G21026_Pyrophorini_indet_Elateridae | 3786 | 3339008 | 1062 | 65.4 | 881.9 | 750 | 10275 | 33 |
| G21027_Elaterinae_indet_Elateridae | 2518 | 1352203 | 633 | 37.39 | 537 | 450 | 4968 | 51 |
| G21028_Dascillidae_indet_Dascillidae | 3884 | 4483998 | 1386 | 86.42 | 1154.5 | 951 | 21351 | 72 |
| G21029_Drilaster_sp_Lampyridae | 3392 | 2247763 | 798 | 47.93 | 662.7 | 561 | 5718 | 45 |
| G21030_Psephenidae_indet_Psephenidae | 3879 | 4114527 | 1281 | 79.02 | 1060.7 | 891 | 20358 | 81 |
| G21031_Rousia_sp_Elateridae | 3926 | 4460042 | 1356 | 85.07 | 1136 | 948 | 19278 | 75 |
| G21035_Anaissus_sp_Elateridae | 3890 | 3793719 | 1194 | 73.25 | 975.2 | 825 | 8874 | 72 |
| G21036_Campyloxenus_pyrothorax_Elateridae | 3779 | 3311373 | 1050 | 66.09 | 876.3 | 738 | 21357 | 33 |
| G21037_Pityobius_anguinus_Elateridae | 3419 | 2624742 | 930 | 56.43 | 767.7 | 648 | 4767 | 54 |
| G21038_Adelocera_sp_Elateridae | 3735 | 3134689 | 1002 | 62.03 | 839.3 | 699 | 7248 | 57 |
| G21039_Psephenidae_indet_Psephenidae | 3917 | 4378515 | 1356 | 83.88 | 1117.8 | 927 | 14139 | 60 |
| G21040_Adelocera_sp_Elateridae | 3758 | 2885868 | 915 | 57.12 | 767.9 | 654 | 6999 | 60 |
| G21041_Adelocera_sp_Elateridae | 3969 | 4477348 | 1350 | 84.71 | 1128.1 | 936 | 15291 | 27 |
| G21042_Telegeusis_sp_Omethidae | 3756 | 3379026 | 1101 | 66.93 | 899.6 | 741 | 11211 | 51 |
| G22002_Plectonotum_laterale_Cantharidae | 3772 | 3285105 | 1038 | 65.02 | 870.9 | 711 | 10272 | 69 |
| G22003_Cerophytum_japonicum_Cerophytiae | 3157 | 1791553 | 666 | 43.4 | 567.5 | 477 | 4440 | 45 |
| G22005_Nyctophyxis_ocellatus_Elateridae | 2536 | 2483994 | 1239 | 69.03 | 979.5 | 834 | 10260 | 39 |
| G22007_Anaspasis_paralella_Elateridae | 2667 | 1437558 | 630 | 38.52 | 539 | 465 | 5142 | 63 |
| G22009_Jurasai_digitusdei_Jurasaidae | 3593 | 3122028 | 1029 | 64.24 | 868.9 | 729 | 11103 | 45 |
| G22015_Protelmidae_indet_Protelmidae | 3816 | 3586388 | 1161 | 70.05 | 939.8 | 771 | 7542 | 48 |
| G22020_Lutrochidae_indet_Lutrochidae | 3883 | 4049319 | 1287 | 78.15 | 1042.8 | 855 | 15969 | 45 |
| G22022_Lutrochidae_indet_Lutrochidae | 3923 | 4456557 | 1377 | 84.97 | 1136 | 942 | 19272 | 69 |
| G22024_Elmidae_indet_Elmidae | 3376 | 3679680 | 1329 | 79.75 | 1090 | 894 | 20361 | 81 |
| G22025_Ototretinae_indet_Lampyridae | 3978 | 3970209 | 1179 | 75.06 | 998 | 864 | 19257 | 48 |
| G22026_Stenocladius_sp_Lampyridae | 3492 | 2465174 | 855 | 52.13 | 705.9 | 600 | 6432 | 57 |
| G22027_Lampyrinae_indet_Lampyridae | 3829 | 3958098 | 1233 | 76.49 | 1033.7 | 843 | 16143 | 60 |
| G19002_Hemiops_sp_Elateridae | 3615 | 2537544 | 849 | 52.19 | 701.9 | 588 | 7071 | 72 |
| Heterocerus_fenestratus_Heteroceridae | 3082 | 1985923 | 756 | 52.3 | 644.4 | 594 | 2331 | 36 |
| Heteromastix_sp_Cantharidae | 3876 | 4440562 | 1389 | 86.71 | 1145.7 | 954 | 8517 | 48 |
| Hypnoidus_riparius_Elateridae | 3849 | 4506433 | 1410 | 88.31 | 1170.8 | 981 | 12078 | 90 |
| G19015_Iberobaenia_minuta_Iberobaeniidae | 3601 | 2584692 | 852 | 53.16 | 717.8 | 606 | 9222 | 63 |
| Ignelater_luminosus_Elateridae | 3834 | 4478535 | 1416 | 87.23 | 1168.1 | 966 | 16131 | 60 |
| R19001_Isorhipis_marmottani_Eucnemidae | 3876 | 4436799 | 1386 | 86.7 | 1144.7 | 960 | 16422 | 90 |
| Konoplatycis_otome_Lycidae | 3381 | 2867979 | 1032 | 65.7 | 848.3 | 732 | 18951 | 66 |
| Lamprigera_yunnana_Lampyridae | 3725 | 4416438 | 1437 | 88.21 | 1185.6 | 972 | 16146 | 48 |
| Lamprohiza_splendidula_Lampyridae | 3623 | 3622619 | 1203 | 75.53 | 999.9 | 855 | 10332 | 78 |
| S1_Lampyris_sardiniae_Lampyridae | 3744 | 4346898 | 1410 | 87.36 | 1161 | 966 | 19794 | 33 |
| Leptinotarsa_decemlineata | 4007 | 4217655 | 1251 | 77.89 | 1052.6 | 882 | 13518 | 99 |
| Leptolycus_sp_Lycidae | 3678 | 2517450 | 825 | 50.45 | 684.5 | 570 | 9786 | 66 |
| Libnetis_sp_Lycidae | 3686 | 4065385 | 1335 | 83.25 | 1102.9 | 933 | 9744 | 84 |
| Limonius_californicus_Elateridae | 3884 | 3469095 | 1032 | 67.04 | 893.2 | 750 | 19278 | 57 |
| R19013_Lissomus_sp_Elateridae | 3815 | 4412208 | 1392 | 87.64 | 1156.5 | 975 | 10494 | 111 |
| Lucidota_atra_Lampyridae | 3730 | 4174680 | 1359 | 84.05 | 1119.2 | 939 | 16143 | 75 |
| Lycoprogenthes_sp_Lycidae | 3664 | 4162624 | 1377 | 85.7 | 1136.1 | 948 | 10935 | 93 |
| Lycostomus_kraatzi_Lycidae | 3430 | 3738999 | 1320 | 83.06 | 1090.1 | 918 | 11460 | 102 |
| Lygistopterus_sanguineus_Lycidae | 3662 | 4124955 | 1350 | 84.68 | 1126.4 | 942 | 16077 | 84 |
| Lyropaeus_optabilis_Lycidae | 3472 | 3717908 | 1305 | 80.22 | 1070.8 | 902.5 | 16779 | 57 |
| Ponyalis_sp_Lycidae | 3625 | 4073273 | 1362 | 84.68 | 1123.7 | 945 | 13185 | 45 |
| R20009_Malthodes_sp_Cantharidae | 3812 | 4188375 | 1317 | 83.2 | 1098.7 | 948 | 7743 | 81 |
| Melanotus_cribricollis_Elateridae | 3663 | 4333035 | 1431 | 88.7 | 1182.9 | 975 | 16149 | 96 |
| Melanotus_villosus_Elateridae | 3666 | 4128162 | 1356 | 85.23 | 1126.1 | 961.5 | 12525 | 90 |
| Micronaspis_floridana_Lampyridae | 3471 | 3346437 | 1188 | 73.52 | 964.1 | 822 | 7674 | 48 |
| Microphotus_sp_Lampyridae | 3534 | 3756876 | 1296 | 79.89 | 1063.1 | 891 | 13296 | 78 |
| G19011_Diplophoenicus_sp_Elateridae | 3921 | 4270214 | 1314 | 81.9 | 1089.1 | 924 | 16146 | 69 |
| Negastrius_arenicola_Elateridae | 3938 | 4586445 | 1416 | 88.52 | 1164.7 | 972 | 12876 | 48 |
| Notolioon_sp_Byrrhidae | 3157 | 3100175 | 1218 | 76.44 | 982 | 843 | 6669 | 72 |
| Omalisus_fontisbellaquei_Elateridae | 3858 | 3722241 | 1149 | 72.1 | 964.8 | 825 | 21357 | 60 |
| Onthophagus_taurus | 4079 | 4968361 | 1443 | 90.68 | 1218 | 1002 | 21006 | 129 |
| R19005_Ototretinae_indet_Lampyridae | 3823 | 4449279 | 1416 | 87.87 | 1163.8 | 963 | 20370 | 96 |
| R20013_Ototretinae_indet_Lampyridae | 3875 | 4415193 | 1380 | 86.13 | 1139.4 | 960 | 8661 | 33 |
| G19008_Oxynopterus_sp_Elateridae | 3919 | 4281941 | 1332 | 82.07 | 1092.6 | 915 | 13353 | 81 |
| G19005_Pachyderes_sp_Elateridae | 3901 | 3713631 | 1143 | 71.34 | 952 | 819 | 13557 | 66 |
| G19006_Parasaphes_sp_Elateridae | 3907 | 4176045 | 1290 | 80.12 | 1068.9 | 897 | 15375 | 66 |
| G19004_Pectocera_sp_Elateridae | 3915 | 3683818 | 1131 | 71.08 | 940.9 | 804 | 10245 | 54 |
| Phausis_reticulata_Lampyridae | 3688 | 4084599 | 1341 | 83.69 | 1107.5 | 939 | 13626 | 33 |
| Phengodes_sp_B_Phengodidae | 3635 | 4019172 | 1341 | 83.29 | 1105.7 | 924 | 18975 | 84 |
| Phengodes_sp_A_Phengodidae | 3858 | 3569784 | 1116 | 68.92 | 925.3 | 780 | 11121 | 84 |
| Photinus_australis_Lampyridae | 3533 | 3651195 | 1275 | 78.72 | 1033.5 | 891 | 10407 | 48 |
| Photinus_carolinus_Lampyridae | 3566 | 3709743 | 1284 | 78.61 | 1040.3 | 882 | 13293 | 45 |
| Photinus_macdermotti_Lampyridae | 3396 | 3667875 | 1344 | 81.52 | 1080.1 | 894 | 15828 | 42 |
| Photinus_pyralis_Lampyridae | 3874 | 4548807 | 1413 | 88.47 | 1174.2 | 969 | 21351 | 54 |
| Photinus_scintillans_Lampyridae | 3775 | 4320327 | 1386 | 86.53 | 1144.5 | 951 | 13296 | 78 |
| Photuris_frontalis_Lampyridae | 3698 | 4182123 | 1383 | 85.49 | 1130.9 | 945 | 19275 | 66 |
| Photuris_sp_Lampyridae | 3531 | 3856203 | 1353 | 82.08 | 1092.1 | 918 | 16146 | 51 |
| Phrixothrix_hirtus_Phengodidae | 2654 | 2030806 | 975 | 62.96 | 765.2 | 618 | 12615 | 51 |
| A01544_Plastocerus_angulosus_Elateridae | 3918 | 4083348 | 1269 | 78.34 | 1042.2 | 867 | 21312 | 33 |
| Platerodrilus_sp_Lycidae | 3218 | 3336606 | 1263 | 78.69 | 1036.9 | 873 | 16410 | 66 |
| Plateros_sp_Lycidae | 3633 | 4182954 | 1392 | 86.31 | 1151.4 | 960 | 10176 | 60 |
| Ponyalis_quadricollis_Lycidae | 3527 | 3826175 | 1314 | 82.25 | 1084.8 | 921 | 9864 | 87 |
| R19004_Ptilodactylidae_indet_Ptilodactylidae | 3951 | 4635972 | 1431 | 89.03 | 1173.4 | 987 | 18279 | 81 |
| Ptosima_undecimmaculata_Buprestidae | 3331 | 3302208 | 1212 | 76.23 | 991.4 | 843 | 16680 | 63 |
| Pyractomena_borealis_Lampyridae | 3727 | 4224120 | 1386 | 85.25 | 1133.4 | 948 | 19272 | 69 |
| Pyrearinus_fragilis_Elateridae | 3857 | 4625235 | 1443 | 90.34 | 1199.2 | 996 | 19623 | 93 |
| Pyrocoelia_pectoralis_Lampyridae | 3735 | 4044383 | 1335 | 81.79 | 1082.8 | 903 | 19593 | 63 |
| R18040_Mangkutanus_sp_Lycidae | 3726 | 4182852 | 1356 | 84.94 | 1122.6 | 933 | 20361 | 48 |
| R20002_Agrypninae_indet_Elateridae | 3741 | 4132254 | 1338 | 83.53 | 1104.6 | 948 | 6807 | 60 |
| R20003_Athous_sp_Elateridae | 3910 | 4505430 | 1386 | 86.9 | 1152.3 | 960 | 10716 | 87 |
| R20004_Selatosomus_aeneus_Elateridae | 3611 | 3768429 | 1272 | 80.84 | 1043.6 | 918 | 6306 | 72 |
| R20005_Ampedus_sineatus_Elateridae | 3841 | 4496166 | 1416 | 88.47 | 1170.6 | 984 | 11157 | 57 |
| R20006_Ampedus_cinnabarinu_Elateridaes | 3873 | 4530978 | 1413 | 88.28 | 1169.9 | 981 | 8532 | 78 |
| R20007_Anostirus_purpureus_Elateridae | 3842 | 4446189 | 1401 | 87.4 | 1157.3 | 969 | 19746 | 81 |
| R20008_Agriotes_sp_Elateridae | 3877 | 4526148 | 1416 | 87.96 | 1167.4 | 984 | 16146 | 93 |
| R20010_Elaterinae_indet_Elateridae | 3861 | 4389435 | 1380 | 85.42 | 1136.9 | 957 | 9579 | 60 |
| R20011_Alaus_myops_Elateridae | 3220 | 2907984 | 1089 | 71.31 | 903.1 | 777 | 15180 | 66 |
| R20014_Limnius_perrisi_Elmidae | 3914 | 4555800 | 1401 | 88.24 | 1164 | 973.5 | 7200 | 51 |
| R20016_Cardiophorus_discicollis_Elateridae | 3906 | 4496634 | 1389 | 86.76 | 1151.2 | 957 | 9075 | 60 |
| R20017_Eubria_palustris_Psephenidae | 3911 | 4563714 | 1422 | 88.23 | 1166.9 | 966 | 10572 | 96 |
| R20018_Phosphaenus_hemipterus_Lampyridae | 3861 | 4448274 | 1392 | 87.06 | 1152.1 | 957 | 10515 | 90 |
| R20019_Dascillus_cervinus_Dascillidae | 3930 | 4495335 | 1386 | 86.6 | 1143.9 | 958.5 | 8064 | 72 |
| R20022_Lampyris_noctiluca_Lampyridae | 3891 | 4466439 | 1386 | 86.61 | 1147.9 | 969 | 8919 | 48 |
| Anischia_sp_Eucnemidae | 3858 | 4054758 | 1281 | 78.39 | 1051 | 861 | 16158 | 72 |
| Platycrepidius_sp_Elateridae | 3961 | 4434126 | 1362 | 83.96 | 1119.4 | 930 | 21357 | 27 |
| Lamellipalpus_pacholatkoi_Lampyridae | 3973 | 3962575 | 1173 | 74.97 | 997.4 | 858 | 10254 | 81 |
| Byrrhinus_sp_Limnichidae | 3913 | 4586040 | 1422 | 88.22 | 1172 | 966 | 15843 | 102 |
| Epilichas_sp_Ptilodactylidae | 3414 | 2257063 | 795 | 48.97 | 661.1 | 534 | 6282 | 48 |
| Rhagonycha_fulva_Cantharidae | 3455 | 3691702 | 1293 | 80.67 | 1068.5 | 930 | 9933 | 96 |
| Rhagopthalmus_sp_Rhagophthalmidae | 3740 | 4352655 | 1401 | 88.01 | 1163.8 | 966 | 21354 | 93 |
| S39_Limnichidae_indet_Limnichidae | 3666 | 3999504 | 1347 | 82.5 | 1091 | 918 | 14982 | 78 |
| Sandalus_niger_Rhipiceridae | 3549 | 3828730 | 1305 | 81.24 | 1078.8 | 912 | 20379 | 48 |
| Scarelus_anthracinus_Lycidae | 3711 | 4182281 | 1365 | 84.9 | 1127 | 942 | 19323 | 60 |
| G19009_Semiotus_sp_Elateridae | 3956 | 4223981 | 1293 | 80.06 | 1067.7 | 879 | 16146 | 96 |
| R19009_Silis_sp_Cantharidae | 3885 | 4550625 | 1407 | 88.47 | 1171.3 | 972 | 16146 | 66 |
| Sinopyrophorus_schimmeli_Sinopyrophoridae | 3788 | 3431362 | 1110 | 68.51 | 905.9 | 748.5 | 6696 | 54 |
| G20013_Sinopyrophorus_sp_Sinopyrophoridae | 3765 | 3246969 | 1059 | 64.74 | 862.4 | 705 | 9570 | 42 |
| Taphes_brevicollis_Lycidae | 3684 | 4182843 | 1386 | 85.24 | 1135.4 | 951 | 16857 | 57 |
| G19001_Sinelater_perroti_Elateridae | 3828 | 3584212 | 1131 | 70.51 | 936.3 | 771 | 13527 | 45 |
| Tribolium_castaneum | 4163 | 5390981 | 1542 | 97.02 | 1295 | 1056 | 21357 | 156 |
| Trixagus_carinifrons_Throscidae | 3409 | 2978667 | 1053 | 68.17 | 873.8 | 810 | 3297 | 36 |

**Supplementary Table S5.** Overview of compiled datasets, statistics and descriptions.

| Dataset name | Type | Genes number | Num. of aligned positions | Num. of taxa | Completenes (C) score | C_max | C_min | Parsimony informative sites | MARE matrix saturation | MARE information content | Bowker's test | Dataset's description |
|---|---|---|---|---|---|---|---|---|---|---|---|---|
| **A-4189-AA** | AA | 4189 | 1 865 514 | 200 | 0.661537 | 0.963269 | 0.241612 | 1 133 846 | 0.881 | 0.39 | 99.75% | AliScore masked E-4224-AA-raw dataset. After masking in each individual gene alignment, sequences with ≥90% missing data were removed and individual gene alignments on both amino acid and nucleotide levels with less than 25% (50) taxa were removed. |
| **B-4189-NT** | NT | 4189 | 5 596 542 | 200 | 0.661539 | 0.96327 | 0.241614 | 3 745 031 | NA | NA | 99.99% | AliScore masked F-4224-NT-raw dataset. After masking in each individual gene alignment, sequences with ≥90% missing data were removed and individual gene alignments on both amino acid and nucleotide levels with less than 25% (50) taxa were removed. |
| **C-4189-NT12** | NT | 4189 | 3 731 028 | 200 | | | | 1 936 237 | NA | NA | 99.96% | Dataset B-4189-NT after removal of third codon position. |
| **D-1892-AA-MARE** | AA | 1892 | 718 512 | 196 | 0.765498 | 0.965888 | 0.329569 | 364 851 | 0.935 | 0.548 | 97.17% | Dataset A-4189-AA after filtering with MARE using default settings. |
| **E-4224-AA-raw** | AA | 4224 | 3 383 300 | 200 | 0.419818 | 0.72923 | 0.156032 | 1 645 101 | 0.883 | 0.364 | 99.87% | Raw data after removal of outlier sequences. |
| **F-4224-NT-raw** | NT | 4224 | 10 149 900 | 200 | 0.419819 | 0.729231 | 0.156033 | 5 375 430 | NA | NA | 99.99% | Raw data after removal of outlier sequences. |
| **G-962-AA-decisive** | AA | 962 | 416 210 | 52 | 0.837545 | 0.974909 | 0.463213 | 201 753 | | | 99.70% | Decisive dataset of 52 selected taxa. Only genes with all taxa kept. |
| **H-962-NT-decisive** | NT | 962 | 1 248 630 | 52 | 0.837547 | 0.974909 | 0.463213 | 737 380 | NA | NA | 100% | Decisive dataset of 52 selected taxa. Only genes with all taxa kept. |
| **I-1451-AA-Bacoca** | AA | 1451 | 817 164 | 200 | 0.652969 | 0.969666 | 0.232087 | 426 540 | 0.833 | 0.444 | 96.56% | Dataset A-4189-AA after removing partitions with an overall RCFV value >=0.1. |
| **J-1451-NT-Bacoca** | NT | 1451 | 2 451 492 | 200 | 0.652971 | 0.969666 | 0.232086 | 1 509 974 | NA | NA | 99.95% | Dataset B-4189-NT after removing partitions with an overall RCFV value >=0.1 (identified at amino acid level) to enable direct comparison. |
| **K-3583-NT-SymTest** | NT | 3583 | 5 045 097 | 200 | 0.651593 | 0.963255 | 0.232604 | 3 382 295 | NA | NA | 99.98% | Genes of dataset B-4189-NT passing IQTree matched-pair test of symetry (pvalue cutoff = 0.05). |
| **L-200-fast-AA-decisive** | AA | 200 | 61 329 | 52 | 0.901704 | 0.973911 | 0.610869 | 42 397 | - | - | - | Subset of dataset G-962-AA-decisive. |
| **M-200-slow-AA-decisive** | AA | 200 | 125 526 | 52 | 0.747399 | 0.969664 | 0.350278 | 51 144 | - | - | - | Subset of dataset G-962-AA-decisive. |

**Supplementary Table S6.** Overview of selected fossil taxa used for node calibrations, with their corresponding soft minimum and soft maximum age constraints. Detailed fossil placement and justifications are provided in Supplementary Figure S33 and Supplementary Text S2.

| Node | Clade | Fossil constraint | Minimum age (Mya) | Maximum age (Mya) | Reference |
|---|---|---|---|---|---|
| 1 | crown Polyphaga | undescribed crown Polyphaga | 237 | 293.6 | Zheng *et al.,* 2018; Montagna *et al.,* 2024 |
| 2 | Rhipiceridae – Dascillidae | *Parelateriformius communis* | 165 | 235 | Yan & Wang 2010; Li *et al.,* 2022 |
| 3 | crown Dascillidae | *Cretodascillus sinensis* | 122.5 | 235 | Jin *et al.,* 2013 |
| 4 | (Dryopidae + Protelmidae) Lutrochidae)) | *Mastigocoleus rhinoceros* | 112.6 | 165 | Tihelka *et al.,* 2022 |
| 5 | Ptilodactylidae – Cneoglossidae | *Aphebodactyla rhetine* | 98.17 | 165 | Chatzimanolis *et al.,* 2012 |
| 6 | crown Elmidae | *Cretohypsilara parva* | 98.17 | 165 | Cai *et al.,* 2018 |
| 7 | Limnichidae – Heteroceridae | *Heterocerites magnus* | 122.2 | 165 | Prokin & Ren 2011 |
| 8 | crown Psephenidae | *Vetujinbrianax cretaceus* | 98.17 | 165 | Cai & Huang 2018 |
| 9 | Artematopodidae – Omethidae | *Sinobrevipogon jurassicus* | 165 | 235 | Cai *et al.,* 2015 |
| 10 | (Cerophytidae + Jurrasaidae) Throscidae)) | *Praelaterium problematicum* | 190.8 | 235 | Yu *et al.,* 2019 |
| 11 | Anischinae – “core” Eucnemidae | *Rheanischia brevicornis* | 122.2 | 235 | Li *et al.,* 2021 |
| 12 | Lycidae – Iberobaeniidae or crown Lycidae | *Burmolycus compactus* | 98.17 | 165 | Bocak *et al.,* 2019 |
| 13 | crown Cantharidae | *Molliberus albae* | 105 | 165 | Peris & Fanti 2018 |
| 14 | Phengodidae – Rhagophthalmidae | *Cretocydistus wittmeri* | 98.17 | 165 | Roza *et al.,* 2023 |

**Supplementary Table S7.** Estimated marginal likelihood of relaxed-clock models with MCMCTree and mcmc3r.

**Method 1:** Datasets A-4189-AA and B-4189-NT reduced to 90% completeness, 200 taxa, resulting in 174,330 amino acid and 523,013 nucleotide positions.

| Data | Model | –log L | S.E. (delta) | Pr(M\|D) |
|---|---|---|---|---|
| **Amino acids** | CLK | -68883.88 | 1.574319 | 0 |
| | ILN | -1632.394 | 2.376573 | 0.00388202 |
| | GBM | -1626.846 | 2.688745 | 0.99611798 |
| **Nucleotides** | **Model** | **–log L** | **S.E. (delta)** | **Pr(M\|D)** |
| | CLK | -269716.3 | 2.712618 | 0 |
| | ILN | -1933.031 | 2.903188 | 0 |
| | GBM | -1630.658 | 2.660464 | 1 |

**Method 2:** Randomly selected 300 genes (selecting from 1914 genes with 195 and more taxa apresent in datasets A-4189-AA and B-4189-NT), resulting in 131,160 amino acid and 393,480 nucleotide positions.

| Data | Model | –log L | S.E. (delta) | Pr(M\|D) |
|---|---|---|---|---|
| **Amino acids** | CLK | -58041.42 | 1.431978 | 0 |
| | ILN | -1570.911 | 2.028879 | 0.02080547 |
| | GBM | -1567.06 | 1.964049 | 0.97919453 |
| **Nucleotides** | **Model** | **–log L** | **S.E. (delta)** | **Pr(M\|D)** |
| | CLK | -173239.5 | 2.614653 | 0 |
| | ILN | -1827.042 | 2.559182 | 1 |
| | GBM | -1872.01 | 2.931212 | 2.95417e-20 |

| | |
|---|---|
| CLK | The geometric Brownian motion (GBM) model 3 auto corelated rates |
| ILN | The independent log-normal (ILN) model 2 independent rates |
| CLK | The strict clock (CLK) model |

**Supplementary Table S8.** Divergence time estimates (Mya) for major Elateriformia lineages and comparison between this study and previous studies.

| **This study** | | | **Age (Mya)** | |
|---|---|---|---|---|
| **Node** | **Analysis** | **Mean** | **Upper CI** | **Lower CI** |
| crown Elateriformia | Dataset_A_NT_AR | 287.13 | 296.05 | 275.97 |
| crown Elateriformia | Dataset_A_AA_AR | 281.59 | 294.22 | 266.73 |
| crown Elateriformia | Dataset_A_NT_IR | 284.91 | 295.38 | 272.19 |
| crown Elateriformia | Dataset_A_AA_IR | 280.99 | 294.07 | 266.04 |
| crown Elateriformia | Dataset_B_NT_AR | 286.55 | 296.24 | 275.48 |
| crown Elateriformia | Dataset_B_AA_AR | 282.39 | 294.53 | 267.93 |
| crown Elateriformia | Dataset_B_NT_IR | 285.13 | 295.61 | 272.66 |
| crown Elateriformia | Dataset_B_AA_IR | 283.22 | 294.61 | 269.01 |
| Byrrhoidea+Elateroidea | Dataset_A_NT_AR | 257.79 | 266.88 | 248.35 |
| Byrrhoidea+Elateroidea | Dataset_A_AA_AR | 252.34 | 264.51 | 240.28 |
| Byrrhoidea+Elateroidea | Dataset_A_NT_IR | 261.75 | 274.14 | 249.15 |
| Byrrhoidea+Elateroidea | Dataset_A_AA_IR | 255.69 | 269.7 | 241.31 |
| Byrrhoidea+Elateroidea | Dataset_B_NT_AR | 256.91 | 266.04 | 247.02 |
| Byrrhoidea+Elateroidea | Dataset_B_AA_AR | 253.18 | 264.52 | 240.81 |
| Byrrhoidea+Elateroidea | Dataset_B_NT_IR | 261.33 | 274.03 | 248.5 |
| Byrrhoidea+Elateroidea | Dataset_B_AA_IR | 258.09 | 272.02 | 244.35 |
| crown Dascilloidea | Dataset_A_NT_AR | 216.12 | 238.57 | 189.24 |
| crown Dascilloidea | Dataset_A_AA_AR | 211.82 | 236.01 | 182.77 |
| crown Dascilloidea | Dataset_A_NT_IR | 212.16 | 236.74 | 179.77 |
| crown Dascilloidea | Dataset_A_AA_IR | 199.95 | 233.26 | 168.51 |
| crown Dascilloidea | Dataset_B_NT_AR | 215.15 | 237.94 | 185.49 |
| crown Dascilloidea | Dataset_B_AA_AR | 213.56 | 236.05 | 181.97 |
| crown Dascilloidea | Dataset_B_NT_IR | 208.88 | 236.62 | 175.71 |
| crown Dascilloidea | Dataset_B_AA_IR | 200.68 | 234.75 | 169.22 |
| crown Byrrhoidea | Dataset_A_NT_AR | 250.93 | 260.04 | 241.38 |
| crown Byrrhoidea | Dataset_A_AA_AR | 243.72 | 255.32 | 231.71 |
| crown Byrrhoidea | Dataset_A_NT_IR | 251.1 | 265.75 | 235.7 |
| crown Byrrhoidea | Dataset_A_AA_IR | 243.75 | 260.09 | 227.26 |
| crown Byrrhoidea | Dataset_B_NT_AR | 250.19 | 259.51 | 240.45 |
| crown Byrrhoidea | Dataset_B_AA_AR | 245.16 | 256.65 | 233.35 |
| crown Byrrhoidea | Dataset_B_NT_IR | 250.95 | 265.4 | 235.15 |
| crown Byrrhoidea | Dataset_B_AA_IR | 246.4 | 262.61 | 230.1 |
| crown Elateroidea | Dataset_A_NT_AR | 250.3 | 259.06 | 241.01 |
| crown Elateroidea | Dataset_A_AA_AR | 242.07 | 253.34 | 230.28 |
| crown Elateroidea | Dataset_A_NT_IR | 254.45 | 266.39 | 241.4 |
| crown Elateroidea | Dataset_A_AA_IR | 246.23 | 260.18 | 232.12 |
| crown Elateroidea | Dataset_B_NT_AR | 249.31 | 258.4 | 239.93 |
| crown Elateroidea | Dataset_B_AA_AR | 242.32 | 253.62 | 231.06 |
| crown Elateroidea | Dataset_B_NT_IR | 253.79 | 266.5 | 241.14 |
| crown Elateroidea | Dataset_B_AA_IR | 247.98 | 261.91 | 234.26 |
| Buprestoidea(Byrrhoidea+Elateroidea) | Dataset_A_NT_AR | 270.8 | 279.77 | 260.72 |
| Buprestoidea(Byrrhoidea+Elateroidea) | Dataset_A_AA_AR | 263.65 | 276.02 | 250.53 |
| Buprestoidea(Byrrhoidea+Elateroidea) | Dataset_A_NT_IR | 272.94 | 284.95 | 260.28 |
| Buprestoidea(Byrrhoidea+Elateroidea) | Dataset_A_AA_IR | 266.35 | 280.61 | 251.73 |
| Buprestoidea(Byrrhoidea+Elateroidea) | Dataset_B_NT_AR | 269.04 | 278.32 | 258.79 |
| Buprestoidea(Byrrhoidea+Elateroidea) | Dataset_B_AA_AR | 264.15 | 275.8 | 251.03 |
| Buprestoidea(Byrrhoidea+Elateroidea) | Dataset_B_NT_IR | 272.25 | 284.39 | 259.38 |
| Buprestoidea(Byrrhoidea+Elateroidea) | Dataset_B_AA_IR | 268.63 | 282.51 | 254.87 |

| **Previous studies** | | **Age (Mya)** | | | | |
|---|---|---|---|---|---|---|
| **Node** | **Study** | **Mean** | **Upper CI** | **Lower CI** | **Median** | **Note** |
| crown Elateriformia | Toussaint_et_al_2016 | NA | 281.49 | 252.78 | 266.56 | |
| crown Elateriformia | McKenna_et_al_2015 | NA | NA | NA | 184.6 | 95% CI numbers not published in original study |
| crown Elateriformia | McKenna_et_al_2019_1 | 242.81 | 267.77 | 204.55 | | 4818-gene dataset |
| crown Elateriformia | McKenna_et_al_2019_2 | 239.35 | 267.09 | 160.4 | | 89-gene dataset |
| crown Elateriformia | Zhang_et_al_2018_1 | 240.84 | 251.34 | 230.19 | | Family level tree calibrations |
| crown Elateriformia | Zhang_et_al_2018_2 | 217.90 | 227.51 | 207.97 | | |
| crown Elateriformia | Cai_et_al_2022 | 254.25 | 260.16 | 248.63 | | |
| crown Dascilloidea | Toussaint_et_al_2016 | NA | 212.39 | 149.25 | 179.92 | |
| crown Dascilloidea | McKenna_et_al_2015 | NA | 155.79 | 82.36 | 120.47 | Values from Table 2. in McKenna et al.,2015 |
| crown Dascilloidea | McKenna_et_al_2019_1 | 154.29 | 202.94 | 108.82 | | 4818-gene dataset |
| crown Dascilloidea | McKenna_et_al_2019_2 | 138.73 | 178.04 | 107.78 | | 89-gene dataset |
| crown Dascilloidea | Zhang_et_al_2018_1 | 132.31 | 154.11 | 112.11 | | Family level tree calibrations |
| crown Dascilloidea | Zhang_et_al_2018_2 | 119.07 | 138.83 | 100.17 | | |
| crown Dascilloidea | Cai_et_al_2022 | 189.37 | 213.43 | 164.2 | | |
| crown Byrrhoidea | Toussaint_et_al_2016 | NA | 254.67 | 222.21 | 238.65 | |
| crown Byrrhoidea | McKenna_et_al_2015 | NA | 176.96 | 142.85 | 160.03 | Values from Table 2. in McKenna et al.,2015 |
| crown Byrrhoidea | McKenna_et_al_2019_1 | NA | NA | NA | | Byrrhoidea paraphyletic |
| crown Byrrhoidea | McKenna_et_al_2019_2 | NA | NA | NA | | Byrrhoidea paraphyletic |
| crown Byrrhoidea | Zhang_et_al_2018_1 | 195.82 | 207.18 | 183.59 | | Family level tree calibrations |
| crown Byrrhoidea | Zhang_et_al_2018_2 | 176.99 | 187.44 | 167.05 | | |
| crown Byrrhoidea | Cai_et_al_2022 | NA | NA | NA | | Byrrhoidea paraphyletic |
| crown Elateroidea | Toussaint_et_al_2016 | NA | 260.12 | 231.35 | 246.02 | |
| crown Elateroidea | McKenna_et_al_2015 | NA | 181.57 | 151.83 | 166.18 | Values from Table 2. in McKenna et al.,2015 |
| crown Elateroidea | McKenna_et_al_2019_1 | 187.7 | 215.28 | 151.75 | | 4818-gene dataset |
| crown Elateroidea | McKenna_et_al_2019_2 | 195.36 | 221.07 | 131.35 | | 89-gene dataset |
| crown Elateroidea | Zhang_et_al_2018_1 | 203.28 | 213.22 | 194.17 | | Family level tree calibrations |
| crown Elateroidea | Zhang_et_al_2018_2 | 183.01 | 191.11 | 174.73 | | |
| crown Elateroidea | Cai_et_al_2022 | 224.84 | 232.86 | 216.69 | | |
| Buprestoidea(Byrrhoidea+Elateroidea) | Toussaint_et_al_2016 | NA | 273.9 | 245.3 | 260.16 | |
| Buprestoidea(Byrrhoidea+Elateroidea) | McKenna_et_al_2015 | NA | | | 178.91 | 95% CI numbers not published in original study |
| Buprestoidea(Byrrhoidea+Elateroidea) | McKenna_et_al_2019_1 | NA | NA | NA | | Byrrhoidea paraphyletic |
| Buprestoidea(Byrrhoidea+Elateroidea) | McKenna_et_al_2019_2 | NA | NA | NA | | Byrrhoidea paraphyletic |
| Buprestoidea(Byrrhoidea+Elateroidea) | Zhang_et_al_2018_1 | 222.56 | 232.76 | 212.87 | | Family level tree calibrations |
| Buprestoidea(Byrrhoidea+Elateroidea) | Zhang_et_al_2018_2 | 201.37 | 211.00 | 192.11 | | |
| Buprestoidea(Byrrhoidea+Elateroidea) | Cai_et_al_2022 | NA | NA | NA | | Byrrhoidea paraphyletic |
| Byrrhoidea+Elateroidea | Toussaint_et_al_2016 | NA | 269.38 | 241.24 | 255.87 | |
| Byrrhoidea+Elateroidea | McKenna_et_al_2015 | NA | | | 175.04 | 95% CI numbers not published in original study |
| Byrrhoidea+Elateroidea | McKenna_et_al_2019_1 | NA | NA | NA | | Byrrhoidea paraphyletic |
| Byrrhoidea+Elateroidea | McKenna_et_al_2019_2 | NA | NA | NA | | Byrrhoidea paraphyletic |
| Byrrhoidea+Elateroidea | Zhang_et_al_2018_1 | 211.95 | 221.94 | 202.59 | | |
| Byrrhoidea+Elateroidea | Zhang_et_al_2018_2 | 190.93 | 199.22 | 182.83 | | |
| Byrrhoidea+Elateroidea | Cai_et_al_2022 | NA | NA | NA | | Byrrhoidea paraphyletic |

**Supplementary Table S9.** Type and age (Mya) of neotenic and putatively neotenic lineages.

| | | | | | | | Median age (Ma) in the pressent study (95% CI), Dataset A nuceotides AR | | Median age (Ma) in the pressent study (95% CI), Dataset A nucleotides IR | | Median age (Ma) in the pressent study (95% CI), Dataset B nuceotides AR | | Median age (Ma) in the pressent study (95% CI), Dataset B nucleotides IR | | Median age (Ma) in the pressent study (95% CI), Dataset A amino acids, AR | | Median age (Ma) in the pressent study (95% CI), Dataset A amino acids, IR | | Median age (Ma) in the pressent study (95% CI), Dataset B amino acids, AR | | Median age (Ma) in the pressent study (95% CI), Dataset B amino acids, IR | |
|---|---|---|---|---|---|---|---|---|---|---|---|---|---|---|---|---|---|---|---|---|---|---|
| **Group** | **Neotenic lineage** | **Split from non-neotenic sister lineage** | **Crown split of neotenic lineage** | **Females** | **Type of female neoteny** | **Males** | **Split of neotenic clade from a sister clade.** | **Crown age of neotenic clade (if more neotenic species were sampled for lineage).** | **Split of neotenic clade from a sister clade.** | **Crown age of neotenic clade (if more neotenic species were sampled for lineage).** | **Split of neotenic clade from a sister clade.** | **Crown age of neotenic clade (if more neotenic species were sampled for lineage).** | **Split of neotenic clade from a sister clade.** | **Crown age of neotenic clade (if more neotenic species were sampled for lineage).** | **Split of neotenic clade from a sister clade.** | **Crown age of neotenic clade (if more neotenic species were sampled for lineage).** | **Split of neotenic clade from a sister clade.** | **Crown age of neotenic clade (if more neotenic species were sampled for lineage).** | **Split of neotenic clade from a sister clade.** | **Crown age of neotenic clade (if more neotenic species were sampled for lineage).** | **Split of neotenic clade from a sister clade.** | **Crown age of neotenic clade (if more neotenic species were sampled for lineage).** |
| 1 | Jurasaidae | Jurasaidae + Cerophytidae | NA | Females known for two species. | Type 4. Larviform females with adults-like-head and legs. | Winged | 156.97 (173.18 to 137.51) | NA | 137.08 (165.38 to 103.74) | NA | 157.78 (174.22 to 138.48) | NA | 138.87 (171.46 to 116.66) | NA | 153.58 (174.16 to 129.61) | NA | 143.87 (173.83 to 112.71) | NA | 158.19 (177.80 to 135.86) | NA | 146.18 (175.71 to 112.87) | NA |
| 2 | Telegeusinae | Telegeusinae + Driloniinae | NA | Females unknown. Only males are collected in high quantities. Females suspected to be neotenic. | - | Winged, shortened elytras | 167.95 (203.38 to 129.15) | NA | 153.48 (198.76 to 111.38) | NA | 166.77 (201.98 to 127.40) | NA | 151.66 (198.01 to 111.23) | NA | 158.83 (189.54 to 124.21) | NA | 145.88 (188.22 to 102.95) | NA | 158.74 (190.51 to 121.15) | NA | 144.28 (187.91 to 99.96) | NA |
| 3 | Iberobaeniidae | Iberobaeniidae + Lycidae | NA | Female unknown, suspected to be neotenic. Only males are collected. | - | Winged | 165.88 (177.77 to 155.31) | NA | 162.73 (171.93 to 151.55) | NA | 165.98 (177.51 to 155.19) | NA | 161.43 (171.66 to 148.49) | NA | 162.92 (171.18 to 153.49) | NA | 162.91 (171.21 to 153.16) | NA | 162.54 (170.71 to 152.93) | NA | 161.55 (170.06 to 150.33) | NA |
| 4 | *Dexoris ruzzieri* | Dexorinae + rest of Lycidae | NA | Female unknown, suspected to be neotenic. Only males are collected. | - | Winged | 122.86 (141.85 to 102.59) | NA | 132.00 (146.09 to 117.51) | NA | 120.81 (140.65 to 100.86) | NA | 125.67 (141.78 to 109.28) | NA | 119.46 (134.40 to 104.11) | NA | 132.49 (145.17 to 119.79) | NA | 117.04 (132.32 to 99.90) | NA | 127.44 (141.15 to 113.62) | NA |
| 5 | Lyropaeinae | Lyropaeinae + sister group | Atennolycus constrictus + rest of Lyropaeinae | Females known in some species. | Type 5. Fully larviform sexually mature females. | Winged | 87.78 (107.15 to 68.66) | 79.56 (98.38 to 61.62) | 97.21 (110.24 to 82.98) | 86.31 (101.57 to 70.84) | 86.76 (106.33 to 67.01) | 78.26 (97.52 to 60.21) | 91.33 (105.42 to 76.15) | 80.71 (96.81 to 64.50) | 81.76 (96.60 to 67.93) | 70.22 (84.05 to 57.80) | 94.14 (106.24 to 82.07) | 79.59 (93.56 to 65.50) | 81.53 (96.96 to 66.66) | 70.04 (84.37 to 56.75) | 90.61 (102.43 to 77.77) | 76.57 (91.03 to 62.56) |
| 6 | *Scarelus anthracinus* | *Scarelus* + rest of Ateliinae | NA | Female unknown. Only males are collected. | - | Winged | 57.83 (75.58 to 41.40) | NA | 51.50 (75.82 to 28.38) | NA | 57.05 (74.69 to 40.22) | NA | 48.36 (72.17 to 26.29) | NA | 50.62 (63.62 to 38.18) | NA | 50.03 (70.75 to 30.60) | NA | 51.20 (64.79 to 38.28) | NA | 48.08 (69.08 to 28.01) | NA |
| 7 | *Leptolycus* sp. | (*Leptolycus*(*Plateros* sp. + rest of Lycinae)) | NA | Female presumambly reported for one species. Only males are collected. | Type 5. Fully larviform sexually mature females. | Winged | 83.49 (103.05 to 65.38) | NA | 91.85 (105.95 to 76.88) | NA | 82.56 (102.56 to 64.03) | NA | 86.10 (101.66 to 70.38) | NA | 74.56 (88.57 to 61.16) | NA | 84.80 (98.28 to 71.16) | NA | 75.00 (89.50 to 60.51) | NA | 81.98 (95.94 to 68.50) | NA |
| 8 | Phengodidae + Rhagophthalmidae | (Lampyridae(Phengodidae + Rhagophthalmidae)) | Phengodidae + Rhagopthalmidae | Females known. | Type 5. Fully larviform sexually mature females. | Winged, most Phengodidae with shortened elytras | 184.01 (192.55 to 175.98) | 162.25 (170.75 to 152.57) | 180.05 (194.87 to 163.00) | 146.22 (166.36 to 124.89) | 182.42 (197.65 to 180.72) | 160.82 (168.98 to 150.09) | 174.67 (191.75 to 157.66) | 139.73 (163.44 to 116.12) | 166.47 (176.96 to 156.19) | 146.53 (159.17 to 133.53) | 164.26 (179.05 to 149.94) | 135.79 (155.13 to 115.45) | 164.02 (174.34 to 153.65) | 142.30 (155.48 to 129.21) | 161.11 (175.80 to 146.41) | 130.08 (150.83 to 109.84) |
| 9 | *Stenocladius* + *Brachypterodrilus* sp. | (*Stenocladius* + *Brachypterodrilus*) rest of Lampyridae)) | Brachypterodrilus sp. + Stenocladius | Females known in some species. | Type 5. Fully larviform sexually mature females. | Winged, Brachypterodrilus shortened elytras | 156.58 (166.65 to 145.67) | 144.35 (156.20 to 132.32) | 156.56 (170.51 to 141.04) | 136.26 (156.24 to 113.70) | 155.23 (165.76 to 143.49) | 143.14 (155.95 to 130.23) | 151.24 (166.74 to 134.59) | 131.38 (153.30 to 109.76) | 134.53 (147.01 to 121.92) | 123.09 (135.89 to 109.74) | 141.55 (156.14 to 127.30) | 122.50 (140.63 to 102.99) | 132.62 (144.99 to 119.74) | 120.61 (133.49 to 107.24) | 137.74 (152.81 to 123.40) | 117.94 (136.02 to 97.41) |
| 10 | *Lamprigera* sp. | *Lamprigera* + Luciolinae | NA | Females known. | Type 5. Fully larviform sexually mature females. | Winged | 113.60 (128.82 to 99.07) | NA | 106.83 (126.56 to 87.63) | NA | 111.13 (127.05 to 94.73) | NA | 101.97 (121.28 to 82.14) | NA | 92.93 (106.79 to 79.65) | NA | 93.61 (110.57 to 75.50) | NA | 89.61 (103.30 to 75.98) | NA | 89.25 (106.76 to 71.51) | NA |
| 11 | Lamprohrizinae | Lamprohrizinae + rest of Lampyridae | Lamprohriza splendidula + Phausis reticulata | Females known. | Type 3. Larviform females with adult-like head, prothorax, and legs; elytra absent, wingless or Type 2. Larviform females with adults-like-head, prothorax, and legs; elytra vestigial, wingless | Winged | 129.59 (140.97 to 117.87) | 69.91 (95.68 to 43.10) | 127.06 (140.52 to 113.18) | 50.57 (78.67 to 25.69) | 127.04 (139.01 to 113.87) | 68.68 (93.72 to 42.71) | 120.65 (135.88 to 105.91) | 48.07 (74.24 to 24.48) | 102.98 (115.22 to 90.53) | 61.91 (77.61 to 45.71) | 109.60 (123.09 to 96.79) | 54.26 (78.33 to 31.37) | 100.83 (113.00 to 88.43) | 60.25 (76.41 to 43.77) | 105.28 (118.83 to 91.99) | 50.62 (74.15 to 27.27) |
| 12 | *Microphotus, Pyrocoelia, Lampyris* | ((*Microphotus* + others) rest of Lampyrinae) | (Microphotus(Pyrocoelia + Lampyris)) | Females known. | Type 3. Larviform females with adult-like head, prothorax, and legs; elytra absent, wingless or Type 2. Larviform females with adults-like-head, prothorax, and legs; elytra vestigial, wingless | Winged | 72.29 (83.11 to 60.84) | 33.05 (44.03 to 22.22) | 79.66 (91.23 to 68.02) | 20.44 (28.21 to 13.15) | 71.13 (82.44 to 58.55) | 31.15 (42.53 to 20.69) | 75.16 (87.19 to 63.54) | 17.95 (24.98 to 11.70) | 55.42 (65.25 to 46.18) | 29.91 (37.64 to 22.18) | 69.16 (79.97 to 58.96) | 28.75 (38.44 to 19.47) | 53.01 (62.65 to 43.43) | 27.46 (35.44 to 20.07) | 65.53 (76.23 to 55.23) | 24.20 (32.85 to 16.04) |
| 13 | *Phosphaenus hemipterus* | (*Phosphaenus*(*Lucidota atra* + G22027)) | NA | Females known. | Type 3. Larviform females with adult-like head, prothorax, and legs; elytra absent, wingless | Males wingless with reduced elytras | 22.14 (28.81 to 15.67) | NA | 27.93 (37.29 to 18.81) | NA | 22.13 (29.18 to 15.52) | NA | 26.65 (35.99 to 18.05) | NA | 20.87 (26.13 to 15.93) | NA | 30.11 (39.17 to 21.58) | NA | 19.66 (25.00 to 14.71) | NA | 28.69 (37.71 to 20.30) | NA |
| 14 | *Photinus scintillans* | *Photinus scintillans* + *P. carolinus* | NA | Female known. | Type 2. Larviform females with adults-like-head, prothorax, and legs; elytra vestigial, wingless | Winged | 13.97 (18.52 to 9.54) | NA | 15.07 (21.33 to 9.18) | NA | 13.64 (18.34 to 9.40) | NA | 14.31 (20.18 to 8.81) | NA | 14.09 (17.98 to 10.52) | NA | 17.33 (23.92 to 11.16) | NA | 12.83 (16.66 to 9.35) | NA | 16.34 (22.71 to 10.43) | NA |
| 15 | *Cebrio* sp. | *Cebrio* sp. - rest | NA | Female known. | Type 2. Larviform females with adults-like-head, prothorax, and legs; elytra vestigial, wingless | Winged | 120.93 (136.22 to 106.34) | NA | 103.81 (121.21 to 84.88) | NA | 123.99 (138.40 to 109.07) | NA | 98.55 (116.25 to 79.41) | NA | 115.02 (128.57 to 101.26) | NA | 85.20 (102.40 to 69.14) | NA | 114.93 (129.38 to 100.33) | NA | 81.65 (98.13 to 65.93) | NA |
| 16 | Drilini | Drilini - rest of Agrypninae | Austroselasia sp. + Drilini | Females known. | Type 4. Larviform females with adults-like-head and legs. | Winged | 106.51 (120.36 to 89.71) | 27.42 (39.74 to 15.79) | 77.87 (95.23 to 62.13) | 29.79 (39.49 to 20.88) | 103.28 (118.64 to 84.85) | 25.16 (36.90 to 13.97) | 72.29 (88.90 to 57.63) | 27.61 (36.41 to 19.28) | 100.93 (115.02 to 85.73) | 28.93 (40.33 to 18.07) | 76.67 (89.12 to 64.00) | 38.08 (48.35 to 28.03) | 94.49 (110.90 to 77.90) | 24.13 (34.98 to 14.46) | 70.88 (83.28 to 59.06) | 34.24 (44.10 to 24.81) |
| 17 | Omalisinae | Omalisinae - rest of Elateridae | Omalisus + Euanoma | Female known in some species. | Type 2. Larviform females with adults-like-head, prothorax, and legs; elytra vestigial, wingless | Winged | 161.49 (173.11 to 150.73) | 110.31 (132.24 to 88.69) | 146.95 (161.72 to 132.41) | 100.72 (125.12 to 73.47) | 160.04 (170.71 to 148.85) | 109.52 (130.77 to 86.91) | 140.27 (156.18 to 123.94) | 95.93 (120.12 to 68.88) | 149.19 (160.41 to 137.58) | 101.07 (119.08 to 81.80) | 121.94 (137.91 to 107.16) | 83.12 (105.51 to 60.28) | 144.72 (157.04 to 132.38) | 97.61 (116.29 to 77.38) | 118.28 (133.25 to 103.57) | 80.61 (102.54 to 58.12) |
| 18 | *Octinodes* sp. | *Octinodes* sp. - *Ampedus* | NA | Females known in some species. | Type 2. Larviform females with adults-like-head, prothorax, and legs; elytra vestigial, wingless | Winged | 97.28 (111.45 to 82.37) | NA | 81.72 (100.24 to 62.61) | NA | 99.52 (113.86 to 85.02) | NA | 76.46 (95.44 to 57.85) | NA | 93.26 (105.93 to 79.77) | NA | 67.64 (83.32 to 52.58) | NA | 92.57 (106.25 to 78.72) | NA | 64.10 (79.23 to 49.77) | NA |
| 19 | *Paulusiella serraticornis* | *Paulusiella serraticornis* + *Hemiops* sp. | NA | Female unknown. Only males are collected in high quantities. | - | Winged | 37.35 (61.51 to 13.77) | NA | 27.87 (44.34 to 13.75) | NA | 38.80 (64.54 to 15.64) | NA | 26.12 (41.25 to 12.34) | NA | 63.06 (89.69 to 36.50) | NA | 37.21 (58.03 to 18.68) | NA | 58.39 (85.40 to 31.97) | NA | 34.05 (54.31 to 17.02) | NA |
| 20 | *Anorus parvicollis* | *Anorus parvicollis* + Dascillidae | NA | Female known. | Type 3. Larviform females with adult-like head, prothorax, and legs; elytra absent, wingless | Winged | 149.58 (180.75 to 119.60) | NA | 141.17 (174.87 to 117.43) | NA | 149.69 (183.48 to 120.30) | NA | 138.53 (171.46 to 116.66) | NA | 146.38 (176.74 to 119.28) | NA | 131.61 (157.27 to 113.75) | NA | 148.51 (179.16 to 119.79) | NA | 131.78 (158.67 to 113.00) | NA |

**Supplementary Table S10.** Divergence time estimates (Mya) for bioluminescent lineages.

| | | | Median age (Mya) in the pressent study (95% CI), Dataset A nuceotides AR | | Median age (Mya) in the pressent study (95% CI), Dataset A nucleotides IR | |
|---|---|---|---|---|---|---|
| **Bioluminescent lineage** | **Split of bioluminescent taxa from non-bioluminescent sister group** | **Crown split of bioluminescent lineage** | **Split of bioluminescent clade from a sister clade.** | **Crown age of bioluminescent clade (if more bioluminescent species were sampled for lineage).** | **Split of bioluminescent clade from a sister clade.** | **Crown age of bioluminescent clade (if more bioluminescent species were sampled for lineage).** |
| Pyrophorini | Pyrophorini-Platycrepidiini | *Ignelater*-rest of Pyrophorini | 71.57 (86.16 to 56.80) | 53.41 (67.01 to 40.24) | 39.79 (52.17 to 27.89) | 26.83 (35.06 to 19.27) |
| Lampyroid clade | Lampyroid-Elateroid clade | Sinopyrophoridae-rest of Lampyroid clade | 204.73 (213.58 to 196.23) | 191.79 (200.59 to 183.66) | 203.23 (217.78 to 187.08) | 189.46 (204.73 to 172.64) |
| NA (*Balgus schnusei*) | *Balgus obsconicus*-*Cussolenis* sp. (without bioluminescent *Balgus schnusei*) | | 91.77 (116.36 to 66.20) | NA | 80.69 (110.31 to 53.41) | NA |
| *Campyloxenus pyrothorax* | *Campyloxenus pyrothorax*-*Tibionema abdominalis* | | 41.12 (59.69 to 22.27) | NA | 32.70 (50.13 to 16.58) | NA |

| | | | Median age (Mya) in the pressent study (95% CI), Dataset B nuceotides AR | | Median age (Mya) in the pressent study (95% CI), Dataset B nucleotides IR | |
|---|---|---|---|---|---|---|
| **Bioluminescent lineage** | **Split of bioluminescent taxa from non-bioluminescent sister group** | **Crown split of bioluminescent lineage** | **Split of bioluminescent clade from a sister clade.** | **Crown age of bioluminescent clade (if more bioluminescent species were sampled for lineage).** | **Split of bioluminescent clade from a sister clade.** | **Crown age of bioluminescent clade (if more bioluminescent species were sampled for lineage).** |
| Pyrophorini | Pyrophorini-Platycrepidiini | *Ignelater*-rest of Pyrophorini | 68.79 (84.71 to 53.10) | 49.91 (64.09 to 36.28) | 36.87 (48.75 to 25.62) | 23.96 (31.44 to 17.16) |
| Lampyroid clade | Lampyroid-Elateroid clade | Sinopyrophoridae-rest of Lampyroid clade | 200.85 (209.17 to 192.17) | 189.12 (197.65 to 180.72) | 196.59 (212.78 to 179.04) | 183.37 (200.52 to 165.95) |
| NA (*Balgus schnusei*) | bioluminescent *Balgus schnusei*) | | 88.55 (112.72 to 63.61) | NA | 76.69 (105.43 to 50.42) | NA |
| *Campyloxenus pyrothorax* | *Campyloxenus pyrothorax*-*Tibionema abdominalis* | | 38.20 (56.24 to 20.95) | NA | 30.39 (46.69 to 15.83) | NA |

| | | | Median age (Mya) in the pressent study (95% CI), Dataset A amino acids, AR | | Median age (Mya) in the pressent study (95% CI), Dataset A amino acids, IR | |
|---|---|---|---|---|---|---|
| **Bioluminescent lineage** | **Split of bioluminescent taxa from non-bioluminescent sister group** | **Crown split of bioluminescent lineage** | **Split of bioluminescent clade from a sister clade.** | **Crown age of bioluminescent clade (if more bioluminescent species were sampled for lineage).** | **Split of bioluminescent clade from a sister clade.** | **Crown age of bioluminescent clade (if more bioluminescent species were sampled for lineage).** |
| Pyrophorini | Pyrophorini-Platycrepidiini | *Ignelater*-rest of Pyrophorini | 73.79 (87.09 to 60.21) | 60.10 (72.57 to 47.57) | 48.25 (60.31 to 37.02) | 37.48 (47.49 to 27.84) |
| Lampyroid clade | Lampyroid-Elateroid clade | Sinopyrophoridae-rest of Lampyroid clade | 186.54 (196.18 to 176.91) | 173.80 (184.17 to 164.00) | 184.40 (199.26 to 170.05) | 172.50 (187.63 to 158.36) |
| NA (*Balgus schnusei*) | bioluminescent *Balgus schnusei*) | | 81.83 (102.33 to 61.89) | NA | 66.32 (89.86 to 43.31) | NA |
| *Campyloxenus pyrothorax* | *Campyloxenus pyrothorax*-*Tibionema abdominalis* | | 41.97 (57.41 to 27.26) | NA | 31.01 (46.25 to 17.11) | NA |

| | | | Median age (Mya) in the pressent study (95% CI), Dataset B amino acids, AR | | Median age (Mya) in the pressent study (95% CI), Dataset B amino acids, IR | |
|---|---|---|---|---|---|---|
| **Bioluminescent lineage** | **Split of bioluminescent taxa from non-bioluminescent sister group** | **Crown split of bioluminescent lineage** | **Split of bioluminescent clade from a sister clade.** | **Crown age of bioluminescent clade (if more bioluminescent species were sampled for lineage).** | **Split of bioluminescent clade from a sister clade.** | **Crown age of bioluminescent clade (if more bioluminescent species were sampled for lineage).** |
| Pyrophorini | Pyrophorini-Platycrepidiini | *Ignelater*-rest of Pyrophorini | 69.17 (83.69 to 54.52) | 54.88 (68.25 to 42.08) | 42.63 (54.97 to 31.67) | 30.95 (40.11 to 22.37) |
| Lampyroid clade | Lampyroid-Elateroid clade | Sinopyrophoridae-rest of Lampyroid clade | 184.08 (194.07 to 175.14) | 171.08 (181.31 to 161.16) | 181.85 (197.14 to 167.35) | 169.45 (184.71 to 155.10) |
| NA (*Balgus schnusei*) | bioluminescent *Balgus schnusei*) | | 76.77 (97.24 to 55.51) | NA | 64.50 (88.66 to 41.13) | NA |
| *Campyloxenus pyrothorax* | *Campyloxenus pyrothorax*-*Tibionema abdominalis* | | 41.33 (57.10 to 26.40) | NA | 30.22 (45.40 to 16.03) | NA |

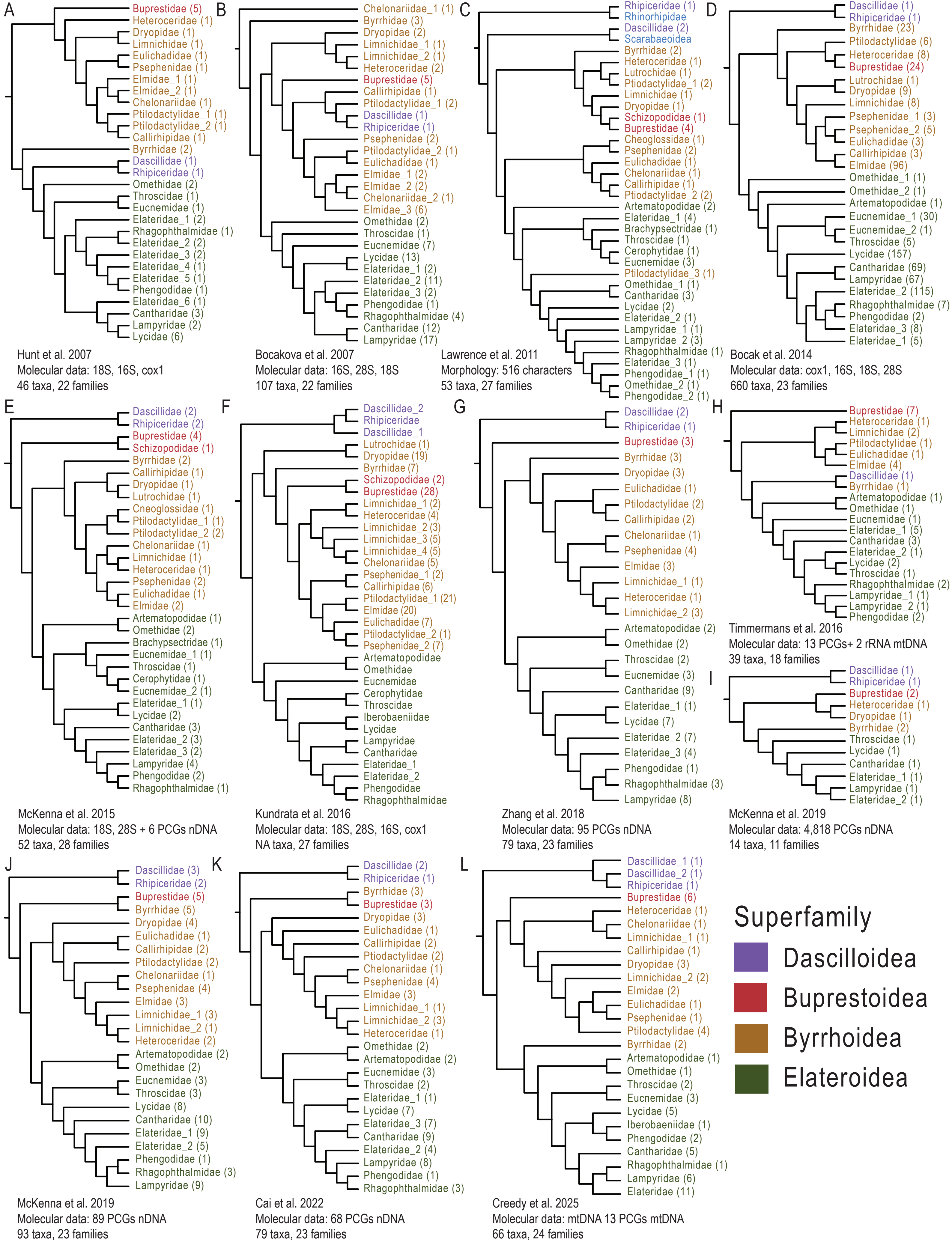

**Supplementary Figure S1.** History of phylogenetic relationships among Elateriformia families from selected studies based on morphology and molecular data. Adapted to match currently recognized family classification. A) Hunt et al., 2007 (Fig. 2); B) Bocakova et al., 2007 (Fig. 4; see also alternative topologies presented); C) Lawrence et al., 2011 (Cladogram 3); D) Bocak et al., 2014 (Fig. 2; see Fig. S1 for minor differences); E) McKenna et al., 2015 (Bayesian tree, Figs. 8–9; see also Maximum Likelihood tree); F) Kundra et al., 2016 (Fig. 2A; see Fig. S3 for 488-taxon tree; numbers of individual samples are not present in details for Elateroidea); G) Zhang et al., 2018 (Fig. 2); H) Timmermans et al., 2016 (Fig. 1; see also Fig. 3); I) McKenna et al., 2019 (Fig. 1); J) McKenna et al., 2019 (Fig. S10); K) Cai et al., 2022 (Fig. 1); L) Creedy et al., 2025 (Fig. 6; see also Fig. 4). Only sections of respective trees are shown. Numbers in parentheses represent the number of species sampled in respective studies. PCGs, protein-coding genes; nDNA, nuclear DNA; mtDNA, mitochondrial DNA. See also Muona 1995; Kundrata et al., 2014; Kusy et al., 2018a, b; Douglas et al. 2021; Hayashi et al., 2024 for further reference.

| | # spp. | % |
|---|---|---|
| Dascilloidea | 150 | 0,30 |
| Buprestoidea | 15007 | 29,73 |
| Byrrhoidea | 4285 | 8,488 |
| Elateroidea | 31043 | 61,49 |
| | 50485 | |

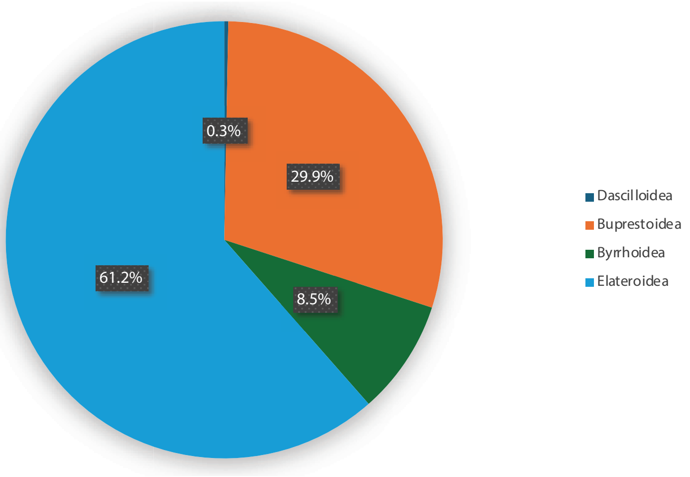

| Byrrhoidea | # spp. | % |
|---|---|---|
| Byrrhidae | 430 | 10,04 |
| Callirhipidae | 150 | 3,501 |
| Chelonariidae | 300 | 7,001 |
| Cladotomidae | 20 | 0,467 |
| Cneoglossidae | 9 | 0,21 |
| Dryopidae | 300 | 7,001 |
| Elmidae | 1500 | 35,01 |
| Eulichalidae | 30 | 0,7 |
| Heteroceridae | 349 | 8,145 |
| Limnichidae | 380 | 8,868 |
| Lutrochidae | 21 | 0,49 |
| Protelmidae | 6 | 0,14 |
| Psephenidae | 290 | 6,768 |
| Ptilodactylidae | 500 | 11,67 |
| | 4285 | |

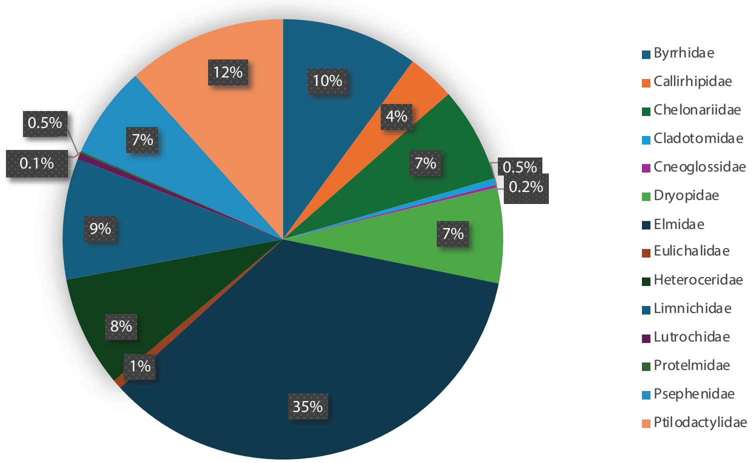

| Elateroidea | # spp. | % |
|---|---|---|
| Artematopodidae | 74 | 0,238 |
| Badmaateridae | 1 | 0,003 |
| Brachypsectridae | 9 | 0,029 |
| Cantharidae | 6735 | 21,7 |
| Cerophytidae | 23 | 0,074 |
| Elateridae | 14600 | 47,03 |
| Eucnemidae | 1707 | 5,50 |
| Iberobaeniidae | 3 | 0,01 |
| Jurasaidae | 7 | 0,023 |
| Lampyridae | 2605 | 8,392 |
| Lycidae | 4629 | 14,91 |
| Omethidae | 64 | 0,206 |
| Phengodidae | 314 | 1,012 |
| Rhagophthalmidae | 71 | 0,229 |
| Sinopyrophoridae | 1 | 0,003 |
| Throscidae | 200 | 0,644 |
| | 31043 | |

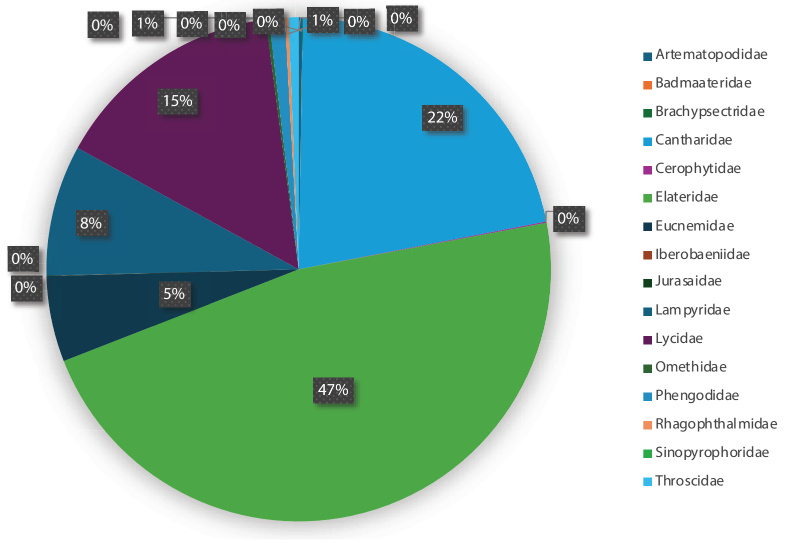

**Figure S2.** Numbers of described species in Elateriformia.

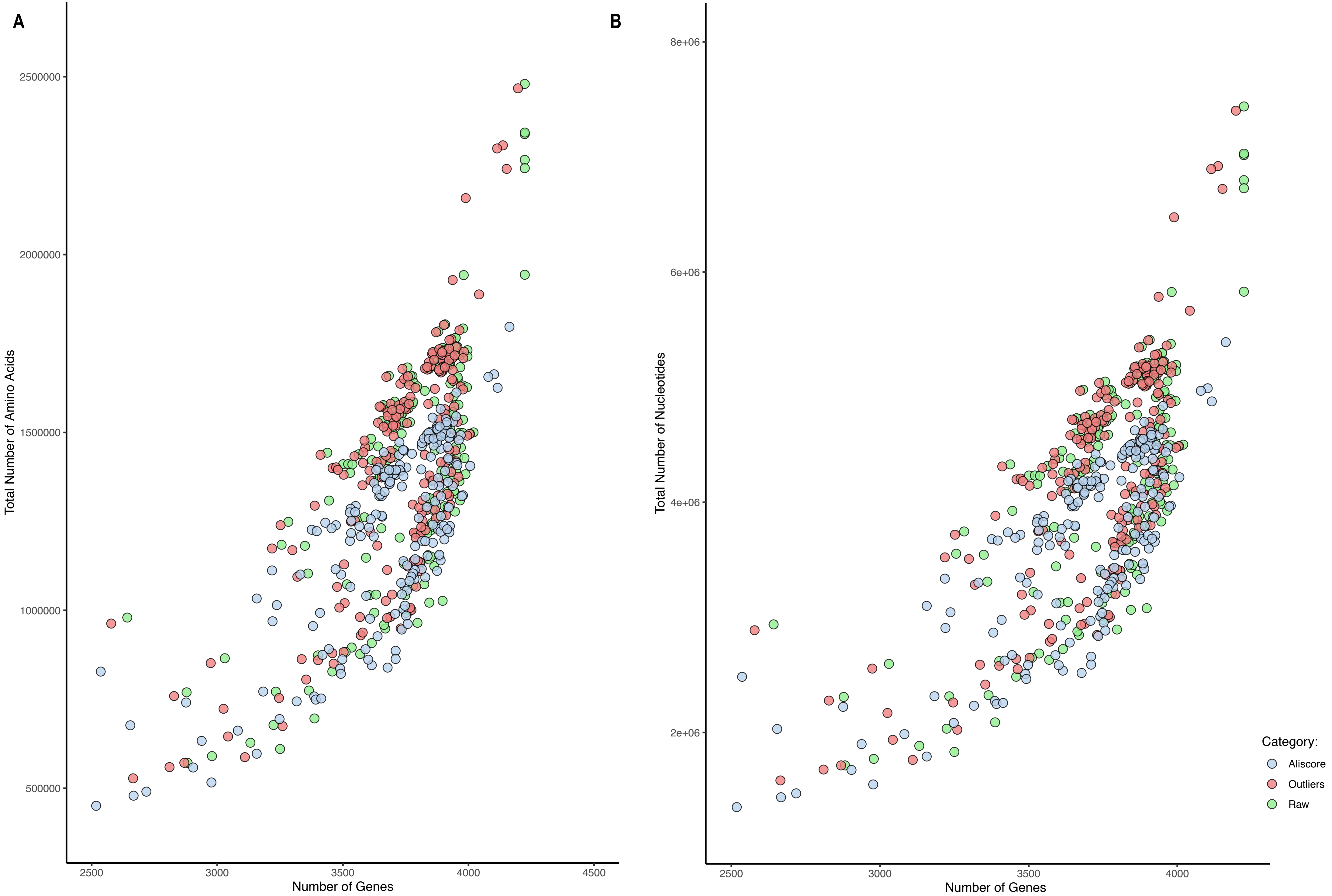

**Supplementary Figure S3.** Number of genes vs. total number of amino acids A) and nucleotides B) recovered for each of the categories. Three processing stages are shown: Raw (initial Orthograph output), Outliers (after filtering outlier sequences), and Aliscore (after outlier removal and ambiguous alignment filtering with Aliscore).

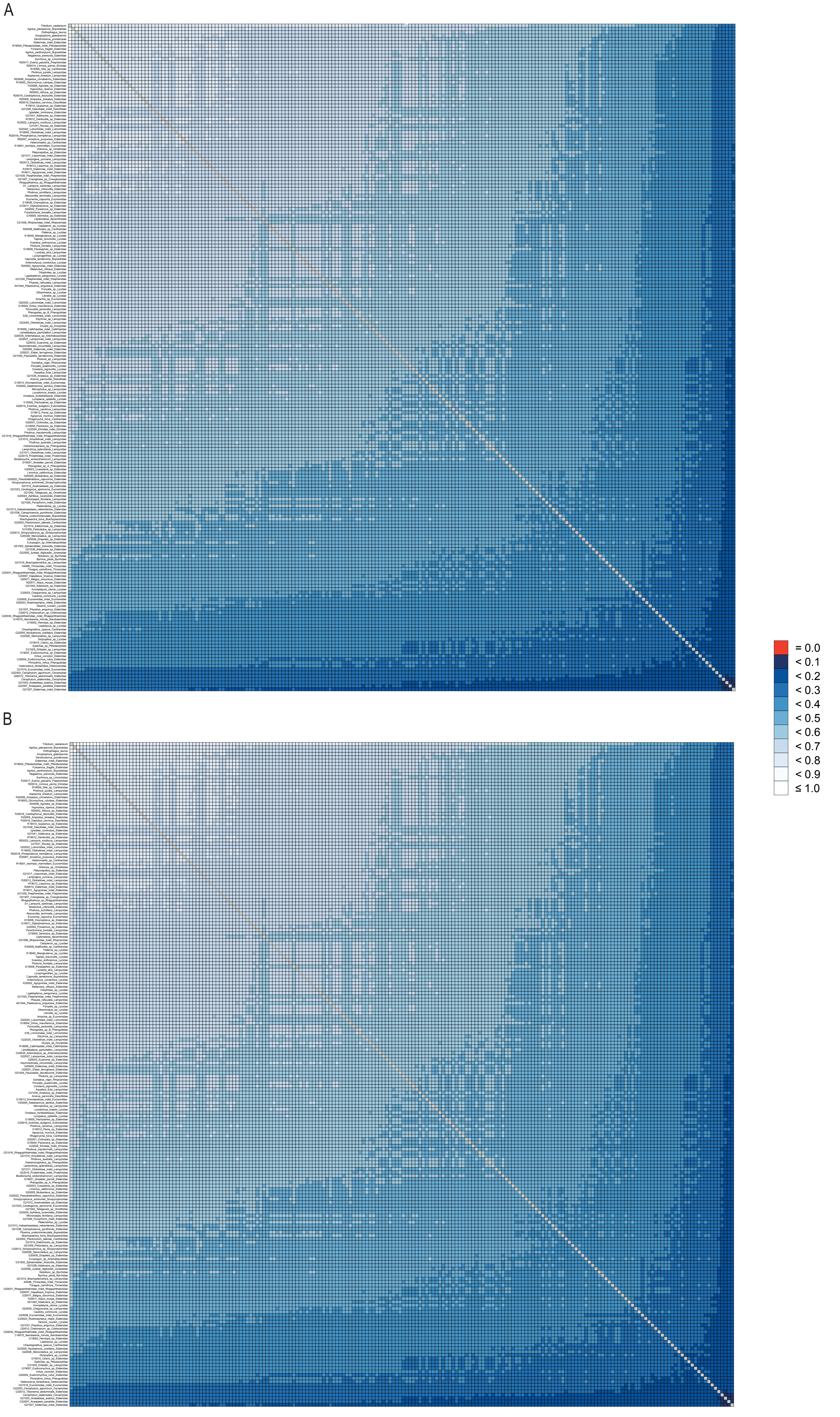

**Supplementary Figure S4.** AliStat heat maps showing pairwise alignment completeness for all species included in supermatrices: A) A-4189-AA, CA score 0.661537 and B) B-4189-NT, CA score for single nucleotides 0.661539 and 0.661535 for codons.

A

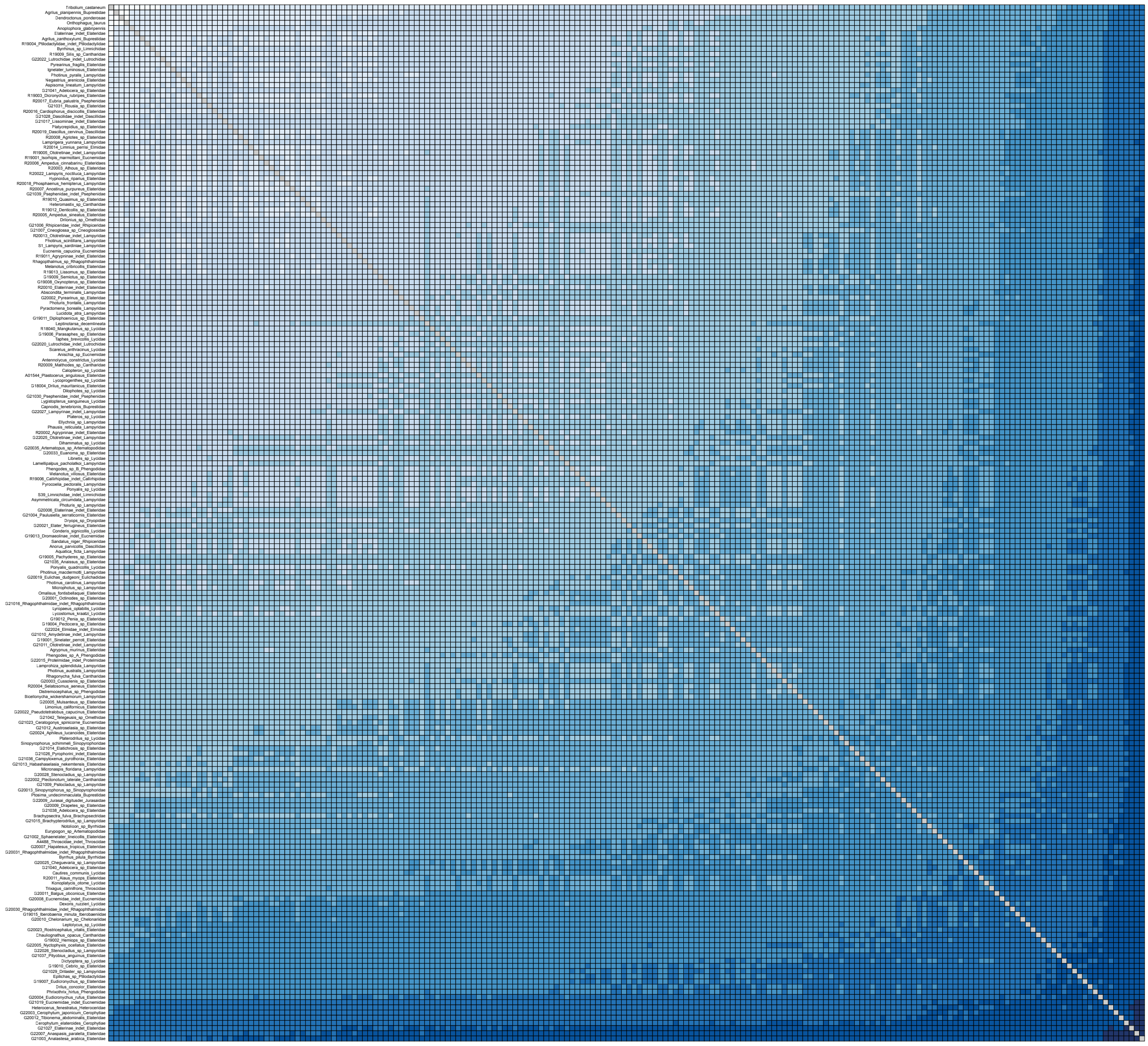

B

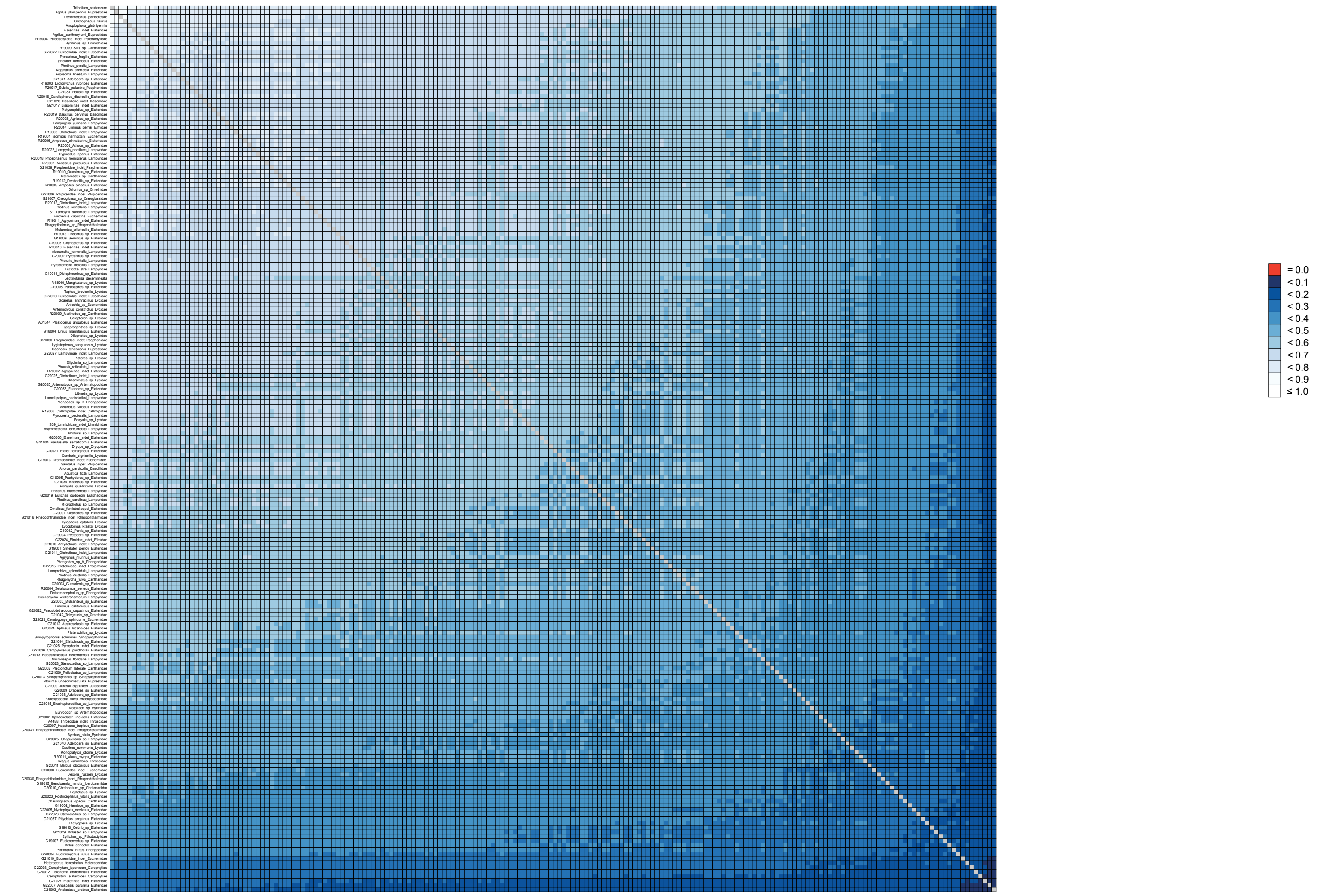

**Supplementary Figure S5.** AliStat heat maps showing pairwise alignment completeness for all species included in supermatrices: A) I-1451-AA-Bacoca, CA score 0.652969 and B) J-1451-NT-Bacoca, CA score 0.652971.

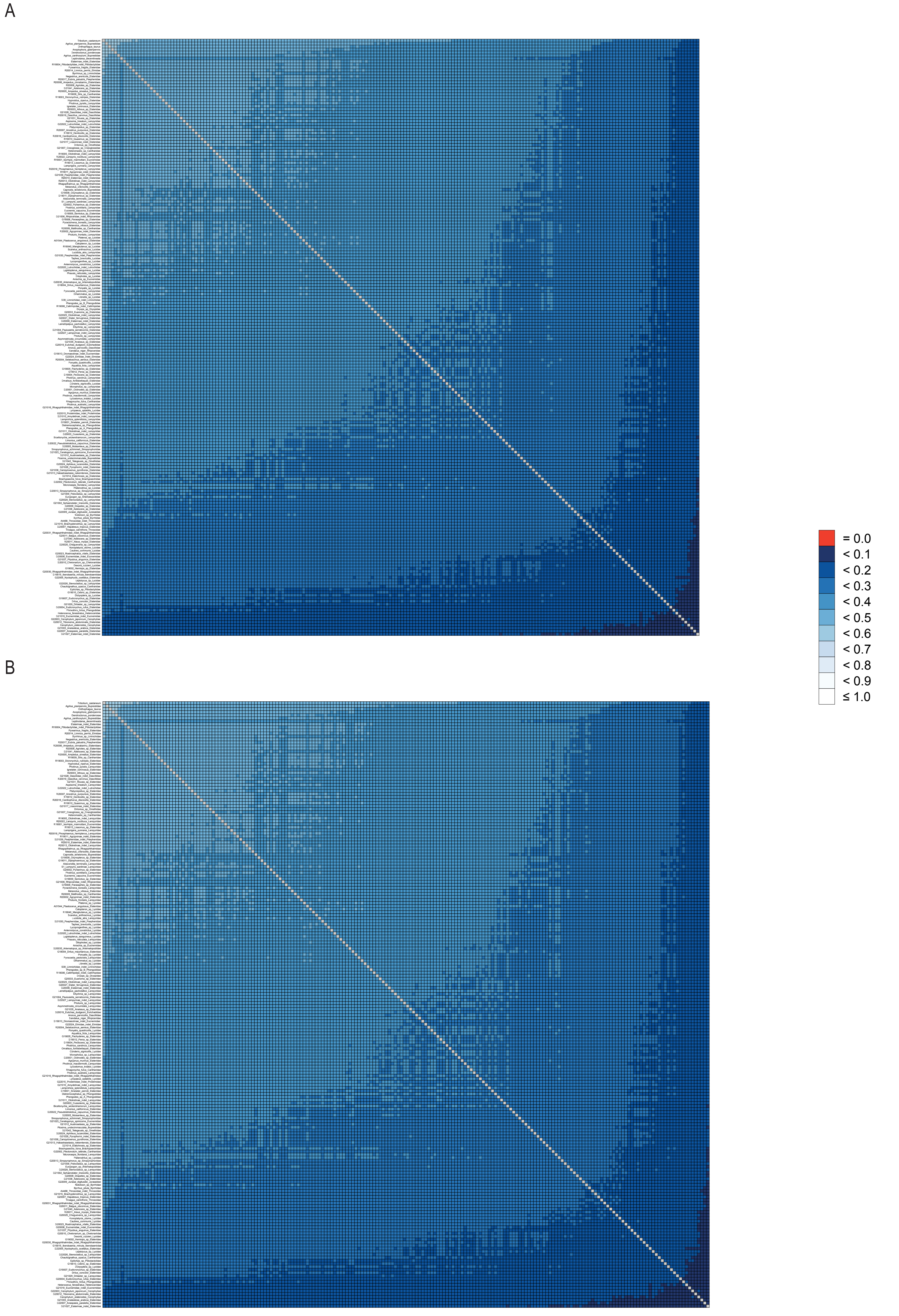

**Supplementary Figure S6.** AliStat heat maps showing pairwise alignment completeness for all species included in supermatrices: A) F-4224-NT-raw, CA score 0.419819 and B) E-4224-AA-raw, CA score0.419818.

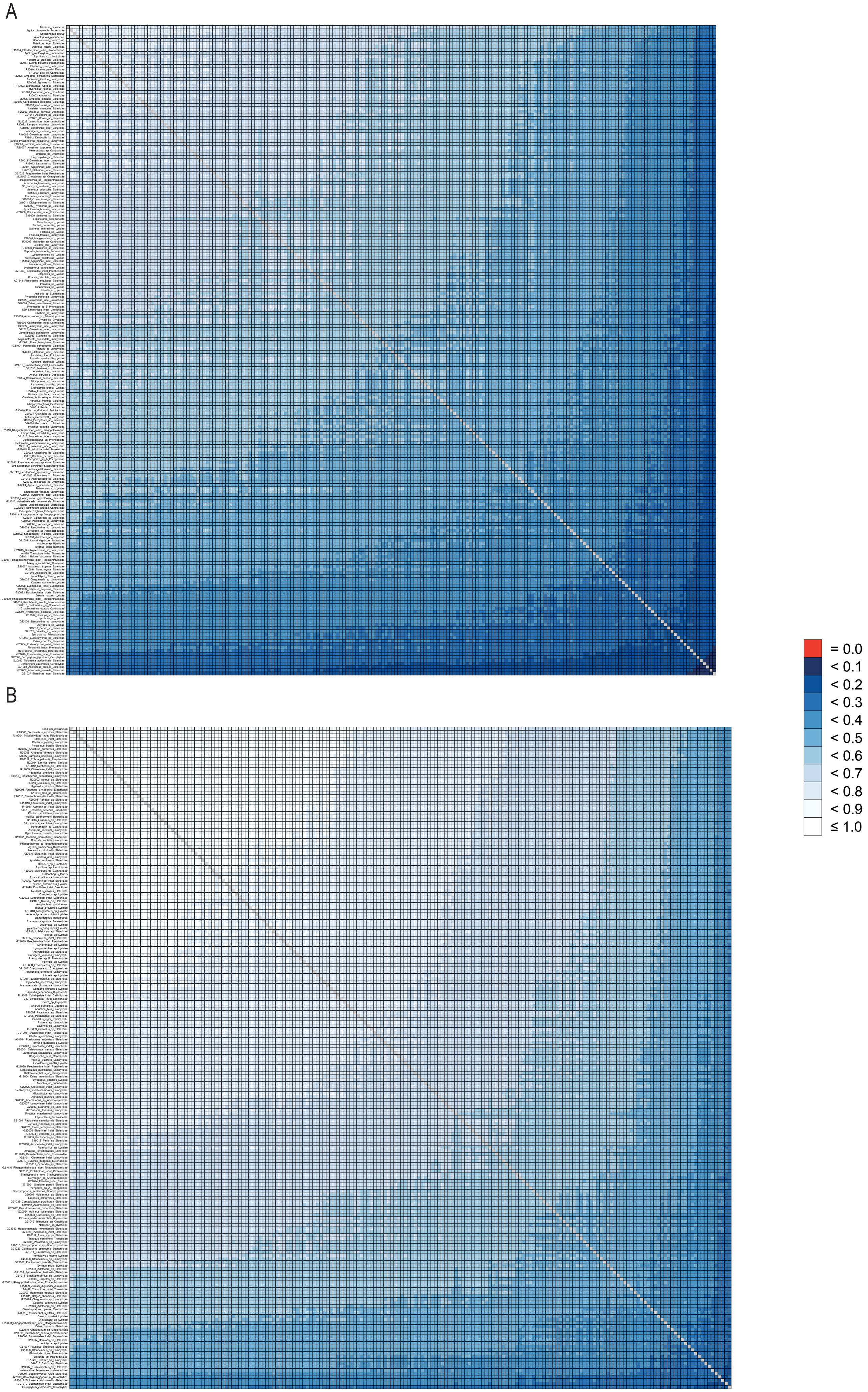

**Supplementary Figure S7.** AliStat heat maps showing pairwise alignment completeness for all species included in supermatrices: A) K-3583-NT-SymTest, CA score 0.651593 and B) D-1892-AA-MARE, CA score 0.765498.

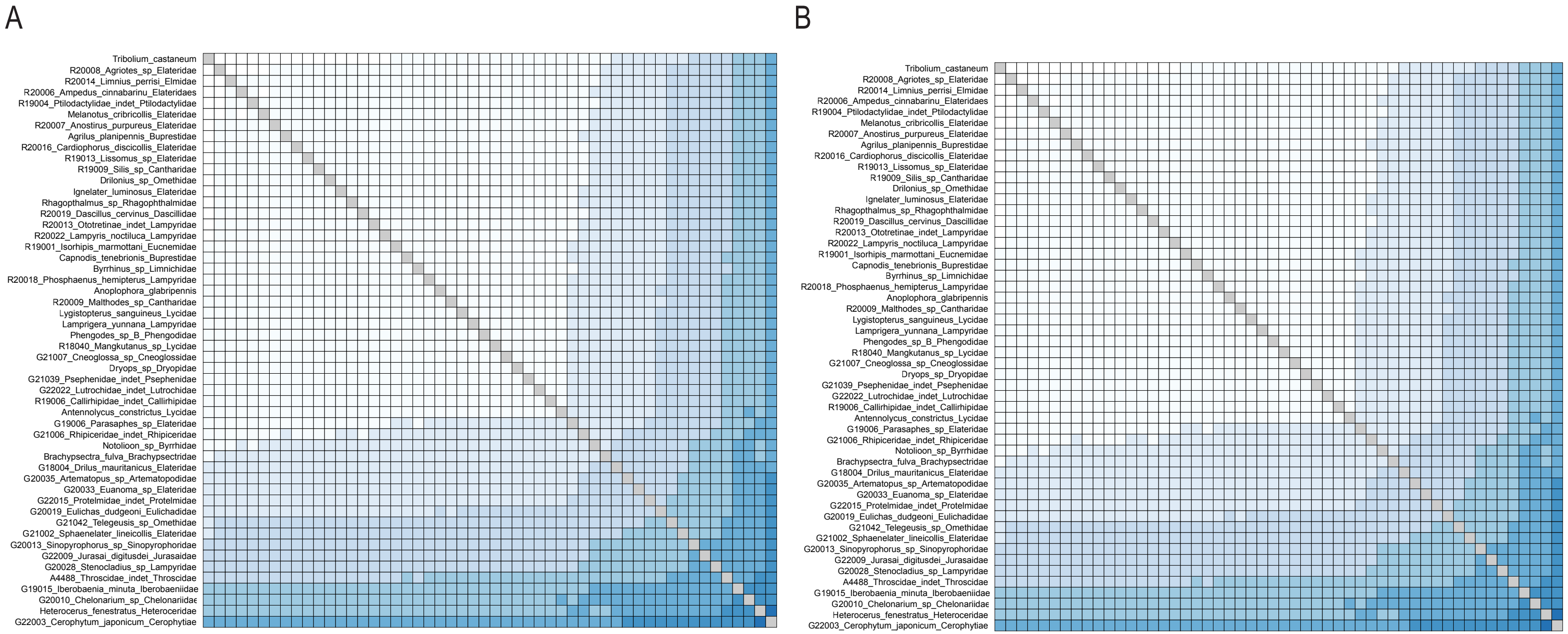

**Supplementary Figure S8.** AliStat heat maps showing pairwise alignment completeness for all species included in supermatrices: A) G-962-AA-decisive, CA score 0.837545 and B) H-962-NT-decisive, CA score 0.837547.

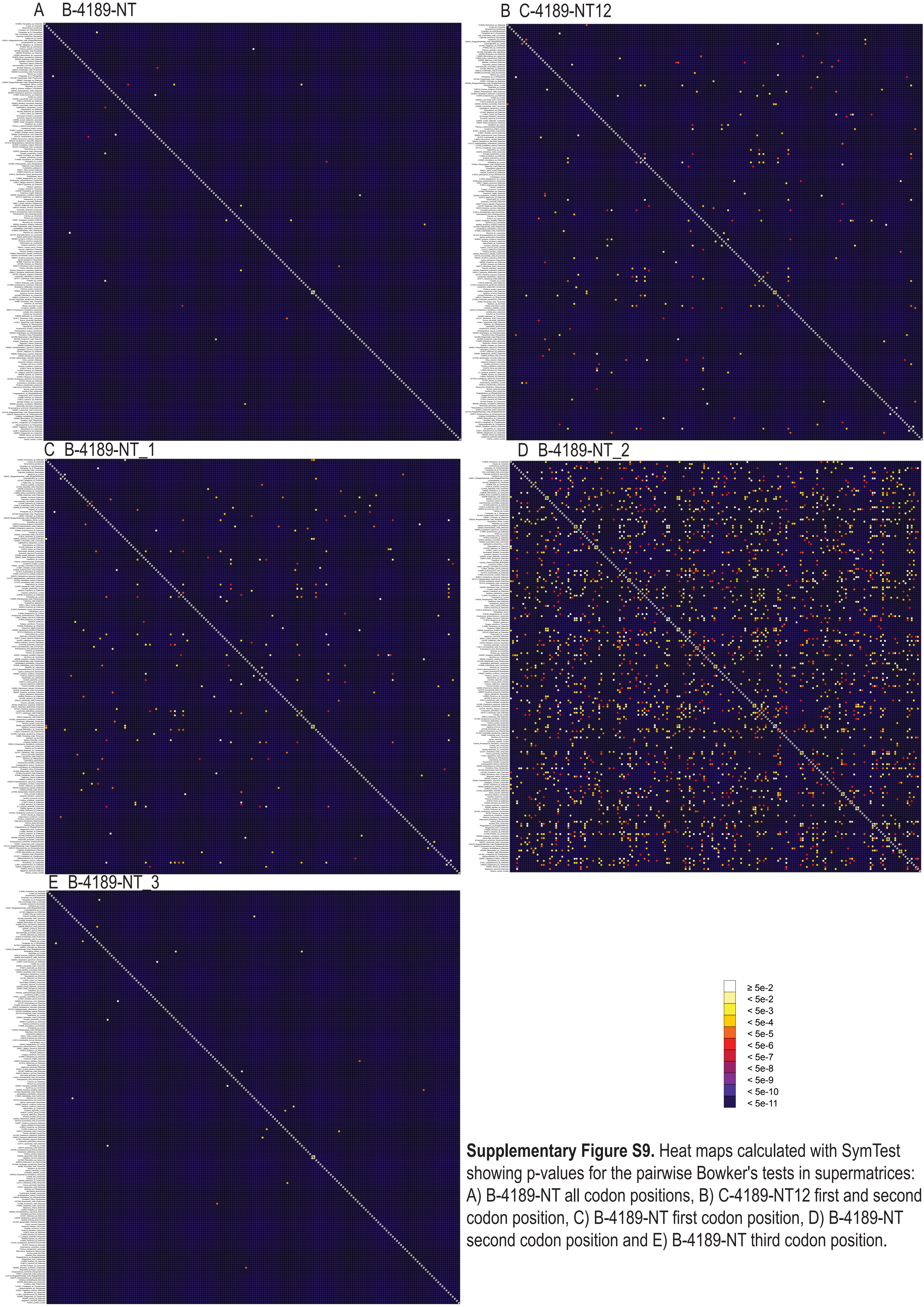

**Supplementary Figure S9.** Heat maps calculated with SymTest showing p-values for the pairwise Bowker's tests in supermatrices: A) B-4189-NT all codon positions, B) C-4189-NT12 first and second codon position, C) B-4189-NT first codon position, D) B-4189-NT second codon position and E) B-4189-NT third codon position.

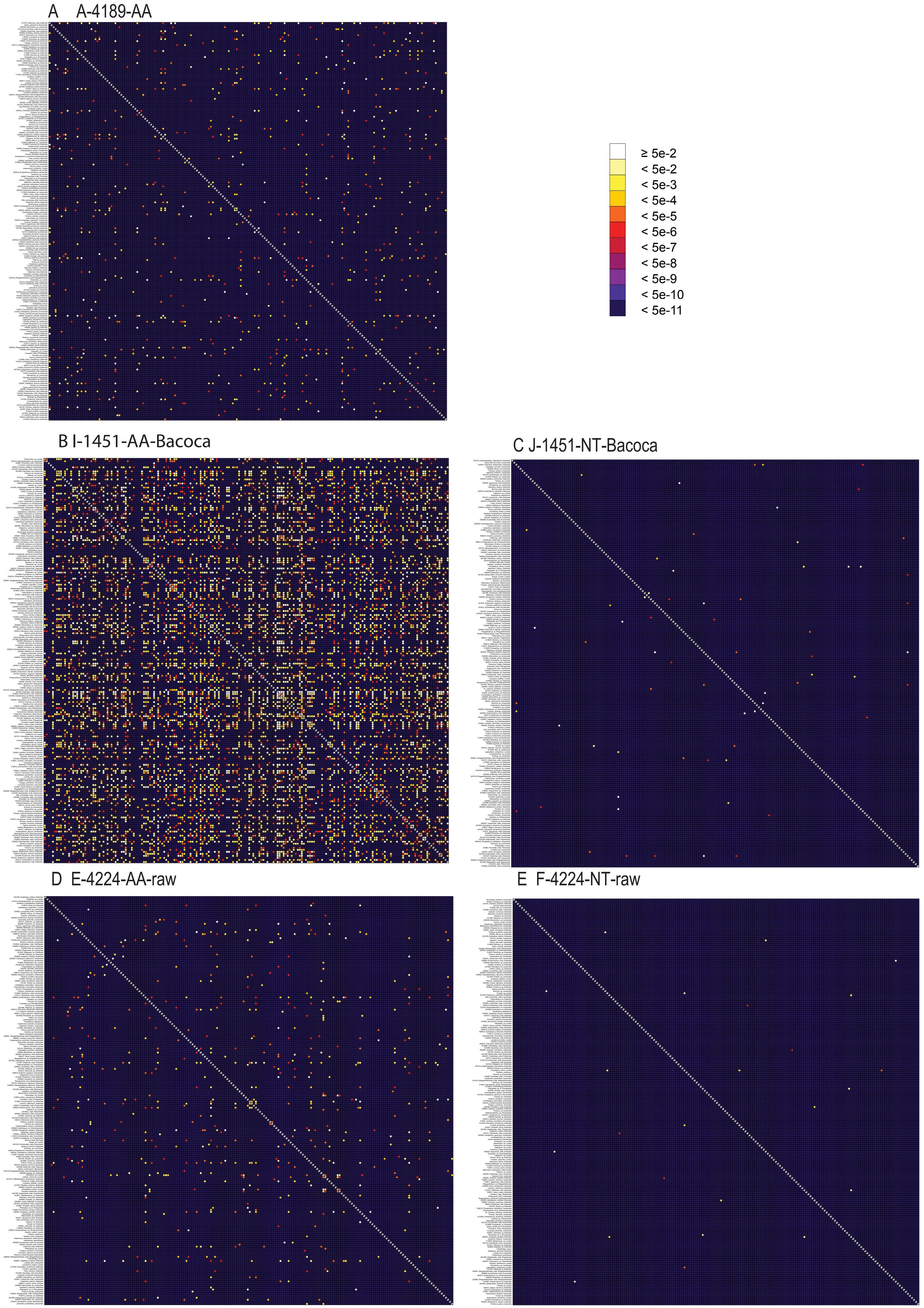

**Supplementary Figure S10.** Heat maps calculated with SymTest showing p-values for the pairwise Bowker's tests in supermatrices: A) A-4189-AA, B) I-1451-AA-Bacoca, C) J-1451-NT-Bacoca, D) E-4224-AA-raw and E) F-4224-NT-raw.

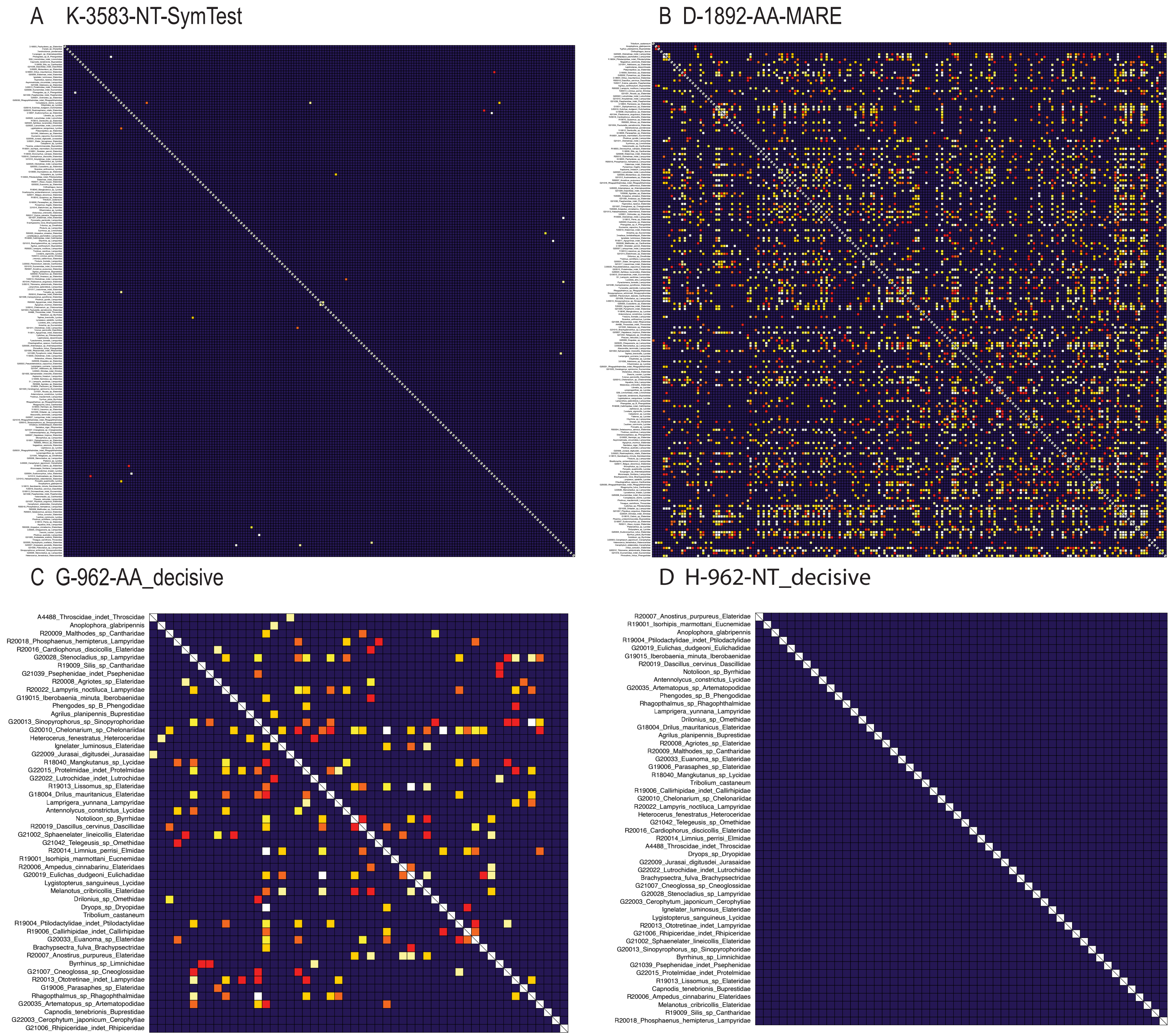

**Supplementary Figure S11.** Heat maps calculated with SymTest showing p-values for the pairwise Bowker's tests in supermatrices: A) K-3583-NT-SymTest, B) D-1892-AA-MARE, C) G-962-AA-decisive and D) H-962-NT-decisive.

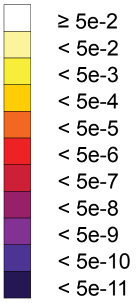

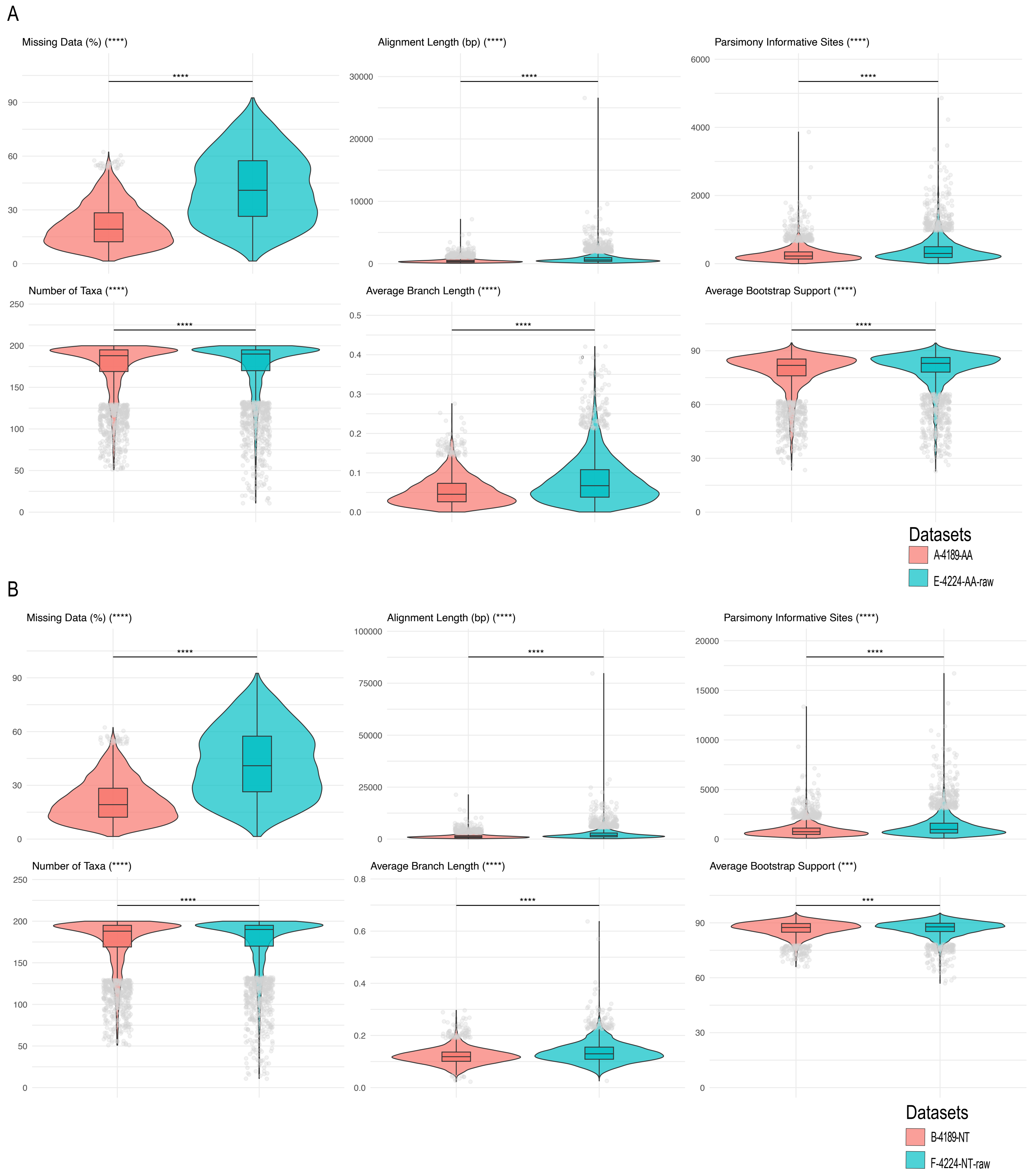

**Supplementary Figure S12.** Comparison of characteristics between datasets 4189 (coral) and 4224 (teal) for amino acid A) and nucleotide B) sequence alignments. Violin plots display distributions of missing data percentage, alignment length, parsimony informative sites, average branch length, average bootstrap support, and number of taxa. Statistical significance determined by Wilcoxon rank-sum tests: *** $p < 0.001$, **** $p < 0.0001$. All metrics showed significant differences between datasets. Grey points represent outliers identified using the IQR method (values beyond Q1 - 1.5×IQR or Q3 + 1.5×IQR).

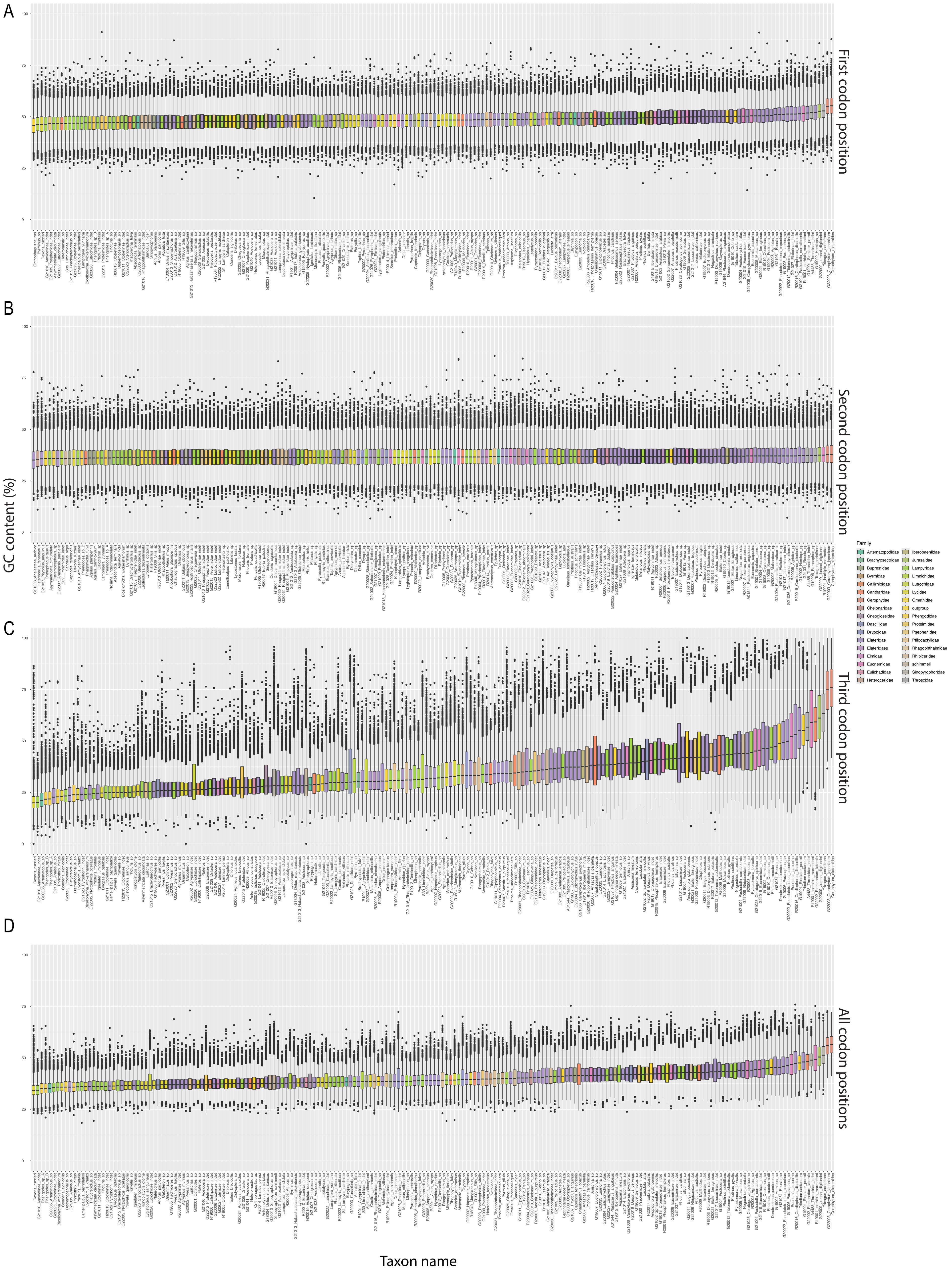

**Supplementary Figure S13.** GC content plot for supermatrix B-4189-NT A) first codon position; B) second codon position; C) third codon position and D) all three codon positions.

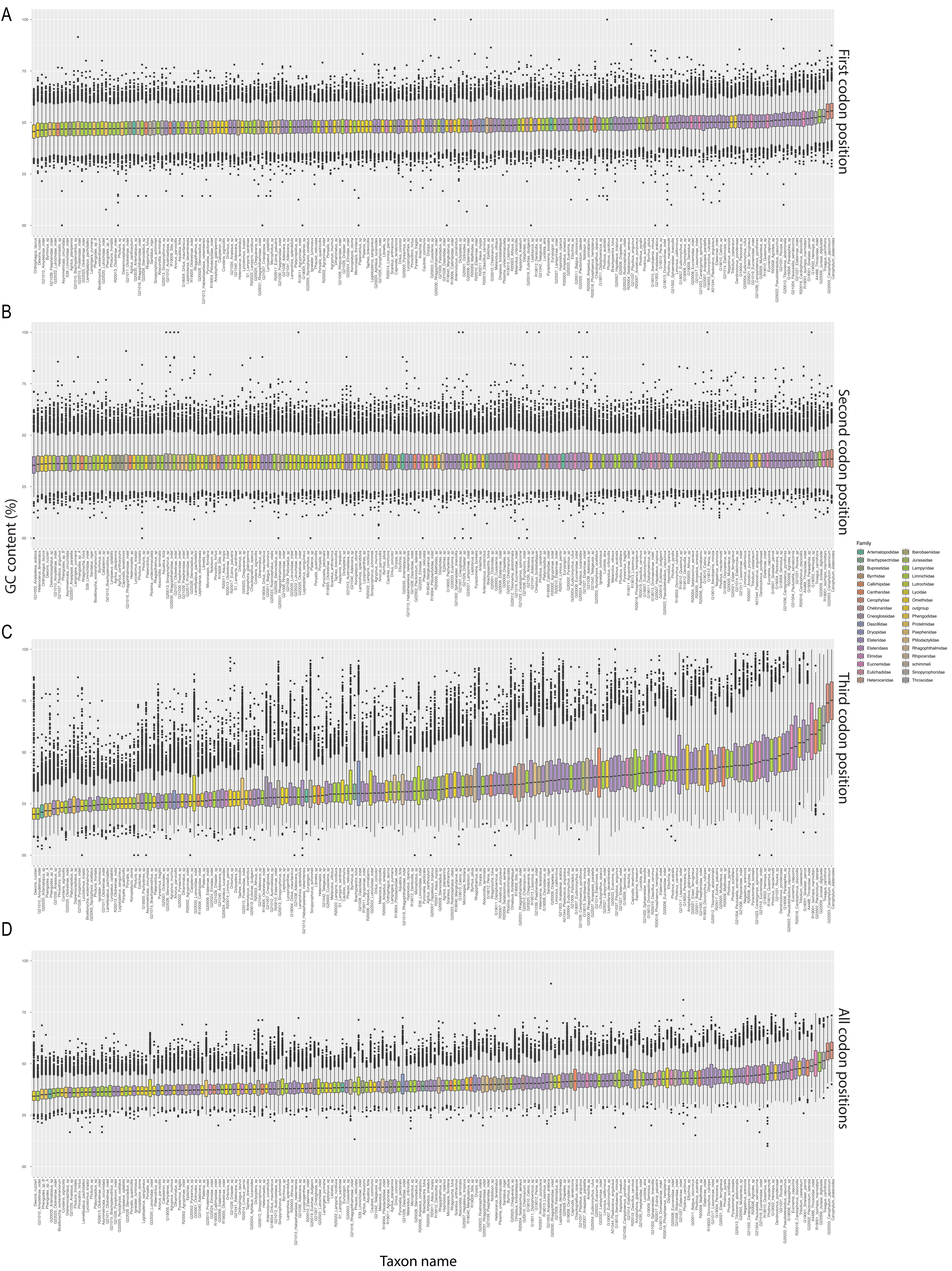

**Supplementary Figure S14.** GC content plot for supermatrix F-4224-NT-raw A) first codon position; B) second codon position; C) third codon position and D) all three codon positions.

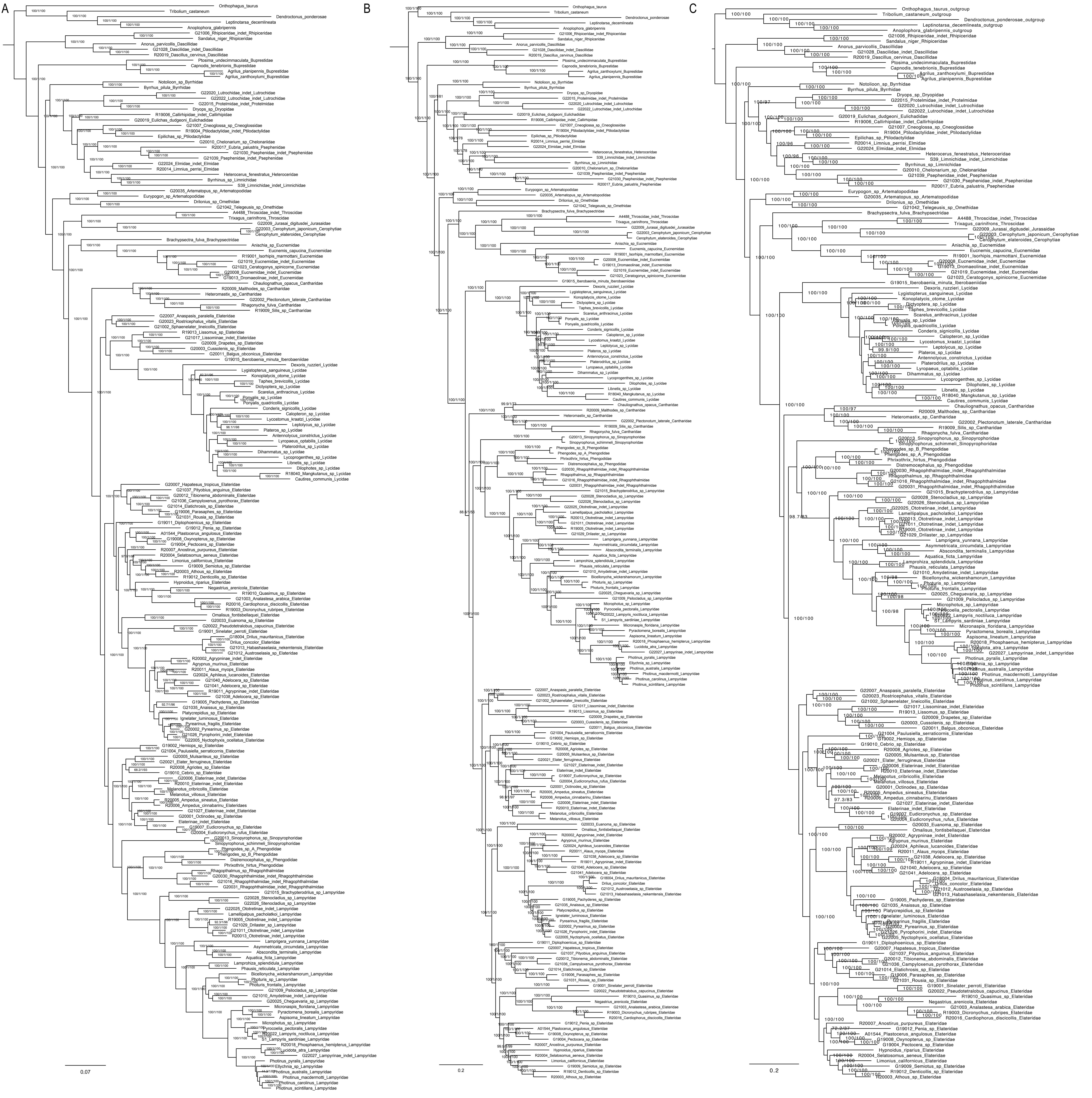

**Supplementary Figure S15.** Maximum likelihood trees from IQ-TREE analyses of the following supermatrices: A) A-4189-AA supermatrix partitioned by genes; B) B-4189-NT supermatrix partitioned by genes with only GTR model considered in model selection step and C) B-4189-NT supermatrix partitioned by genes with all available models considered in model selection step -m MFP. The depicted branch support values are based on SH-aLRT and ultrafast bootstrap (3000 replicates each).

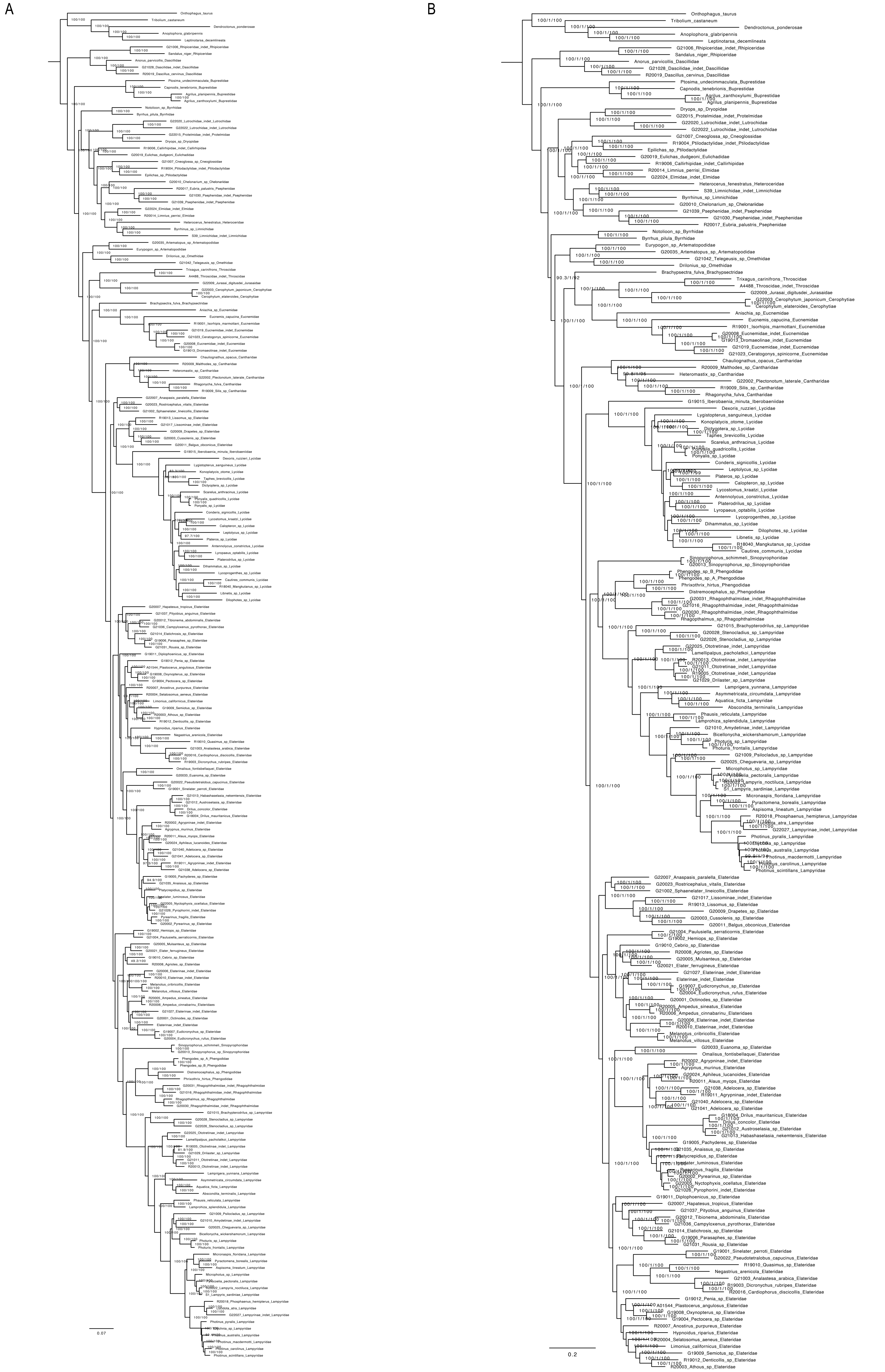

**Supplementary Figure S16.** Maximum likelihood trees from IQ-TREE analyses of the following supermatrices: A) A-4189-AA supermatrix analysed as single partition and B) B-4189-NT supermatrix analysed as single partition with only GTR model considered in model selection step. The depicted branch support values are based on SH-aLRT and ultrafast bootstrap (3000 replicates each). And approximate Bayes test in B).

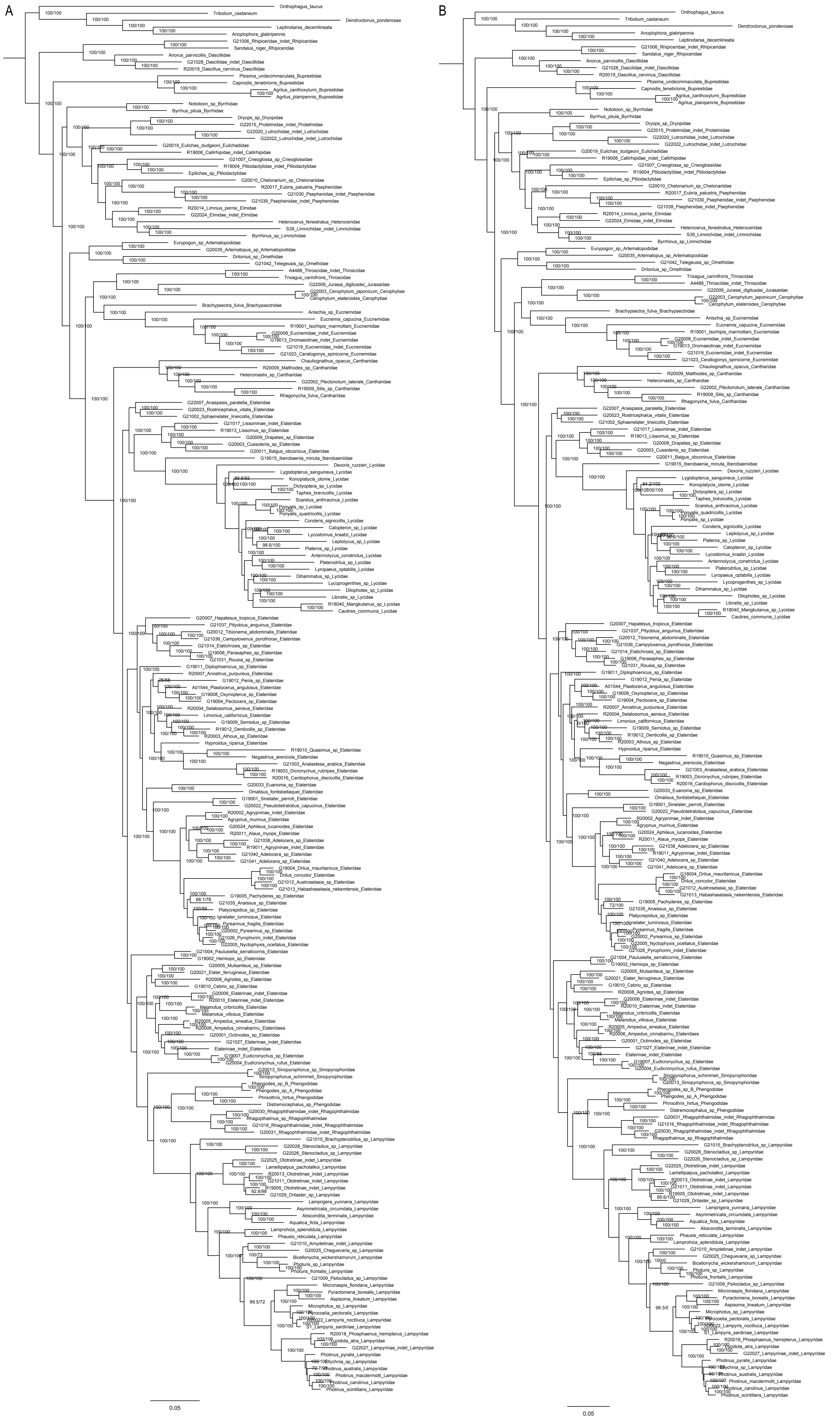

**Supplementary Figure S17.** Maximum likelihood trees from IQ-TREE analyses of the supermatrix C-4189-NT12 A) partitioned by genes and B) analysed as a single partition. Only the GTR model was considered in the model selection step. The depicted branch support values are based on SH-aLRT and ultrafast bootstrap (3000 replicates each).

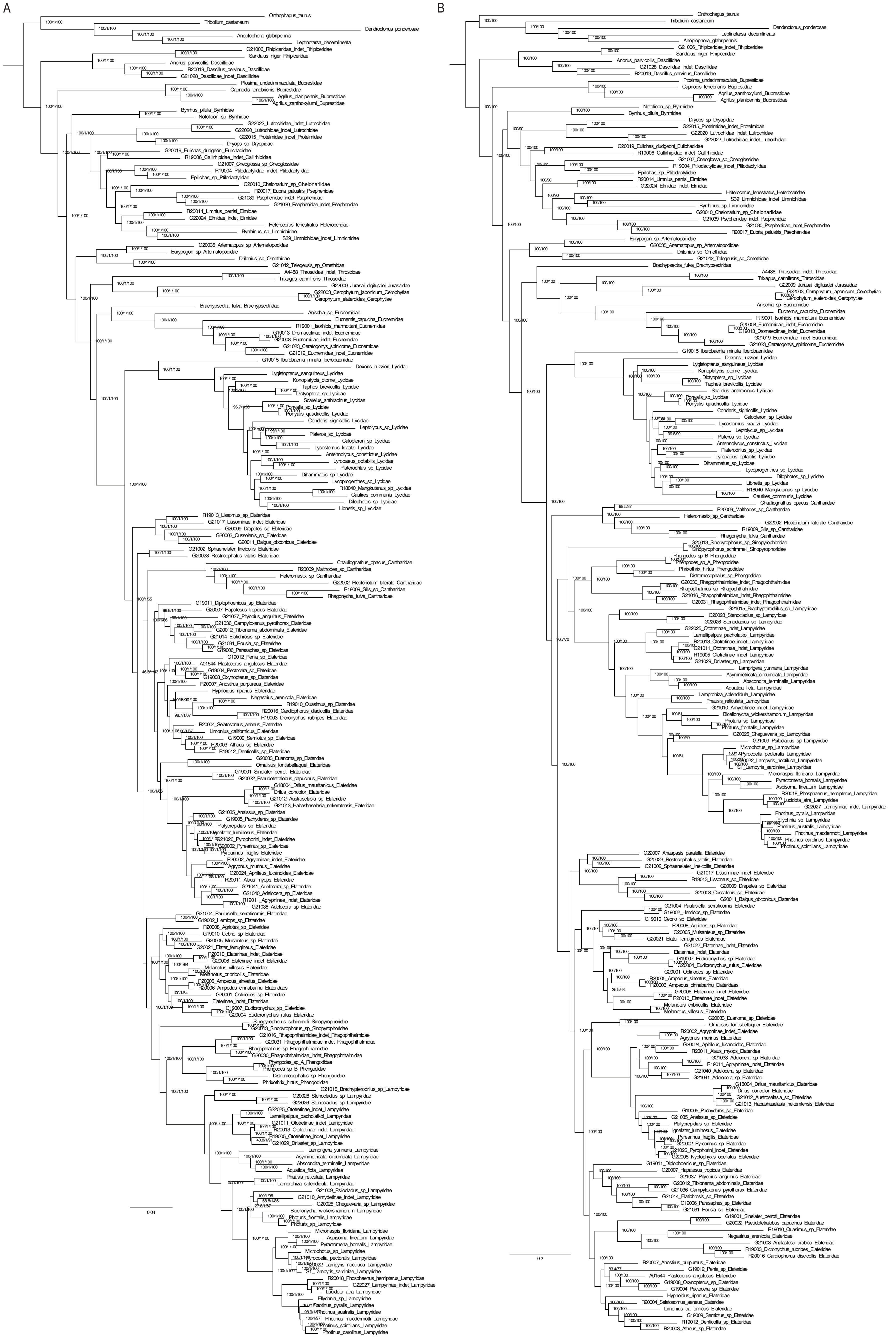

**Supplementary Figure S18.** Maximum likelihood trees from IQ-TREE analyses of the following supermatrices: A) D-1892-AA-MARE supermatrix partitioned by genes and B) K-3583-NT-SymTest supermatrix partitioned by genes with only GTR model considered in model selection step. The depicted branch support values are based on SH-aLRT and ultrafast bootstrap (3000 replicates each).

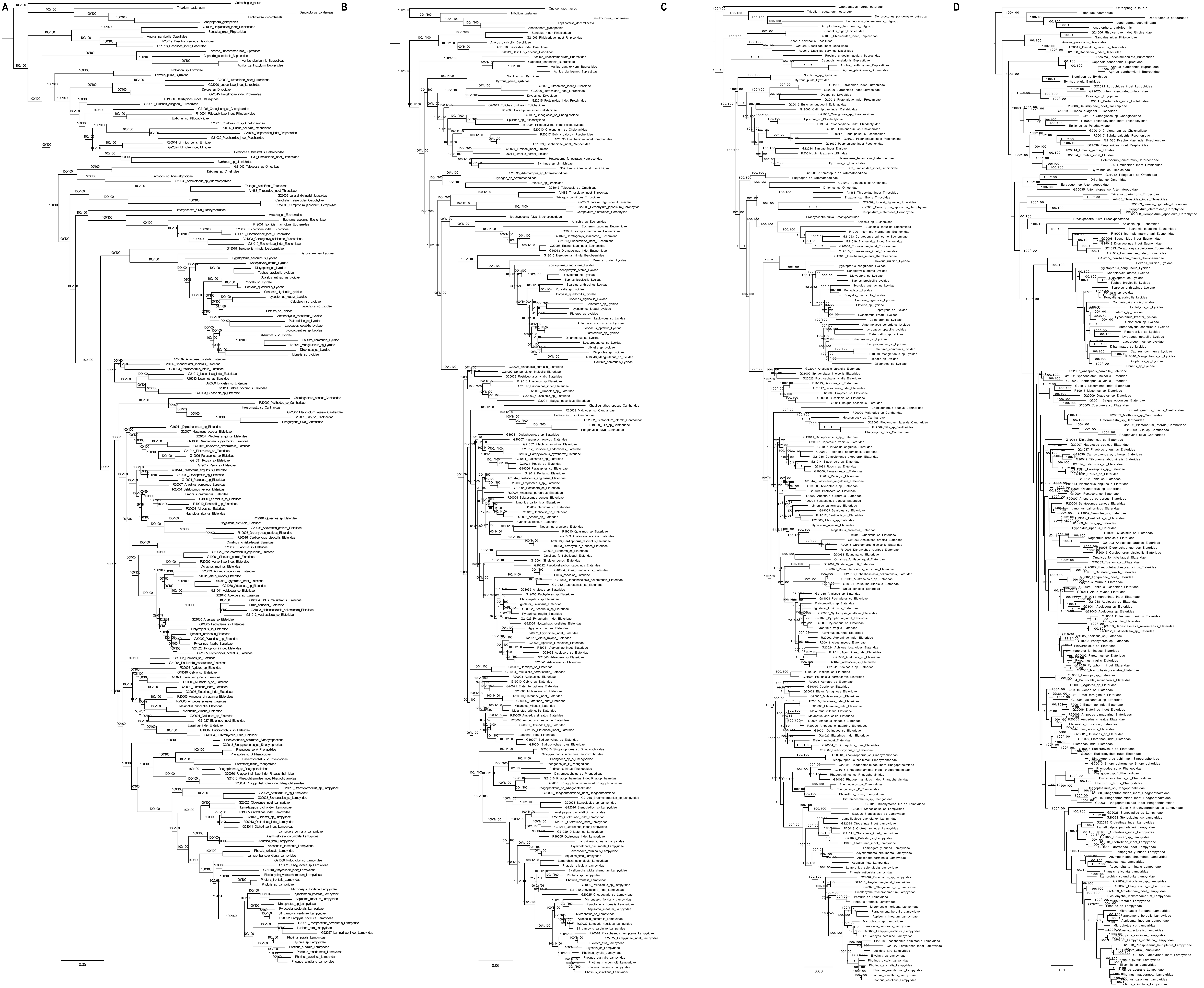

**Supplementary Figure S19.** Maximum likelihood trees from IQ-TREE analyses of the following supermatrices: A) I-1451-AA-Bacoca supermatrix analysed as single partition; B) I-1451-AA-Bacoca supermatrix partitioned by genes; C) I-1451-AA-Bacoca supermatrix with optimalized partitioning scheme options: –merge –rclusterf 10 and –rcluster-max set to default; D) I-1451-AA-Bacoca supermatrix analysed under heterogeneous posterior mean site frequency (PMSF) model.The depicted branch support values are based on SH-aLRT and ultrafast bootstrap (3000 replicates each).

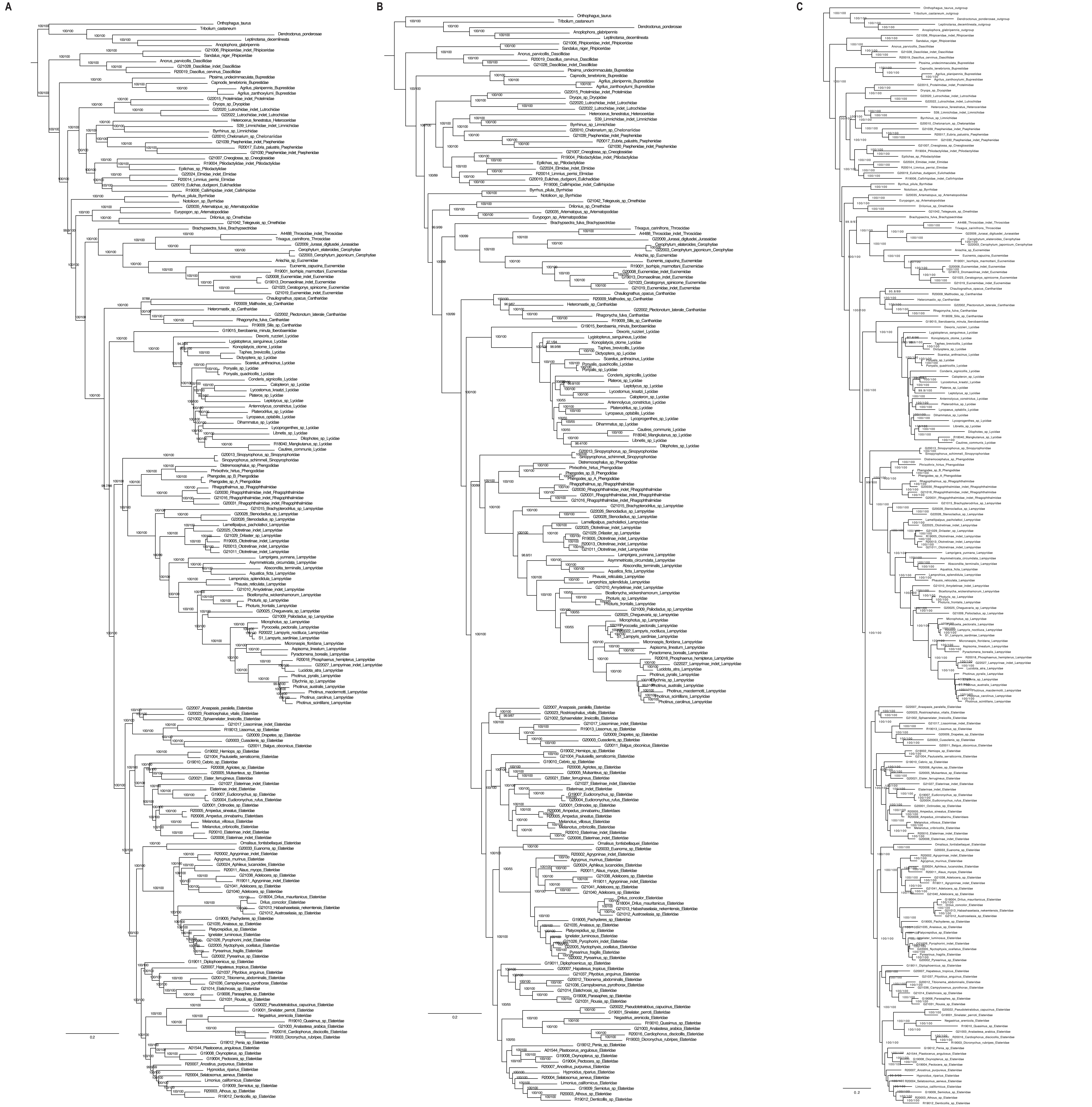

**Supplementary Figure S20.** Maximum likelihood trees from IQ-TREE analyses of the following supermatrices: A) J-1451-NT-Bacoca supermatrix partitioned by genes; B) J-1451-NT-Bacoca supermatrix analysed as single partition; C) J-1451-NT-Bacoca supermatrix with optimalized partitioning scheme options: –merge –rclusterf 10 and –rcluster-max set to default. The depicted branch support values are based on SH-aLRT and ultrafast bootstrap (3000 replicates each).

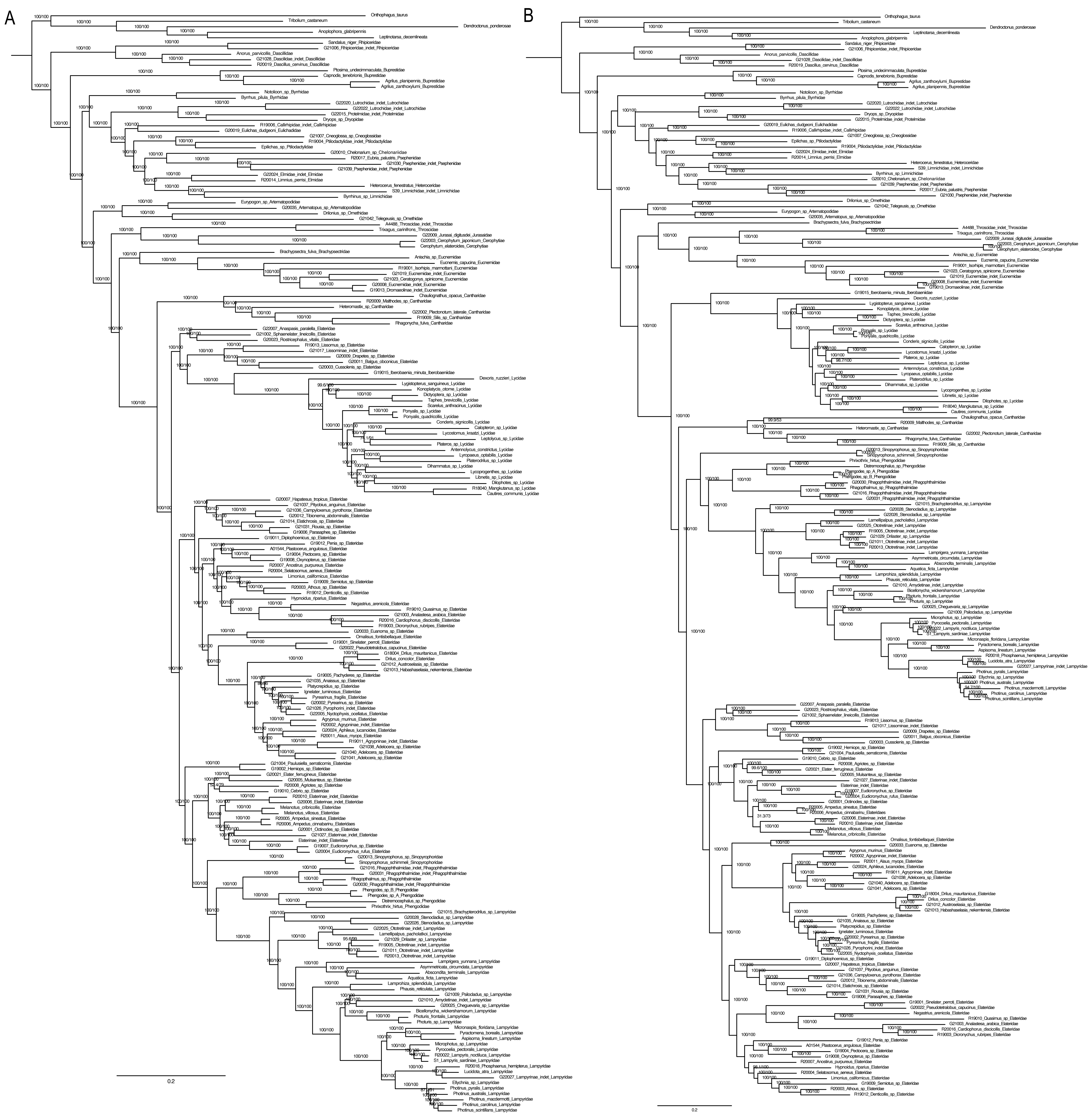

**Supplementary Figure S21.** Maximum likelihood trees from IQ-TREE analyses of the following supermatrices: A) E-4224-AA-raw supermatrix partitioned by genes; B) F-4224-NT-raw supermatrix partitioned by genes with only GTR model considered in model selection step. The depicted branch support values are based on SH-aLRT and ultrafast bootstrap (3000 replicates each).

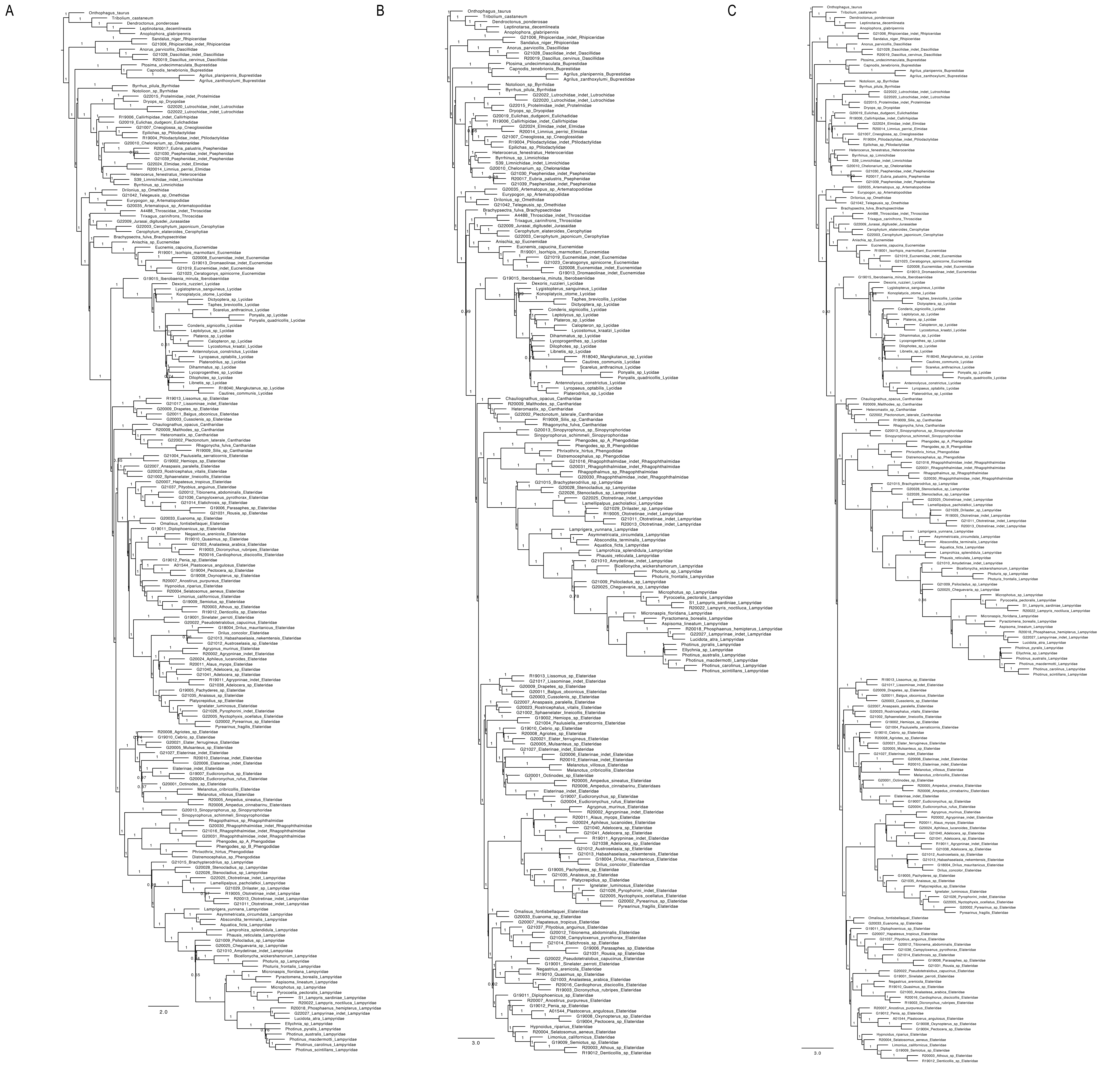

**Supplementary Figure S22.** Coalescent species trees inferred by ASTRAL from individual gene trees. Maximum likelihood gene trees were constructed using IQ-TREE with datasets A) A-4189-AA; B) B-4189-NT and C) K-3583-NT-SymTest. Branches with ultra-fast bootstrap support ≤50 were collapsed prior to coalescent analysis. Branch lengths are shown in coalescent units, and node support values represent local posterior probabilities.

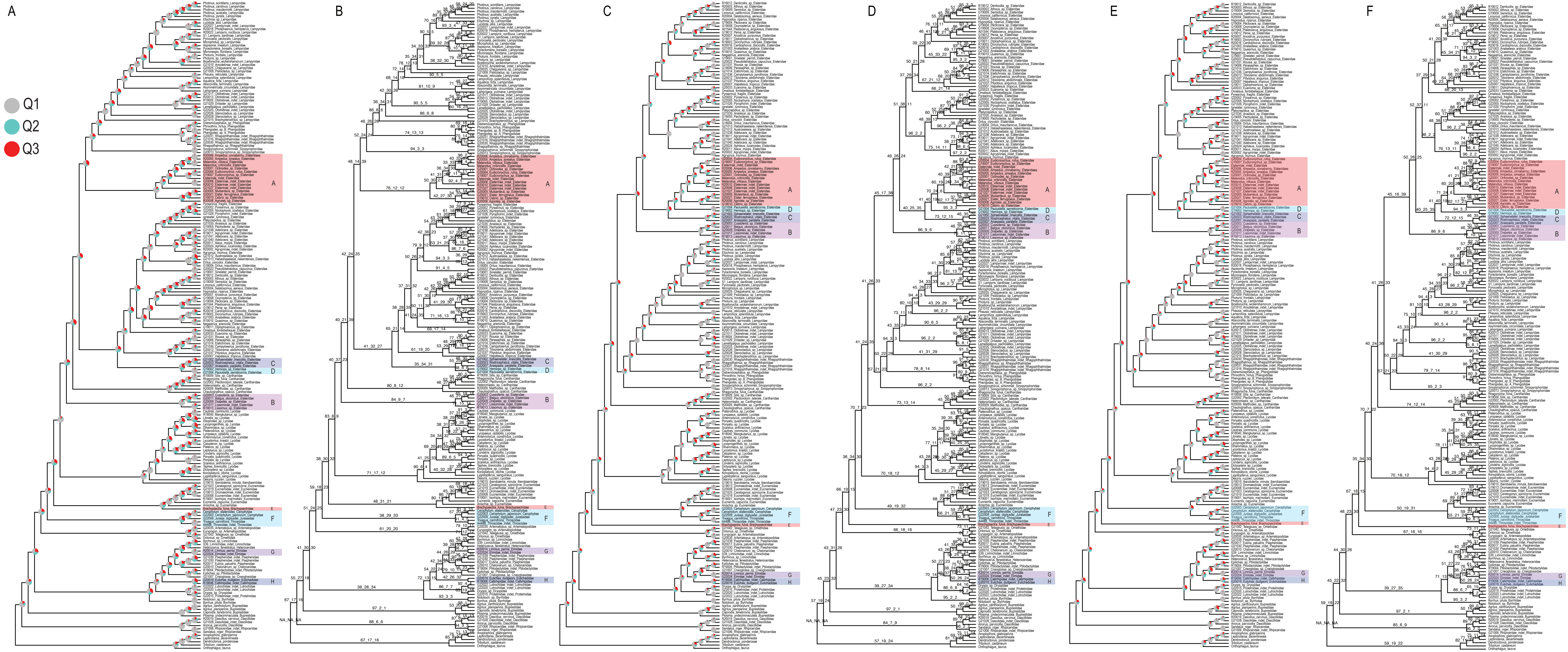

**Supplementary Figure S23.** Coalescent species trees inferred by ASTRAL from individual gene trees. Maximum likelihood gene trees were constructed using IQ-TREE with datasets A), B) A-4189-AA; C), D) B-4189-NT and E), F) K-3583-NT-SymTest. Branches with ultra-fast bootstrap support ≤50 were collapsed prior to coalescent analysis. Branch lengths are in coalescent units. Node support is shown as quartet frequencies (q1: main topology; q2, q3: alternative topologies) displayed as percentages in numerical or pie chart format.

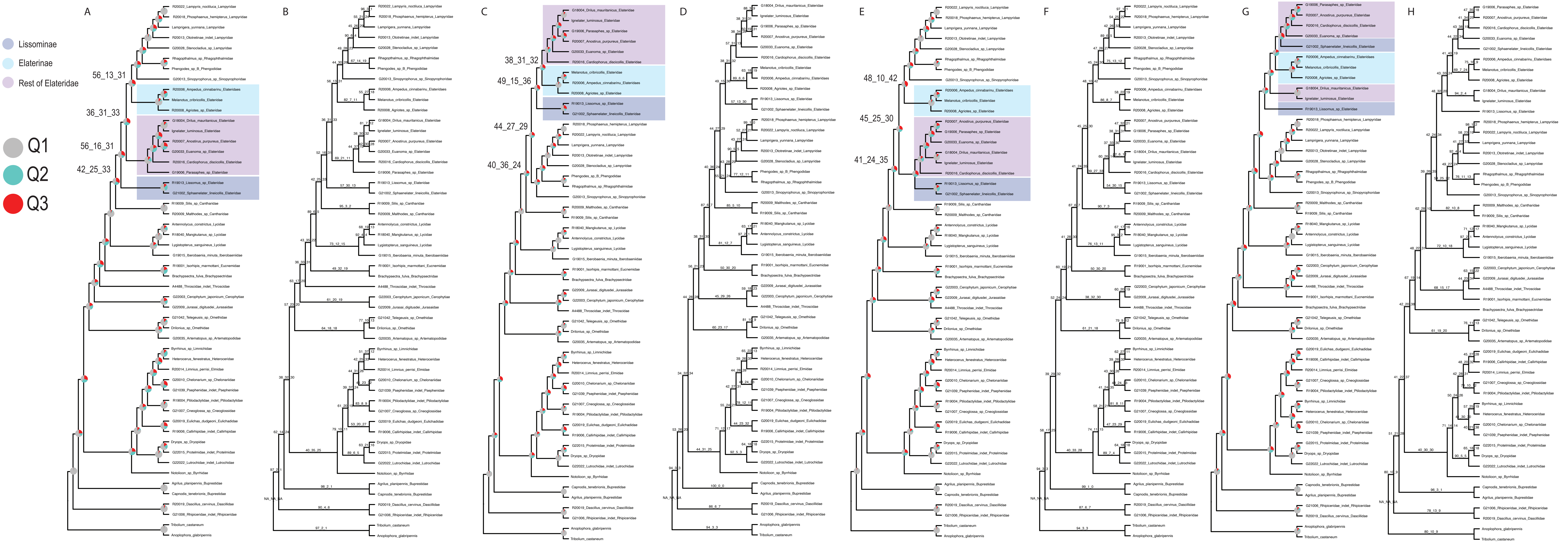

**Supplementary Figure S24.** Coalescent species trees inferred by ASTRAL from subsets of individual gene trees. Maximum likelihood gene trees were constructed using IQ-TREE with A), B) 200 slowest-evolving genes from dataset G-962-AA-decisive; C), D) 200 fastest-evolving genes from dataset G-962-AA-decisive; E), F) all genes from dataset G-962-AA-decisive and G), H) all genes from dataset H-962-NT-decisive. Branches with ultra-fast bootstrap support ≤50 were collapsed prior to coalescent analysis. Branch lengths are in coalescent units. Node support is shown as quartet frequencies (q1: main topology; q2, q3: alternative topologies) displayed as percentages in numerical or pie chart format.

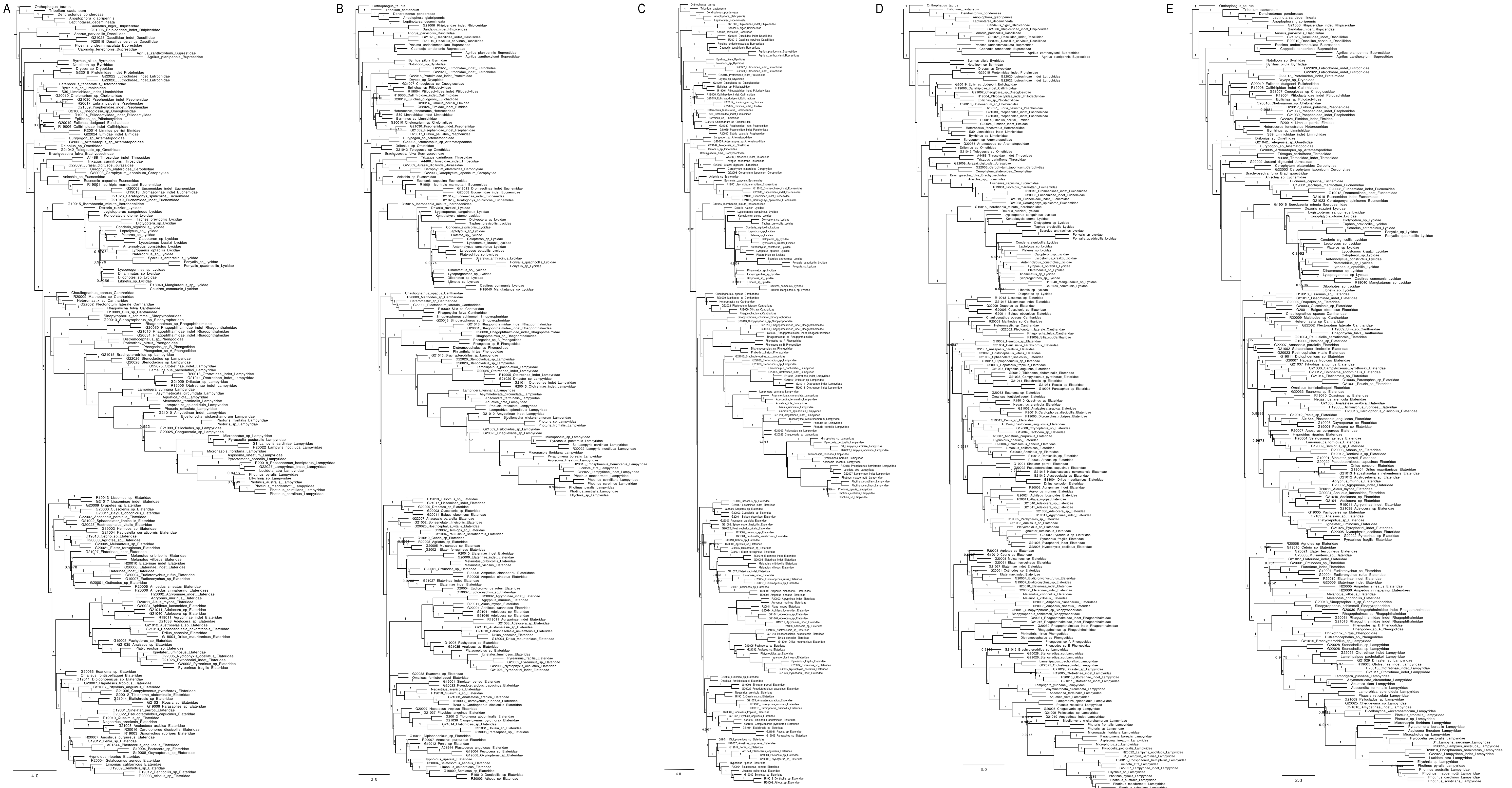

**Supplementary Figure S25.** Coalescent species trees inferred by wASTRAL from individual gene trees. Maximum likelihood gene trees were constructed using IQ-TREE with datasets A) F-4224-NT-raw; B) B-4189-NT; C) K-3583-NT-SymTest; D) E-4224-AA-raw and E) A-4189-AA. Gene trees were used without collapsing poorly supported branches. Branch lengths are shown in coalescent units, and node support values represent local posterior probabilities.

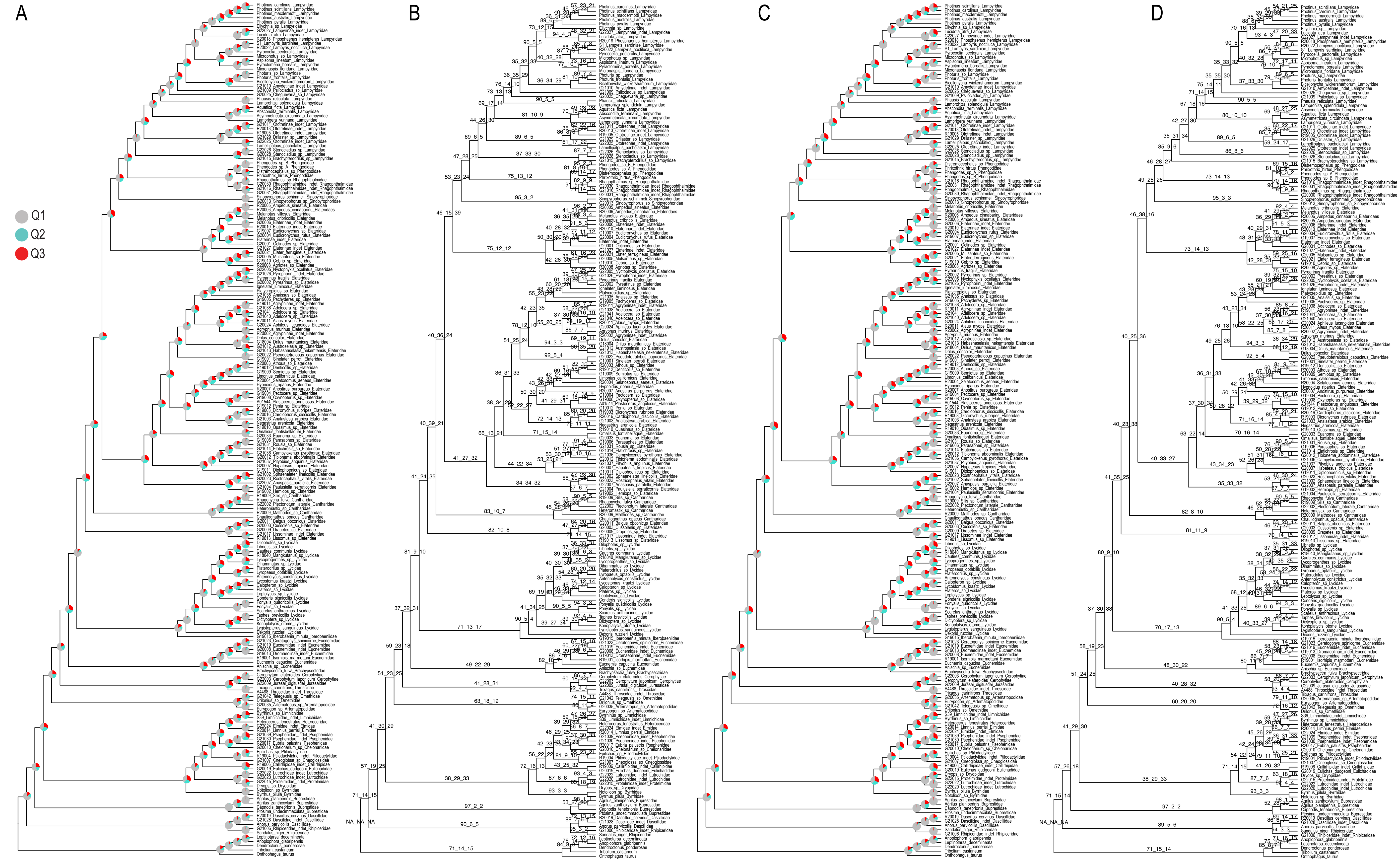

**Supplementary Figure S26.** Coalescent species trees inferred by wASTRAL from individual gene trees. Maximum likelihood gene trees were constructed using IQ-TREE with datasets A), B) E-4224-AA-raw and C), D) A-4189-AA.
Gene trees were used without collapsing poorly supported branches. Branch lengths are in coalescent units. Node support is shown as quartet frequencies (q1: main topology; q2, q3: alternative topologies) displayed as percentages in numerical or pie chart format.

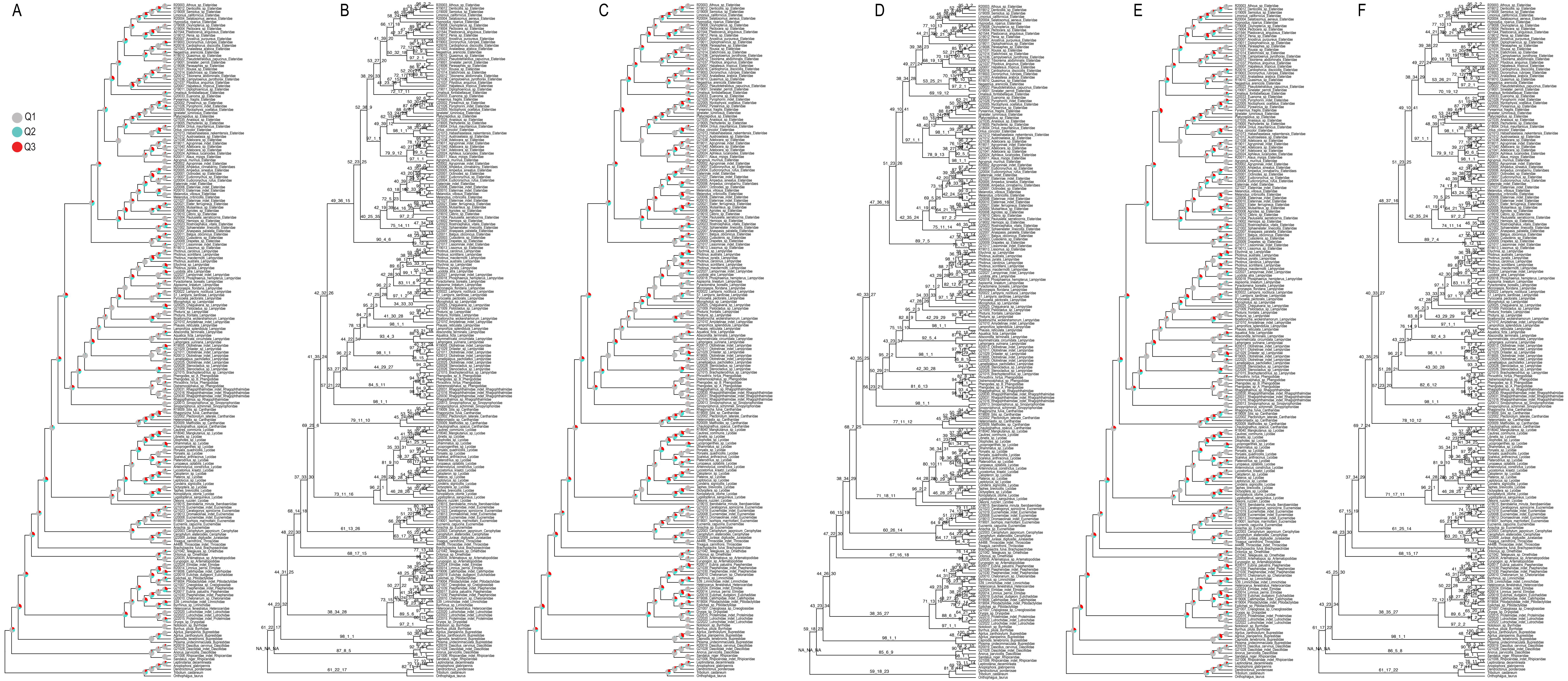

**Supplementary Figure S27.** Coalescent species trees inferred by wASTRAL from individual gene trees. Maximum likelihood gene trees were constructed using IQ-TREE with datasets A), B) F-4224-NT-raw; C), D) B-4189-NT and E), F) K-3583-NT-SymTest. Gene trees were used without collapsing poorly supported branches. Branch lengths are in coalescent units. Node support is shown as quartet frequencies (q1: main topology; q2, q3: alternative topologies) displayed as percentages in numerical or pie chart format.

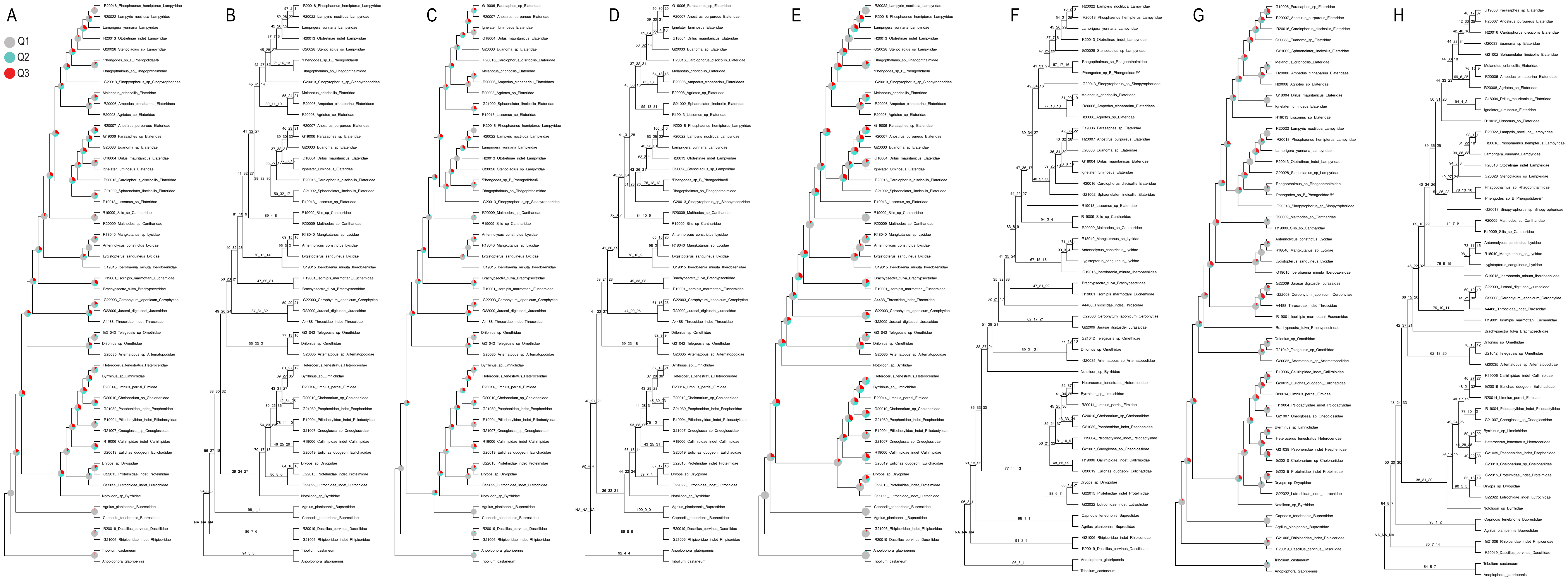

**Supplementary Figure S28.** Coalescent species trees inferred by wASTRAL from subsets of individual gene trees. Maximum likelihood gene trees were constructed using IQ-TREE with A), B) all genes from dataset G-962-AA-decisive; C), D) 200 fastest-evolving genes from dataset G-962-AA-decisive; E), F) 200 slowest-evolving genes from dataset G-962-AA-decisive and G), H) all genes from dataset H-962-NT-decisive. Gene trees were used without collapsing poorly supported branches. Branch lengths are in coalescent units. Node support is shown as quartet frequencies (q1: main topology; q2, q3: alternative topologies) displayed as percentages in numerical or pie chart format.

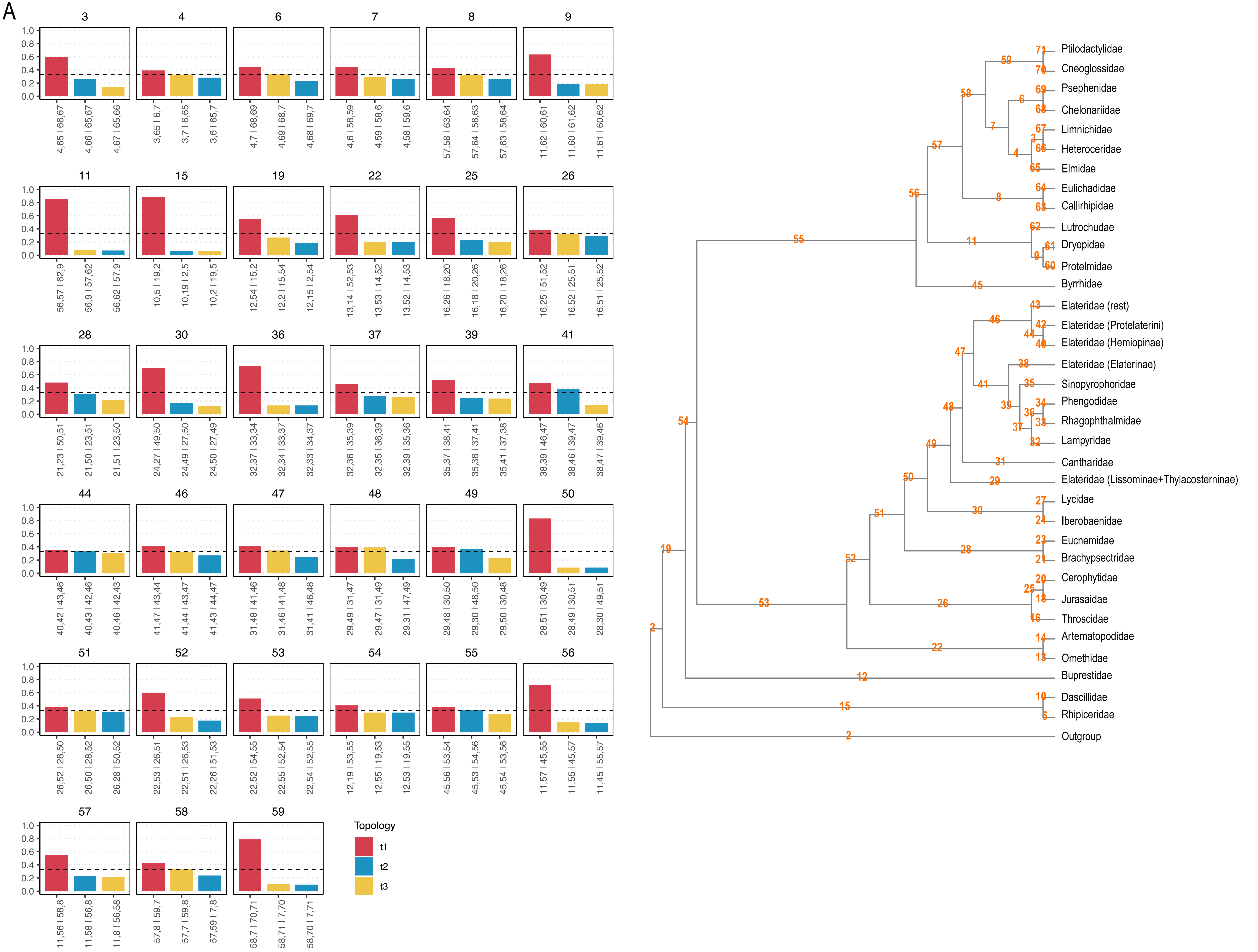

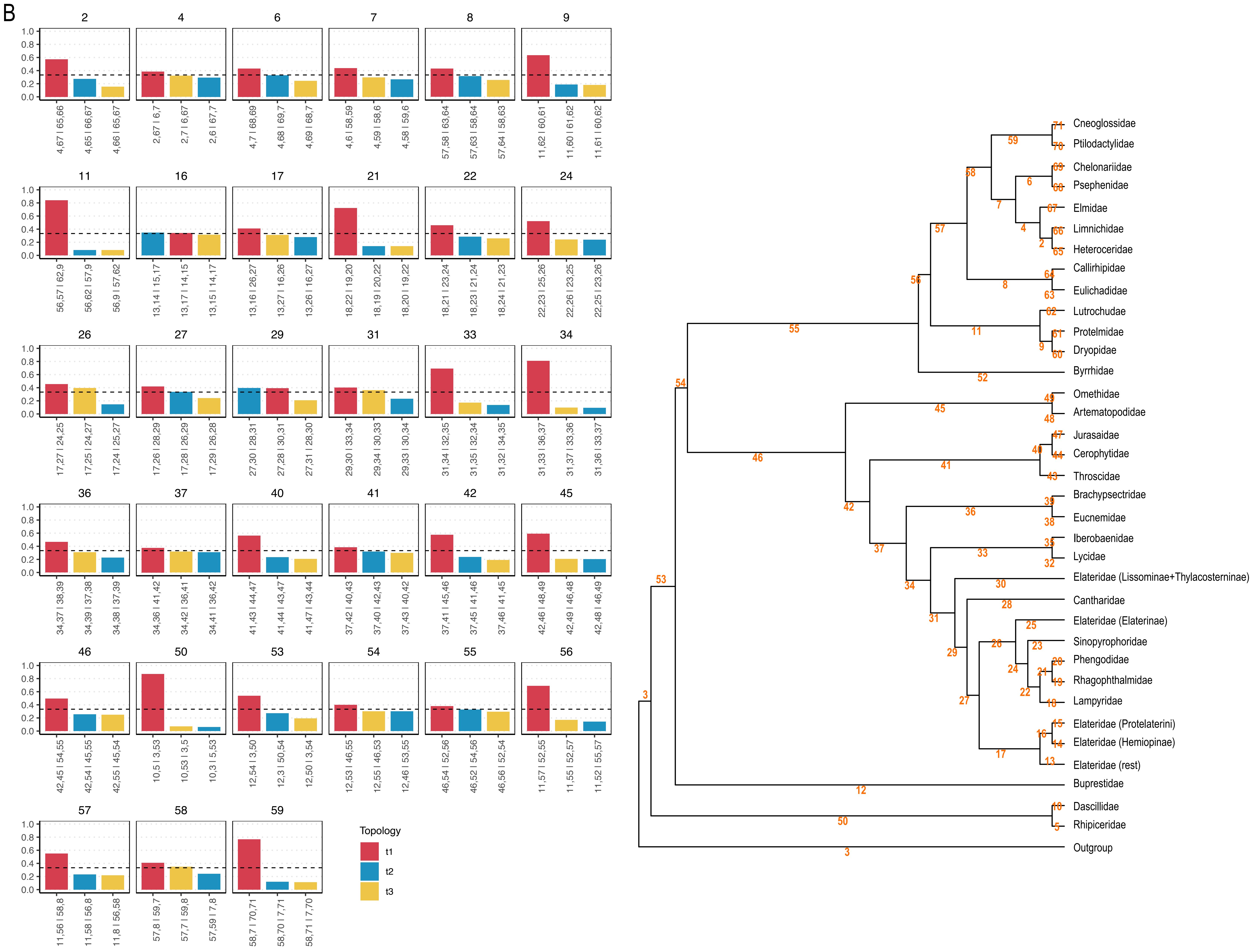

**Supplementary Figure S29.** DiscoVista relative frequency analyses of datasets: A) A-4189-AA branches with ultra-fast bootstrap support ≤50 were collapsed in individual gene trees; B) E-4224-AA-raw gene trees were used without collapsing poorly supported branches.

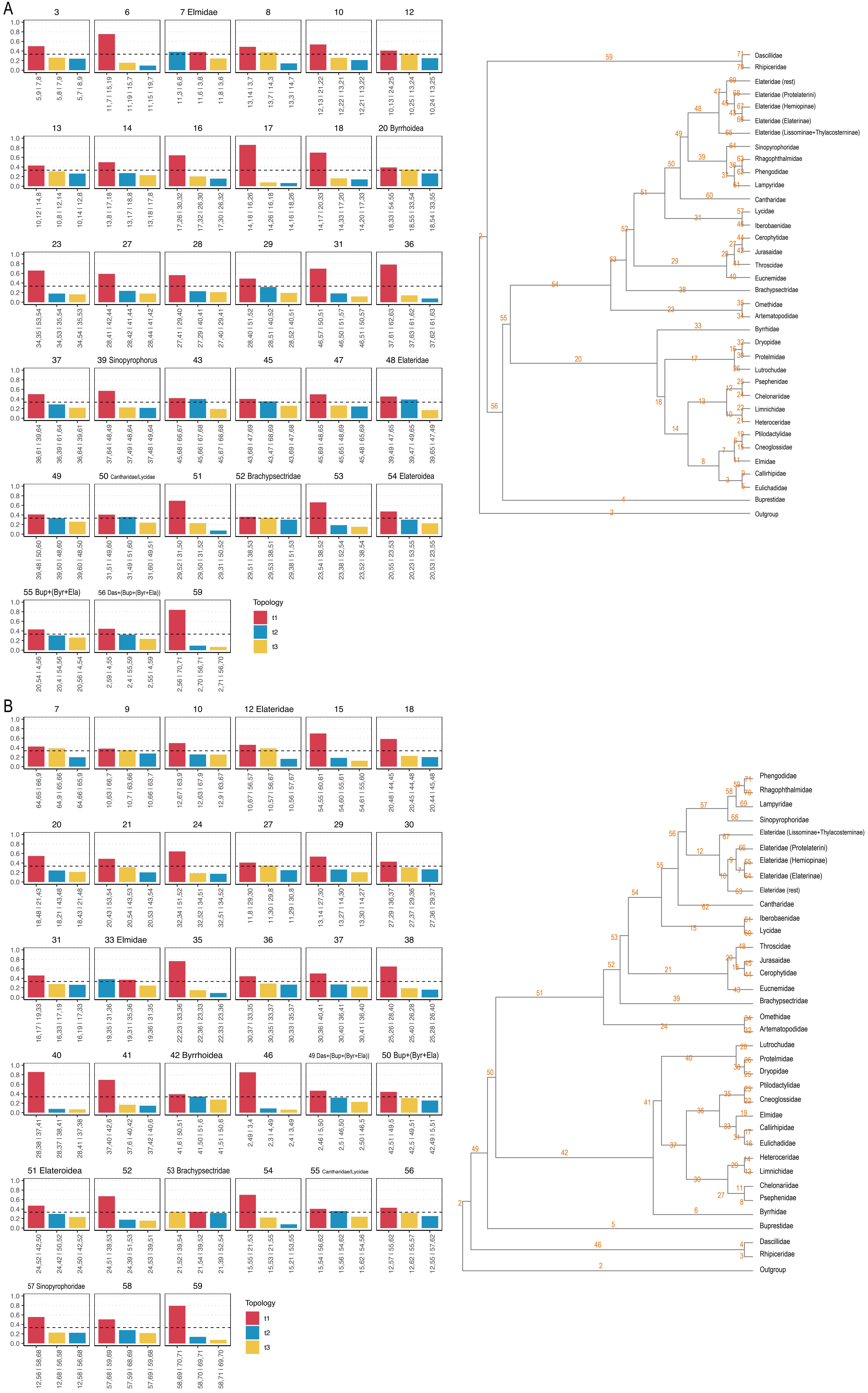

**Supplementary Figure S30.** DiscoVista relative frequency analyses of datasets: A) B-4189-NT branches with ultra-fast bootstrap support ≤50 were collapsed in individual gene trees; B) F-4224-NT-raw gene trees were used without collapsing poorly supported branches.

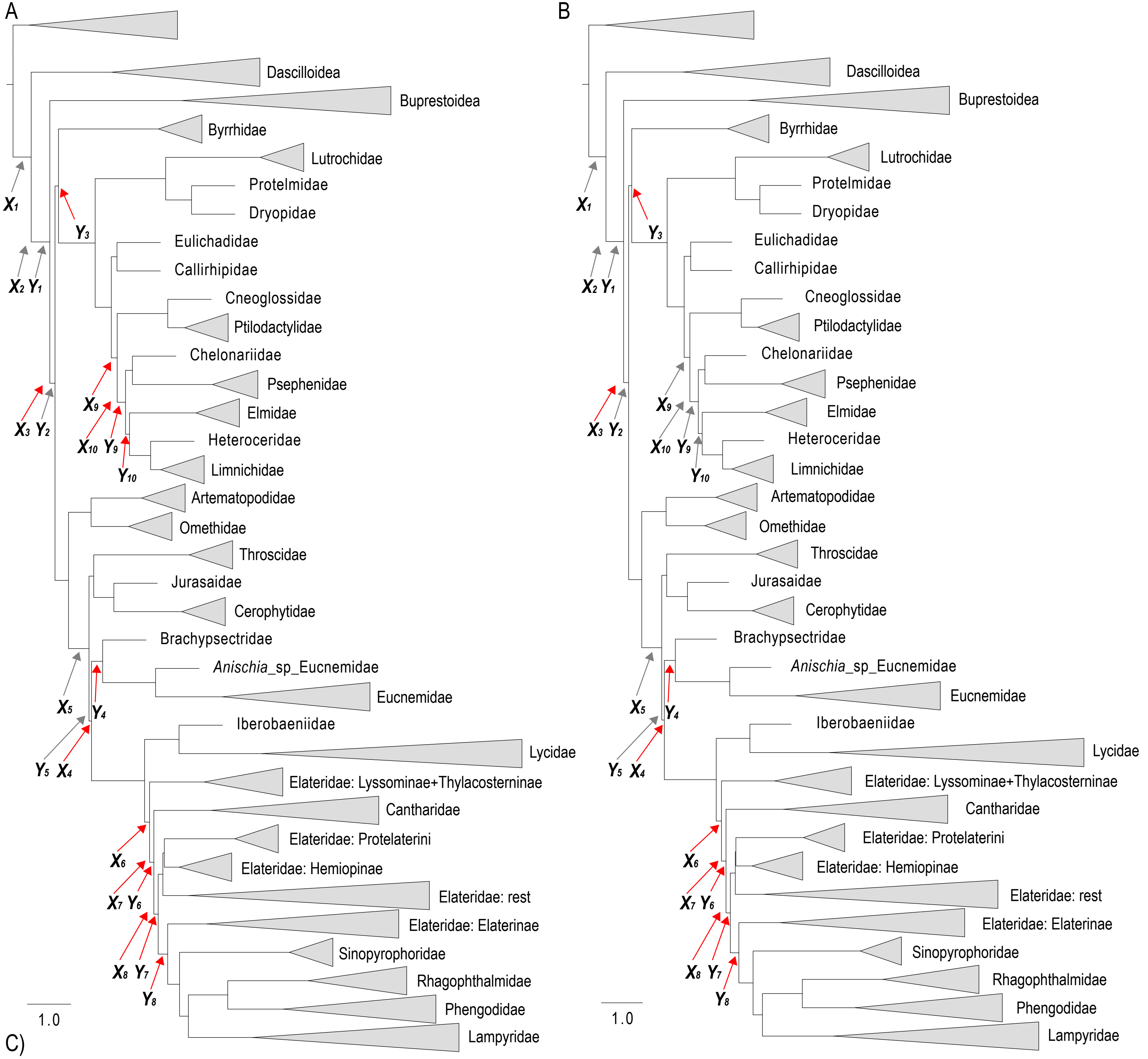

C)

| | Clade 1 | Clade 2 | A-4189-AA x | A-4189-AA a(x) | A-4189-AA y | A-4189-AA Is anomalous y < a(x) | E-4224-AA-raw x | E-4224-AA-raw a(x) | E-4224-AA-raw y | E-4224-AA-raw Is anomalous y < a(x) |
|---|---|---|---|---|---|---|---|---|---|---|
| 1 | Byrrhoidea/Elateroidea+Buprestoidea+Dascilloidea | Byrrhoidea/Elateroidea+Buprestoidea | 0.417423 | -0.110578 | 0.428404 | FALSE | 0.416109 | -0.109866 | 0.427925 | FALSE |
| 2 | Byrrhoidea/Elateroidea+Buprestoidea | Byrrhoidea/Elateroidea | 0.428404 | -0.116411 | 0.119971 | FALSE | 0.427925 | -0.116161 | 0.121589 | FALSE |
| 3 | Byrrhidae/Byrrhoidea+Elateroidea | Byrrhidae/Byrrhoidea | 0.119971 | 0.253500 | 0.078648 | TRUE | 0.121589 | 0.248300 | 0.074830 | TRUE |
| 4 | Brachypsectridae/Eucnemidae+Elateroidea | Brachypsectridae/Eucnemidae | 0.057890 | 0.589261 | 0.256734 | TRUE | 0.062417 | 0.549100 | 0.265690 | TRUE |
| 5 | Eucnemidae/Brachypsectridae+Elateroidea+Jurasaidae/Cerophytidae/Throscidae | Eucnemidae/Brachypsectridae+Elateroidea | 0.470071 | -0.136875 | 0.057890 | FALSE | 0.483896 | -0.143144 | 0.062417 | FALSE |
| 6 | Lissominae/Thylacosterninae+Cantharidae/Elat/Elat/Lampyroid | Lissominae/Thylacosterninae | 0.115435 | 0.268420 | 0.097601 | TRUE | 0.126821 | 0.232408 | 0.109712 | TRUE |
| 7 | Elat+Elat/Lampyroid+Cantharidae | Elat+Elat/Lampyroid | 0.097601 | 0.336999 | 0.099037 | TRUE | 0.109712 | 0.288601 | 0.104042 | TRUE |
| 8 | Lampyroid+Elaterinae+rest of Elateridae | Lampyroid+Elaterinae | 0.099037 | 0.330802 | 0.218358 | TRUE | 0.104042 | 0.310211 | 0.207024 | TRUE |
| 9 | Chelo/Pse+Elmi/Hetero/Limni+Cneoglossidae/Ptilodactylidae | Chelo/Pse+Elmi/Hetero/Limni | 0.135227 | 0.208789 | 0.191433 | TRUE | 0.139070 | 0.198711 | 0.203103 | FALSE |
| 10 | Hetero/Limni+Elmidae+Chelo/Pse | Hetero/Limni+Elmidae | 0.191433 | 0.093038 | 0.091023 | TRUE | 0.203103 | 0.075172 | 0.095363 | FALSE |

**Supplementary Figure S31.** Pairs of short successive internal branches create conditions predicted by anomaly zone theory. wASTRAL species tree topologies for datasets A) A-4189-AA and B) E-4224-AA-raw show internal branch lengths in coalescent units, derived from individual maximum-likelihood gene tree reconstructions by IQTree. Branch length values for all consecutive branch pairs (x and y) are provided in the data table C), with values a(x) computed following Degnan and Rosenberg (2006) Equation 4. Gene tree incongruence is predicted under anomaly zone conditions when y < a(x). Branch pairs satisfying this anomaly zone condition are marked in the topologies by red arrows.

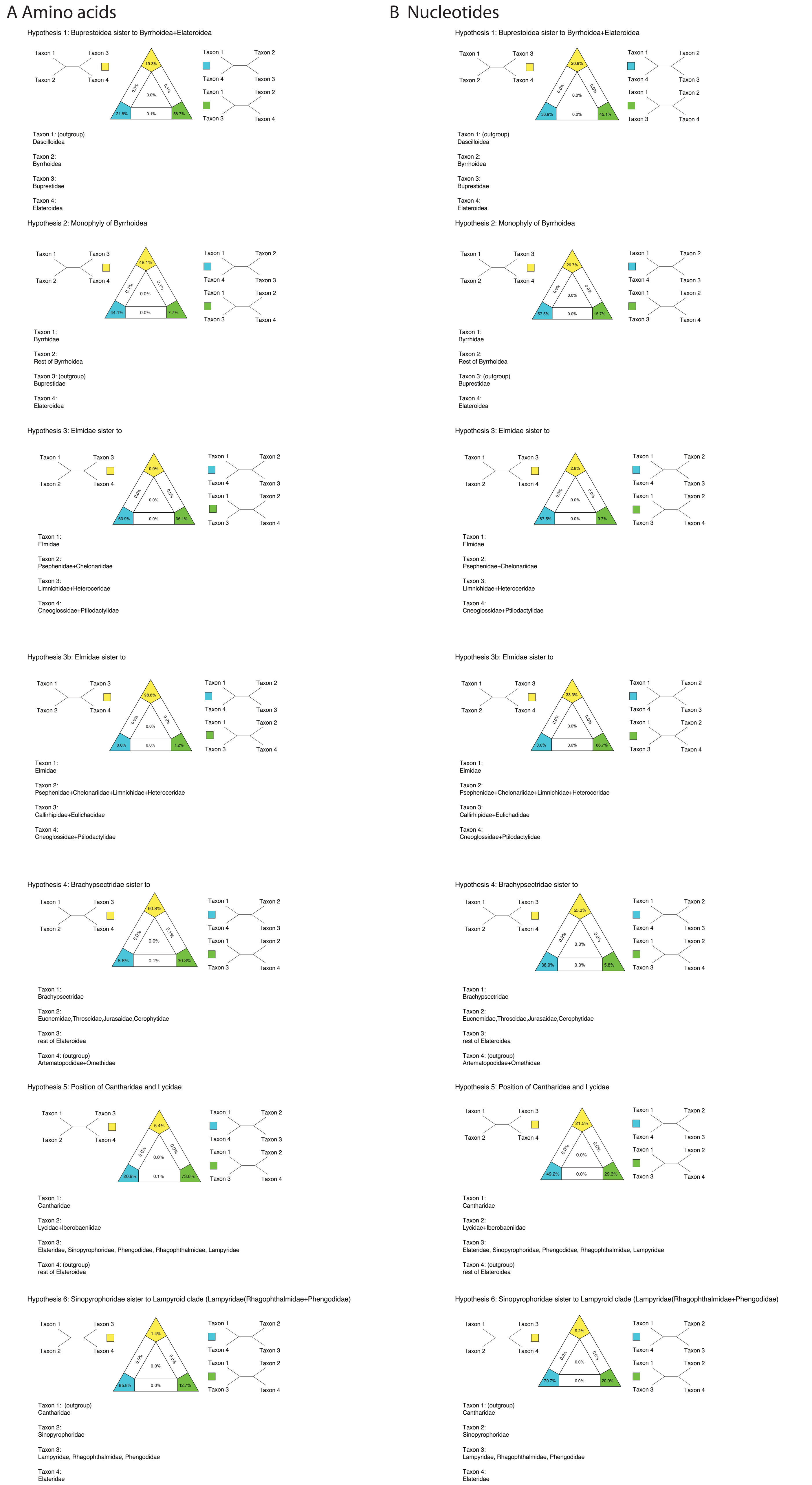

**Supplementary Figure S32.** Results of Four cluster likelihood mapping (FcLM) analyses for a selection of phylogenetic hypotheses applied at the amino acid and nucleotide level for datasets: A) A-4189-AA and B) B-4189-NT.

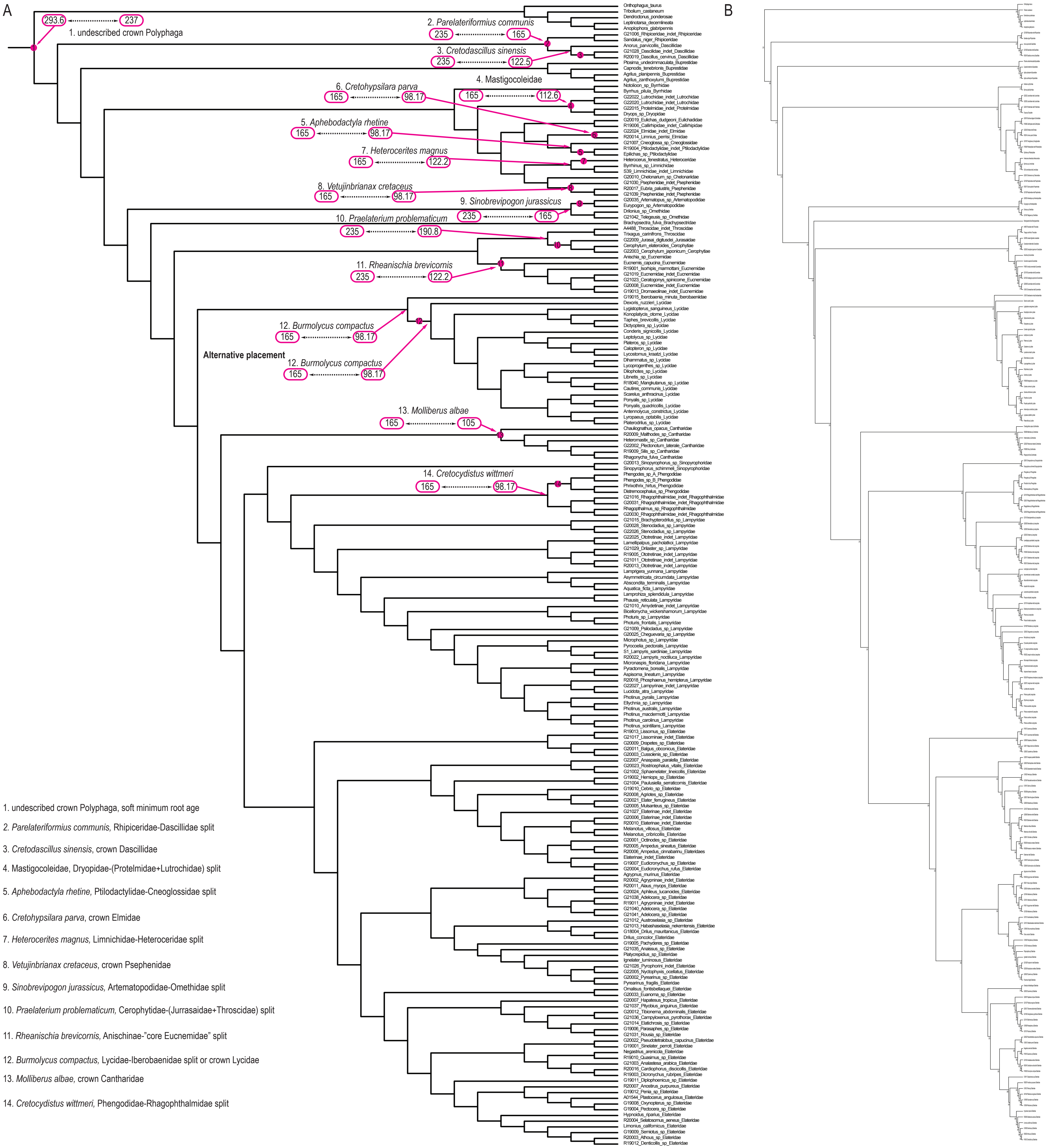

**Supplementary Figure S33.** A) The cladogram with preferred topology (Fig. 5) with fossil calibration points (arrows) used for molecular dating. Pink ovals and dotted lines indicate soft minimum and maximum age constraints, respectively. Numbered fossils (1–14) are listed with their placements; see Supplementary Text S1 for detailed fossil justifications. B) Same tree as (A) but with node numbers displayed.

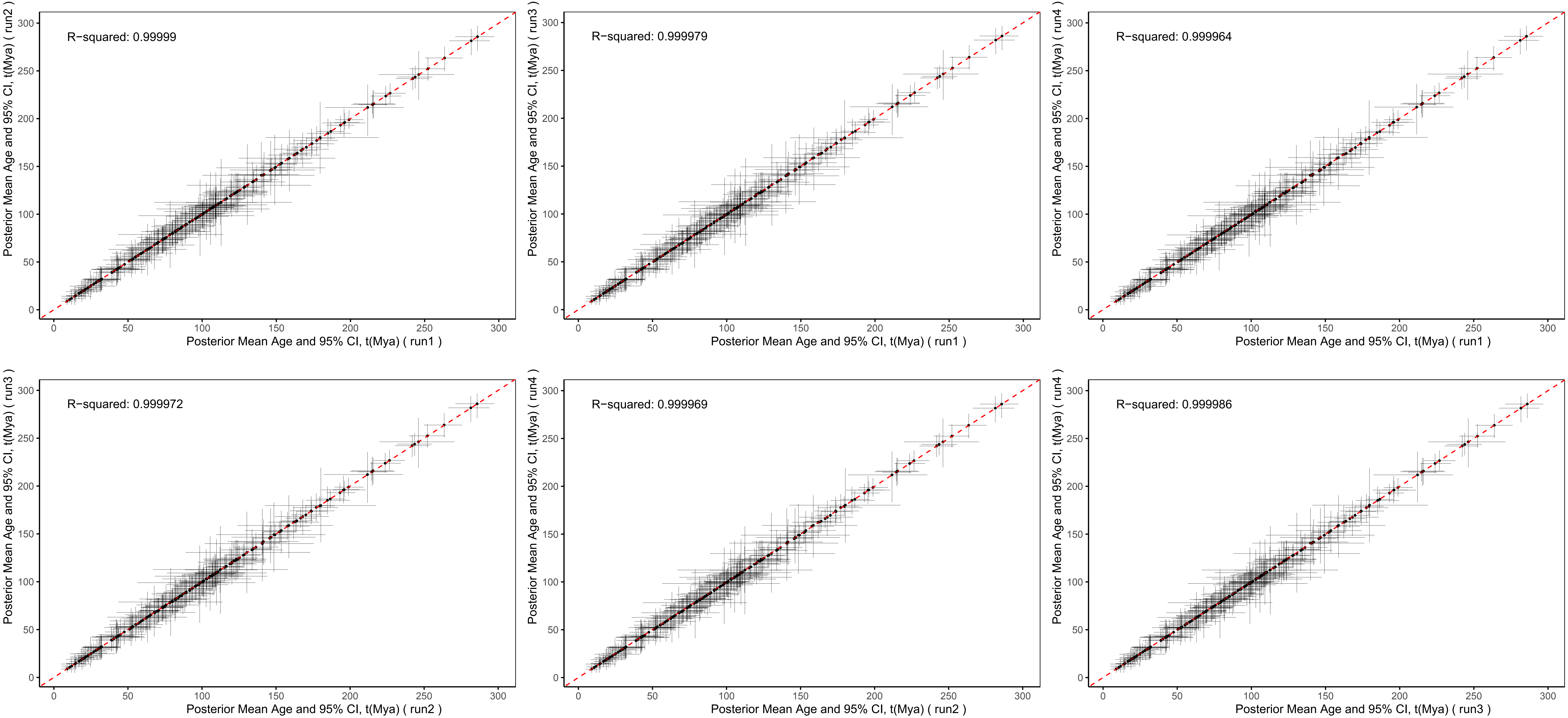

**Supplementary Figure S34.** Convergence assessment of four independent MCMCTREE runs for dataset A at the amino acid level under the autocorrelated rates (AR) clock model. Scatter plots compare posterior mean ages and 95% credibility intervals (CI) between pairs of independent runs. Each plot shows one run on the x-axis versus another on the y-axis. High $R^2$ values (>0.999) across all pairwise comparisons indicate convergence among chains. Mya - million years ago.

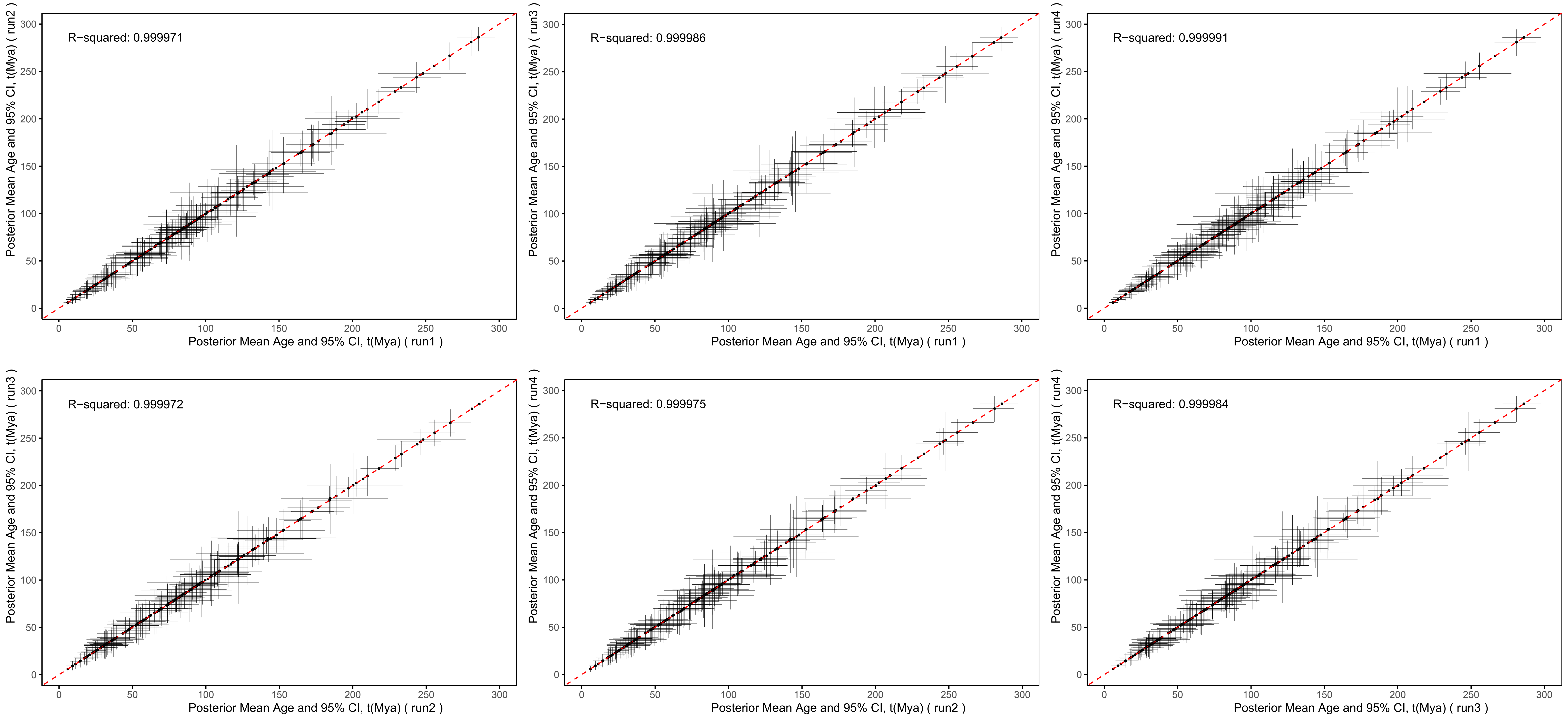

**Supplementary Figure S35.** Convergence assessment of four independent MCMCTREE runs for dataset A at the amino acid level under the independent rates (IR) clock model. Scatter plots compare posterior mean ages and 95% credibility intervals (CI) between pairs of independent runs. Each plot shows one run on the x-axis versus another on the y-axis. High $R^2$ values (>0.999) across all pairwise comparisons indicate convergence among chains. Mya - million years ago.

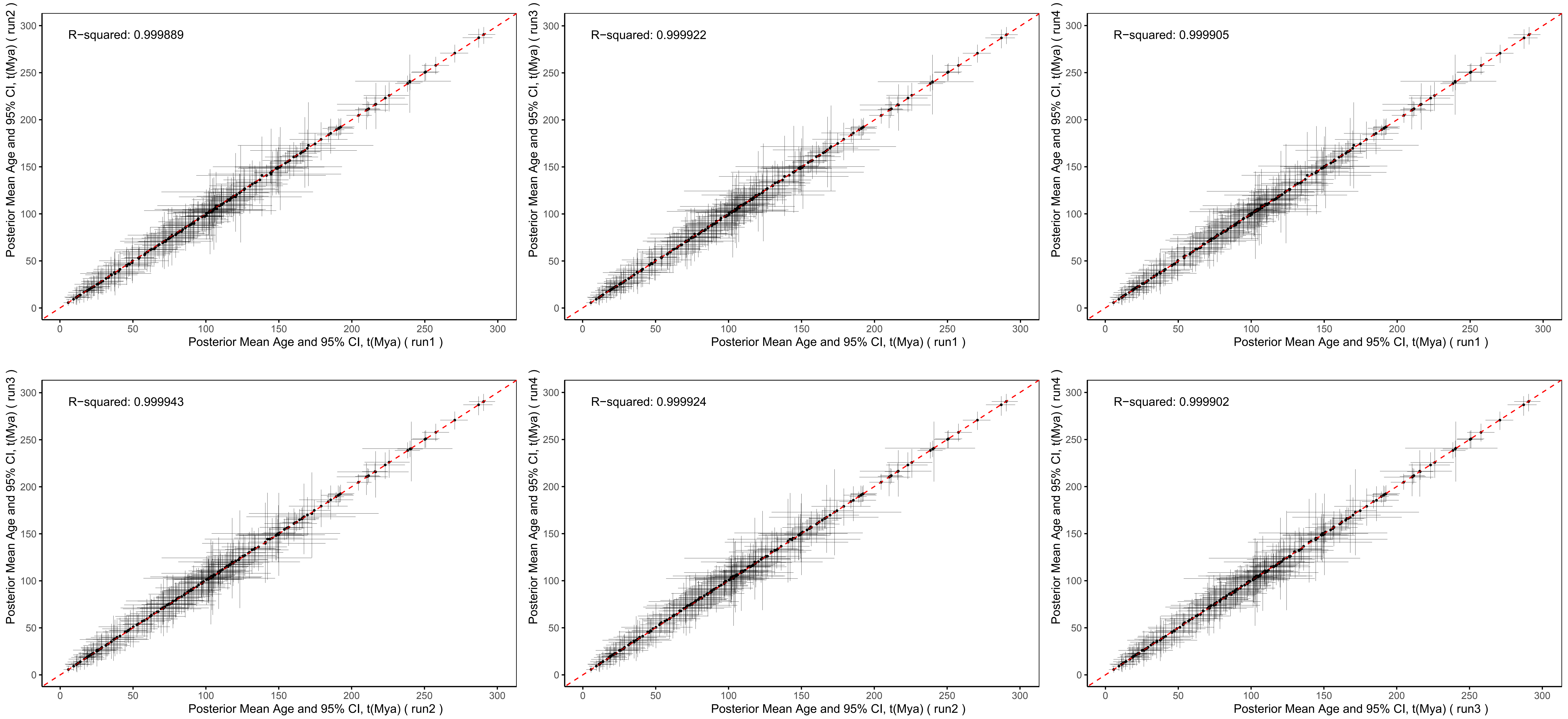

**Supplementary Figure S36.** Convergence assessment of four independent MCMCTREE runs for dataset A at the nucleotide level under the autocorrelated rates (AR) clock model. Scatter plots compare posterior mean ages and 95% credibility intervals (CI) between pairs of independent runs. Each plot shows one run on the x-axis versus another on the y-axis. High $R^2$ values (>0.999) across all pairwise comparisons indicate convergence among chains. Mya - million years ago.

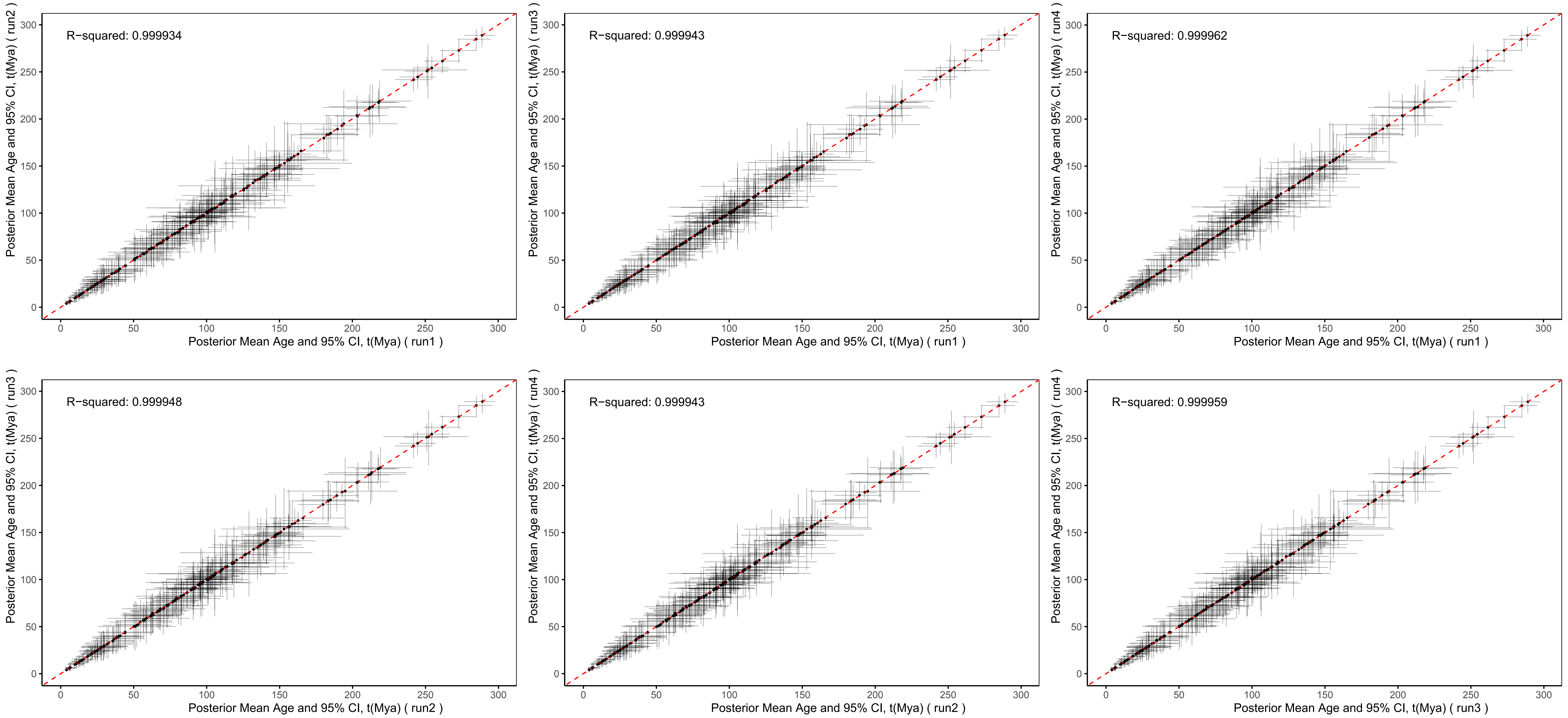

**Supplementary Figure S37.** Convergence assessment of four independent MCMCTREE runs for dataset A at the nucleotide level under the independent rates (IR) clock model. Scatter plots compare posterior mean ages and 95% credibility intervals (CI) between pairs of independent runs. Each plot shows one run on the x-axis versus another on the y-axis. High $R^2$ values (>0.999) across all pairwise comparisons indicate convergence among chains. Mya - million years ago.

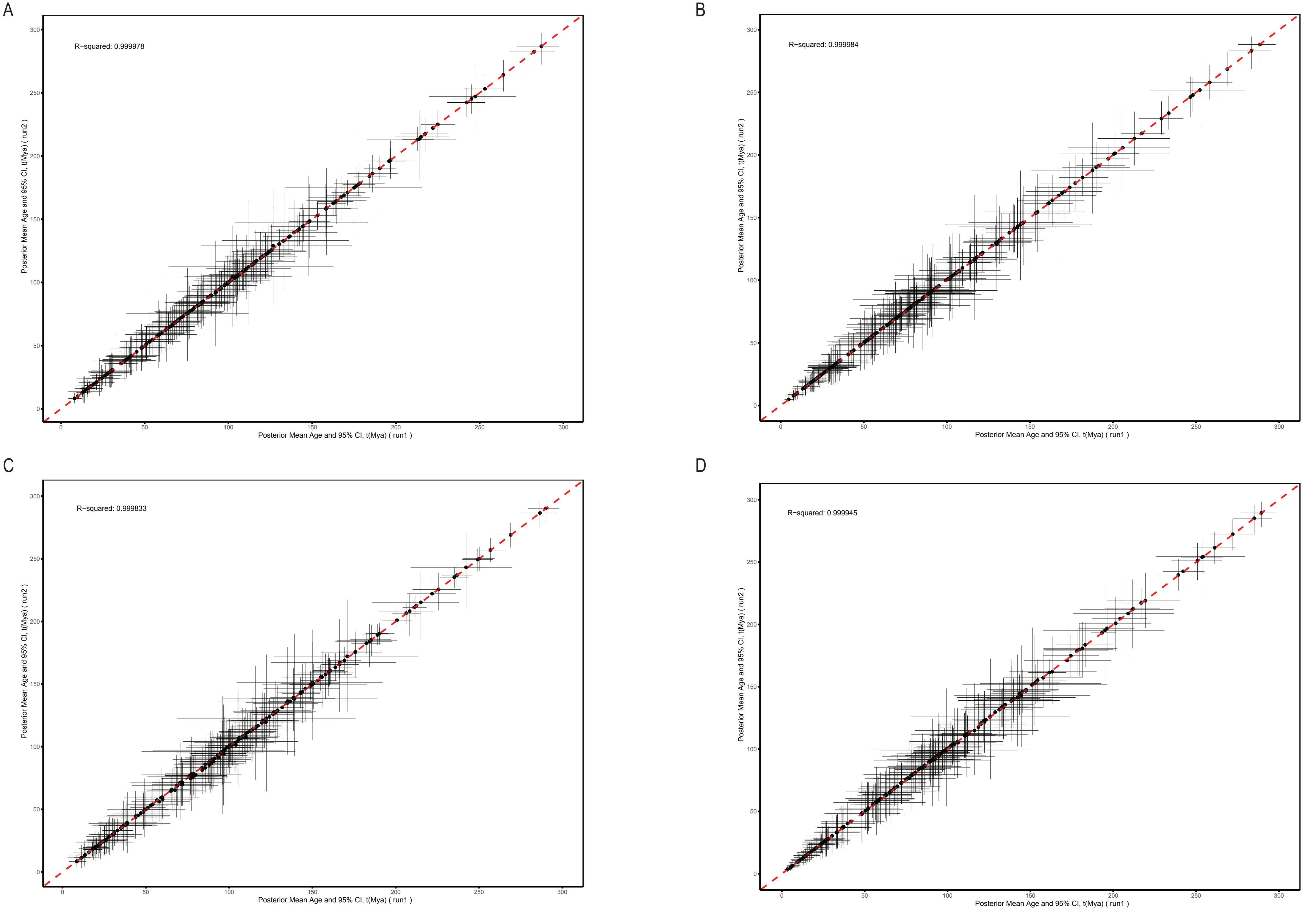

**Supplementary Figure S38.** Convergence assessment of two independent MCMCTREE runs for dataset B under different data types and clock models. Scatter plots compare posterior mean ages and 95% credibility intervals (CI) between the two runs for: A) amino acid sequences under the autocorrelated rates (AR) clock model, B) amino acid sequences under the independent rates (IR) clock model, C) nucleotide sequences under the autocorrelated rates (AR) clock model, and D) nucleotide sequences under the independent rates (IR) clock model. High $R^2$ values (>0.999) across all comparisons indicate convergence between runs. Mya - million years ago.

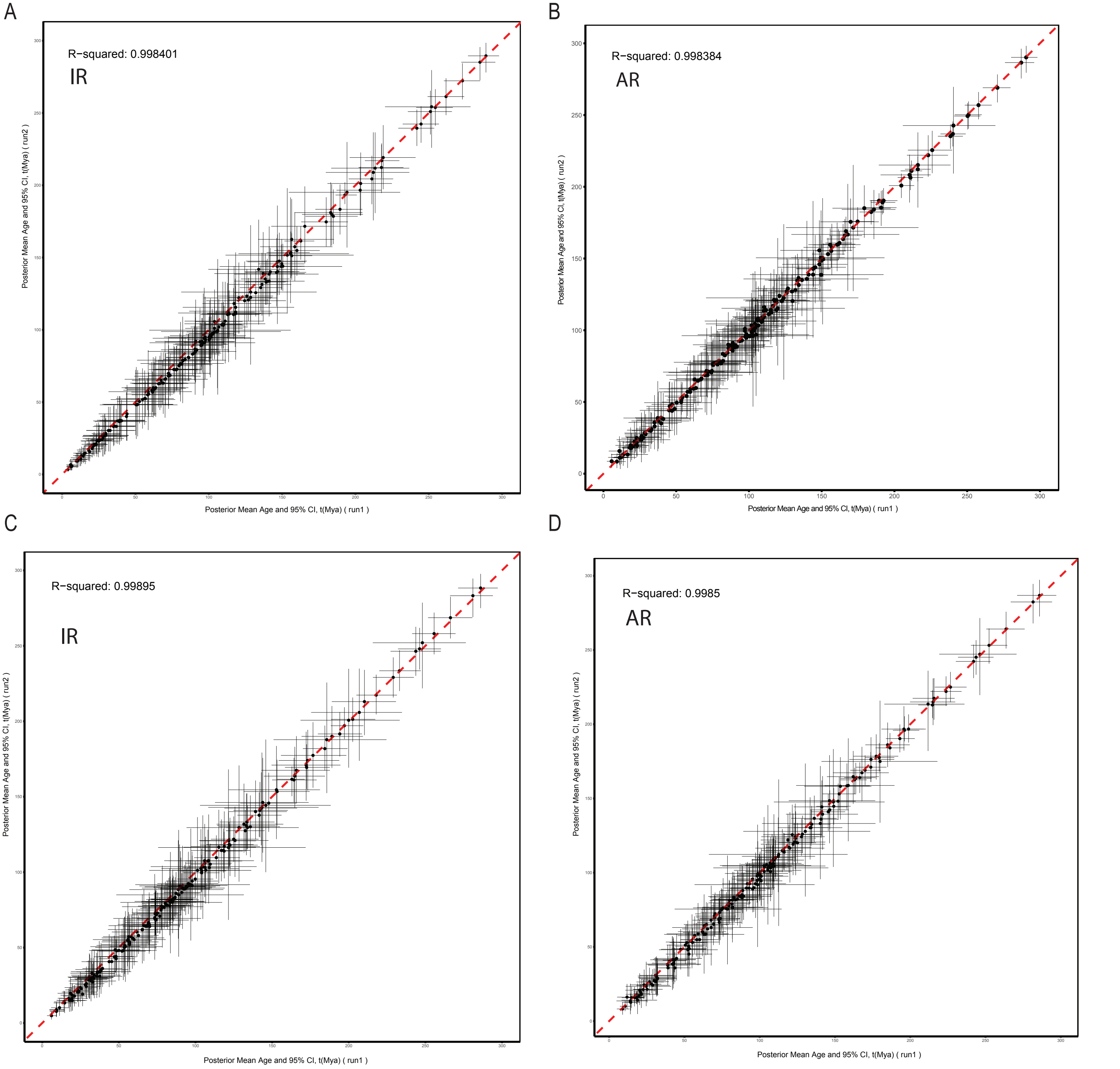

**Supplementary Figure S39.** Convergence assessment comparing merged MCMCTREE runs between datasets A and B under different data types and clock models. Scatter plots compare posterior mean ages and 95% credibility intervals (CI) between dataset A (four merged independent runs-run1) and dataset B (two merged independent runs-run2) for: A) nucleotide sequences under the independent rates (IR) clock model, B) nucleotide sequences under the autocorrelated rates (AR) clock model, C) amino acid sequences under the independent rates (IR) clock model, and D) amino acid sequences under the autocorrelated rates (AR) clock model. High $R^2$ values (>0.998) across all comparisons indicate strong agreement in age estimates between the two datasets despite differences in data composition and number of runs.

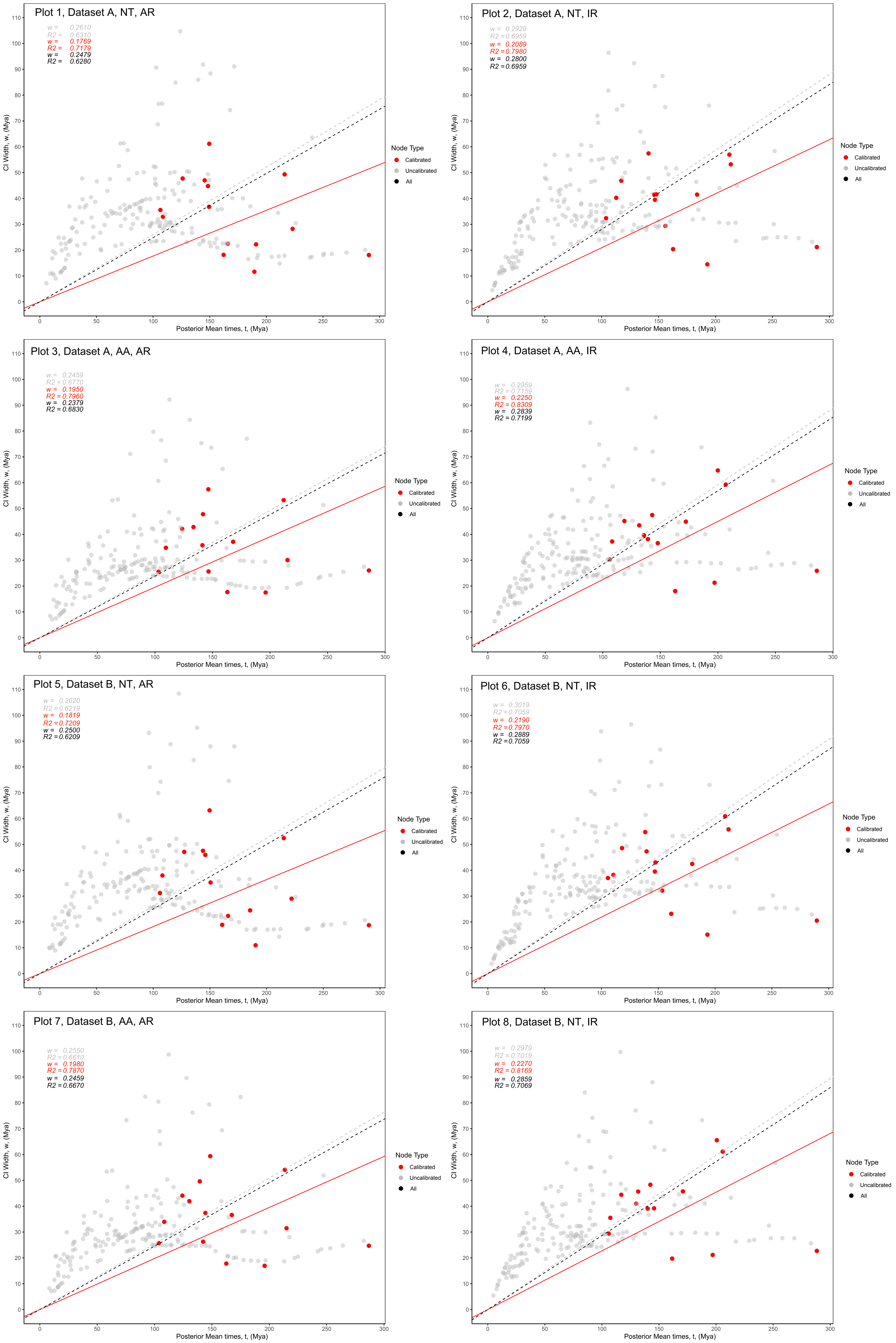

**Supplementary Figure S40.** Infinite-sites plots. Posterior CI width is plotted against mean posterior divergence times, dataset A and B on both NT = nucleotides and AA = amino acid level under AR = autocorrelated rates and IR = independent rates clock models. Red dots represent nodes with fossil calibrations, grey dots represent the rest of nodes. Solid red line: regression through the origin fitted to the red dots. Grey dashed line: regression through the origin fitted to the grey dots. Black dashed line: regression through the origin fitted to all dots.

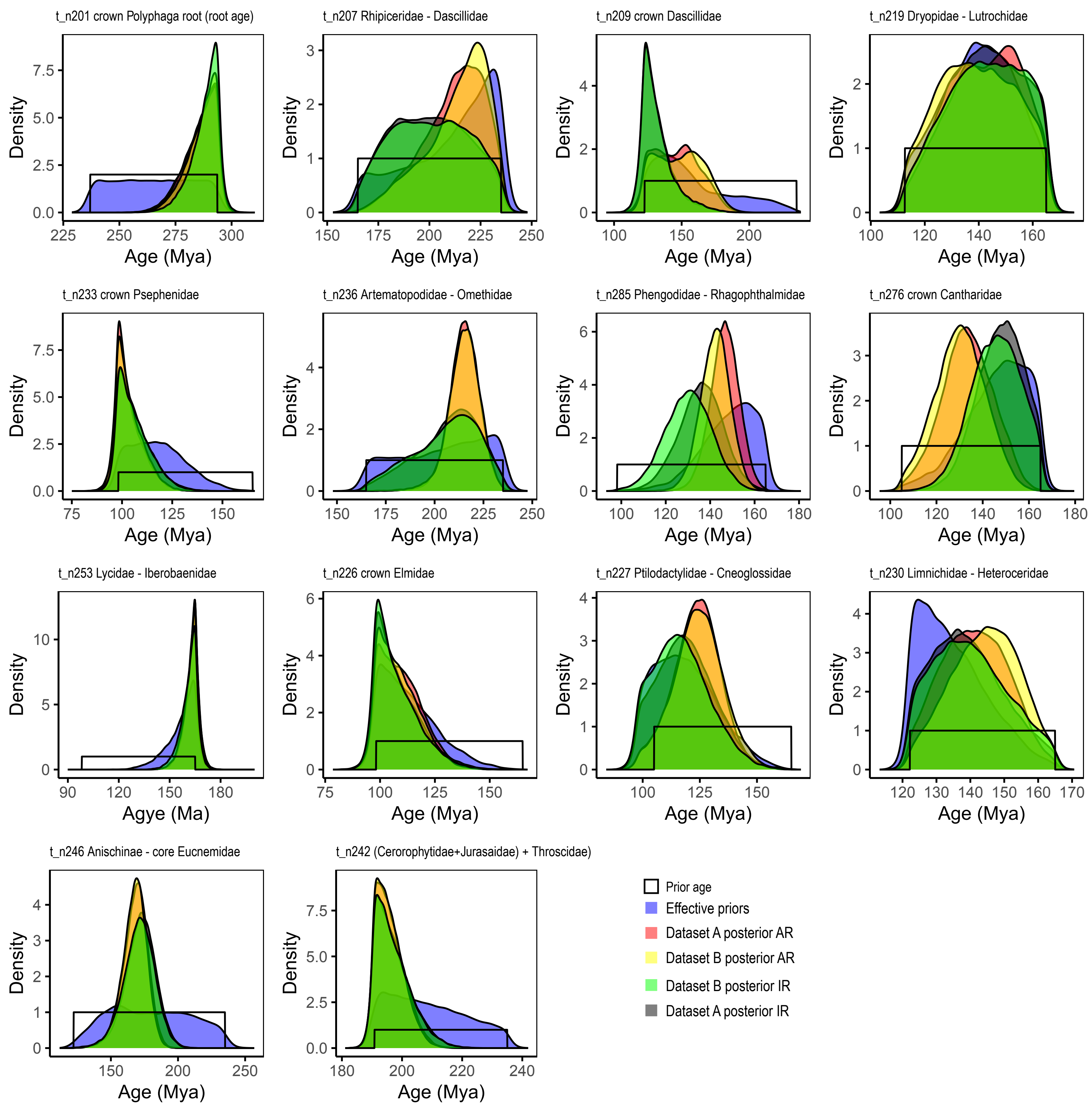

**Supplementary Figure S41.** Density plots of age estimates of calibrated nodes across all time tree analyses performed. Datasets A and B on amino acid level, AR = autocorrelated rates and IR = independent rates clock models. Color legend: solid black line = prior age; blue = effective priors; red = posterior Dataset A, autocorrelated rates; yellow = posterior Dataset B, autocorrelated rates; green = posterior Dataset B, independent rates; grey = posterior Dataset A, independent rates.

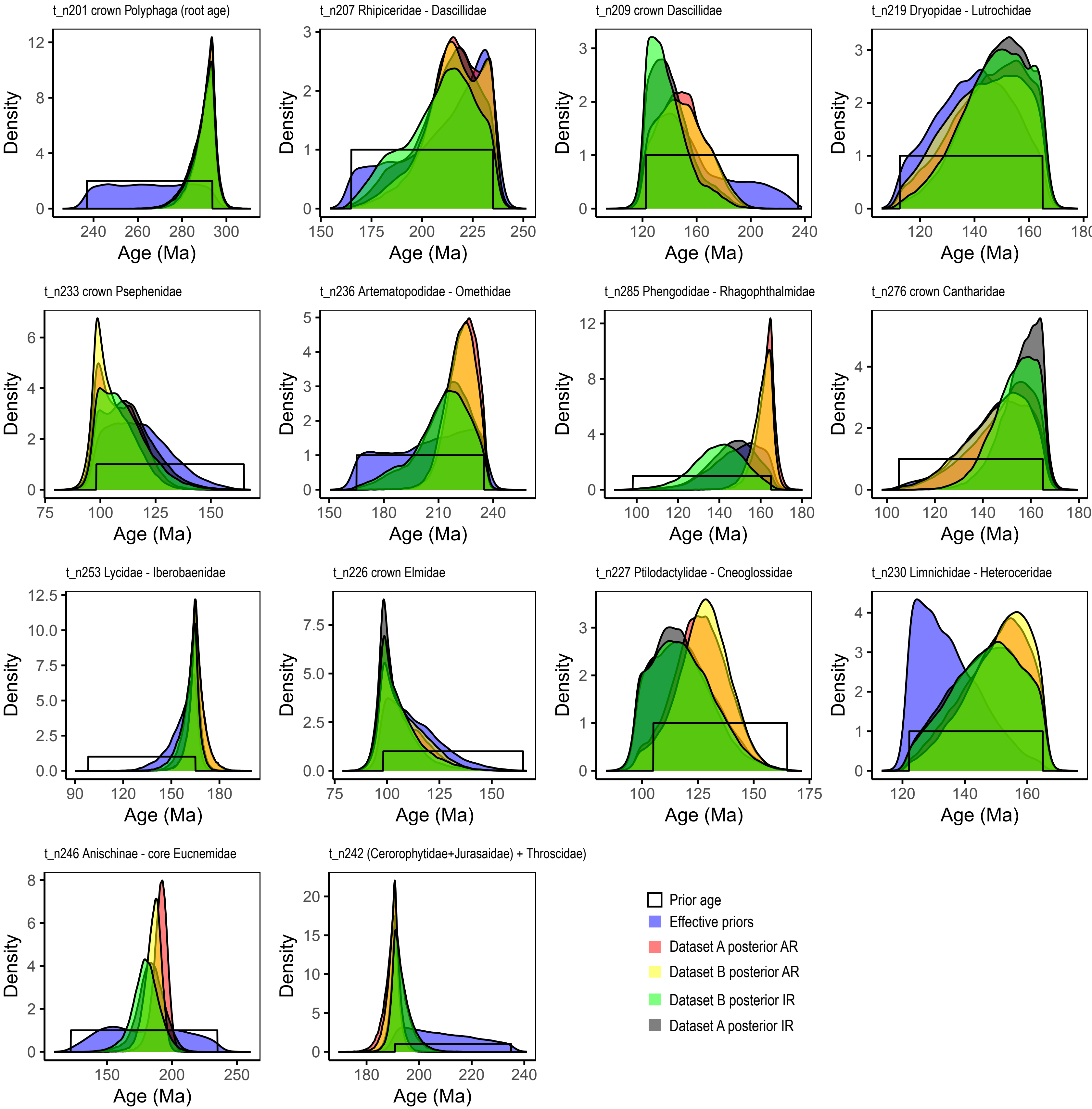

**Supplementary Figure S42.** Density plots of age estimates of calibrated nodes across all time tree analyses performed. Datasets A and B on nucleotide level, AR = autocorrelated rates and IR = independent rates clock models. Color legend: solid black line = prior age; blue = effective priors; red = posterior Dataset A, autocorrelated rates; yellow = posterior Dataset B, autocorrelated rates; green = posterior Dataset B, independent rates; grey = posterior Dataset A, independent rates.

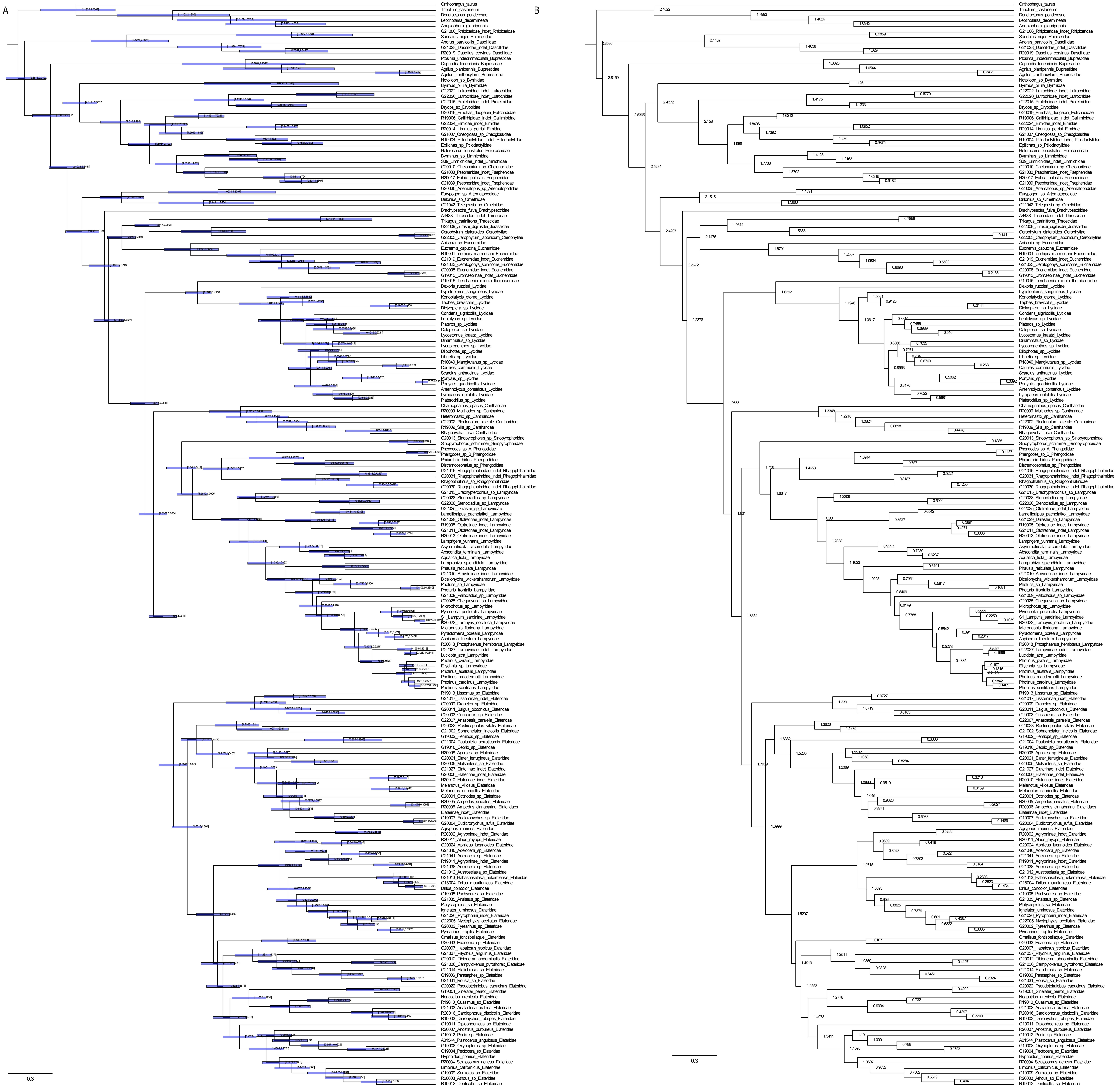

**Supplementary Figure S43.** Divergence time estimation results. Preferred topology analyzed with Dataset A at the amino acid level under autocorrelated rates. A) Blue horizontal bars represent 95% confidence intervals with lower and upper bound ages labeled at bar endpoints. B) Mean node ages given in units of 100 Mya (e.g., 1.2511 = 125.11 million years ago).

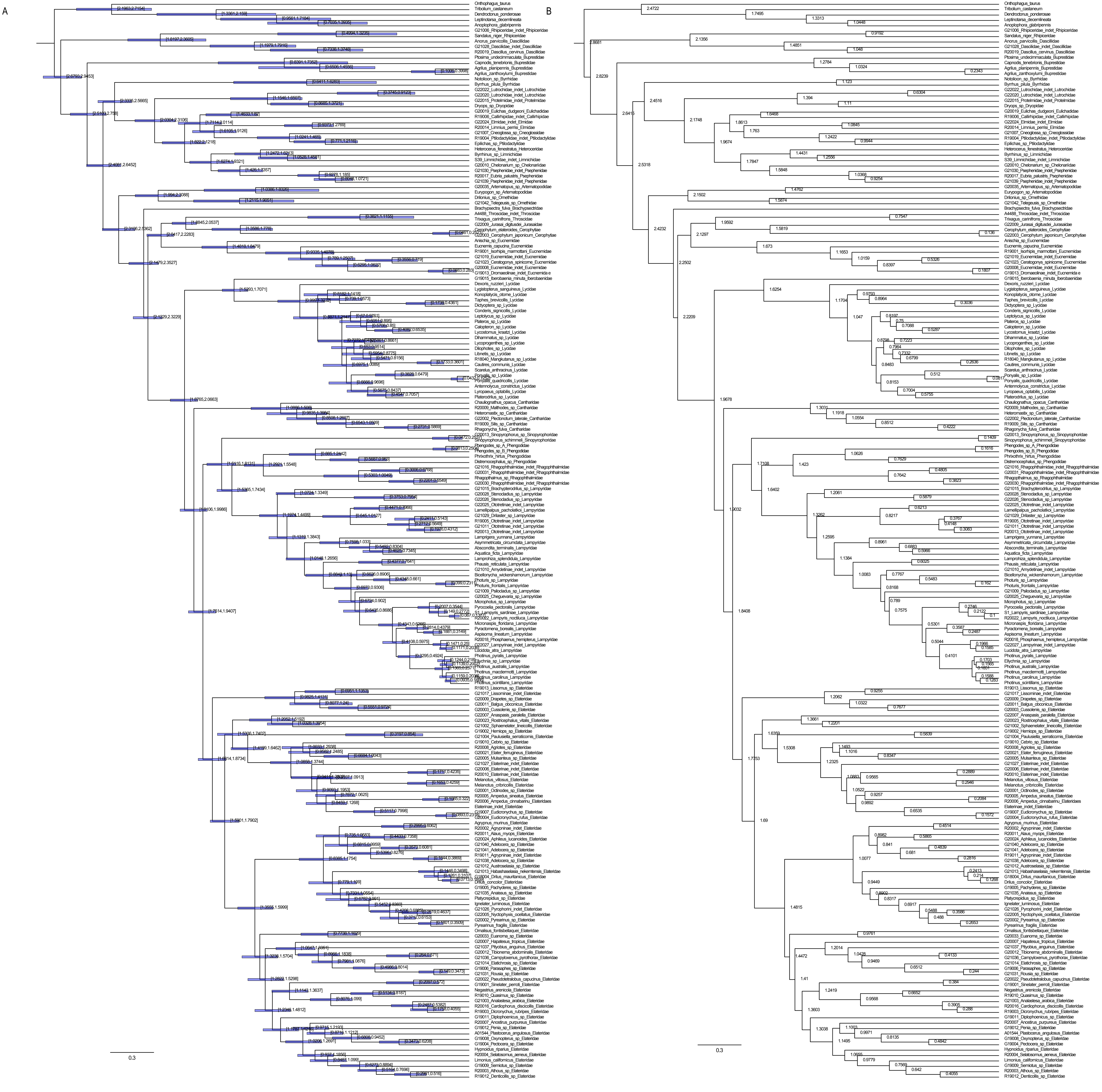

**Supplementary Figure S44.** Divergence time estimation results. Preferred topology analyzed with Dataset B at the amino acid level under autocorrelated rates. A) Blue horizontal bars represent 95% confidence intervals with lower and upper bound ages labeled at bar endpoints. B) Mean node ages given in units of 100 Mya (e.g., 1.2511 = 125.11 million years ago).

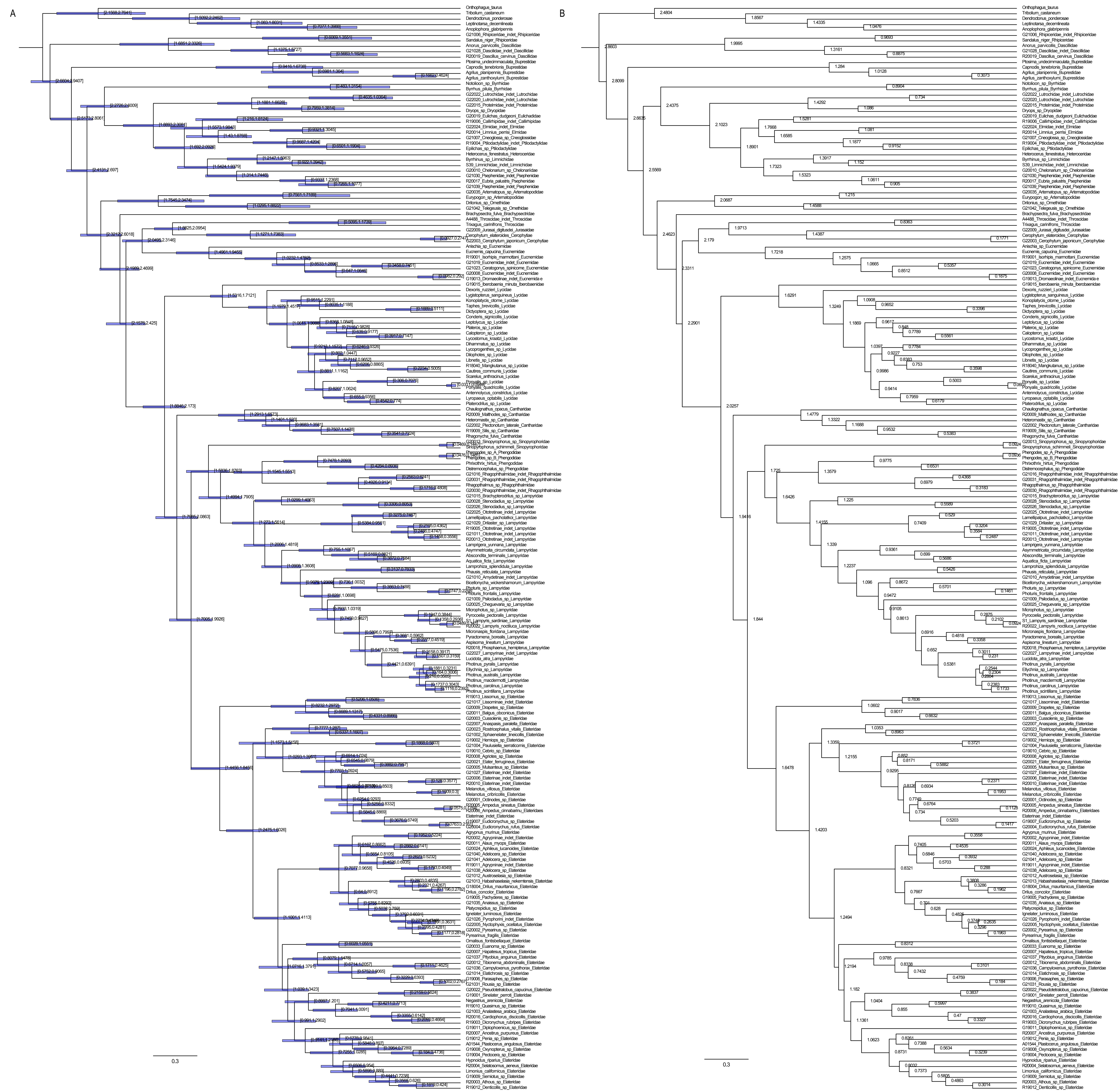

**Supplementary Figure S45.** Divergence time estimation results. Preferred topology analyzed with Dataset A at the amino acid level under independent rates. A) Blue horizontal bars represent 95% confidence intervals with lower and upper bound ages labeled at bar endpoints. B) Mean node ages given in units of 100 Mya (e.g., 1.2511 = 125.11 million years ago).

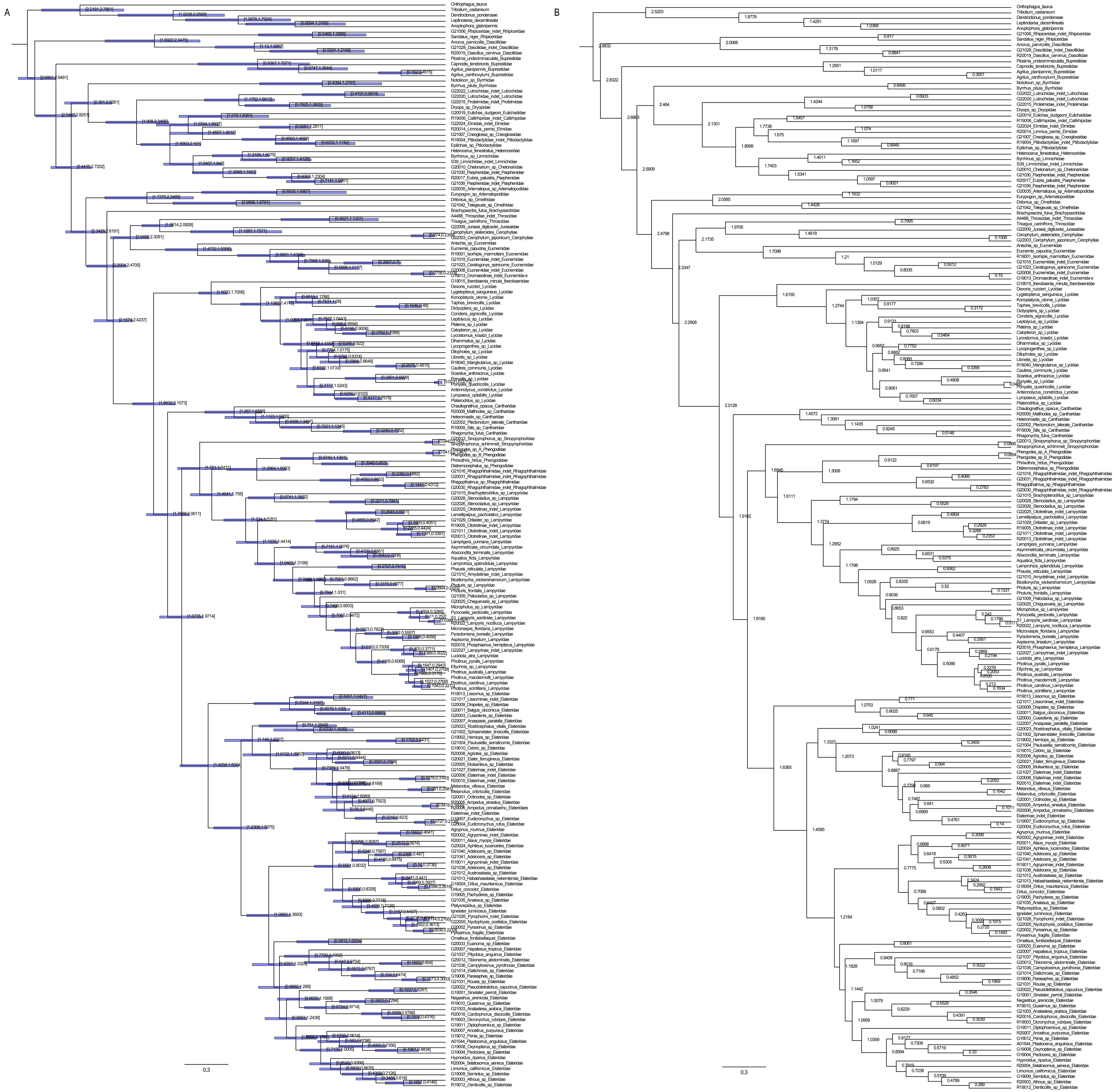

**Supplementary Figure S46.** Divergence time estimation results. Preferred topology analyzed with Dataset B at the amino acid level under independent rates. A) Blue horizontal bars represent 95% confidence intervals with lower and upper bound ages labeled at bar endpoints. B) Mean node ages given in units of 100 Mya (e.g., 1.2511 = 125.11 million years ago).

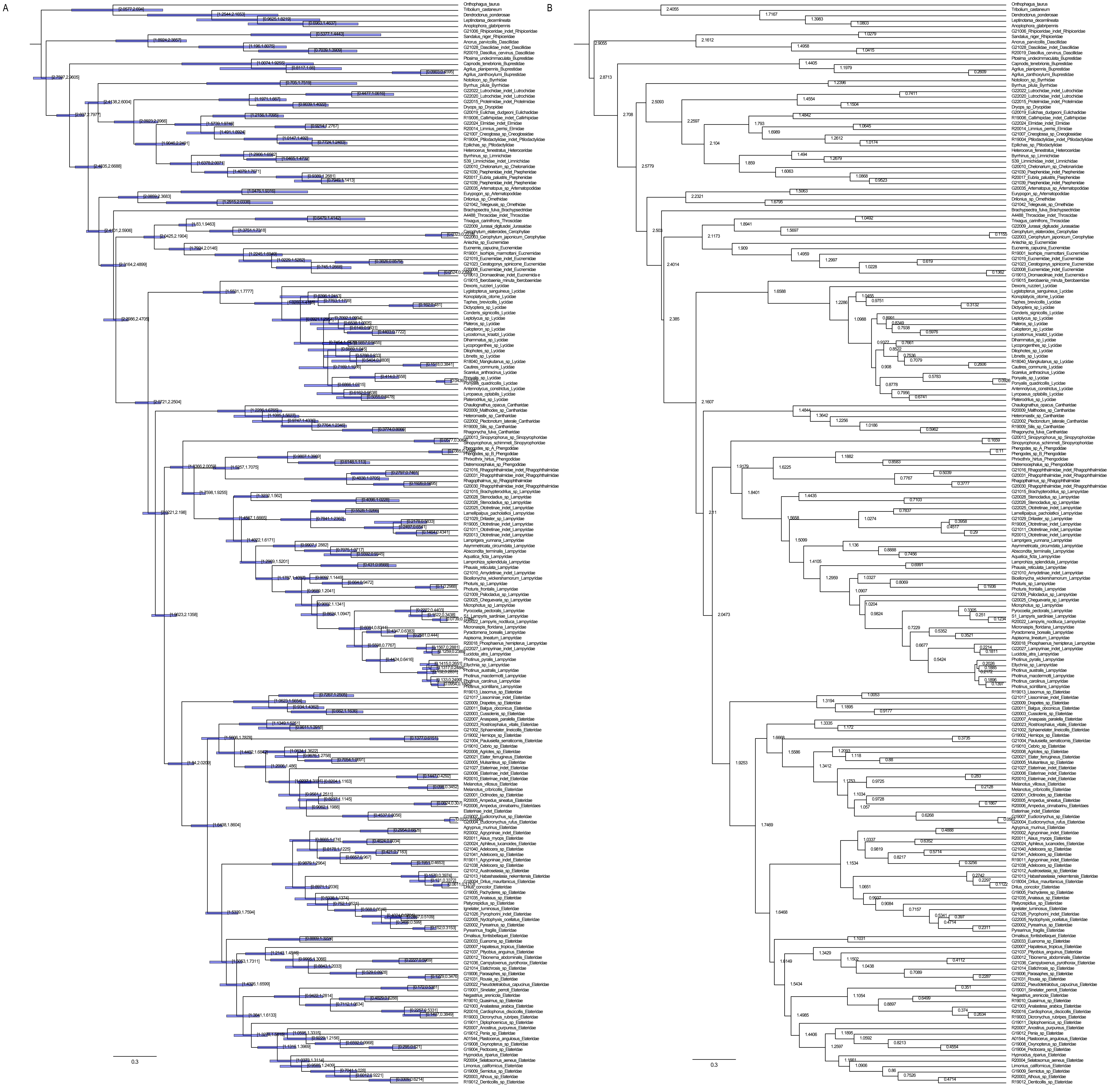

**Supplementary Figure S47.** Divergence time estimation results. Preferred topology analyzed with Dataset A at the nucleotide level under autocorrelated rates. A) Blue horizontal bars represent 95% confidence intervals with lower and upper bound ages labeled at bar endpoints. B) Mean node ages given in units of 100 Mya (e.g., 1.2511 = 125.11 million years ago).

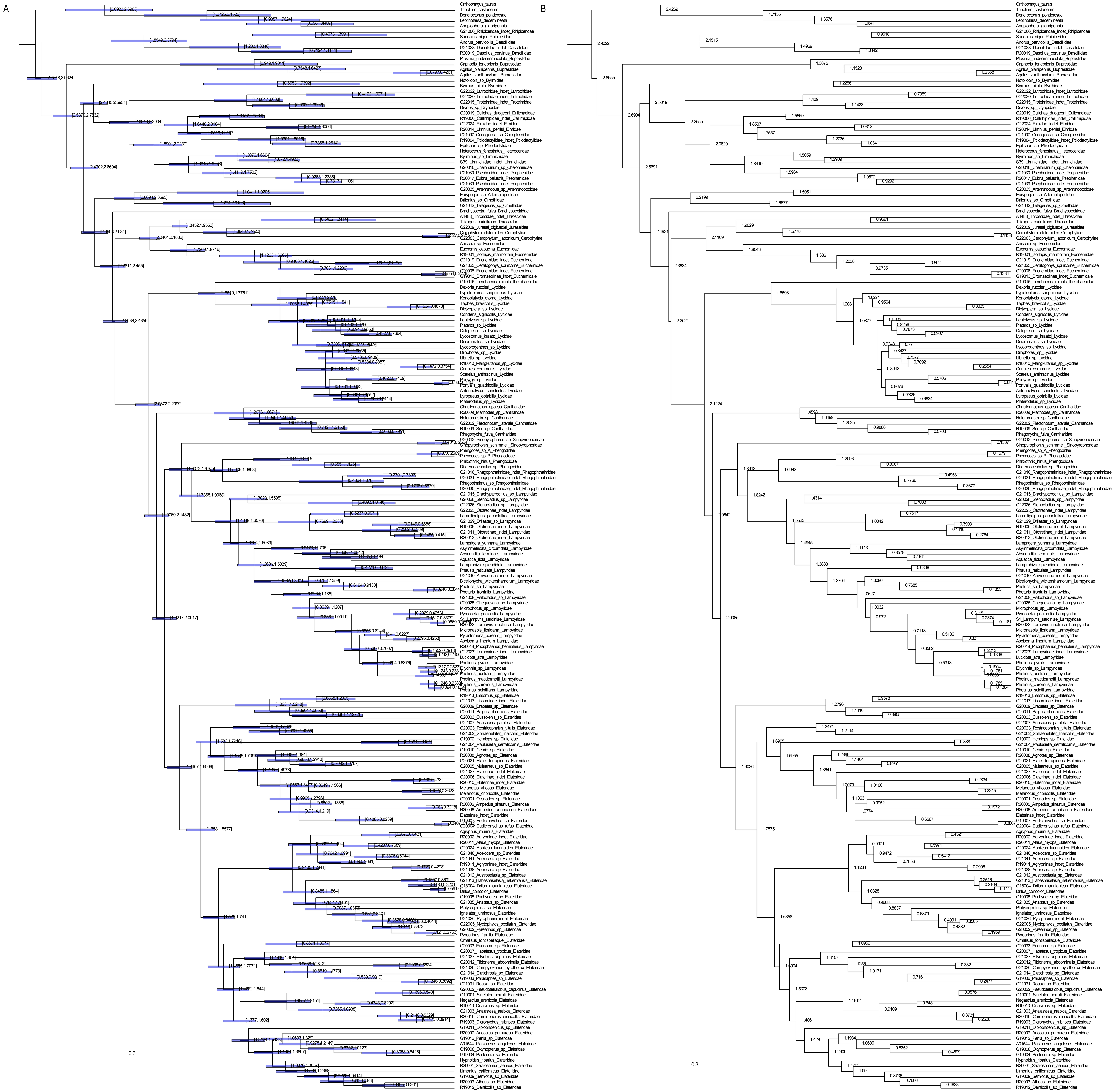

**Supplementary Figure S48.** Divergence time estimation results. Preferred topology analyzed with Dataset B at the nucleotide level under autocorrelated rates. A) Blue horizontal bars represent 95% confidence intervals with lower and upper bound ages labeled at bar endpoints. B) Mean node ages given in units of 100 Mya (e.g., 1.2511 = 125.11 million years ago).

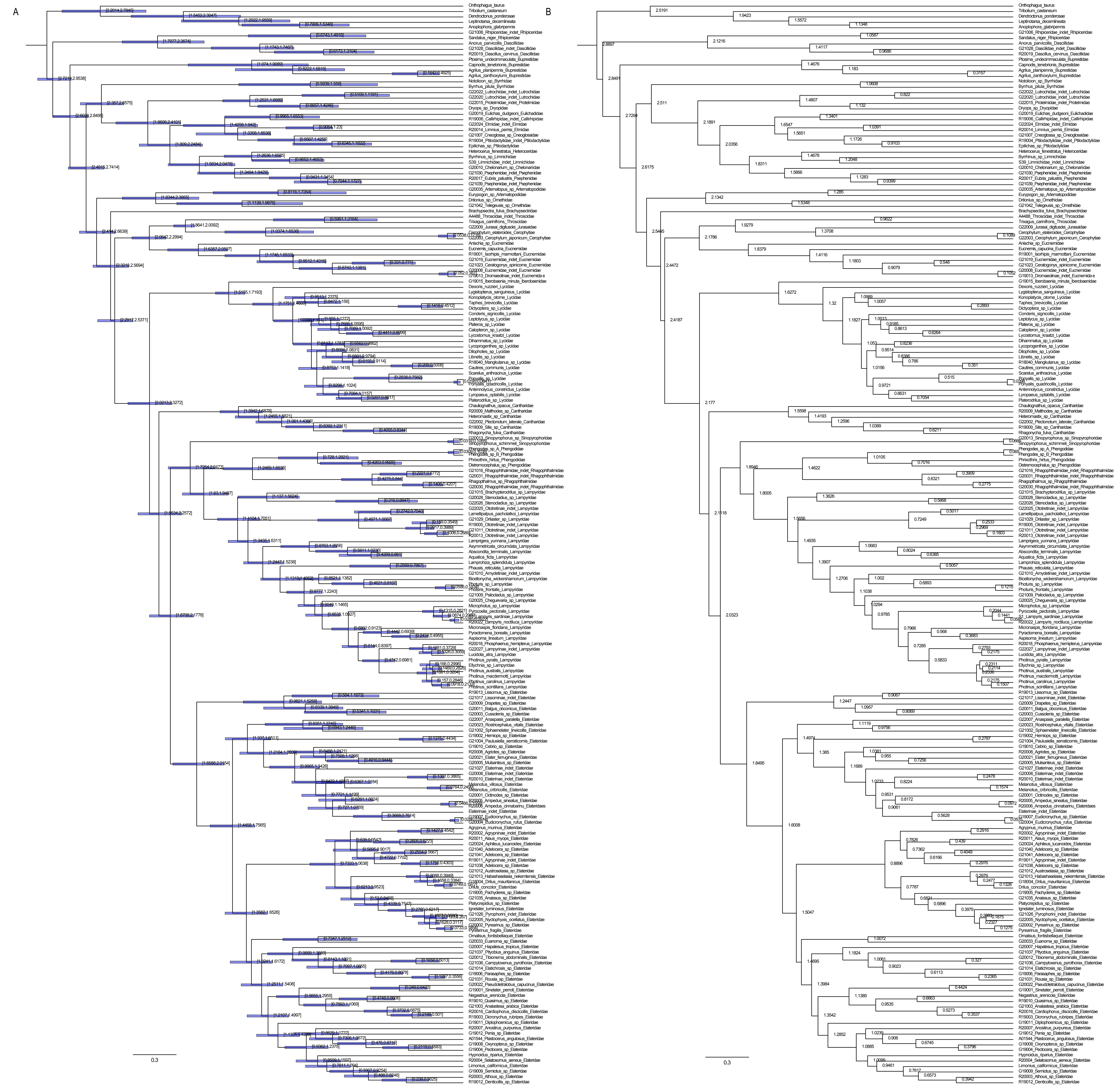

**Supplementary Figure S49.** Divergence time estimation results. Preferred topology analyzed with Dataset A at the nucleotide level under independent rates. A) Blue horizontal bars represent 95% confidence intervals with lower and upper bound ages labeled at bar endpoints. B) Mean node ages given in units of 100 Mya (e.g., 1.2511 = 125.11 million years ago).

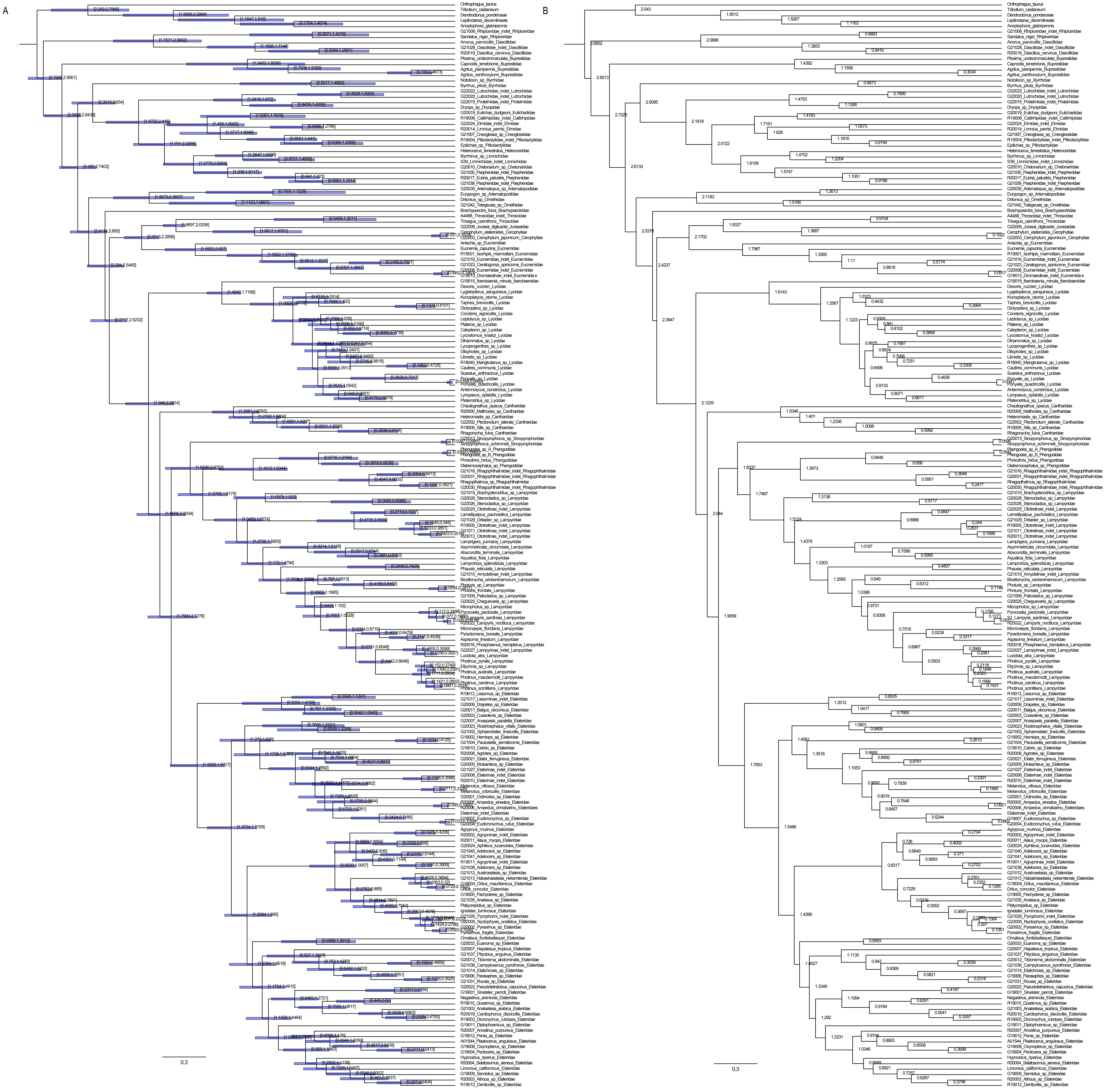

**Supplementary Figure S50.** Divergence time estimation results. Preferred topology analyzed with Dataset B at the nucleotide level under independent rates. A) Blue horizontal bars represent 95% confidence intervals with lower and upper bound ages labeled at bar endpoints. B) Mean node ages given in units of 100 Mya (e.g., 1.2511 = 125.11 million years ago).

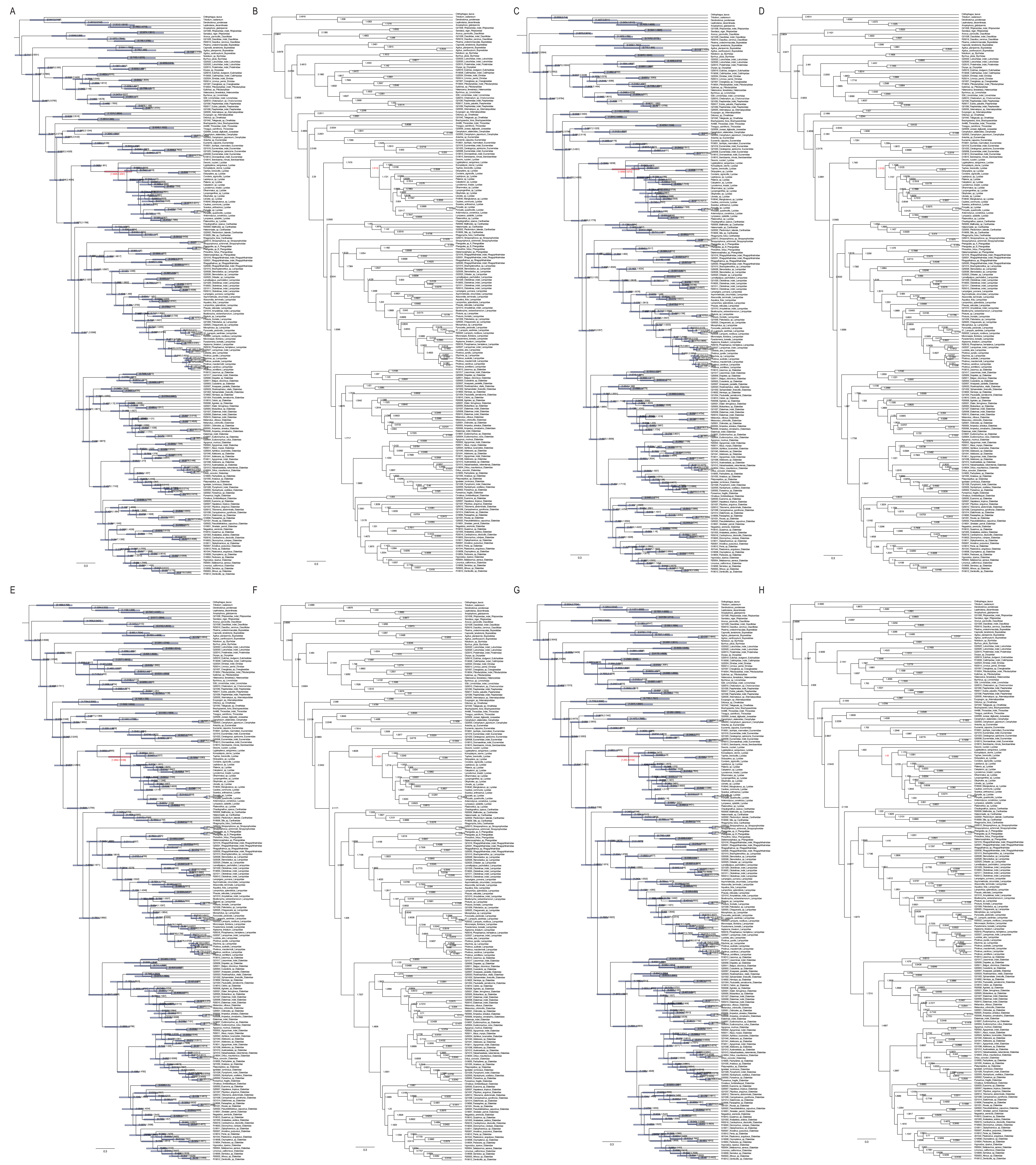

**Supplementary Figure S51.** Divergence time estimation results. Preferred topology analyzed with alternative Lycidae fossil calibration placement under two relaxed clock models (autocorrelated rates and independent rates) and two datasets (Dataset A and Dataset B) analysed at amino acid level. Crown age of Lycidae highlighted in red. For each analysis, left panels (A, C, E, G) display 95% confidence intervals as blue/red horizontal bars with lower and upper bound ages labeled at bar endpoints. Right panels (B, D, F, H) show mean node ages given in units of 100 Mya (e.g., 1.2511 = 125.11 million years ago). A–B) Dataset A under autocorrelated rates. C–D) Dataset B under autocorrelated rates. E–F) Dataset A under independent rates. G–H) Dataset B under independent rates.

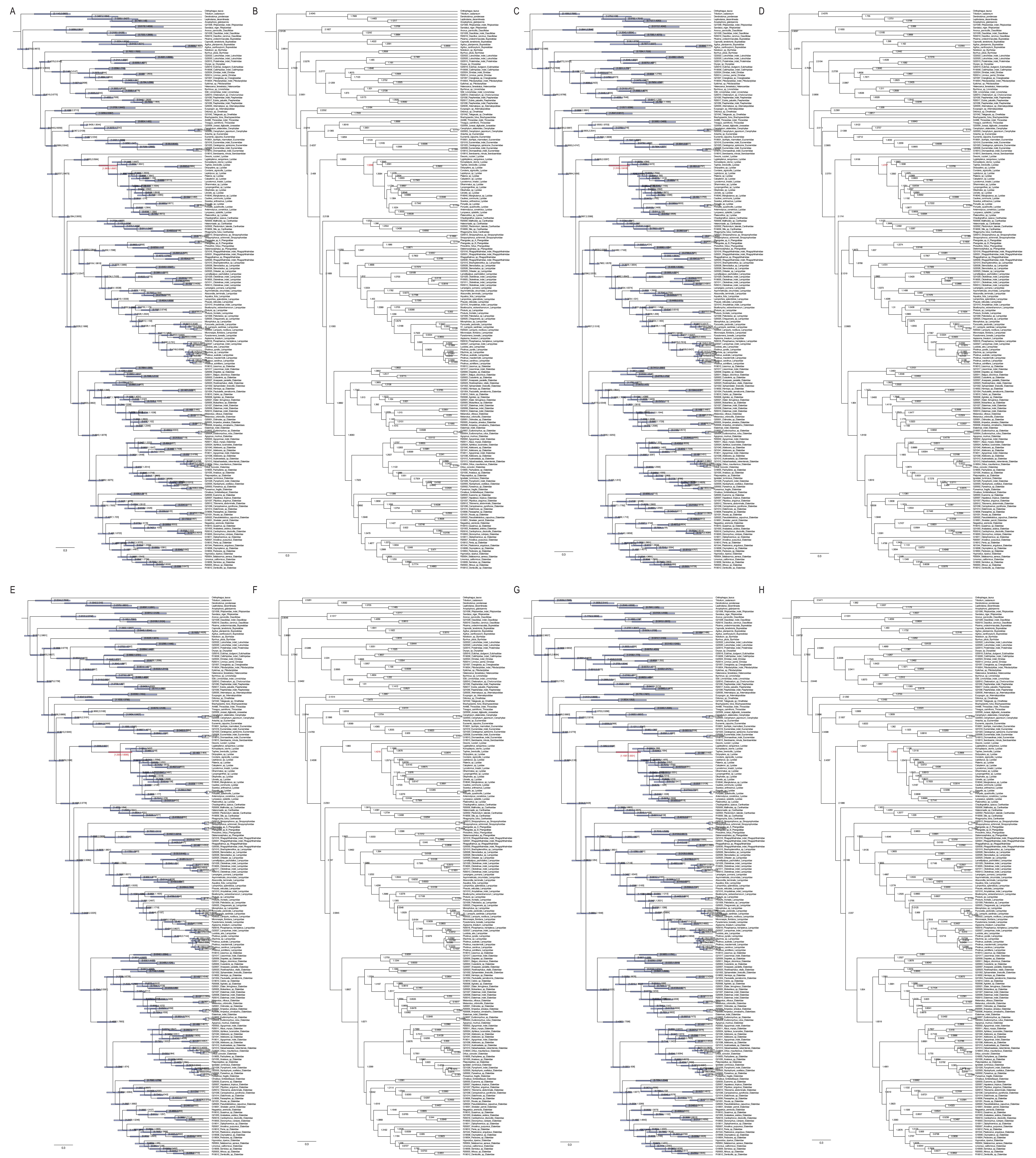

**Supplementary Figure S52.** Divergence time estimation results. Preferred topology analyzed with alternative Lycidae fossil calibration placement under two relaxed clock models (autocorrelated rates and independent rates) and two datasets (Dataset A and Dataset B) analysed at nucleotide level. Crown age of Lycidae highlighted in red. For each analysis, left panels (A, C, E, G) display 95% confidence intervals as blue/red horizontal bars with lower and upper bound ages labeled at bar endpoints. Right panels (B, D, F, H) show mean node ages given in units of 100 Mya (e.g., 1.2511 = 125.11 million years ago). A–B) Dataset A under autocorrelated rates. C–D) Dataset B under autocorrelated rates. E–F) Dataset A under independent rates. G–H) Dataset B under independent rates.

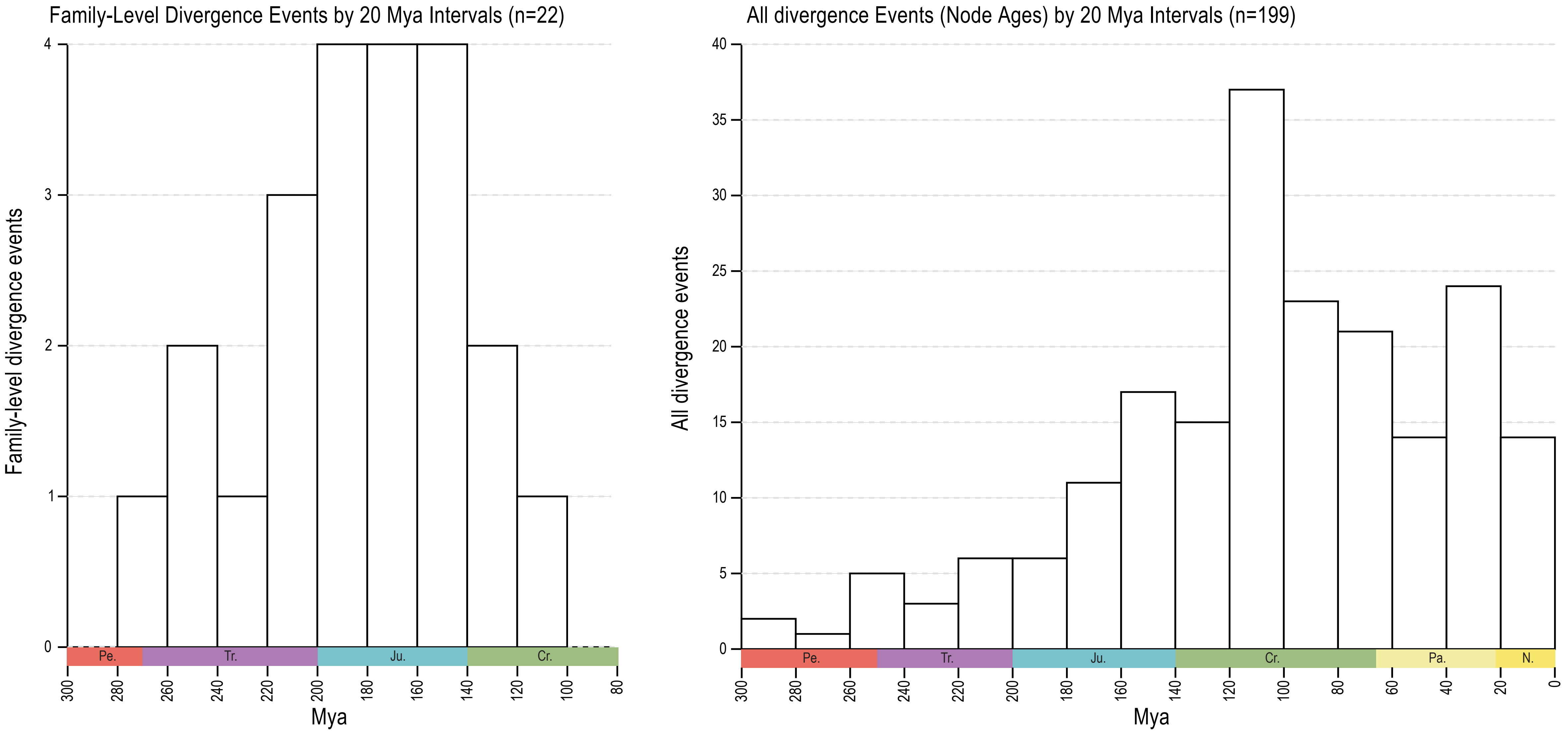

**Supplementary Figure S53.** Family-Level Divergence Events by 20 Mya Intervals (n=22); All divergence Events (Node Ages) by 20 Mya Intervals (n=199). Based on results analyzed with Dataset A at the nucleotide level under autocorrelated rates.

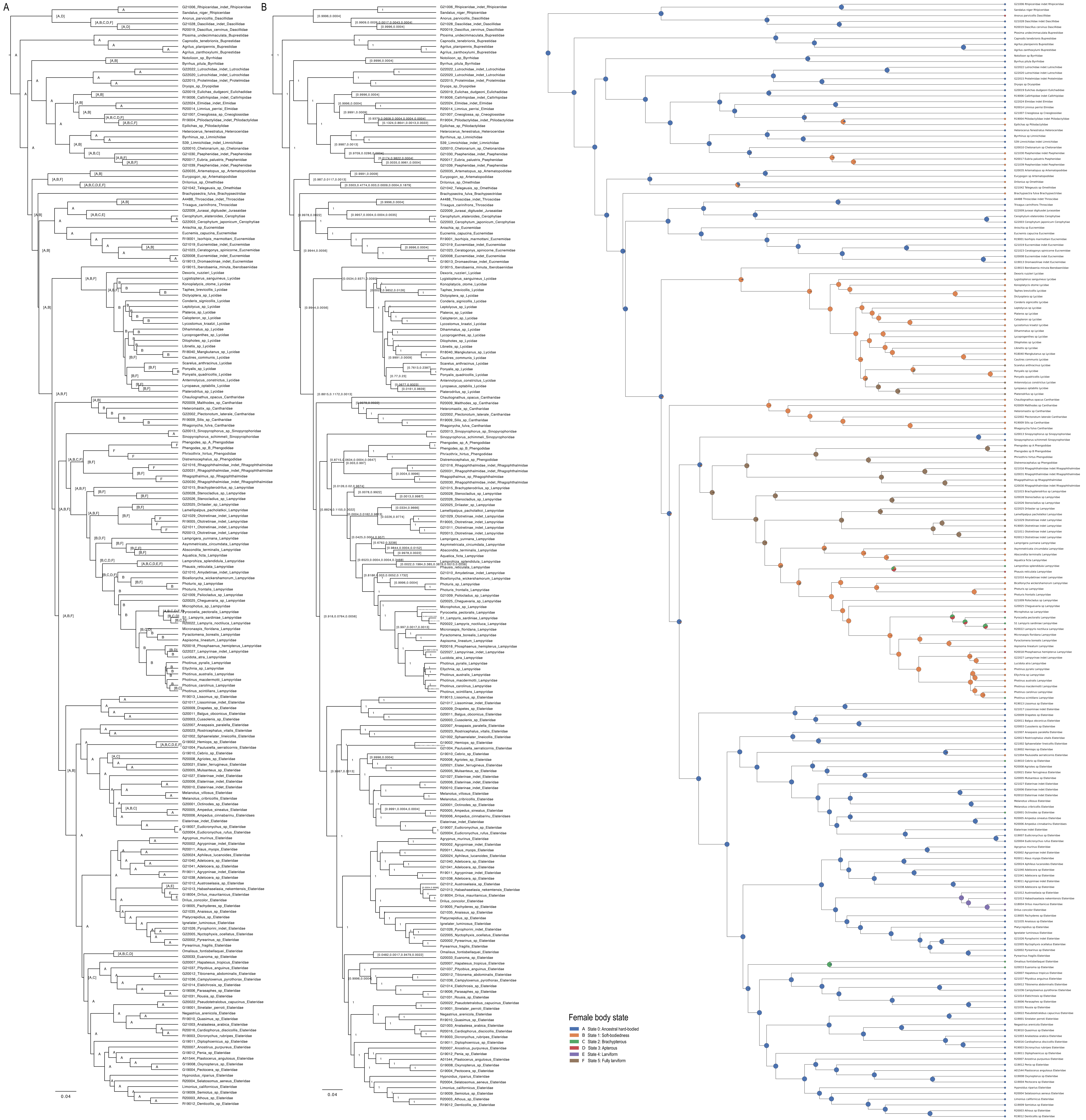

**Supplementary Figure S54.** Ancestral state reconstruction of neoteny using Bayesian inference (BEAST). Trees A) and B) (left) display the estimated states and their probability values for the internal nodes. Tree C) (right) represents the collation of these data, with the color-coded nodes corresponding to the reconstructed female body states (see "Female body state" legend).

**Supplementary Text S1.** List of currently recognized families and subfamilies of Elateriformia, with indication of neoteny, soft-bodied forms, and extinct taxa.

**Legend:**
▲ Ontogenetic modifications have been proven or hypothesized in at least some representatives.
▵ At least some representatives are soft-bodied
† Extinct

### Dascilloidea Guérin-Méneville, 1843

Dascillidae Guérin-Méneville, 1843
- Dascillinae Guérin-Méneville, 1843
- ▲ Karumiinae Escalera, 1913

Rhipiceridae Latreille, 1834

### Buprestoidea Crowson, 1955

Buprestidae Leach, 1815
- Julodinae Lacordaire, 1857
- Polycestinae Lacordaire, 1857
- Galbellinae Reitter, 1911
- Chrysochroinae Laporte, 1835
- Buprestinae Leach, 1815
- Agrilinae Laporte, 1835

Schizopodidae LeConte, 1859

### Byrrhoidea Latreille, 1804

(incl. Dryopoidea Billberg, 1820 (1817) *sensu* Cai et al. 2022, here proposed)

Byrrhidae Latreille, 1804
- Byrrhinae Latreille, 1804
- Syncalyptine Mulsant et Rey, 1869
- Amphicyrtinae LeConte, 1861

Callirhipidae Emden, 1924
Chelonariidae Blanchard, 1845
Cladotomidae Pic, 1914
Cneoglossidae Champion, 1897
Dryopidae Billberg, 1820
Elmidae Curtis, 1830
- Elminae Curtis, 1830
- Larainae LeConte, 1861

Eulichadidae Crowson, 1973
Heteroceridae MacLeay, 1825
- Heterocerinae MacLeay, 1825
- Elythomerinae Pacheco, 1964

Limnichidae Erichson, 1846
- Hyphalinae Britton, 1971
- Cephalobyrrhinae Champion, 1925
- Thaumastodinae Champion, 1925
- Limnichinae Erichson, 1846

Lutrochidae Kasap & Crowson, 1975
Protelmidae Jeannel, 1851
▵ Psephenidae Lacordaire, 1854
- Afroeubriinae Lee et al., 2007
- Eubriinae Lacordaire, 1857
- Eubrianacinae Jacobson, 1913

Psephenoidinae Hilton, 1929
Psepheninae Lacordaire, 1854
△ Ptilodactylidae Laporte, 1836
Anchytarsinae Champion, 1897
Ptilodactylinae Laporte, 1836
Aploglossinae Champion, 1897
Araeopidiinae Lawrence, 1991
† Mastigocoleidae Tihelka, et al., 2022

**Elateroidea Leach, 1815**

Artematopodidae Lacordaire, 1857
Electribiinae Crowson, 1975
Allopogoniinae Crowson, 1973
Artematopodinae Lacordaire, 1857
Badmaateridae Grebennikov et al., 2025
△ Brachypsectridae LeConte & Horn, 1883
△ Cantharidae Imhoff, 1856 (1815)
△ Cantharinae Imhoff, 1856 (1815)
△ Silinae Mulsant, 1862
△ Dysmorphocerinae Brancucci, 1918
△ Heteromastiginae Motyka et al., 2023
△ Malthininae Kiesenwetter, 1852
△ Chauliognathinae LeConte, 1861
Cerophytidae Latreille, 1834
▲ Elateridae Leach, 1815
▲ Agrypninae Candeze, 1857
Cardiophorinae Candeze, 1860
Dendrometrinae Gistel, 1848
▲ Elaterinae Leach, 1815
Hapatesinae Kusy et al., 2021
▲ Hemiopinae Fleutiaux, 1941
Hypnoidinae Douglas et al., 2021
Lissominae Laporte, 1835
Morostominae Dolin, 2000
Negastriinae Nakane & Kishii, 1956
Oestodinae Hyslop, 1917
▲ Omalisinae Lacordaire, 1857
Physodactylinae Lacordaire, 1857
Pityobiinae Hyslop, 1917
† Protagrypninae, Dolin 1973
Subprotelaterinae Fleutiaux, 1920
Tetralobine Laporte, 1840
Thylacosterninae Fleutiaux, 1920
Eucnemidae Eschscholtz, 1829
Perothopinae Lacordaire, 1857
Phyllocerinae Reitter, 1905
Pseudomeninae Muona, 1993
Palaeoxeninae Muona, 1993
Phlegoninae Muona, 1993
Anischiinae Fleutiaux, 1936
Melasinae Fleming, 1821
Eucneminae Eschscholtz, 1829
Dromaeolinae Beauleau, 1919
Schizophilinae Muona, 1993

▲ Iberobaeniidae Bocak et al., 2016
▲ Jurasaidae Rosa et al., 2020
▲ Lampyridae Latreille, 1817
    ▲ Luciolinae Lacordaire, 1857
    ▲ Pterotinae LeConte, 1861
    ▲ Ototretinae McDermott, 1964
    ▲ Lamprohizinae Kazantsev, 2010
    △ Cyphonocerinae Crowson, 1972
    △ Psilocladinae McDermott, 1964
    △ Amydetinae Olivier, 1907
    △ Cheguevariinae Kazantsev, 2006
    △ Photurinae Lacordaire, 1857
    ▲ Lampyrinae Rafinesque, 1815
    ▲ Cladodinae Bocakova et al., 2022
    △ Chespiritoinae Ferreira et al., 2020
▲ Lycidae Laporte, 1836
    ▲ Dexorinae Bocak & Bocakova, 1989
    △ Calochrominae Lacordaire, 1857
    △ Erotinae Leconte, 1881
    ▲ Ateliinae Kleine, 1928
    ▲ Lycinae Laporte, 1836
    ▲ Lyropaeinae Bocak et Bocakova, 1989
    ▲ Metriorrhynchinae Kleine, 1926
    ▲ Xenolycinae Ferreira et al., 2025
▲ Omethidae LeConte, 1861
    △ Driloniinae Crowson, 1972
    △ Omethinae LeConte, 1861
    △ Matheteinae LeConte, 1881
    ▲ Telegeusinae Leng, 1920
▲ Phengodidae LeConte, 1861
    ▲ Phengodinae LeConte, 1861
    ▲ Mastinocerinae LeConte, 1881
    ▲ Penicilliphorinae Paulus, 1975
    ▲ Cydistinae Paulus, 1972
    ▲ Cenophenginae Roza et al., 2024
▲ Rhagophthalmidae Olivier, 1907
Sinopyrophoridae Bi & Li, 2019
Throscidae Laporte, 1840
† △ Cretophengodidae
† Mysteriomorphidae Alekseev et Ellenberger, 2019
† △ Berendtimiridae Winkler, 1987
† △ *Anoeuma lawrencei* Li et al., 2021

**Supplementary Text S2.** Detailed justifications of selected fossil calibrations.

Below we provide detailed justifications for the fossil calibrations used in our MCMCTree divergence time analyses. For fossils previously used as calibrations by Toussaint et al. (2017) and/or Cai et al. (2022) we re-evaluated their justifications following the criteria of Parham et al. (2012). Where our assessment fully agrees with a previous study, we explicitly state this and refer the reader to that source. Where we deviate in the calibrated node, age constraints, or interpretation, we explain the reason for the difference and provide updated justifications. For fossils not utilized in previous studies, we provide original justifications following the same criteria. In all cases we attempted to provide additional comments, assessments of newly published literature, and relevant additional fossil occurrences beyond those discussed in prior work.

**Fossil Calibration 1**
**Calibrated:** Crown Polyphaga (root age in MCMCTree)
**Taxon:** Unnamed Middle Triassic crown polyphagan fossil NIGP162053 presented in Zheng et al. (2018; in Figure 3M)
**Geological deposit/locality:** Ladinian deposit, Tongchuan entomofauna, Hejiafang, Shaanxi, central China
**Soft minimum age:** 237 Mya
**Soft maximum age:** 293.6 Mya
**Reference:** Zheng, D., Chang, S. C., Wang, H., Fang, Y., Wang, J., Feng, C., ... & Wang, B. (2018). Middle-Late Triassic insect radiation revealed by diverse fossils and isotopic ages from China. Science advances, 4(9), eaat1380.
**Justification of Parham et al. (2012) criteria:**
**CR1 (single/multiple OTUs with museum numbers):** A single specimen (NIGP162053); Nanjing Institute of Geology and Palaeontology, Chinese Academy of Sciences, Nanjing, China.
**CR2 (apomorphy-based or phylogenetic analysis):** Presence of apomorphies. The fossil displays features of Polyphaga including a robust body with a distinct prosternal intercoxal process and produced hind pronotal angles. Additionally, the antennae bear a distinct antennal club, a feature widely distributed among numerous polyphagan lineages. Together, these characters clearly represent a crown group polyphagan.
**CR3 (agreement of morphology and molecular data, phylogenetic justification):** Polyphaga is supported as a monophyletic group by extensive molecular and morphological data, forming the sister lineage to the remaining three beetle suborders (Adephaga, Archostemata, Myxophaga). Elateriformia represents one of the earliest diverging lineages within Polyphaga (Beutel et al., 2024), and the unnamed fossil from the Tongchuan Formation can thus be reliably placed within crown Polyphaga based on its characters.
**CR4 (detailed locality and stratigraphy data provided):** Locality and stratigraphy data provided in the original paper, double-checked for accuracy with The Paleobiology Database (https://paleobiodb.org/).
**CR5 (radioisotopic age or numeric age references given):** Age derived from The Paleobiology Database (https://paleobiodb.org/). The soft minimum age (237 Mya) is based on isotopic dating of the insect-bearing layer of the Tongchuan Formation to 237–238 Mya (Zheng et al. 2018). The soft maximum (293.6 Mya) is derived from the oldest beetle fossil

*Coleopsis archaica* (Kirejtshuk et al. 2014) and the age of the Sakmarian period (Schneider & Werneburg 2006, 2012).

**Fossil Calibration 2**
**Calibrated:** Split between Rhipiceridae – Dascillidae
**Taxon:** *Parelateriformius communis*
**Geological deposit/locality:** Haifanggou Formation at Daohugou, Ningcheng County, Inner Mongolia, northeastern China
**Soft minimum age:** 165 Mya
**Soft maximum age:** 235 Mya
**Reference:** **1)** Yan, E. V., & Wang, B. (2010). A new genus of elateriform beetles (Coleoptera, Polyphaga) from the Jurassic of Daohugou, China. Paleontological Journal, 44(3), 297-302. **2)** Li, Y. D., Jin, Z. Y., Ślipiński, A., Huang, D. Y., & Cai, C. Y. (2022). Parelateriformius from the Middle–Late Jurassic of China reinterpreted as the earliest Dascillidae (Coleoptera: Dascilloidea). Palaeoentomology, 5(6), 545-568.
**Justification of Parham et al. (2012) criteria:**
**CR1:** Additional morphologically distinct specimens assigned to the same genus exist. Specimen numbers NIGP nos. 151537a, 151537b (part and counterpart, nearly complete adult beetle, legs missing, antennae partly missing); Nanjing Institute of Geology and Palaeontology, Chinese Academy of Sciences, Nanjing, China.
**CR2:** Presence of apomorphies. The fossil displays key similarities with Dascillidae: distinct frontoclypeal suture, head lacking posterior constriction, large protuberant eyes, exposed antennal insertions, 11-segmented serrate antennae, pronotum with complete lateral carina, long prosternum in front of procoxae, complete prosternal process between coxae, transverse procoxal cavities with lateral notches exposing protrochantins, narrowly separated mesocoxal cavities, metaventrite with discrimen and katepisternal sutures almost complete, transverse hind coxae with large coxal plates, 5-segmented tarsi, simple claws lacking an empodium, and 5 abdominal ventrites. This places it within crown Dascilloidea (Cai et al. 2022).
**CR3:** Reliably assigned to Dascilloidea/Dascillidae. Dascilloidea is supported as a monophyletic group.
**CR4:** Locality and stratigraphy data provided in the original papers, double-checked for accuracy with The Paleobiology Database (https://paleobiodb.org/).
**CR5:** Age derived from The Paleobiology Database (https://paleobiodb.org/). The soft minimum age (165 Ma) is based on Ar/Ar and SHRIMP U-Pb dating of intermediate-acid volcanic rock overlying the Daohugou fossiliferous beds (Chen et al. 2004), widely adopted by palaeoentomologists. The soft maximum (235 Ma) is derived from the age of the Madygen Formation at Dzhailyau-Cho, Kyrgyzstan (237 ± 2 Ma; Voigt et al. 2017). Although beetles dominate the Madygen entomofauna, polyphagans are rare and represented only by stem group taxa such as *Peltosyne triassica* and *P. varyvrosa* (Ponomarenko 1977; Yan et al. 2018).
**Overlap note:** This fossil is also used in Cai et al. (2022) [Node 9] for the same node with identical age constraints.

**Fossil Calibration 3**
**Calibrated:** Crown Dascillidae
**Taxon:** *Cretodascillus sinensis,* Dascillidae
**Geological deposit/locality:** Liutiaogou, Yixian Formation, China

**Soft minimum age:** 122.5 Mya
**Soft maximum age:** 235 Mya
**Reference:** Jin, Z., Slipinski, A., Pang, H., & Ren, D. (2013). A new Mesozoic species of soft-bodied plant beetle (Coleoptera: Dascillidae) from the Early Cretaceous of Inner Mongolia, China with a review of fossil Dascillidae. In Annales Zoologici (Vol. 63, No. 3, pp. 501-509). Museum and Institute of Zoology, Polish Academy of Sciences.
**Justification of Parham et al. (2012) criteria:**
**CR1:** A single specimen (CNU-COL-LB-2010330). Full museum data in the reference above.
**CR2:** Presence of apomorphies diagnostic for Dascillidae.
**CR3:** Reliably assigned to Dascillidae. Dascillidae is supported as a monophyletic group.
**CR4:** Locality and stratigraphy data provided in the original paper, double-checked for accuracy with The Paleobiology Database (https://paleobiodb.org/).
**CR5:** Age derived from The Paleobiology Database (https://paleobiodb.org/). The soft minimum age (122.5 Mya) is based on high-precision $^{40}Ar/^{39}Ar$ dating of the Yixian Formation: the uppermost part of the main fossil bed was dated at 122.9 ± 0.7 Mya (Chang et al. 2009), providing a conservative minimum of 122.5 Mya (consistent with Toussaint et al. 2017). The soft maximum (235 Mya) is derived from the Madygen Formation at Dzhailyau-Cho, Kyrgyzstan (237 ± 2 Mya; Voigt et al. 2017), see Fossil Calibration 2 for details.
**Overlap note:** This fossil is also used in Toussaint et al. (2017) [syen12198, FC 14] as calibration for stem Dascillidae with the same minimum age (122.5 Ma).

**Fossil Calibration 4**
**Calibrated:** Split of (Lutrochidae (Dryopidae + Protelmidae))
**Taxon:** *Mastigocoleus rhinoceros*, Mastigocoleidae
**Specimen:** Holotype SMNS 66552; Staatliches Museum für Naturkunde Stuttgart, Germany
**Geological deposit/locality:** Nova Olinda Member, Crato Formation, Chapada do Araripe, northeastern Brazil; Aptian–Albian (Lower Cretaceous)
**Soft minimum age:** 112.6 Mya
**Soft maximum age:** 165 Mya
**Reference:** Tihelka, E., Jaech, M. A., Kundrata, R., Li, Y. D., Engel, M. S., Lozano-Fernandez, J., ... & Cai, C. (2022). Mastigocoleidae fam. nov., a new Mesozoic beetle family and the early evolution of Dryopoidea (Coleoptera). Insect Systematics and Diversity, 6(3), 3.
**Justification of Parham et al. (2012) criteria:**
**CR1:** Multiple species described and placed in the newly established family Mastigocoleidae. Holotype SMNS 66552; Staatliches Museum für Naturkunde Stuttgart, Germany.
**CR2:** Presence of apomorphies reliably placing the taxon within the clade (Lutrochidae (Dryopidae + Protelmidae)).
**CR3:** Monophyly of the clade uniting Lutrochidae and Dryopidae is supported by our molecular phylogenetic analyses as well as the phylogenetic analysis of Tihelka et al. (2022), which recovered Mastigocoleidae as sister to (Dryopidae + Lutrochidae).
**CR4:** Locality and stratigraphy data provided in the original paper, double-checked for accuracy with The Paleobiology Database (https://paleobiodb.org/).
**CR5:** Age derived from The Paleobiology Database (https://paleobiodb.org/). The soft minimum age (112.6 Mya) is based on the upper boundary of the Aptian (113.0 ± 0.4 Mya), giving a conservative minimum of 112.6 Mya. The soft maximum (165 Mya) is based on the Daohugou biota, which lacks representatives of this clade.

**Additional fossils:** See the reference above.

**Fossil Calibration 5**
**Calibrated:** Split of Ptilodactylidae – Cneoglossidae
**Taxon:** *Aphebodactyla rhetine*, Ptilodactylidae
**Specimen:** Holotype KU-NHM-ENT Bu-006 (male, amber inclusion); Fossil Insect Collection, Division of Entomology, Natural History Museum, University of Kansas, Lawrence, Kansas, USA.
**Geological deposit/locality:** Burmese amber, Tanai Village, Hukawng Valley, Kachin State, Myanmar; latest Albian
**Soft minimum age:** 98.17 Mya
**Soft maximum age:** 165 Mya
**Reference:** Chatzimanolis, S., Cashion, M. E., Engel, M. S., & Falin, Z. H. (2012). A new genus of Ptilodactylidae (Coleoptera: Byrrhoidea) in mid-Cretaceous amber from Myanmar (Burma). Geodiversitas, 34(3), 569-574.
**Justification of Parham et al. (2012) criteria:**
**CR1:** A single specimen (holotype KU-NHM-ENT Bu-006, male). Full museum data in the reference above.
**CR2:** Presence of apomorphies. *Aphebodactyla rhetine* can be placed within Ptilodactylidae based on the cordiform mesoscutellum, pseudotetramerous tarsi, and 11-segmented pectinate antennae in the male. The fossil cannot be assigned to any extant subfamily as it intermingles putatively derived features of several ptilodactylid subfamilies, and is tentatively considered a stem-group member of the family (Chatzimanolis et al. 2012).
**CR3:** Ptilodactylidae is recovered as a monophyletic group sister to Cneoglossidae in our molecular phylogenetic analyses, consistent with previous molecular studies. The placement of *Aphebodactyla rhetine* within Ptilodactylidae, as justified under CR2, thus reliably positions this fossil at the split between Ptilodactylidae and Cneoglossidae.
**CR4:** Locality and stratigraphy data provided in the original paper, double-checked for accuracy with The Paleobiology Database (https://paleobiodb.org/).
**CR5:** Age derived from The Paleobiology Database (https://paleobiodb.org/). The soft minimum age (98.17 Mya) is based on radiometric dating of the Burmese amber-bearing horizon at 98.79 ± 0.62 Mya (Shi et al. 2012), taken as a minimum age estimate (Mao et al. 2018). The soft maximum (165 Mya) is based on the Daohugou biota, which lacks representatives of this clade (Chen et al. 2004).
**Additional fossils:** Many undescribed specimens from Burmese amber (personal collection of authors).

**Fossil Calibration 6**
**Calibrated:** Crown Elmidae
**Taxon:** *Cretohypsilara parva*, Elmidae
**Specimen:** Holotype NIGP163429 (male, amber inclusion); Nanjing Institute of Geology and Palaeontology, Chinese Academy of Sciences, Nanjing, China.
**Geological deposit/locality:** Burmese amber, Hukawng Valley, northern Myanmar; earliest Cenomanian
**Soft minimum age:** 98.17 Mya
**Soft maximum age**: 165 Mya

**Reference:** Cai, C., Maier, C. A., & Huang, D. (2018). A new riffle beetle in Upper Cretaceous amber from northern Myanmar (Coleoptera: Elmidae). Cretaceous Research, 89, 154-159.

**Justification of Parham et al. (2012) criteria:**

**CR1:** A single specimen (holotype NIGP163429, male). Full museum data in the reference above.

**CR2:** Combination of diagnostic characters. *Cretohypsilara parva* is confidently placed in Elmidae based on the following combination of characters: widely separated antennae, large protuberant eyes, slender legs with long simple tarsi, all tarsi 5-segmented with the last tarsomere longer than the other four combined, very long pretarsal claws, completely carinate lateral pronotal margins, prosternum expanded anteriorly forming a turtleneck, five abdominal ventrites, and male genitalia with two parameres and median piece forming three elongate lobes as in extant Elmidae (Cai et al. 2018). The fossil bears similarity to the subfamily Larainae based on its subovate body covered with dense pubescence, clubbed antennae, and exposed protrochantins, but cannot be confidently attributed to either extant subfamily and is therefore treated as Elmidae *incertae sedis*.

**CR3:** Elmidae is recovered as a monophyletic group in our molecular phylogenetic analyses. The placement of *Cretohypsilara parva* within crown Elmidae, as justified under CR2, is unambiguous. It is noteworthy that Cai et al. (2022) used *Elmadulescens rugosus* from the Lower Cretaceous El Soplao amber to calibrate the node Elmidae – (Limnichidae + Heteroceridae). However, the familial placement of *Elmadulescens rugosus* was questioned by Bukejs et al. (2015), who transferred the species to Coleoptera *incertae sedis,* rendering it an unreliable calibration for Elmidae. We therefore use *Cretohypsilara parva*, whose placement within Elmidae is unambiguous, as our calibration for crown Elmidae.

**CR4:** Locality and stratigraphy data provided in the original paper, double-checked for accuracy with The Paleobiology Database (https://paleobiodb.org/).

**CR5:** Age derived from The Paleobiology Database (https://paleobiodb.org/). The soft minimum age (98.17 Mya) is based on U-Pb zircon dating of the Burmese amber-bearing horizon at 98.79 ± 0.62 Mya (Shi et al. 2012), taken as a soft minimum age estimate (Mao et al. 2018). The soft maximum (165 Mya) is derived from the age of the Daohugou biota, constrained by Ar/Ar and SHRIMP U-Pb dating to approximately 165 Mya (Chen et al. 2004), a deposit that preserves diverse Elateriformia but lacks any definitive representatives of Elmidae.

**Fossil Calibration 7**

**Calibrated:** Split of Limnichidae – Heteroceridae

**Taxon:** *Heterocerites magnus*, Heteroceridae

**Geological deposit/locality:** Chaomidian (Huanbanjigou), Yixian Formation, Shangyuan, Beipiao City, Liaoning Province, China

**Soft minimum age:** 122.2 Mya

**Soft maximum age:** 165 Mya

**Reference:** Prokin, A. A., & Ren, D. (2011). New species of variegated mud-loving beetles (Coleoptera: Heteroceridae) from mesozoic deposits of China. Paleontological Journal, 45, 284-286.

**Justification of Parham et al. (2012) criteria:**

**CR1:** Multiple specimens (holotype CNU-COL-LB-2010023, impression of adult beetle with partly preserved legs and mandibles; Key Lab of Insect Evolution and Environmental

Changes, Capital Normal University, Beijing, China). Full museum data in the reference above.
**CR2:** Based on careful examination of type photos for presence of apomorphies. The fossil can be placed in Heteroceridae on the basis of its densely pubescent body, large head with prominent mandibles and labrum, characteristically modified antennae with the distal half forming a loosely serrated club, a series of spines at the outer edges of the tibiae, short mesosternum, and stridulate ridges on ventrite 1. The dark elytra possess three pairs of light spots, a pattern similar to those found in extant members of the family (Clarke 1973; Vanin et al. 2016; Cai et al. 2022).
**CR3:** Reliably assigned to Heteroceridae. The monophyly of Heteroceridae is well supported.
**CR4:** Locality and stratigraphy data provided in the original paper, double-checked for accuracy with The Paleobiology Database (https://paleobiodb.org/).
**CR5:** Age derived from The Paleobiology Database (https://paleobiodb.org/). The soft minimum age (122.2 Mya) is based on $^{40}Ar/^{39}Ar$ dating of the Yixian Formation: the lowest part of the main fossil bed was dated at 124.1 ± 0.3 Mya and the uppermost at 122.9 ± 0.7 Mya (Chang et al. 2009), with 122.2 Mya adopted as a conservative minimum consistent with Cai et al. (2022). The soft maximum (165 Mya) is based on the Daohugou biota, which lacks representatives of this family.
**Overlap note:** This fossil is also used in Cai et al. (2022) [Node 14, Crown Heteroceridae] with a minimum of 122.2 Mya, and in Toussaint et al. (2017) [FC 17] with a minimum of 125.5 Mya for the same node. We follow the age of Cai et al. (2022).

**Fossil Calibration 8**
**Calibrated:** Crown Psephenidae
**Taxon:** *Vetujinbrianax cretaceus,* Psephenidae
**Geological deposit/locality:** Lowermost Cenomanian, amber mine near Noije Bum Village, Tanaing Town, Hukawng Valley, northern Myanmar
**Soft minimum age:** 98.17 Mya
**Soft maximum age:** 165 Mya
**Reference:** Cai, C., & Huang, D. (2018). First Mesozoic water-penny beetles from Upper Cretaceous Burmese amber (Coleoptera: Psephenidae). Cretaceous Research, 91, 202-207.
**Justification of Parham et al. (2012) criteria:**
**CR1:** Single clear specimen (NIGP168215, male); Nanjing Institute of Geology and Palaeontology, Chinese Academy of Sciences, Nanjing, China. Full museum data in the reference above.
**CR2:** The fossil can be placed in Psephenidae on the basis of an apparently soft body, pectinate antennae in males, exposed protrochantins, relatively long maxillary palps, and simple 5-segmented tarsi. It can be further assigned to the extant subfamily Eubrianacinae based on the presence of plesiomorphic characters including a short prosternal process and paired tibial apical spurs, similar to the extant Oriental genus *Jinbrianax* (Lee et al. 2007; Cai & Huang 2018). It represents the oldest fossil record of Eubrianacinae and the first definitive adult fossil of Psephenidae from the Mesozoic.
**CR3:** Reliably assigned to Psephenidae, subfamily Eubrianacinae. The monophyly of Psephenidae and placement of Eubrianacinae are well supported.
**CR4:** Locality and stratigraphy data provided in the original paper, double-checked for accuracy with The Paleobiology Database (https://paleobiodb.org/).

**CR5:** Age derived from The Paleobiology Database (https://paleobiodb.org/). The soft minimum age (98.17 Mya) is based on radiometric dating of volcanoclastic matrix from the amber-bearing horizon at 98.79 ± 0.62 Mya (Shi et al. 2012), interpreted as a minimum age estimate (Mao et al. 2018). The discovery of an ammonite in the amber confirms it is at most Albian in age (Yu et al. 2019). The soft maximum (165 Mya) is based on the Daohugou biota, which lacks representatives of this clade (Chen et al. 2004).
**Overlap note:** This fossil is also used in Cai et al. (2022) [Node 12, Crown Psephenidae] with identical age constraints (soft minimum 98.17 Mya, soft maximum 165 Mya).
**Additional fossils:** *Pseudomataeopsephus* and *Succinumanax birmaniasis* and suspected larvae fossil from Lower Barremian of Spain mentioned in Bao et al., (2018).

Li, Y. D., Huang, D., & Cai, C. (2022). Pseudomataeopsephus, a new genus of water penny beetles from mid-Cretaceous Burmese amber (Coleoptera: Psephenidae). Cretaceous Research, 130, 105055.
Bao, T., Rust, J., & Wang, B. (2018). Systematics, phylogeny and taphonomy of Cretaceous Psephenidae (Insecta: Coleoptera) from Burmese amber. Palaeontographica Abteilung A, 310, 131-159.

**Fossil Calibration 9**
**Calibrated:** Split of Artematopodidae – Omethidae
**Taxon:** *Sinobrevipogon jurassicus*, Artematopodidae
**Geological deposit/locality:** Haifanggou Formation at Daohugou, Ningcheng County, Inner Mongolia, northeastern China
**Soft minimum age:** 165 Mya
**Soft maximum age:** 235 Mya
**Reference:** Cai, C. Y., Lawrence, J. F., Slipinski, A., & Huang, D. Y. (2015). Jurassic artematopodid beetles and their implications for the early evolution of Artematopodidae (Coleoptera). Systematic Entomology, 40(4), 779-788.
**Justification of Parham et al. (2012) criteria:**
**CR1:** Multiple specimens (holotype NIGP160704; paratypes NIGP160705, NIGP160706; holotype nearly completely preserved adult beetle with both dorsal and ventral characters visible including hindwings); Nanjing Institute of Geology and Palaeontology, Chinese Academy of Sciences, Nanjing, China. Full museum data in the reference above.
**CR2:** Phylogenetic analysis. *Sinobrevipogon jurassicus* exhibits defining features of Artematopodidae, including paired carinae on the prosternum and an internal apical interlocking tongue on the ventral side of each elytron. It differs from modern members by having the mesocoxal cavities closed by the mesepimeron and the anterolateral edge of the metanepisternum. A morphological phylogenetic analysis nonetheless recovered *Sinobrevipogon* within the crown Artematopodidae (Cai et al. 2015; 2020).
**CR3:** Reliably assigned to Artematopodidae. The monophyly of Artematopodidae is well supported.
**CR4:** Locality and stratigraphy data provided in the original paper, double-checked for accuracy with The Paleobiology Database (https://paleobiodb.org/).
**CR5:** Age derived from The Paleobiology Database (https://paleobiodb.org/). The soft minimum is derived from the age of the Daohugou biota, constrained by Ar/Ar and SHRIMP U-Pb dating to approximately 165 Mya (Chen et al. 2004). The soft maximum (235 Mya) is derived from the Madygen Formation at Dzhailyau-Cho, Kyrgyzstan (237 ± 2 Ma; Voigt et al. 2017).

**Overlap note:** This fossil is also used in Cai et al. (2022) [Node 15, Artematopodidae–Omethidae] with a minimum of 165 Mya, and in Toussaint et al. (2017) [FC 18] with a minimum of 157.3 Ma. We follow the age of Cai et al. (2022).
**Additional fossils:** *Tarsomegamerus mesozoicus* (Cai et al.,2015) from Jurassic.

**Fossil Calibration 10**
**Calibrated:** Split of (Throscidae (Cerophytidae + Jurassaidae))
**Taxon:** *Praelaterium problematicum*, Cerophytidae
**Geological deposit/locality:** Dgil'skaya Formation, Issyk-Kul', Tong District, Osh Region, Kyrgyzstan; Lower Jurassic (ca. 201.6–189.6 Mya)
**Soft minimum age:** 190.8 Ma
**Soft maximum age:** 235 Ma
**Reference:** Yu et al. (2019)
**Justification of Parham et al. (2012) criteria:**
**CR1:** The holotype of *Praelaterium problematicum* (PIN No. 358/351) is figured in Yu et al. (2019) as Figure 4, with both a line drawing (4A) and photograph (4B). However, the preservation is limited. It is described as a damaged imprint of mostly the ventral aspect, without antennae, legs, or scutellar shield. The figures are of sufficient quality to verify the key characters used for placement in Cerophytidae, particularly the prosternal process shape and coxal cavity structure, though the incompleteness of the specimen means not all diagnostic characters can be confirmed from the figures alone.
**CR2:** Placement within Cerophytidae is supported by the shape of the prosternal process, broad behind the coxae, then abruptly narrowed to a blade-like apex extending into the mesoventral cavity, which Yu et al. (2019) identify as the principal synapomorphy of Cerophytidae observable in fossils. Additional supporting characters include complete lateral pronotal carinae, circular procoxal cavities, narrow parallel-sided metanepisterna, and transverse metacoxae with complete metacoxal plates. The specimen was included in the phylogenetic analysis and recovered within Cerophytidae.
**CR3:** Yu et al. (2019) formally reassign *Praelaterium* from Praelateriidae (which they synonymize with Cerophytidae) to Cerophytidae based on detailed morphological reanalysis, and the taxon is included in their phylogenetic analysis. The family assignment is therefore well-supported, though it is worth noting that the holotype's preservation is incomplete, and several characters are coded as uncertain in the data matrix. The assignment rests substantially on the prosternal process structure.
**CR4:** Locality and stratigraphy data provided in the original paper, double-checked for accuracy with The Paleobiology Database (https://paleobiodb.org/).
**CR5:** Age derived from The Paleobiology Database (https://paleobiodb.org/). The minimum age (190.8 Mya) is based on the age of the Dgil'skaya Formation (Lower Jurassic, ca. 201.6–189.6 Mya; Kotov 2007), with 190.8 Mya as a conservative minimum. The soft maximum (235 Mya) is derived from the Madygen Formation, Kyrgyzstan (237 ± 2 Mya; Voigt et al. 2017).
**Additional fossils:**
*Elaterocoleus oculatus* from same deposit as *P. problematicum*.
*Wongaroo amplipectorale* from Talbragar Fossil Fish Bed, NSW, Australia; Kimmeridgian (ca. 155.7–150.8 Mya).
*Jurassophytum cleidecostae* from Daohugou, Inner Mongolia; Jiulongshan Formation, Middle Jurassic (ca. 164.7–155.7 Mya).

High diversity of Cerophytidae is additionally described from the Cretaceous (see Yu et al. 2019).

**Fossil Calibration 11**
**Calibrated:** Split of Anischiinae – "core" Eucnemidae
**Taxon:** *Rheanischia brevicornis*, Eucnemidae
**Geological deposit/locality:** Yixian Formation, Chaomidian Village, Liaoning Province, China; Lower Cretaceous (Jehol Biota)
**Soft minimum age:** 122.2 Mya
**Soft maximum age:** 235 Mya
**Reference:** Li, H., Chang, H., Muona, J., Zhao, Y., & Ren, D. (2021). Subfamily Anischiinae (Coleoptera: Eucnemidae) in early Cretaceous of Northeast China. Insects, 12(2), 105.
**Justification of Parham et al. (2012) criteria:**
**CR1:** The holotype (CNU-COL-LB2018344) and paratype (CNU-COL-LB2018313) are figured in Li et al. (2021), with photographs in dorsal and ventral views (Figures 1 and 3) and detailed anatomical line drawings (Figure 2). The figures are of sufficient quality to verify the key diagnostic characters used to place the fossil within Eucnemidae and Anischiinae, including the pronotal interlocking cavities, elytral excretory pores, and antennal structure.
**CR2:** Combination of diagnostic characters reliably placing the fossil within Eucnemidae, subfamily Anischiinae.
**CR3:** Reliably assigned to Eucnemidae.
**CR4:** Locality and stratigraphy data provided in the original paper, double-checked for accuracy with The Paleobiology Database (https://paleobiodb.org/).
**CR5:** Age derived from The Paleobiology Database (https://paleobiodb.org/). The soft minimum age (122.2 Mya) is based on $^{40}Ar/^{39}Ar$ dating of the Yixian Formation: the uppermost part of the main fossil bed was dated at 122.9 ± 0.7 Mya (Chang et al. 2009), providing a conservative minimum of 122.2 Mya. The soft maximum (235 Mya) is derived from the Madygen Formation, Kyrgyzstan (237 ± 2 Mya; Voigt et al. 2017), see Fossil Calibration 2 for details.
**Additional fossils:** Family is represented by diverse numbers of fossils spanning from Middle Jurassic to Cretaceous see additional references.
*Paradesmatus baiae* Chang, Kirejtshuk & Ren, 2009 from Daohugou beds, Jiulongshan Formation, Inner Mongolia; Middle Jurassic.
*Beattieellus jurassicus* Oberprieler et al., 2016 from Talbragar Fish Bed, NSW, Australia; Kimmeridgian–Tithonian (151 ± 4 Mya).
Additional references: Muona et al. (2020); Chang et al. (2009, 2010); Oberprieler et al. (2016); Huali et al. (2016).

Muona, J., Chang, H., & Ren, D. (2020). The clicking Elateroidea from Chinese Mesozoic deposits (Insecta, Coleoptera). Insects, 11(12), 875.
Chang, H., Kirejtshuk, A., & Ren, D. (2010). New fossil elaterids (Coleoptera: Polyphaga: Elateridae) from the Jehol biota in China. Annals of the Entomological Society of America, 103(6), 866-874.
Chang, H., Kirejtshuk, A. G., Ren, D., & Shih, C. (2009). First fossil click beetles from the middle Jurassic of Inner Mongolia, China (Coleoptera: Elateridae). In Annales zoologici (Vol. 59, No. 1, pp. 7-14). Museum and Institute of Zoology, Polish Academy of Sciences.

Oberprieler, R. G., Ashman, L. G., Frese, M., & Ślipiński, A. (2016). The first elateroid beetles (Coleoptera: Polyphaga: Elateroidea) from the Upper Jurassic of Australia. Zootaxa, 4147(2), 177-191.
Huali, C., Muona, J., Hanyong, P., Li, X., Chen, W., Teräväinen, M., ... & Songhai, J. (2016). Chinese Cretaceous larva exposes a southern Californian living fossil (Insecta, Coleoptera, Eucnemidae). Cladistics, 32(2), 211-214.

**Fossil Calibration 12**
**Calibrated:** Split of Lycidae – Iberobaeniidae (conservatively) or crown Lycidae
**Taxon:** *Burmolycus compactus*, Lycidae
**Geological deposit/locality:** Lowermost Cenomanian, amber mine near Noije Bum Village, Tanaing Town, Hukawng Valley, northern Myanmar
**Soft minimum age:** 98.17 Mya
**Soft maximum age:** 165 Mya
**Reference:** Bocak, L., Li, Y., & Ellenberger, S. (2019). The discovery of Burmolycus compactus gen. et sp. nov. from the mid-Cretaceous of Myanmar provides the evidence for early diversification of net-winged beetles (Coleoptera, Lycidae). Cretaceous research, 99, 149-155.
**Justification of Parham et al. (2012) criteria:**
**CR1:** Single specimen (holotype MB.I 7319, female; amber inclusion; Museum für Naturkunde, Berlin, Germany). Full museum data in the reference above.
**CR2:** Combination of diagnostic characters. *Burmolycus compactus* can be placed in Lycidae on the basis of 11-segmented antennae, a ridged pronotum, a very short and transverse prosternum, 5-segmented tarsi, and lobed tarsomeres 2–4 (Bocak & Bocakova 2008, 2010). It can further be assigned to the extant subfamily Dexorinae based on a prolonged cranium, frontally inserted antennae, antennomere 2 relatively long compared to antennomere 3, hypognathous head, rounded mouthpart cavity, reduced labium with 1 or 2-segmented labial palpi, and femora obliquely attached to trochanters (Bocak et al. 2019).
**CR3:** Reliably assigned to Lycidae, subfamily Dexorinae. Monophyly of Lycidae is well supported.
**CR4:** Locality and stratigraphy data provided in the original paper, double-checked for accuracy with The Paleobiology Database (https://paleobiodb.org/).
**CR5:** Age derived from The Paleobiology Database (https://paleobiodb.org/). The minimum age (98.17 Ma) is based on radiometric dating of the volcanoclastic matrix from the amber-bearing horizon at 98.79 ± 0.62 Mya (Shi et al. 2012), interpreted as a minimum age (Mao et al. 2018). The soft maximum (165 Mya) is based on the Daohugou biota, which lacks representatives of this family (Chen et al. 2004).
**Overlap note:** This fossil is also used in Cai et al. (2022) [Node 17, Osslimus (Elateridae) – Lycidae] with identical age constraints.
**Additional fossils:** Other species have also been recovered from Burmese amber, further supporting its use as a calibration point for this node (Tihelka et al. 2019; Kazantsev 2019).

**Fossil Calibration 13**
**Calibrated:** Crown Cantharidae
**Taxon:** *Molliberus albae*, Cantharidae
**Geological deposit/locality:** El Soplao amber deposit, Celis municipality, Cantabria, Spain (El Soplao Cave, Celis, Cantabria)
**Soft minimum age:** 105 Mya

**Soft maximum age:** 165 Mya
**Reference:** Peris, D., & Fanti, F. (2018). Molliberus albae gen. et sp. nov., the oldest Laurasian soldier beetle (Coleoptera: Cantharidae) from the Lower Cretaceous Spanish amber. Cretaceous Research, 91, 263-268.
**Justification of Parham et al. (2012) criteria:**
**CR1:** Single specimen (holotype CES-522, amber inclusion; laboratory of the El Soplao Cave, Celis-Rábago, Cantabria, Spain). Full museum data in the reference above.
**CR2:** Combination of diagnostic characters. *Molliberus albae* can be placed in the extant subfamily Cantharinae based on maxillary palpomeres different in length to the last securiform segment, long elytra, symmetrical last abdominal segments, prothorax without lateral processes, the presence of tibial spurs, and the fourth tarsomere undivided (Peris & Fanti 2018). It thus represents the earliest member of crown Cantharidae.
**CR3:** Reliably assigned to Cantharidae. The monophyly of Cantharidae is well supported.
**CR4:** Locality and stratigraphy data provided in the original paper, double-checked for accuracy with The Paleobiology Database (https://paleobiodb.org/).
**CR5:** Age derived from The Paleobiology Database (https://paleobiodb.org/). The minimum age (105 Mya) is based on the amber from El Soplao being early to middle Albian in age as determined by biostratigraphy of the Las Peñosas Formation (Najarro et al. 2009; Peñalver & Delclòs 2010). The soft maximum (165 Mya) is based on the Daohugou biota, which lacks representatives of this family (Chen et al. 2004).
**Overlap note:** This fossil is also used in Cai et al. (2022) [Node 19, Crown Cantharidae] with identical age constraints.
**Additional fossils:** Many species are described from Cretaceous Burmese amber.

**Fossil Calibration 14**
**Calibrated:** Split of Phengodidae – Rhagophthalmidae
**Taxon:** *Cretocydistus wittmeri*, Phengodidae
**Geological deposit/locality:** Burmese amber, Hukawng Valley, Kachin State, Myanmar; lowermost Cenomanian (ca. 98.79 ± 0.62 Mya)
**Soft minimum age:** 98.17 Mya
**Soft maximum age:** 165 Mya
**Reference:** Roza, A. S., Kusy, D., Lian, Z., & Kundrata, R. (2023). The first Phengodidae fossil (Coleoptera: Elateroidea): Cretocydistus wittmeri gen. et sp. nov. from the mid-Cretaceous Burmese amber. Palaeoentomology, 6.
**Justification of Parham et al. (2012) criteria:**
**CR1:** Single, perfectly visible specimen.
**CR2:** Combination of diagnostic characters reliably placing the fossil within Phengodidae.
**CR3:** Reliably assigned to Phengodidae. Phengodidae are monophyletic with early-branching Cydistinae and sister lineage to Rhagophthalmidae.
**CR4:** Locality and stratigraphy data provided in the original paper, double-checked for accuracy with The Paleobiology Database (https://paleobiodb.org/).
**CR5:** Age derived from The Paleobiology Database (https://paleobiodb.org/). The minimum age (98.17 Mya) is based on radiometric dating of the Burmese amber-bearing horizon at 98.79 ± 0.62 Mya (Shi et al. 2012). The soft maximum (165 Mya) is based on the Daohugou biota, which lacks representatives of this family.
